# Mapping AMPA and NMDA receptors at mouse brain hippocampal synapses by cryo-ET

**DOI:** 10.64898/2026.11.02.763697

**Authors:** Cathy J Spangler, Aya Matsui, Hoyoung Kim, Elizaveta V Shiltseva, Gunasekaran Dhandapani, Dennis Weingarten, Rachel Arnold, Yui Sugiyama, Skyler Jackman, Johannes Elferich, Eric Gouaux

## Abstract

AMPA and NMDA receptors are fundamental to synaptic transmission through the detection of released neurotransmitter. These ion channels open in response to glutamate binding, resulting in depolarization of the postsynaptic membrane and influx of calcium. Receptor localization plays a key role in synaptic strength, and changes in receptor number and location underpin synaptic plasticity, learning and memory. Here we apply receptor subunit-specific antibody fragments and multiplexed gold nanoparticle conjugates, together with cryo-electron tomography (cryo-ET), to elucidate the *in-situ* molecular organization of AMPA and NMDA receptors at hippocampal synapses in adult mouse brain tissue. AMPA receptors are predominantly found in densely packed liquid-like clusters, with interspersed NMDA receptors and adjacent NMDA receptor clusters. By direct localization of individual receptors, we provide molecular-resolution insight into the distribution of AMPA and NMDA receptors across the synapse, at peri and extrasynaptic sites, and within endocytosed vesicles. Visualization of multiple presynaptic vesicle fusion events suggests that receptors are near, but not always directly adjacent to, transmitter release sites. By mapping AMPA and NMDA receptors, and glutamate release sites, we provide receptor-resolution insight into the structure and function of brain hippocampal synapses.

## Main

Chemical synapses provide the linchpins of signaling between neurons in the brain, transforming the release of neurotransmitter from presynaptic boutons into electrical and chemical signals in opposing postsynaptic compartments^1^. Not only are the number and function of synapses central to the development and function of the nervous system, but the potentiation or depression of signaling at synapses underlies nearly all experience-based activities, including learning and memory^2^. On a molecular level, synaptic strength is modulated by the number, localization and conductance of synaptic receptors^3^. Neurological conditions, including autism and schizophrenia^4^, involve perturbation in synaptic number, while diseases, such as Alzheimer’s and NMDA receptor encephalitis, involve disruption in synaptic receptor function^5,6^.

Glutamatergic synapses, which mediate the majority of fast excitatory neurotransmission, harbor AMPA receptors (AMPARs) and NMDA receptors (NMDARs)^7^. Despite AMPA and NMDA receptors both being activated by glutamate, their relative affinities differ by nearly 3 orders of magnitude, with AMPARs and NMDARs activated by millimolar and micromolar concentrations of glutamate, respectively^7^. Moreover, NMDARs are blocked by magnesium in a voltage-dependent manner, thus requiring initial AMPAR activation and membrane depolarization to enable conductance of ions^8,9^. Influx of calcium through NMDARs activates multiple signaling pathways, one of which includes calcium–calmodulin (CaM)-dependent protein kinase II (CaMKII). Activated CaMKII promotes long-term potentiation and AMPAR insertion, whereas long-term depression involves AMPAR removal at synapses^10^. To provide a foundation upon which to understand the complex choreography of AMPAR and NMDAR signal transduction at hippocampal synapses, we sought to define the relationships between glutamate release sites and receptor localizations in three-dimensions, at a nanometer, receptor-level resolution.

Over the last several decades, our understanding of AMPAR and NMDAR localization at synapses has evolved dramatically from extensive studies using freeze fracture electron microscopy^11^, immunogold labeling electron microscopy^12^, super-resolution fluorescence imaging^13^, and cryo-ET without and with gold nanoparticle labeling^14–17^. These studies have provided progressively refined insight into the numbers of AMPARs and NMDARs at synapses, the presence of receptor clusters, and the locations of the receptors and clusters within the synaptic cleft. Indeed, prior studies have suggested that AMPARs in hippocampal synapses are preferentially localized on the periphery of the synapse, while NMDARs are more centrally positioned^14,18^. Additional elegant fluorescence and electron microscopy experiments suggest that sites of vesicular release and receptor detection are aligned with one another, in synaptic nanocolumns^19,20^. While previous studies have provided invaluable insight into vesicle and receptor localization, they are also limited by intrinsic resolution constraints, the use of large primary and secondary antibodies, or AuNPs bound to flexible linkers on the labeling reagent. Here we have brought together cryo-preservation and focused ion beam milling preparation of mouse brain hippocampal tissue, a suite of monomeric and dimeric receptor-specific Fab-AuNP conjugates, the implementation of an algorithm to improve the local alignment of the tilt series, and the analysis of nearly 300 tomograms, to precisely localize AMPARs and NMDARs in brain CA3-CA1 hippocampal synapses^21,22^. Together with the visualization of vesicular fusion events, we define the spatial relationship between receptor populations and sites of neurotransmitter release at unprecedented resolution.

## Results

### Cryo-ET of hippocampal synapses

We prepared 40 µm sections of the CA1 stratum radiatum region (**Fig. 1a**) labeled with Fab-AuNP conjugates to target subunits of AMPARs and NMDARs, and a competitive antagonist and a positive allosteric modulator to stabilize the receptor in a non-desensitized, Y-shaped conformation (**Extended Data Fig. 1**). The excised slice contains primarily CA3-CA1 Schaffer collateral synapses, though interneuron and other excitatory synaptic inputs to the CA1 region could also be sampled within our data. To ensure that synaptic function was retained in slices prepared under the buffer and cryoprotectant conditions used in the cryo-ET experiments, we carried out electrophysiology experiments and determined that the amplitude and frequency of spontaneous EPSCs (sEPSCs) in ACSF and HEPES solutions were not significantly different (**Fig. 1b-c**). Addition of dextran did not affect the frequency and amplitude of sEPSCs (**Supplementary Fig. 1**), while 5% w/v sucrose increased their frequency, but not amplitude (**Supplementary Fig. 1**). To minimize the exposure of the slices to sucrose, and thus retain endogenous synaptic activity, we carried out high pressure freezing (HPF) within 10 sec of applying the sucrose-containing cryo-protectant solution.

**Fig. 1:**
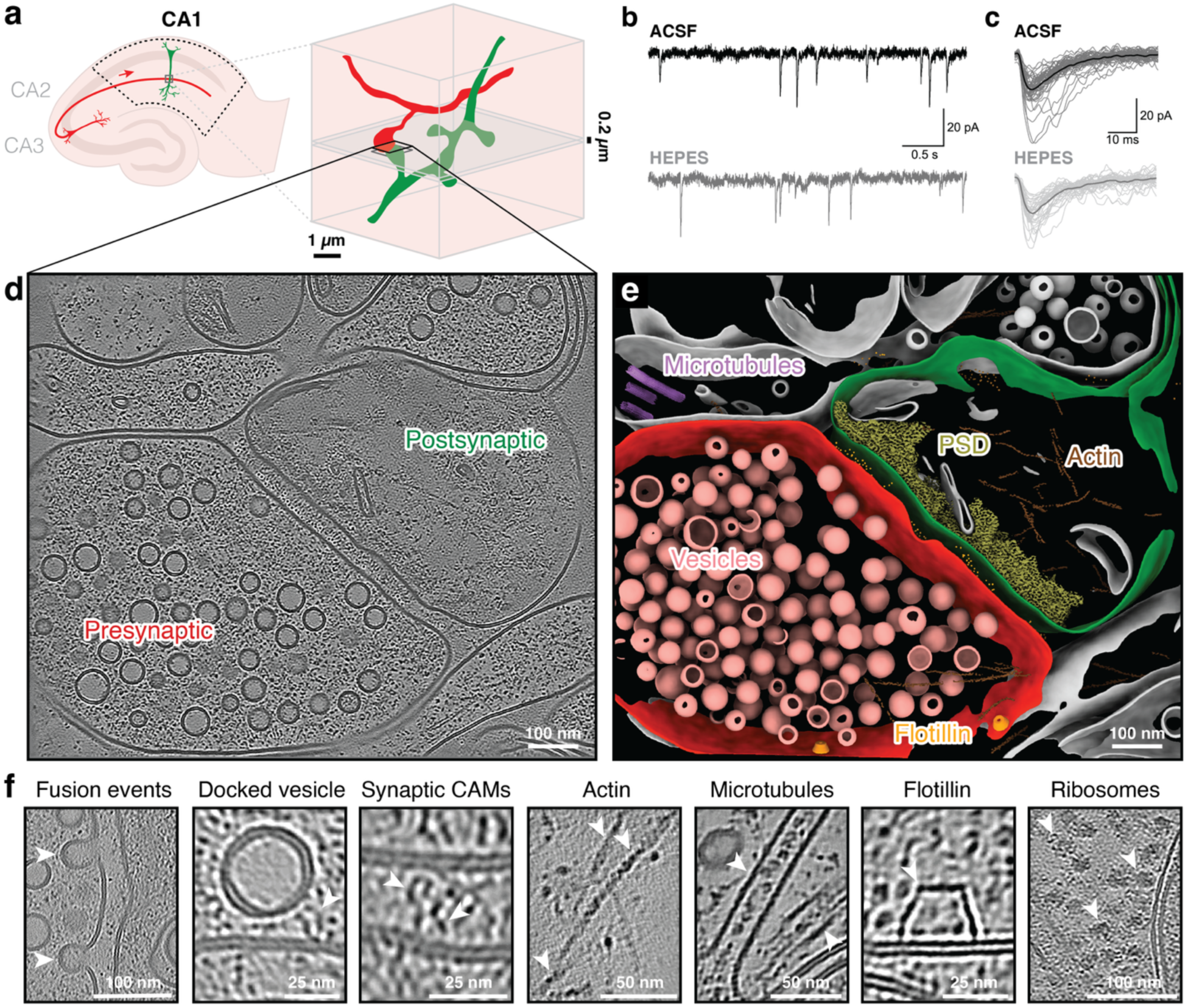
Cryo-ET of functional hippocampal slices reveals macromolecular assemblies at synapses. **a**, Hippocampal slice region excised for high pressure freezing (dotted black outline; presynaptic cell red; postsynaptic cell green; left), with a to-scale tomogram volume (black; 1 x 1 x 0.2 µm^3^; right). **b,** Recordings of spontaneous excitatory postsynaptic currents (sEPSCs) from CA1 neurons in hippocampal brain slices with ACSF or HEPES extracellular solutions. **c**, Traces of sEPSC events in each condition, with averaged traces shown in bold. **d**, Slice through denoised tomogram containing an excitatory hippocampal CA1 synapse. **e**, Annotated segmentation of tomogram shown in panel **d**. **f**, Examples of molecular features observed in hippocampal CA1 synapse tomograms (white arrows; CAMs = cell adhesion molecules).

We next prepared cryo-focused ion beam (cryo-FIB) milled lamella from the HPF tissue slices and collected tilt series for tomogram reconstruction at regions containing synaptic features, largely guided by vesicles within presynaptic compartments (**Extended Data Table 1**). A semi-automated pipeline for processing the tilt series included machine learning-based denoising, membrane segmentation, and an in-house developed tilt series alignment optimization that improved localization of AuNPs (**Extended Data Fig. 2a-b, Supplementary Fig. 2-3**). Ultimately, we obtained a dataset of 283 tomograms with well-aligned synaptic features (**Supplementary File 1**). Synapses in unlabeled and Fab-AuNP labeled samples had similar cleft widths, and we observe no indication that labeling perturbs or is hindered by local cleft geometry (**Supplementary Fig. 4**).

Presynaptic boutons harboring vesicles and postsynaptic compartments with heterogeneous PSD features bordered AuNP-labeled synaptic cleft regions (**Fig.1d-e, Supplementary Vid. 1**). Putative fusing and docked vesicles were observed adjacent to synaptic clefts, which were densely packed with protein-like features, some of which resemble transsynaptic cellular adhesion molecules (CAMs)^23^ (**Fig. 1f**). F-actin was frequently identified near the PSD in postsynaptic compartments (**Supplementary Fig. 5**), in agreement with the established role of actin at spine synapses and in PSD complexes^24^. The resolution obtained from cryo-FIB samples enabled direct visualization of microtubules filled with luminal particles at or near synapses^25^ and flotillin cages^26^, most often observed on presynaptic cellular membranes (**Supplementary Fig. 6**). Consistent with the occurrence of local protein synthesis at synapses^27^, we observed ribosomes in approximately a third of postsynaptic compartments. Most ribosomes were cytosolic rather than membrane-bound, and in selected instances were positioned closely to PSD scaffold components^28^ (**Supplementary Fig. 7**). We did not identify ribosomes within presynaptic regions.

### AMPA receptors populate densely packed clusters throughout the synaptic cleft

To enable localization of single receptors at synapses, we covalently conjugated an AuNP to the heavy chain C-terminus of a high affinity anti-GluA2 15F1 Fab (15F1 “C-term” Fab-AuNP) that binds to the R1 lobe of the amino terminal domain (ATD)^29^. The majority of hippocampal AMPARs contain two copies of GluA2 in the central “B/D” positions of the tetrameric core^30^, resulting in 2-fold symmetric, 2:1 15F1 Fab: receptor binding^31^ (**Fig. 2a, Extended Data Fig. 1a**). In tomograms labeled with the 15F1 Fab-AuNP, we observe the majority of AuNPs in the synaptic cleft region, located between the presynaptic and postsynaptic cleft membranes (**Fig. 2b, Supplementary Vid. 1**). From a set of 5,566 synaptic cleft-localized AuNPs across 71 synapses, we find that AuNPs are 8.8 ± 2.0 nm and 22.6 ± 2.1 from the center of the pre-and postsynaptic membranes, respectively (**Fig. 2c**). The distances between the AuNPs and the postsynaptic membrane agree with reconstructions of purified hippocampal AMPAR – 15F1 Fab-AuNP complexes, while the distances between the AuNPs and the presynaptic membrane show that there is sufficient space to accommodate the Fab-AuNP label^31^.

**Fig. 2:**
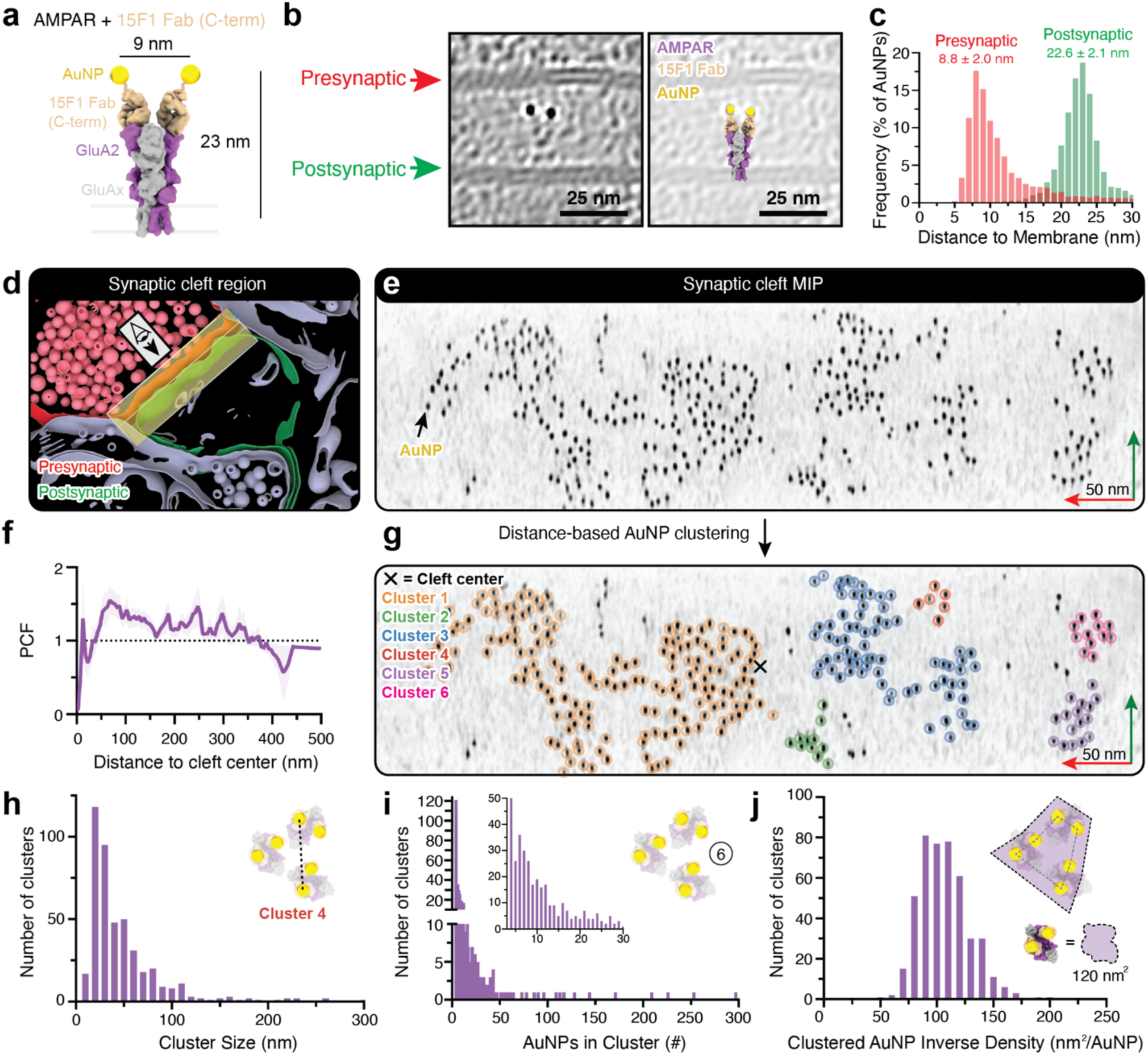
Most synaptic AMPA receptors are in densely packed clusters. **a**, Model of AMPAR bound to Fab-AuNP. **b**, Denoised tomogram slice through a synaptic cleft region containing a pair of AuNPs labeling an AMPAR (left), with overlaid Fab-AuNP: AMPAR model (right). **c**, Histogram of AuNP distances to the pre-and postsynaptic membranes in 15F1 “C-term” Fab-AuNP labeled synapses (n=5,566 AuNPs). **d**, Region included in synaptic cleft minimum intensity projection (MIP). **e**, MIP from the region boxed in **d**. **f**, Pair correlation function (PCF) between synaptic cleft center and AuNP positions in 15F1 “C-term” Fab-AuNP labeled synapses. The mean and SEM are shown (bin size 5; n=63 synaptic regions). **g**, Distance-based clustering (cutoff = 16 nm) of AuNP positions in cleft from **d**. **h**, Histogram of AuNP cluster sizes, estimated as the maximum AuNP-AuNP distance within a cluster (bin size 10). **i**, Histogram of the number of AuNPs per cluster (bin size 1), with inset highlighting the 4-30 AuNP range. **j**, Histogram of clustered AuNP inverse density (bin size 10). The estimated cross-sectional area of a single AMPA receptor is 120 nm^2^. In panels **h-i**, indicated measurements are illustrated on AuNP cluster 4 from panel **g**, for which putative receptor positions are shown.

Traditional AMPAR immunogold labeling methods require primary and secondary antibody binding, which are relatively large molecules relative to the spatial constraints within 20-30 nm wide synaptic clefts. To obviate the need for large labeling reagents, neuronal culture systems using affinity-tagged receptor subunits and AuNP-conjugated binders have been developed, achieving labeling that closely matched estimates derived from electrophysiological recordings of quantal events and single channel conductance^12,16^. In slices labeled with the 15F1 “C-term” Fab-AuNP conjugate, the median density of cleft-localized AuNPs was 2,236 AuNPs/µm^2^ and ranged from 290-7,000 AuNPs/µm^2^ (**Extended Data Fig. 1d**). As we expect this label to bind 2:1 to the majority of hippocampal AMPARs, the resulting receptor density is 145-3,500 AMPAR/µm^2^. Because most AMPARs at hippocampal excitatory synapses contain at least one GluA1 subunit^30^, we separately labeled slices with a GluA1-specific 11B8 “C-term” Fab-AuNP to have an additional probe of receptor localization. Using the 11B8 label, we observed a similar median and range of synaptic AuNP densities (1,974 and 708-5,997 AuNPs/µm^2^) as the GluA2-targeted label, along with AuNP-membrane distances consistent with structures of 11B8: AMPAR complexes^29,30^ (**Extended Data Fig. 1a-d, Supplementary Fig. 8**). These densities are in line with functional estimates of CA1 pyramidal synapse AMPAR densities^32–34^, indicating that remarkably high labeling efficiency is achieved with our Fab-AuNP conjugates. Because the binding mode of the 11B8 Fab to the GluA1 subunits in the A/C positions places the AuNPs farther from the central axis of the receptor, and because hippocampal AMPARs have a heterogeneous stoichiometry of GluA1 incorporation^30^, the 11B8 Fab-AuNP is not as faithful of a reporter of AMPAR localization as 15F1 Fab-AuNP conjugates. Therefore, we utilized the GluA2-targeted 15F1 label to localize AMPA receptors in the remaining experiments.

Prior studies have suggested that AMPARs in hippocampal synapses are preferentially localized on the periphery of the synapse^12,35^. To visualize the distribution of AMPARs within synaptic cleft regions in our tomograms (see Methods), we generated minimum intensity projections (MIPs) of the synaptic cleft oriented “face-on” to the postsynaptic membrane, in which the GluA2-targeted AuNP labels appear as ∼2 nm spots (**Fig. 2d-e**). Although our tomograms sample random slices through synapses, we expect that given the large number of tomograms we can identify clear cleft localization trends. Indeed, pairwise correlation and Ripley’s K-and H-function analyses (**Fig. 2f, Extended Data Fig. 1e, Supplementary Fig. 8c**) show no obvious relationship between the cleft center and AuNPs in GluA2 or GluA1 labeled synapses, implying that AMPARs at CA1 hippocampal synapses are equivalently distributed throughout the cleft and do not show a preferential localization to the center or the periphery.

Using a distance-based clustering algorithm, we assigned AuNPs as “clustered” or “unclustered”, enforcing that defined clusters contain a minimum of 4 AuNPs (**Fig. 2g**). Most AuNPs were located within irregularly shaped clusters of 20-100 nm in their largest dimension and containing 4-16 AuNPs (∼2-8 receptors), although some were substantially larger, with 100 or more AuNPs (**Fig. 2h-i, Supplementary Fig. 9**). Our estimates of cluster size are partially overlapping with those estimated by super-resolution fluorescence imaging of hippocampal synapses labeled with the same 15F1 antibody^36^. However, the spatial resolution of our labeling and imaging approach allows for high confidence quantitation of receptor counts and cluster areas, providing unparalleled insight into the extent of AMPAR colocalization. Within clusters, the density of AMPARs is approximately 3 to 6-fold higher (7,000-13,000 AuNPs/µm^2^) than the entire synaptic cleft (see Methods). This corresponds to a clustered AuNP inverse density of 60-170 nm^2^/AuNP, which relative to the 120 nm^2^ cross-sectional area of a single AMPAR indicates remarkably dense packing within clusters (**Fig. 2j**). The AMPAR packing, which is likely further increased by the presence of other synaptic proteins, directly restricts receptor diffusion and limits access to receptor conformations with large-scale splaying movements of the receptor ATD^37^. In addition, the close packing of receptors further underscores the importance of small, judiciously chosen labeling reagents.

### A minority of AMPARs reside outside of clusters or at extrasynaptic sites

The current model of AMPAR trafficking involves insertion of receptors or receptor subunits at extrasynaptic sites, followed by lateral diffusion of the receptors or subunits to the synapse, and receptor assembly and anchoring to scaffolding proteins within postsynaptic ‘slots’^38^. With our receptor labeling approach, we can determine whether a GluA2-containing receptor is within or outside of a cluster, and we can also exploit the distance between the AuNP and the postsynaptic membrane to infer the assembly and conformational state of the AMPARs. This is possible because prior high resolution x-ray and cryo-EM studies have shown that the competitive antagonist and allosteric modulator used in the slice preparation for this study stabilize AMPA receptors in a single Y-shaped conformation^39^. In fact, most receptor-bound AuNPs exhibited AuNP-postsynaptic membrane spacings consistent with the Y-shaped conformation^39^. We nevertheless identified a small number of AuNPs that were closer to the postsynaptic membrane than expected for a GluA2-containing AMPAR in a non-desensitized conformation (**Fig. 3a**).

**Fig. 3:**
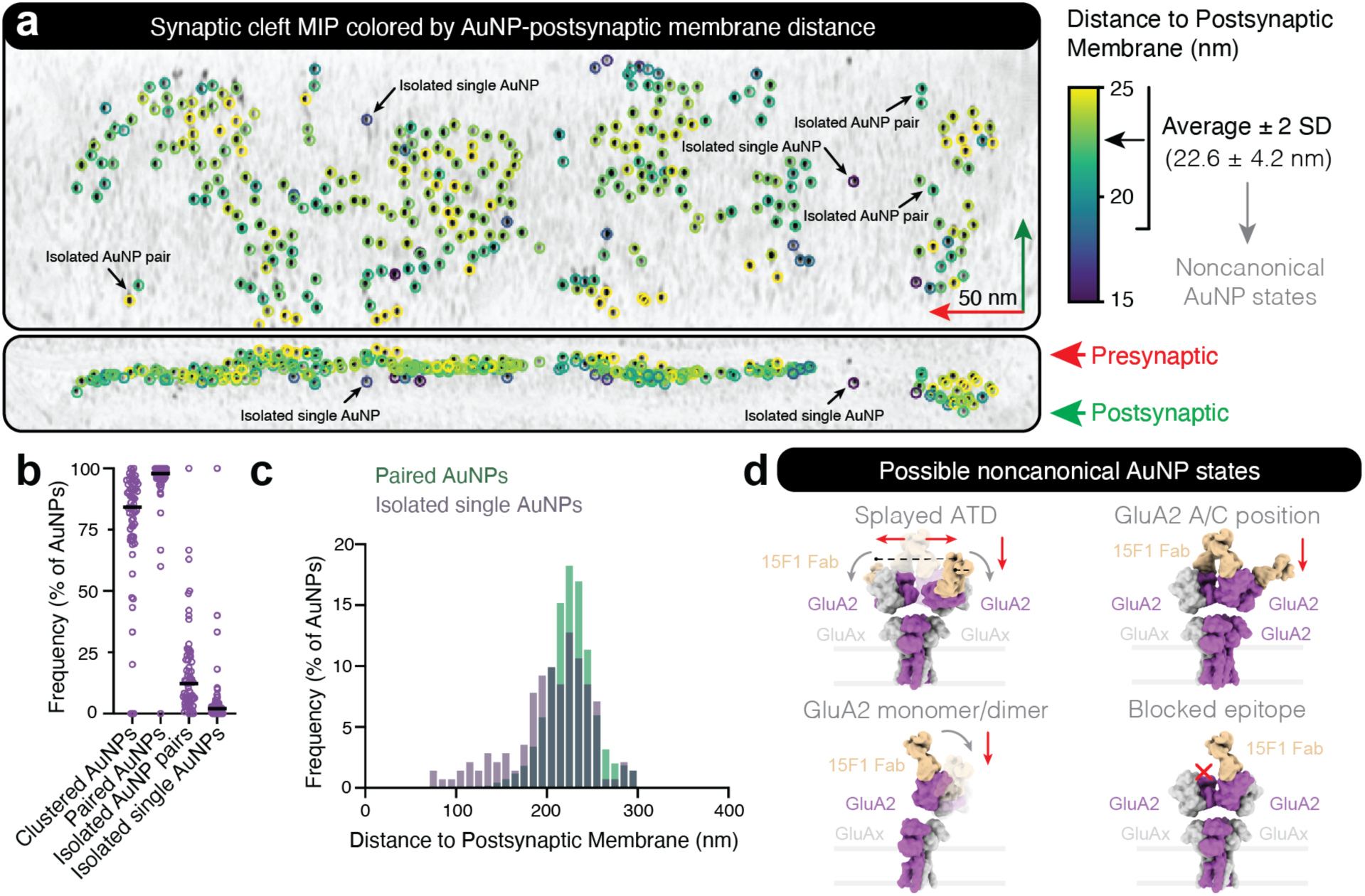
AuNPs pinpoint rare isolated receptors, receptor subunits, and non-resting state conformations. **a**, Postsynaptic membrane to AuNP distance-based coloring of AuNPs in synaptic cleft MIP viewed normal (top) and parallel (bottom) to the postsynaptic membrane. **b**, Median frequency of AuNP states in anti-GluA2 15F1 “C-term” Fab-AuNP labeled synapses (n=63 synaptic regions; see Methods). **c**, Frequency histogram (bin size 10) of postsynaptic membrane to AuNP distances for paired (n=5,417) and isolated single (n=149) AuNPs. **d**, Possible receptor-bound noncanonical AuNP states represented by smaller postsynaptic membrane to AuNP distances or isolated AuNPs.

We therefore asked whether the AuNPs with smaller than average postsynaptic membrane distances participated in expected receptor labeling pairs within clusters or represented isolated or unpaired AuNPs. On average, 84% of all AuNPs were found in clusters and 98% of those were in putative receptor pairs, 12% were in isolated AuNP pairs, and only 2% were clearly isolated single AuNPs (**Fig. 3b**). Interestingly, the distribution of AuNP-postsynaptic membrane distances for the isolated single AuNPs included a higher relative proportion of abnormally small distances than paired AuNPs (**Fig. 3c**). Manual inspection of these “low” AuNP cases confirmed that some were isolated single AuNPs, while others were contained within AuNP clusters (**Fig. 3a**). We speculate that this population could represent receptors with “splayed” ATDs^40^, GluA2 subunits in the “A/C” positions that orient the Fab-AuNP closer to the membrane^29^, isolated receptor subunits, or partially assembled receptors^41^(**Fig. 3d**). We also note that isolated single AuNPs with canonical AuNP-membrane spacing may represent incompletely labeled receptors due to Fab epitope accessibility, unbinding, or incomplete initial labeling. Nevertheless, our data demonstrates that the overwhelming majority of synaptic AMPARs are clustered assembled receptors in canonical resting states, with only a small minority of AuNPs that may represent unclustered assembled receptors or receptor subunits.

We captured several synapses with AuNP labels in positions consistent with peri-or extrasynaptic receptor localization (see Methods), at both head and neck regions of dendritic spines (**Fig. 4a-b, Extended Data Fig. 3a, Supplementary Vid. 1**). In these tomograms, the non-synaptic AuNPs consisted of 5-25% of the total AuNP population, adopted pairwise AuNP-AuNP distances similar to that seen for synaptic receptors, and resided at distances from the postsynaptic membrane consistent with resting, Y-shaped receptor states (**Fig. 4c-e**). We also observed occasional paired AuNPs near undefined membranes, potentially belonging to postsynaptic compartments from synapses outside the field of view, dendritic shafts, or even non-neuronal cells. We discovered several examples of AuNP clusters on internalizing membrane regions (**Fig. 4f**) or within intracellular postsynaptic membrane compartments (**Fig. 4g-h**). Because the 15F1 Fab-AuNP was applied to tissue, and the label does not have a facile mechanism to gain cell entry, we expect that the AuNPs observed within these postsynaptic compartments are the consequence of receptor internalization. The groups of internalized labels contain between 14-41 AuNPs, showing that AMPAR trafficking can occur in groups of at least 7-20 receptors. Together, these snapshots suggest that AMPARs largely exist in clusters within peri-and extra-synaptic regions, and that receptor recycling involves internalization of receptor clusters.

**Fig. 4:**
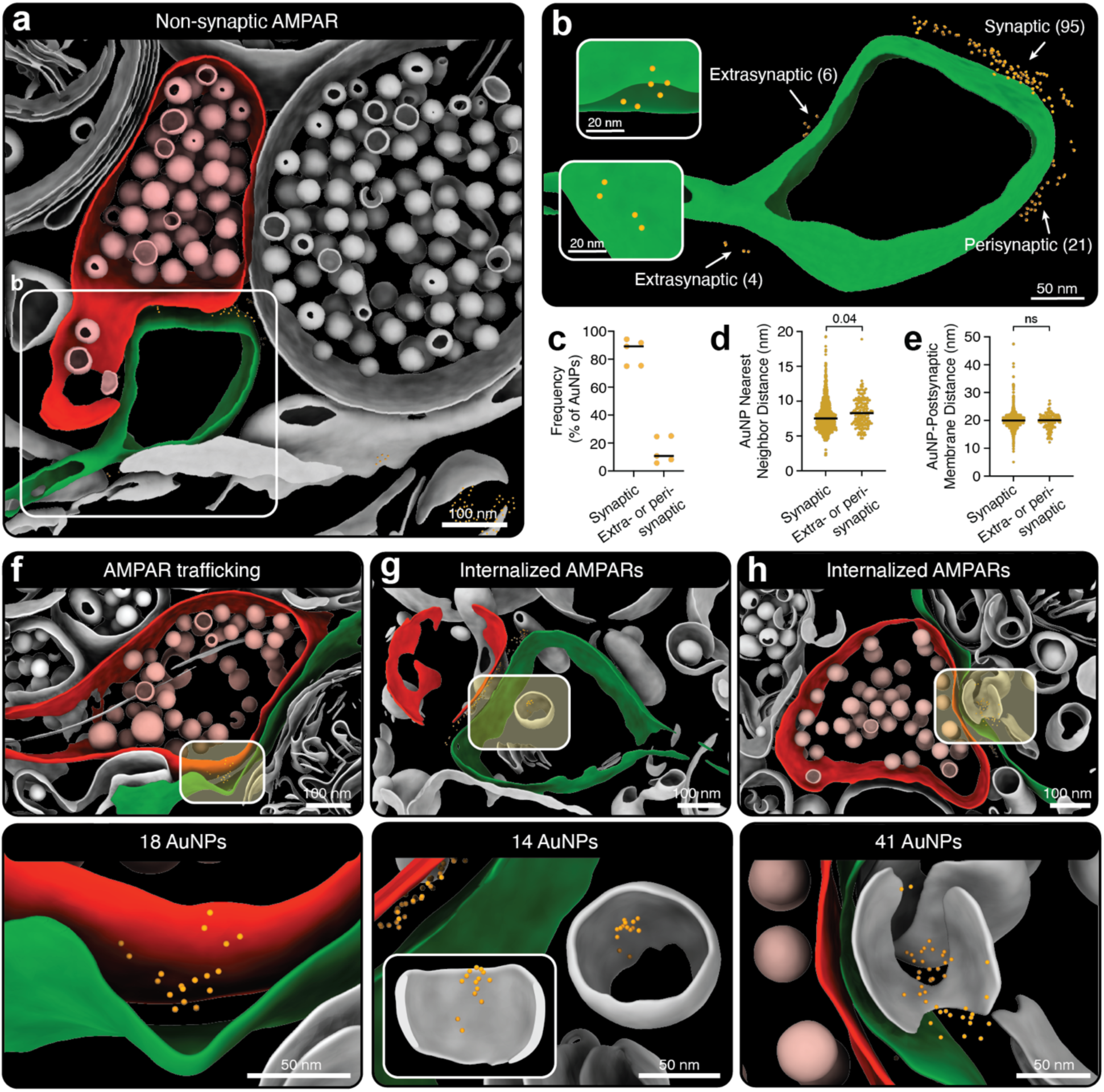
Most non-synaptic AMPA receptors are in canonical resting states within clusters. **a**, Segmentation of anti-GluA2 15F1 “C-term” Fab-AuNP labeled synaptic tomogram. **b**, Isolated segmentation of postsynaptic membrane from panel **a** with indicated AuNP localizations. The number of AuNPs in each region is denoted and views normal to the membrane at extrasynaptic receptor patches are shown. **c**, Median frequency of AuNPs in synaptic versus extra-or perisynaptic regions in tomogram subset containing resolvable non-synaptic AuNPs (n=5 tomograms). **d**, Median AuNP to AuNP nearest neighbor distance for synaptic and extra-or perisynaptic AuNPs (n=976 synaptic, 165 extra-or perisynaptic AuNPs). **e**, Median AuNP to postsynaptic membrane distance for synaptic and extra-or perisynaptic AuNPs. **f**, Membrane invagination with adjacent AuNP labels, indicative of putative AMPA receptor trafficking. **g-h**, Intracellular compartment-localized AuNPs indicative of internalized AMPA receptors.

### AMPA receptors within clusters are closely packed in a liquid-like state

Organization of AMPARs into synaptic clusters is supported by multiple lines of evidence, yet much less is known about the molecular mechanisms orchestrating cluster formation. One potential mechanism is the interaction of AMPARs with PSD95, the most abundant PSD scaffold protein at glutamatergic synapses^42^, via auxiliary TARP subunits in native AMPAR complexes. But other mechanisms exist, including crosslinking by extracellular scaffolds such as neuronal pentraxins and noelins^43,44^. In addition, the high density of AMPARs observed in our data suggest that AMPARs may directly interact with each other within clusters, further raising the question of what interactions and mechanisms drive cluster formation and whether the clustered receptors adopt ordered arrays.

Flexible attachment of the AuNPs, as well as overlapping intra-and inter-receptor AuNP spacings, made it impossible to unambiguously assign AuNP pairs to receptor positions in dense clusters labeled with the 15F1 “C-term” Fab-AuNP conjugate. We therefore developed a rigidly linked conjugate in which the AuNP is divalently coupled to 2 cysteine residues on a constant domain α-helix^31^. This 15F1 “H4K-2Cys” Fab-AuNP conjugate yields a well-defined 7 nm intra-receptor AuNP pair spacing, thus allowing us to define discrete receptors with confidence (**Fig. 5a-b, Extended Data Fig. 1a, Supplementary Vid. 2**). A dataset collected using this labeling reagent resulted in 55 tomograms in which we could assign 1,254 AuNP pairs to receptor positions (**Fig. 5c-d, Extended Data Table 2, Extended Data Fig. 4a-c**). We performed subtomogram averaging (STA) of these positions, which confirmed density corresponding to an AMPAR bound to two 15F1 “H4K-2Cys” Fab-AuNP conjugates (**Fig. 5e, Extended Data Fig. 4d-g**). Subsequent STA and AuNP-guided docking of a native GluA2/GluA1 di-heteromeric AMPAR resting-state receptor structure (PDB ID 7LDD) into tomograms revealed examples of inter-receptor interfaces between GluA2-GluA2, GluAx-GluAx, and GluA2-GluAx subunits, where GluAx corresponds to GluA1 or GluA3 (**Fig. 5f**). These included putative ATD-ATD and LBD-LBD interactions, as well as juxtaposition of transmembrane auxiliary subunits. These varied ensembles of subunit-subunit contacts suggest that there are not any specific receptor-receptor interactions that predominate in receptor clusters.

**Fig. 5:**
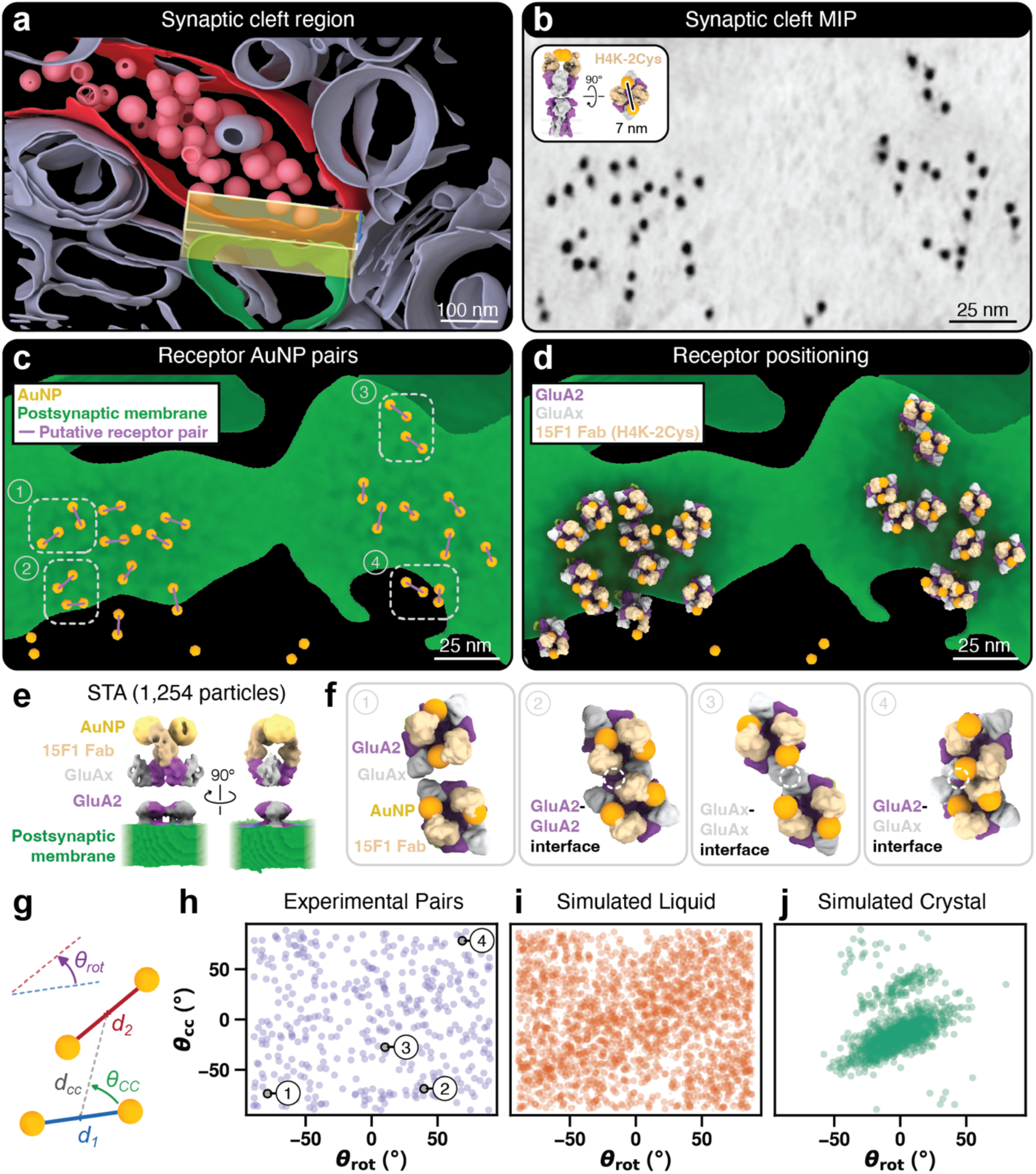
AMPA receptors cluster in non-regular assemblies. **a**, Segmentation of a 15F1 “H4K-2Cys” Fab-AuNP labeled tomogram. **b**, MIP from the region boxed in panel **a**. Inset shows a model of AMPAR bound to rigidly conjugated 15F1 “H4K-2Cys” Fab-AuNP. **c**, View normal to cleft postsynaptic membrane with lines indicating assigned intra-receptor AuNP pairs. **d**, AuNP and membrane-guided assignment of AMPA receptor poses (see Methods). **e,** Composite subtomogram average (STA) map of 15F1 “H4K-2Cys” Fab-AuNP: AMPAR complex. **f**, Examples of adjacent AMPA receptors demonstrate diversity of possible receptor-receptor interfaces. **g**, Definitions for pairwise packing analyses of neighboring AMPA receptors (see Methods). Scatter plots of θ_rot_ vs. θ_cc_ for (**h**) experimental pose (points 1-4 are from examples noted in panels **c,f**), (**i**) simulated liquid, or (**j**) crystalline receptor pairs.

To determine whether specific receptor-receptor species were preferentially enriched within clusters, we systematically investigated the relative orientation of AuNP pairs, as proxies for AMPARs, which we described by two angles: θ_rot_, indicating whether pairs are rotated relative to another, and θ_cc_, indicating the relative orientation of pair centers (**Fig. 5g**). Both angles appeared random and independent of each other, similar to a simulation of a liquid-like arrangement of receptors (**Fig. 5h-i**). Simulation of AMPARs in a 2D-crystalline lattice with defined interactions, on the other hand, resulted in a prevalence of a specific combination of θ_rot_ and θ_cc_ angles (**Fig. 5j**). To exclude bias arising from our assignment of AuNPs into receptor pairs and subsequent STA, we performed a graph-analysis of all combinations of 4 AuNPs, which also showed consistence of our data with a liquid-like model (**Supplementary Fig. 10**). In agreement with this analysis, STA did not reveal clear receptor-associated densities other than those consistent with known AMPAR auxiliary subunits (**Fig. 5e, Extended Data Fig. 4f**). We therefore conclude that the clustering of AMPARs is dominated by conformationally mobile scaffolding proteins, which is further supported by our observation of heterogeneous protein-like features in the PSD adjacent to receptors within tomograms (**Supplementary Fig. 5a**).

### AMPA and NMDA receptors intermingle

Activation of AMPARs at glutamatergic synapses drives rapid depolarization of the postsynaptic membrane potential that, in turn, activates NMDARs^8,9,45^. Similar to AMPARs, NMDARs cluster on dendritic spines, with prior studies suggesting they occupy spatially distinct domains more central within the PSD that are separated by 50-250 nm from AMPAR clusters^12,46^. Using our Fab-AuNP labeling platform, we set out to define this co-organization of AMPARs and NMDARs at synapses with single-receptor resolution. Electrophysiological recordings of hippocampal slices in the ACSF or HEPES slice incubation and cryoprotectant solutions exhibited canonical AMPAR and NMDAR currents, as well as AMPAR/NMDAR ratios^32,47^ (**Fig. 6a-b, Supplementary Fig. 1**), thus demonstrating physiologically representative synaptic receptor activity.

**Fig. 6:**
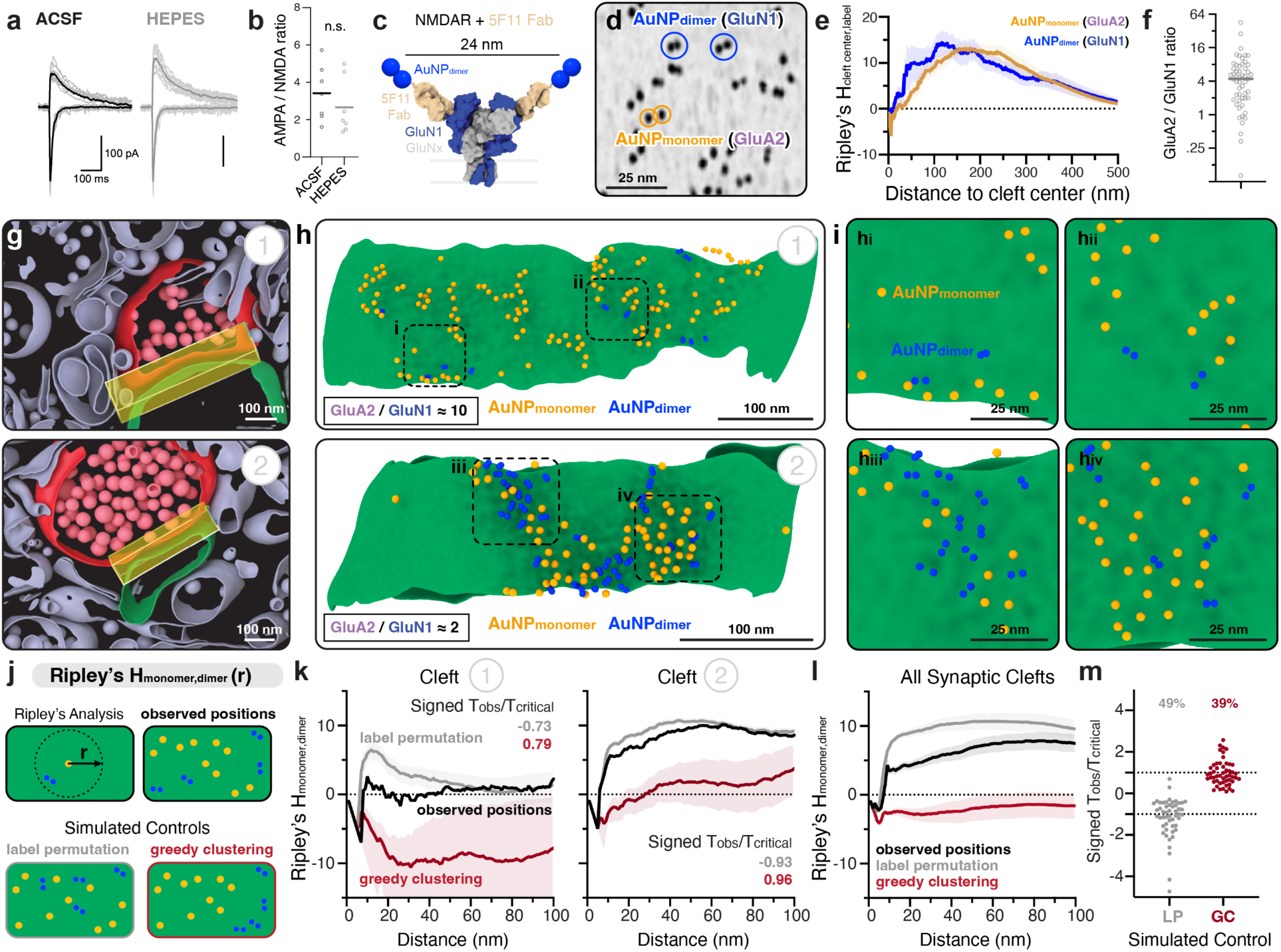
AMPA and NMDA receptor clusters intermingle. **a**, Recordings of evoked AMPA and NMDA receptor currents from neurons in CA1 hippocampal brain slices. **b**, AMPA/NMDA ratios from recordings in panel **a**. **c**, Model of NMDAR bound to 5F11 “C-term” Fab-AuNP_dimer_. **d**, MIP of synaptic region labeled with anti-GluA2 15F1 Fab-AuNP_monomer_ and anti-GluN1 5F11 Fab-AuNP_dimer_. **e**, Bivariate Ripley’s H_cleft center, label_ functions between the synaptic cleft center and AuNP_monomer_ or AuNP_dimer_ positions (n=53 synaptic regions; see Methods). Mean and standard error of the mean (SEM) are shown. **f**, Median GluA2/GluN1 ratio derived from AuNP_monomer_ and AuNP_dimer_ counts (n=53 synaptic regions). **g**, Segmentations of dual labeled tomograms. **h**, Views normal to postsynaptic membranes in **g**, with annotated AuNP_monomer_ and AuNP_dimer_ positions. AuNPs are represented at ∼2x scale for visibility. **i**, Views of boxed regions in panel **h**. **j**, Bivariate Ripley’s H_monomer, dimer_ functions were calculated for observed label positions and two simulated null models: label permutation and greedy clustering (see Methods). **k**, Ripley’s H_monomer, dimer_ for observed, label permutation, and greedy clustering positions at indicated synaptic regions in panel **h**. The mean and standard deviation are shown. **l**, Ripley’s H_monomer, dimer_ mean and SEM from all synaptic regions pooled (n=53 synaptic regions). **m**, Maximum absolute difference test T_observed_/T_critical_ values, multiplied by the direction of change, for each synaptic region compared to label permutation (LP) or greedy clustering (GC) simulated models (see Methods). The percentage of synaptic regions for which the null hypothesis was rejected (outside of-1 to 1 range) is reported.

To validate our strategy to specifically label NMDARs, we harnessed the high affinity, highly specific 5F11 Fab that labels the two obligatory GluN1 subunits in the distal “A/C” positions of the receptor^48^ (**Extended Data Fig. 1a, Extended Data Fig. 5a**). A dataset on hippocampal slices solely labeled with an engineered 5F11 “C-term” Fab-AuNP conjugate contained a median synaptic AuNP density of 2,018 AuNPs/µm^2^ (ranging from 105-6,568 AuNPs/µm^2^) and AuNP-postsynaptic membrane distances consistent with cleft widths and structures of 5F11 Fab: NMDAR complexes, which result in a ∼5 nm “lower” average AuNP-postsynaptic membrane spacing than 15F1 Fab C-term: AMPAR complexes^48^ (**Extended Data Fig. 1c-d, Extended Data Fig. 5b**). In contrast to previous reports of NMDAR enrichment at the center of synapses, we found that GluN1-targeted AuNPs were found throughout imaged cleft regions, with only a modest, statistically insignificant trend towards more central localization than GluA2-targeted AuNPs (**Extended Data Fig.1e, Extended Data Fig. 5c, Supplementary Table 1**).

Clusters of NMDAR-targeted AuNPs were larger and more spread out in comparison to the AMPAR-AuNP clusters, in accord with the larger spacing (∼24 nm) between AuNPs in the 5F11 Fab-AuNP: NMDAR complex^48^. Distance-based analysis of the GluN1-targeted AuNPs estimated cluster sizes at 25-200 nm with 4-16 AuNPs (∼2-8 receptors) (**Extended Data Fig. 5d-e**). The inverse density of AuNPs within these clusters was 100-300 nm^2^/AuNP (**Extended Data Fig. 5f**), approximately twice that observed for GluA2 clusters. However, we expect that the wider intra-receptor AuNP spacing results in effectively larger clusters and a concomitant underestimation of receptor density. We observed many identifiable examples of 5F11 Fab-AuNPs with pair spacings consistent with the 5F11 Fab: NMDAR structure, in both isolated and clustered pairs (**Extended Data Fig. 5g**). Only at one synapse did we observe AuNP pairs on the extracellular surface of a presynaptic compartment, indicating two individual NMDARs (**Extended Data Fig. 5g**). This demonstrates that in the hippocampus of adult mice, presynaptic NMDARs, if present, are rare^49^. Nevertheless, a minority of cleft-localized AuNPs could also represent presynaptic NMDAR localization, which cannot be unequivocally distinguished based on AuNP-membrane spacing.

To simultaneously localize AMPARs and NMDARs, we co-labeled tissue with the GluA2-and GluN1-specific Fab-AuNP conjugates, aiming to differentiate AuNP-receptor pairings by label-specific AuNP-postsynaptic membrane distances. The dual labeled samples had an increased median AuNP density in the synaptic cleft (4,317 AuNPs/µm^2^) and a bimodal distribution of AuNP-postsynaptic membrane distances with peaks matching those from equivalent independently labeled samples (**Extended Data Fig. 1c-d**). However, the extent of overlap in these distributions precluded confident classification of individual AuNPs as either GluA2 or GluN1-bound labels. To overcome this, we utilized a dimeric AuNP label that can be distinguished from AuNP monomers within tomograms^22^ (**Supplementary Fig. 11**) and prepared a 5F11 Fab-AuNP dimer conjugate (**Fig.6c-d, Extended Data Fig. 1b**). The dual labeled tomograms showed specific and distinguishable AuNP monomer and dimer labeling (**Extended Data Fig. 6a-b, Supplementary Vid. 3)**, with AuNP nearest neighbor distances, including those between individual dimer AuNPs, similar to those observed in purified monomer or dimer Fab-AuNP samples. Most importantly, the AuNP-postsynaptic membrane distances for assigned AuNP monomer and dimer picks matched those from the individually labeled tissue samples (**Supplementary Fig. 11**).

The number, proportion, and localization of AMPARs and NMDARs influence the magnitude of the membrane depolarization, ion influx, and activation of downstream signaling pathways that regulate receptor expression and trafficking, contributing to plasticity at glutamatergic synapses^50^. In our dual labeled data, bivariate Ripley’s spatial analysis indicated a subtle, but statistically insignificant, shift of the AuNP dimer distribution towards the center of imaged cleft regions compared to monomers **(Fig. 6e, Supplementary Table 1**). The median ratio of monomer to dimer AuNPs, and by extension the ratio of AMPARs to NMDARs, was 4.4 and ranged from 0.1-45.3 (**Fig. 6f**), remarkably consistent with electrophysiology-derived receptor ratios^32,47,51^. Unexpectedly, most dual-labeled synapses contained monomer and dimer AuNPs that were closely intermingled, thus demonstrating colocalization of AMPARs and NMDARs (**Fig. 6g-i)**. At the same time, we also observed many cases of partially exclusive clustering, where one receptor type largely predominated within a particular synaptic region (**Fig. 6g-i**). We hypothesize that the intermingling of AMPARs and NMDARs facilitates NMDAR-dependent modulation of AMPAR activity by, for example, the phosphorylation of AMPAR GluA1 subunits by CaMKII^52^. We also observed several examples of internalized receptor clusters containing both monomers and dimers, showing that AMPARs and NMDARs can traffic together (**Extended Data Fig. 3b)**. We used bivariate Ripley’s K-and H-function analyses to compare observed AuNP monomer and dimer positions to simulated sets of “label permutated” or “greedy clustered” positions in which monomer or dimer labels were reassigned randomly or by preferential self-clustering, respectively (**Fig. 6j, Extended Data Fig. 7a-c**). The mean Ripley’s function for observed positions lies in between the two modeled extremes, but closer to the label permutation model (**Fig. 6k-m, Extended Data Fig. 7d-e**). On the one hand, our analysis shows that AMPARs and NMDARs are substantially intermingled, yet on the other hand it also shows that receptors can preferentially self-associate.

### AMPA and NMDA receptors are heterogeneously positioned relative to fusion events

Previous studies on AMPARs and NMDARs have detected alignment of receptor clusters with presynaptic vesicle fusion machinery within 50-80 nm diameter nanodomains^12,36,53^. However, the extent of alignment varies by synapse, receptor subtype, and release site, and it remains unclear precisely how such an alignment is choreographed at the molecular scale. Our tomograms contain numerous examples of presynaptic vesicles, including “docked” subsets with tether-like densities connecting to presynaptic or other vesicle membranes, examples with dark intravesicular densities and larger diameters reminiscent of dense core vesicles, and a few containing unidentified rod-shape densities (**Extended Data Fig. 8c-f**). In about 15% of imaged synaptic regions, we observed putative fusion events in the vicinity of AuNP-labeled receptors in the cleft, with diameters expected for fusing presynaptic vesicles and without identifiable clathrin coats indicative of endocytic pits (**Fig. 7a-b, Extended Data Fig. 8a-b, Supplementary Vid. 4**). We expect that these events represent spontaneous fusion or hypertonic sucrose solution-triggered asynchronous release events, both of which come from the same vesicle pool as action potential-evoked fusion events^54,55^. We took advantage of these fortuitously captured events to investigate the localization of receptors relative to fusion sites, leveraging the sub-nanometer precision provided by the Fab-AuNP labels.

**Fig. 7:**
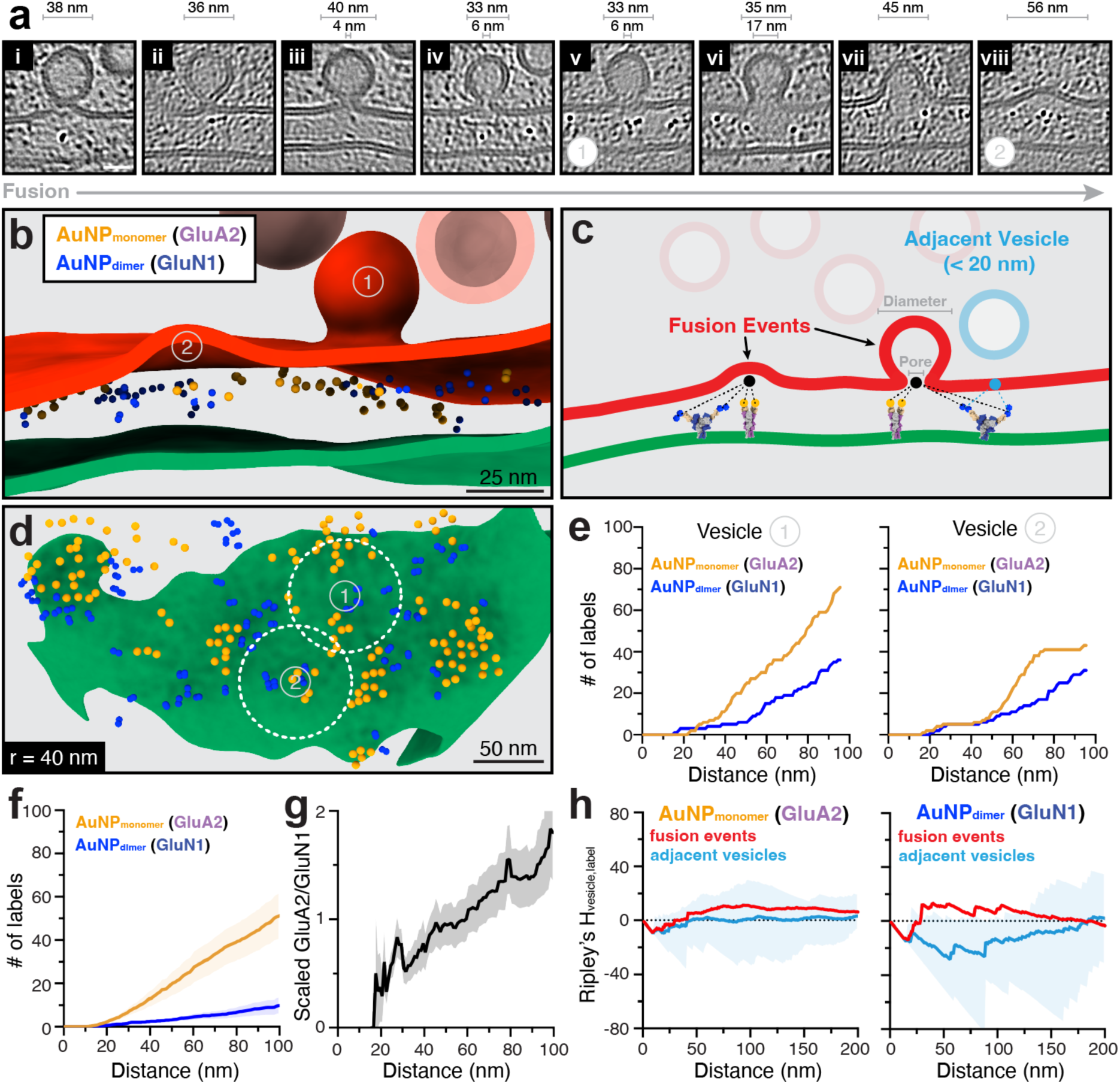
Fusion events have variable alignment with receptor clusters. **a**, Denoised tomogram slices with fusion events at synaptic clefts in AMPAR and NMDAR dual labeled hippocampal slices. Fusing vesicle and pore diameter, when present, are denoted. Scale bar is 20 nm. **b**, Segmentation of dual labeled synaptic tomogram with two fusion events (v and viii from panel **a**). **c**, Illustration of fusion event and adjacent vesicle definitions and fusion site-to-AuNP label measurements. **d**, View normal to postsynaptic membrane from panel **b**, with fusion event locations overlaid. AuNPs are shown at ∼2x scale for visibility. **e**, Cumulative frequency plots showing the number of AuNP labels within a given distance from each fusion event in panel **d**. **f**, Cumulative label number as a function of distance to fusion sites for all fusion events in dual labeled dataset (n=11 fusion events). **g**, GluA2/GluN1 ratio from data in panel **f**, scaled to the overall ratio for each synaptic region. Mean and standard error of the mean per fusion event shown for panels **f**, **g**. **h**, Bivariate Ripley’s H_vesicle, label_ between GluA2 (left) or GluN1 (right)-specific AuNP labels and fusion event (n=13) or adjacent vesicle (n=54) sites on presynaptic membrane. Mean and 95% envelope are shown.

Within anti-GluA2 15F1 Fab-AuNP labeled samples, an average of 42 AuNPs (∼21 receptors) were within a 100 nm radius of fusion events, ranging from 8-73 AuNPs across 23 observed events (**Fig. 7c, Extended Data Fig. 9a-f**). A bivariate Ripley’s analysis indicated a weak statistical association between AuNPs and fusion event sites that were not observed with non-fusing presynaptic membrane-adjacent vesicle sites (**Extended Data Fig. 9g-h**). To assess the spatial dependence of these results, we shifted fusion sites 40 nm along the presynaptic membrane from their original location or randomly reassigned fusion and AuNP labels within the combined point set. On average, the Ripley’s function for observed fusion sites was similar to that for the simulated 40 nm shifted sites. For the label permutated sites, the observed function was lower at distances below 50 nm and higher at distances above 50 nm. We expect that this is due to the label permutation simulation most often assigning fusion sites to an observed AuNP position within a receptor cluster, resulting in consistently higher local alignment with AuNPs than the observed fusion events. However, in all cases, differences between the observed and simulated Ripley’s were not statistically significant, likely in part due to the high variability observed between synapses (**Extended Data Fig. 9g-h, Supplementary Table 1**). Our data therefore suggests only a small enrichment of GluA2-containing AMPARs at the observed fusion event sites. Nevertheless, the juxtaposition of fusion event sites and AMPAR clusters are sufficiently close so as to enable activation of a physiologically relevant number of AMPARs by single vesicle release events^32,47^.

Extending these fusion event analyses to GluA2 and GluN1 dual labeled samples, which together contained 13 fusion events, we observed an average of 44 monomer and 10 dimer AuNP labels within a 100 nm radius of fusion sites, corresponding to approximately 22 AMPARs and 5 NMDARs (**Fig. 7d-f, Extended Data Fig. 10a**). These numbers are in line with electrophysiological estimates of receptor activation^32,47^, implying that receptors are activated within ∼100 nm of fusion sites, on average. Interestingly, the ratio of GluA2 to GluN1, scaled to the overall ratio at each synaptic region, increased with distance from fusion events, suggesting closer alignment of GluN1 with fusion events than GluA2 (**Fig. 7g**). In agreement with this observation, bivariate Ripley’s analyses of fusion sites and monomer or dimer AuNP positions shows a larger, though still statistically insignificant, difference between observed fusion sites and adjacent vesicle sites, 40 nm shifted or label permutated controls for dimers than is observed for monomers (**Fig. 7h, Extended Data Fig. 10b-c, Supplementary Table 1**). Thus NMDARs show a more modest preferential alignment with the observed fusion events than AMPARs, which again raises the possibility that these observed release events represent asynchronous fusion^12^.

## Discussion

Here we used engineered Fab – AuNP complexes to localize AMPARs with single receptor resolution, and exploit newly developed Fab – dimeric AuNP labels to enable the colocalization of AMPARs and NMDARs, thus extending our understanding of synaptic receptor organization from synaptic preparations and primary neuronal cultures^14–16,56^ to *bona fide* brain hippocampal synapses. The AMPAR clusters are variable in size, from groups of two receptors to ensembles of one hundred receptors, generally organized in densely packed groups that preclude an abundance of additional proteins within the interior of the clusters. We find no evidence of crystalline-like order within the clusters, with receptor organization best described as ‘liquid-like’ and thus consistent with prior experiments showing no preponderance for higher order assembly of receptor extracellular domains^57^. Together with the large distribution of cluster sizes and the absence of repetitive and specific receptor-receptor interactions, these synaptic receptor clusters appear distinct from higher-order transient structures that have been observed with other membrane protein model systems^58^. As previously described, cluster formation is likely driven, at least in part, by postsynaptic scaffolding proteins^59^. The functional consequences of the dense packing of receptors, especially in the large clusters, is manifold and includes hinderance of diffusional escape of the interior receptors, restriction of large conformational rearrangements of the receptors’ extracellular domains, and possible slowing of transmitter diffusion.

To gain a more complete understanding of receptor organization at synapses, we harnessed the Fab – monomeric AuNPs together with our recently developed Fab – dimeric AuNP conjugates to simultaneously localize AMPARs and NMDARs, discovering that these two classes of receptors are interwoven in clusters across the synaptic cleft. This result is in contrast to previous models of receptor localization by super-resolution fluorescence imaging of primary neuronal cultures that suggested their spatially distinct localization^46^. Although the immediate colocalization of AMPARs and NMDARs is not a prerequisite for the membrane depolarization relief of NMDAR magnesium block, it may nevertheless play important roles in binding partner and phosphorylation events that take place between AMPARs and NMDARs within the postsynaptic density, thus facilitating receptor ‘cross talk’. The most parsimonious explanation of AMPAR and NMDAR colocalization involves their coupling via PSD-95, perhaps in addition to extracellular scaffolding proteins^43^. Nevertheless, it is also possible that additional layers of postsynaptic proteins participate in the AMPAR and NMDAR colocalization. In satisfying accord with electrophysiological estimates of receptor number^32–34,47^, we observe an average of 20-25 AMPARs and 5 NMDARs within 100 nm of vesicle fusion events. These putative asynchronous release sites are occasionally, but not consistently, aligned with receptor clusters, and spatial analyses indicate greater alignment of these sites with NMDARs than AMPARs. Indeed, a prior study in which AMPARs and NMDARs were separately localized via electron microscopy suggested that AMPARs and NMDARs are preferentially aligned with synchronous and asynchronous vesicle release sites, respectively, in line with their sequential co-dependent activation^12^.

Our work has established the receptor-level organization of AMPARs and NMDARs at hippocampal synapses, together with the juxtaposition of vesicle fusion events. These studies have shed new light on the architecture of glutamatergic synapses, yet we acknowledge several limitations. Because our lamella sample 150-200 nm thick synaptic subregions, they typically represent between 50-80% of an average CA1 synaptic surface area in the stratum radiatum^60^. Sample preparation steps prior to HPF inevitably leave time for receptor trafficking that can affect labeling efficiency, and we also cannot exclude the possibility that some amount of immunolabel unbinding or epitope blocking occurs. We also emphasize that confident identification of these small AuNP labels and multiplexed label distinction require high-quality tomogram alignment. We overcame this challenge by using the AuNPs to guide realignment of synaptic regions, which greatly improved local alignments and demonstrated that these labels have a dual purpose as target identifiers and local fiducials. Notwithstanding these limitations, our studies highlight how new methods using small multiplexed gold nanoparticles, cryo-FIB milling, and cryo-ET, enable the localization of two distinct molecular complexes within constricted and crowded regions of intact brain tissue, opening new avenues to a molecular-level understanding of brain function.

## Methods

### Hippocampal slice electrophysiology

Acute coronal hippocampal slices were prepared from 2–3-month-old male and female mice. Animals were anesthetized with isoflurane and euthanized before brains were rapidly placed in ice-cold cutting solution containing 93 mM *N*-methyl-D-glucamine, 93 mM HCl, 25 mM NaHCO₃, 10 mM glucose, 20 mM HEPES, 10 mM MgCl₂, 5 mM L-ascorbic acid, 3 mM sodium pyruvate, 2.5 mM KCl, 2 mM thiourea, 1.2 mM NaH₂PO₄, and 0.5 mM CaCl₂. The solution was continuously bubbled with 95% O₂/5% CO₂. Coronal slices (270 µm) were prepared using a Leica VT1200S vibratome. The CA3 region was removed with a scalpel, after which slices were transferred to a holding chamber containing oxygenated cutting solution and allowed to recover for 10 min at 35 °C. Slices were subsequently maintained at room temperature in artificial cerebrospinal fluid (ACSF) containing 125 mM NaCl, 25 mM NaHCO₃, 10 mM glucose, 2.5 mM KCl, 1.25 mM NaH₂PO₄, 2 mM sodium pyruvate, 3 mM myo-inositol, 0.4 mM ascorbic acid, 1 mM MgCl₂, and 2 mM CaCl₂, continuously bubbled with 95% O₂/5% CO₂ until recording. All animal procedures were performed in accordance with the guidelines of Oregon Health & Science University Institutional Animal Care and Use Committees (IACUC). The protocol was approved by the IACUC of Oregon Health and Science University (IP00000905).

Whole-cell recordings were performed on upright microscopes (Zeiss AxioExaminer D1 or Olympus BX51WI) equipped with 60× water-immersion objectives. Slices were transferred to a recording chamber (Siskiyou) and continuously perfused with oxygenated ACSF (∼4 mL/min) at room temperature (22 ± 2 °C). Recordings were obtained using either an EPC10/2 patch-clamp amplifier (HEKA Elektronik) or a Multiclamp 700B amplifier (Molecular Devices). Borosilicate glass recording pipettes (2–4 MΩ) were filled with an internal solution containing 35 mM CsF, 100 mM CsCl, 10 mM EGTA, 10 mM HEPES, and 5 mM QX-314 (325 mOsm; pH 7.30 adjusted with CsOH). Data were acquired at 50 kHz and low-pass Bessel-filtered at 2.9 kHz.

Neurons were voltage-clamped at −70 mV or +40 mV to record spontaneous AMPA receptor-mediated or NMDA receptor-mediated EPSCs, respectively. For evoked EPSC recordings, a glass, unipolar stimulating electrode filled with ACSF was positioned in the stratum radiatum at least 100 µm from the recorded neuron. NMDAR-mediated evoked responses were quantified 60 ms after the stimulus to avoid contamination with the AMPAR-mediated response. To evaluate synaptic function under high-pressure freezing buffer conditions (see “Fab-AuNP immunolabeling and high pressure freezing of hippocampal slices” section), recordings were initiated in oxygenated ACSF perfused at 4 mL/min. The bath solution was then sequentially exchanged with HEPES-based buffer followed by HEPES buffer containing 20% w/v dextran. Finally, the recording chamber was rapidly exchanged by manually applying 4 mL of HEPES buffer containing 20% dextran and 5% sucrose. Picrotoxin (100 µM) was included in all bath solutions to isolate AMPA receptor-and NMDA receptor-mediated synaptic responses. EPSC measurements and spontaneous EPSC detection were performed as previously described^61^ using custom-written routines in Igor Pro 9 (WaveMetrics).

### Synthesis of monomeric AuNPs

AuNPs (∼2.2 nm diameter) utilized for preparation of 15F1 “C-term”, 15F1 “H4K-2Cys”, 11B8 “C-term”, and 5F11 “C-term” Fab-AuNP conjugates were synthesized as previously described^21,62,63^, using a 7:1 molar ratio of 3-mercaptobenzoic acid (3-MBA) to gold (III) chloride trihydrate (HAuCl_4_·3H_2_O). Slightly smaller (∼1.5 nm diameter) AuNPs used in preparation of the anti-GluN1 5F11 “C-term” Fab-AuNP_dimer_ and 15F1 “H1-2Cys” conjugates were synthesized following similar methods and using a 5:1 molar ratio of 3-MBA to HAuCl_4_·3H_2_O ^22,64,65^. Dried AuNP pellets were resuspended in water to 5-10 mg/mL. Each independent preparation was run on a 10% glycerol, 12% polyacrylamide gel electrophoresis (PAGE) gel to assess homogeneity and stored at 4°C until use. Under these conditions, the AuNPs remain stable for at least one year.

### Preparation of anti-GluA2 15F1, anti-GluA1 11B8, and anti-GluN1 5F11 Fab-AuNP conjugates

The anti-GluA2 15F1 Fab “C-term” conjugate was designed with an extended heavy chain containing a free cysteine for AuNP conjugation (C-terminal sequence: KVDKK-IVPRDAGAKP**<u>C</u>**) and with a 3C cleavage site and Twin Strep-tag at the C-terminus for affinity purification. The anti-GluA1 “C-term” 11B8 Fab^29^ and the anti-GluN1 5F11 “C-term” Fab^48^ were engineered based on this 15F1 scaffold by appending an identical 11-amino-acid C-terminal sequence (IVPRDAGAKP**<u>C</u>**) to introduce a similarly reactive cysteine on the heavy chain for site-specific AuNP conjugation. All Fab conjugates were expressed in Sf9 insect cells using a baculovirus expression system, purified from cell supernatant over Strep-Tactin resin, and conjugated to AuNPs as previously described^21^. The Fab-AuNP conjugates were PEGylated by incubation with 0.5 mM of PEG550-SH (Creative PEGWorks) at 37°C for 60 minutes. Excess PEG550-SH was removed by serial concentration and dilution in TBE buffer (100 mM Tris base, 100 mM boric acid, 2.5 mM EDTA), and the final PEGylated Fab-AuNP conjugates were stored at 4°C.

The concentration of PEGylated Fab-AuNP conjugates was estimated by gel electrophoresis, comparing the heavy and light chain Fab bands from the PEGylated Fab-AuNP sample and a concentration range of unconjugated identical Fab standards run in denaturing buffer on a Tris-Glycine gel. Using this gel-based quantitation, we estimate the molar extinction coefficient at 280 nm of these Fab-AuNP conjugates to be approximately 1.5 µM^-1^cm^-1^ (with a typical 280 to 500 nm absorbance ratio of ∼2) by nanodrop measurement (DeNovix DS-11). Additionally, the binding efficiency of Fab-AuNP conjugates was assessed by fluorescence-detection size-exclusion chromatography (FSEC) ^66^ by comparing the extent of purified GFP-tagged antigen peak shift between Fab-AuNP conjugate and Fab standards. Using these metrics, we observe that the PEGylated Fab-AuNP complexes maintain antigen binding at similar affinities as the unconjugated Fabs. These PEGylated Fab-AuNP conjugates can be stored at 4°C for at least a year without noticeable degradation or loss of binding efficiency.

The optimized “rigidly conjugated” anti-GluA2 15F1 “H4K-2Cys” and “H1-2Cys” Fab conjugates were prepared in essentially the same way as the described above for the “C-term” conjugates, and their design and AuNP conjugation are described in detail elsewhere^31^. Briefly, the 15F1 “H4K-2Cys” Fab contained heavy chain mutations S192C, S196C, and E197K and 15F1 “H1-2Cys” contained light chain mutations S122C and T126C for AuNP conjugation to multiple cysteines on structured α-helical regions on the Fab constant domain.

### Preparation of dimeric AuNPs (AuNP_dimer_) and the 5F11 Fab AuNP_dimer_ conjugate

The AuNP_dimer_ and the anti-GluN1 5F11 “C-term” Fab**-**AuNP_dimer_ conjugate were prepared as previously described^22^. To verify formation of functional anti-GluN1 5F11 Fab**-**AuNP_dimer_ conjugate, binding to antigen and conjugate concentration were assessed by FSEC and SDS-PAGE, respectively, as described for the other Fab-AuNP conjugates. After PEGylation, the conjugated Fab AuNP_dimer_ was immediately used for brain-slice labeling experiments.

### Acute mouse hippocampus slice preparation for cryo-ET

Adult male and female vGlut1-mScarlet (HET)/PSD95-EGFP (HET) mice (6-30 weeks) were used for the acute hippocampus slice preparation as described previously ^21^. Mice were anesthetized with isoflurane and brains were quickly removed after decapitation. Tissue blocks containing hippocampus were placed in a vibratome (Leica, VT1200). Horizontal brain slices (40 µm) were prepared at room temperature in one of two solutions. One solution was a HEPES-based buffer containing 150 mM NaCl, 2.5 mM KCl, 2 mM CaCl_2_, 2 mM MgCl_2_, 10 mM HEPES, and 0.005 mM MK-801 (pH 7.3). A second solution was artificial cerebrospinal fluid (ACSF) buffer containing 126 mM NaCl, 2.5 mM KCl, 1.2 mM MgCl_2_, 2.4 mM CaCl_2_, 1.2 mM NaH_2_PO_4_, 25 mM NaHCO_3_, 11 mM D-glucose, and 0.005 mM MK-801. The ACSF buffer was bubbled with 95%O_2_/5%CO_2_.

### Fab-AuNP immunolabeling and high pressure freezing of hippocampal slices

Hippocampal brain slices were incubated with the corresponding Fab-AuNP conjugate (2 µg/mL or ∼ 40 nM) in HEPES buffer or 95%O_2_/5%CO_2_ oxygenated ACSF solution for 1 hour at room temperature on an orbital shaker (180 rpm). The AMPA receptor antagonist, ZK-200775 (1 µM)^67^, and the positive allosteric modulator, RR2b^68^ (1 µM), were included throughout the experiments. After three washes of 15 mins each, CA1 regions of the hippocampi were manually dissected with a scalpel knife and were then incubated in the HEPES buffer or in the 95%O_2_/5%CO_2_ oxygenated ACSF solution containing 20% dextran for 30 min.

Prior to high pressure freezing (HPF), planchettes and grids were pretreated as described previously^21^. The sample and 6 mm flat specimen planchettes were assembled at the loading station of a Leica EM ICE. Two CA1 tissue slices were placed on 200 mesh gold extra thick carbon film grids (CF200-Au-ET; EMS) with apical dendrites facing toward the center of the grid and CA1 cell layers ‘vertical’, aligned with a rim mark located at the ‘12 o’clock’ position of the grid. This slice orientation enabled us to target Schaffer collaterals, where CA3 axons synapse onto CA1 apical dendrites, based on the relative grid location.

Cryoprotectant solution (20% dextran, 5% sucrose) and PEGylated 10 nm gold fiducials (CP11-10-PA-1K-DI water-50, Nanopartz) were applied to the sample prior to HPF to limit crystalline ice formation and aid tilt series alignment, respectively. A volume of 100 µL of gold fiducials was spun down at 15,000 rpm for 20 min and the supernatant was removed. After adding 100 µL of the HEPES or ACSF buffer, the centrifugation step was repeated. After removing the supernatant, 30 µL of HEPES or ACSF buffer containing 20% dextran and 5 % sucrose was added to the fiducial pellet. For samples prepared in HEPES buffer, a droplet with 1-2 µL of HEPES base cryoprotectant solution with gold fiducials was applied directly on top of the CA1 slices ∼5-10 sec immediately before HPF. For samples prepared in ACSF buffer, CA1 slices were incubated in 95%O_2_/5%CO_2_ oxygenated ACSF containing 20% dextran including gold fiducials (∼20 nM) for approximately 30 mins prior to HPF. In this case, 1-2 µL of oxygenated low Na ACSF solution containing 75 mM NaCl, 2.5 mM KCl, 1.2 mM MgCl_2_, 2.4 mM CaCl_2_, 1.2 mM NaH_2_PO_4_, 25 mM NaHCO_3_, and 11 mM D-glucose with 20% dextran and 5% sucrose was applied on top of the sample immediately before HPF. For all samples, after applying the cryoprotectant solution, a second planchette was placed on top of the sample grid with flat side down and the sample was subject to HPF. After HPF, sample grids were clipped using a cryo-FIB autogrid ring and C-clip, with the carbon layer side facing the autogrid ring and the sample side facing the C-clip.

### Cryo-FIB milling of tissue samples

The HPF hippocampal brain slice grids were milled on an Aquilos 2 cryo-FIB/SEM microscope (Thermo Fisher Scientific). After taking SEM tile images of an entire grid, sputter coating with a platinum layer (30 mA current for 30 s at 10 Pa pressure) and gas injection system (GIS) deposition of a trimethyl(methylcyclopentadienyl) platinum layer (25 sec) were applied. The grids were oriented with the Schaffer collaterals parallel to the ion beam path such that the resulting lamella targets could be chosen 2-4 grid squares away from the center line on both left and right sides and up to 12 grid squares in a vertical orientation, depending upon the smoothness of the surface of the grid squares. The HPF brain tissue lamellae were made using the ‘waffle’ method^21,69^. At a 90° milling angle, two rectangular trench cuts were milled with an ion beam current of 15 nA until the front and back of the lamella target site were completely cleared. The sample shuttle was moved to the regular milling orientation, and then the bottom sections of the lamella site were removed. At the 20° milling angle, the notch patterns were milled at the side of the lamella sites. Another GIS deposition for 20 sec and sputter coating for 15 sec were then applied.

Automated milling was performed using AutoTEM software with the milling angle of 20° and a set final lamella thickness of 140-160 nm. AutoTEM settings used for each of the steps were the following; rough milling at the ion beam current of 1 nA using rectangle patterns with a pattern offset of 1 µm, medium milling at 0.5 nA using cleaning cross sections (CCS) with a pattern offset of 0.8 µm, fine milling at 0.3 nA using CCS with a pattern offset of 0.6 µm, finer milling at 0.1 nA using CCS with a pattern offset of 0.4 µm, polishing 1 at 50 pA using CCS with over and under tilt of 0.5° and a pattern offset of 150 nm, and polishing 2 at 30 pA using CCS. After automated milling, the grids were tilted to +0.5° and-0.5° of the milling angle and manually polished with a cleaning cross section at 30 pA ion beam current to produce the final lamella. The milled grids were retrieved from the Aquilos 2 and stored in liquid nitrogen until they were imaged.

### Cryo-ET tilt series collection

Cryo-ET data were collected on FEI Titan Krios 300 keV transmission electron microscopes equipped with either Thermo Scientific Falcon4i and Selectris X or Gatan K3 and BioContinuum HD detectors and energy filters, respectively. One microscope was also configured with a spherical aberration corrector. Synaptic regions were targeted based on manual inspection of medium magnification (∼2500-6500x) montages of each lamella, looking for putative presynaptic compartments, identified by densely packed presynaptic vesicle features, next to putative postsynaptic compartments with darker features, indicative of PSD or actin. Tilt series were collected at each target using a dose-symmetric tilt scheme^70^ with a 3° tilt step starting normal to the lamella at-20° and going to maximum tilts of-68° and 28° with SerialEM^71^ or Tomography 5 (Thermo Scientific). Movies were collected at pixel sizes of 1.9-2.3 Å, a defocus range of-2.5 to-7 µm, and a total exposure of 120-150 e^-^/A^2^ (**Extended Data Table 1**).

### Tomogram reconstruction, segmentation, and membrane annotation

Movies were motion corrected and binned to a pixel size of 2.5 Å using MotionCor3^72^. Tilt series were initially reconstructed using AreTomo^73^, and both motion-corrected tilt series and AreTomo reconstructions were individually reviewed to identify tomograms that contained clear synapses. Selected tilt series were then aligned in Etomo (IMOD)^74^ by patch tracking (with a patch size of 200, 200 and fractional overlap of 0.7, 0.7) and also by fiducial tracking, if sufficient 10 nm gold fiducials were present. The tracking strategy resulting in the best alignment was chosen by manual inspection of both results and used for further processing. CTF parameters were estimated using Ctfplotter^75^ and tomograms were reconstructed in IMOD by regular weighted back projection with dose-weighting and 3D CTF correction and at a binned pixel size of 1 nm. Even and odd frame split versions of each tomogram were prepared in the same way and used for denoising model training. Alternatively, motion correction, CTF estimation, and tomogram reconstruction for the 15F1 Fab-AuNP “H4K-2Cys” dataset were performed with WarpTools^76^, using alignments from Etomo.

Reconstructed tomograms were denoised using DeepDeWedge^77^, which was trained on even and odd frame split versions of 3 to 5 tomograms per dataset. Membranes were segmented from the DeepDeWedge denoised tomograms with MemBrain-seg^78^, using the pretrained MemBrain_seg_v10_alpha.ckpt model checkpoint. Segmented membranes were imported into Blender using the MolecularNodes plugin^79^, and the segmentations were manually thresholded and cleaned to separate cellular compartments and remove noise. Membranes corresponding to presynaptic compartments, presynaptic vesicles, and postsynaptic compartments were manually selected and exported as independent meshes for further analyses. Presynaptic vesicle segmentations were only included if at least half of the vesicle was visibly segmented. To export fusing vesicle segmentation regions, the presynaptic membrane was duplicated and the region corresponding to putative fusing vesicle membrane was manually cropped for export.

### Synaptic cleft definition and MIP generation

Synaptic cleft regions were defined based on the annotated presynaptic and postsynaptic membrane meshes. For each pair of presynaptic and postsynaptic membranes, vertices within 10–40 nm of the opposing membrane were defined as “synaptic”. For automated generation of synaptic cleft minimum intensity projection (MIP) visualizations, only synaptic membrane vertices with a normal pointing towards the paired membrane were retained. The midpoint of these filtered mesh centroids was used to define the center of the synaptic cleft, and a direction vector **z_cleft_** was defined by averaging the vectors from 100 randomly chosen postsynaptic vertices to their closest presynaptic vertex. An orthogonal coordinate system for the synaptic cleft was calculated from this vector:

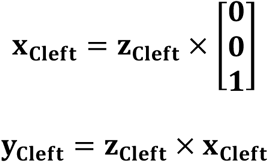

where × denotes the vector crossproduct.

To define the extent of a subvolume containing the synaptic cleft, a rotation matrix **R_cleft_** with rows **x_cleft_**, **y_cleft_**, and **z_cleft_** was used to transform all presynaptic and postsynaptic mesh vertices after centering them on the synaptic cleft center. The extent of the output subvolume along each axis was set to twice the maximum absolute transformed coordinate plus a 10 nm padding. The full transform from tomogram coordinates to subvolume voxel coordinates **M_cleft_** was then composed as a translation of the synaptic cleft center to the origin, followed by rotation by **R_cleft_**, followed by a translation placing the origin at the center of the output subvolume. The synaptic cleft subvolume was then extracted from the tomogram by resampling with the inverse of **M_cleft_**, using trilinear interpolation, via the affine_transform_image_3d function of the torch-transform-image package from teamtomo tools^80^. Three orthogonal minimum intensity projections (MIPs; XY, XZ, and YZ) were created from the extracted subvolume, forming the synaptic cleft minimum intensity projection (MIP) visualization. Picked AuNP positions (see below) were transformed using **M_cleft_** and overlaid on the MIP visualizations, which were inspected to assess the quality of both local tomogram alignment and AuNP picking.

### Tomogram AuNP monomer picking

A template for AuNP detection was generated from three tomograms of purified 15F1 “C-term” Fab-AuNP conjugate, collected with identical imaging parameters as tissue tomograms (binned voxel size 1 nm, equivalent tilt scheme and defocus range). AuNPs in these tomograms were identified using the findbeads3d program in IMOD^74^, yielding 28,409 detections. Subtomograms of 12 pixels were extracted at each detection and averaged to generate a template that captured characteristic aberrations present in the tomographic data. Template matching was performed using pytom_match_pick^81^ with the averaged AuNP template, without rotational search and with spectral whitening. Initial candidate positions were identified as local maxima with a cross-correlation score above 0.3 and a minimum inter-peak distance of 2 pixels. To obtain sub-pixel accurate AuNP positions, subtomograms extracted at each candidate position were fit with a 3D Gaussian model. The center of the fitted Gaussian provided refined coordinates and the Gaussian amplitude served as an intensity metric for each AuNP feature.

Neither the template matching score nor the Gaussian amplitude alone provided reliable discrimination of true AuNP detections, however, and optimal thresholds varied substantially between tomograms and synaptic regions due to differences in detector type, specimen thickness, and local alignment quality. Therefore, a range of template matching score and Gaussian amplitude thresholds were manually screened to maximize the accuracy of automated AuNP picking for each synaptic region. Afterwards, all template-based picks were manually checked and remaining false positives and negatives were corrected. In tomograms with visible platinum redeposition artifacts from cryo-FIB milling at the edges of the lamella, all putative AuNP picks within ∼6 nm of the tomogram edge were excluded from analysis (**Supplementary Fig. 3**). Additionally, AuNPs in tomogram subregions with locally poor alignment were excluded if they could not be confidently distinguished from noise or sufficiently improved by local realignment (see “AuNP-guided local tomogram realignment” section).

### AuNP-guided local tomogram realignment

When tomograms contained elongated AuNP features, suggestive of poor local alignment of the tilt series in our regions of interest (**Supplementary Fig. 2a**), we carried out a local realignment that leveraged the AuNP features. For tomograms containing only monomeric AuNPs, the center point of the volume of interest was manually identified or set by the computationally determined synaptic cleft center point as detailed above. Within a cubic volume extending 120 nm from the center point, coordinates of the local minimal intensity voxel were identified by inverting the tomogram intensities and applying the peak_local_max routine from scikit-image^82^. Alternatively, the AuNP coordinates given by 3D template matching and manual curation of AuNPs as described above were used (**Supplementary Fig. 2b**). For dual-labeled tomograms, coordinates for three dimensional Gaussian fitting were obtained from monomer positions provided by DeepETPicker or from manual picking. A three-dimensional Gaussian model was fit to each peak to determine sub-pixel precise centers and standard deviations (σx, σy, σz). Points above the 95th percentile in any dimension were removed, and to distinguish genuine AuNPs from noise and gold fiducial markers, sigma values were constrained using thresholds corresponding to the estimated AuNP size, taking into account the missing wedge distortion (2 × 2 × 5 nm) (**Supplementary Fig.2c**).

The refined AuNP centers were reprojected into the tilt series coordinate system using the alignment parameters generated by IMOD. A synthetic tilt series was generated by rendering a disk with a radius of 1.75 nm at each reprojected AuNP position (**Supplementary Fig.2d**). For each tilt angle, a cross-correlation map between the synthetic and experimental tilt images was calculated with a maximum search range of 25 nm. The position of optimal alignment was determined by fitting a two-dimensional Gaussian to the cross-correlation peak, and the resulting translational offsets were used to update the tilt-series alignment parameters (**Supplementary Fig. 2e**) before subsequent reconstruction (**Supplementary Fig.2f**). This pipeline was implemented in python via the scipy^83^ and teamtomo packages^80^. Realigned tomograms were reprocessed through the previously described tomogram denoising, membrane segmentation, and annotation workflow. Tomograms with independent AuNP-guided locally optimized alignments that focused on unique synaptic regions were subsequently analyzed as separate tomograms.

### Tomogram AuNP monomer and dimer picking

Monomeric and dimeric AuNPs were manually picked from tomograms using a Napari ^84^-based 3D visualization tool. Initially, 50 particles were selected for each class and used to train DeepETPicker^85^, where monomeric AuNPs were assigned to Class 1 and dimeric AuNPs to Class 2. Manual picking was performed on tomograms with a voxel size of 1 nm, whereas DeepETPicker training and inference were conducted on tomograms resampled to a voxel size of 0.5 nm. In the DeepETPicker configuration, the parameters label_diameter and ocp_diameter were set to 9 and (12, 20), respectively. Training and inference were iteratively performed while removing missing or incorrect picks to refine the dataset. The final training set consisted of 1,311 particles for Class 1 and 661 particles for Class 2. For dimeric AuNPs, DeepETPicker provided the center coordinates of each dimer pair. To determine the individual AuNP positions within each dimer, a local intensity-based refinement strategy was applied. Using the tomogram (voxel size 0.5 nm), a local patch with a radius of 10 voxels was extracted around each detected dimer center coordinate. Within this patch, the two darkest local intensity minima were identified within a 5 × 5 × 5 neighborhood and assigned as the coordinates of the two individual AuNPs. The initial DeepETPicker center coordinate was used only to localize the dimer for this refinement step. After identifying the two individual AuNPs, the centers of the two AuNPs were further refined by local intensity-based Gaussian fitting, and the final dimer center was recalculated as the midpoint between the two Gaussian-fitted AuNP centers. For dimers added manually during dataset curation, the approximate centers of the two AuNPs were manually identified, refined using the same Gaussian fitting procedure, and the dimer center was likewise defined as the midpoint between the two Gaussian-fitted AuNP centers. Using these DeepETPicker results as a starting point, we manually checked all picks to correct monomer and dimer assignments, false positives and negatives, and to remove any picks within ∼6 nm of tomogram edges when redeposition was present.

To independently assess the accuracy of the final monomer and dimer picks, we employed three separate validation tests. First, we compared the nearest neighbor distances between AuNPs in each set with those from purified samples of high-density 15F1 “C-term” Fab-AuNP or low-density 5F11 “C-term” Fab-AuNP dimer conjugates independently frozen on single particle cryo-EM grids in thin ice (**Supplementary Fig. 11a-b**). The dimeric AuNPs had a characteristic spacing of ∼4 nm and the AuNP monomers rarely get closer than 5 nm at high concentrations. Agreement between these nearest neighbor distributions in the purified and in-tissue datasets, with minimal distribution overlap, supports our monomer dimer label classifications in tissue. Second, we assessed the distance between monomer and dimer classified AuNPs and the postsynaptic membranes at synapses. Based on the unique epitopes and binding orientation of the 15F1 and 5F11 Fabs, and supported by our independent labeling experiments with 15F1 “C-term” Fab and 5F11 “C-term” Fab-AuNP monomer conjugates, we expect the AuNP to postsynaptic membrane distance distributions of 15F1-conjugated and 5F11-conjugated AuNPs to be differentially centered at 23 and 18 nm, respectively (**Extended Data Fig. 1c**). These values are in close agreement with those observed for each AuNP in the monomer or dimer-assigned classes in the dual labeled dataset, using the distances to both AuNPs within each dimer (**Supplementary Fig. 11c**). As a third validation test, four independent annotators manually picked monomer and dimer AuNPs positions for a representative subset of dual labeled data that included five synaptic regions. Among the group-annotated picks, 91.5% of assignments were in complete agreement and another 7.5% were consistent for 3 of 4 annotators. Only 1% of the picks were considered completely ambiguous, with annotators split in half on monomer or dimer class assignment. In comparison to the “final” monomer dimer AuNP classification assignments for this subset, 91.5% were in complete agreement among all 4 annotators, 7.1% had agreement between 3 of 4 annotators, and 1.4% agreed with 2 or less annotators (**Supplementary Fig. 11d**).

### Subtomogram averaging of AMPA receptors

To define putative receptor positions in the 15F1 “H4K-2Cys” Fab-AuNP labeled dataset, AuNPs within 17-23 nm of the postsynaptic membrane and with a pair distance of 5-9 nm were identified as potential pairs bound to a single receptor (**Extended Data Fig. 4a**). Possible receptor positions were defined by taking the vector between the AuNP pair center and the closest point on the postsynaptic membrane and picking the point along the vector 6 nm off the membrane, at the expected center of the receptor density. This vector was defined as the z axis and the x axis was set along a vector between the two AuNPs within the pair. From this oversampled set of putative receptor positions, a picked particle set was determined by keeping the maximum number of picks with the following constraints: 1) every AuNP could only be assigned to one particle pick and 2) all particle centers had to be at least 5 nm from each other, limiting picks that would have obvious steric clashes. Determination of these “maximum reasonable pick” solutions was automated with an integer linear programming algorithm^86^, which solved for the globally optimal solution under the assigned constraints for each synaptic region (**Extended Data Fig. 4b-c**).

Using this approach, 1,254 particles were picked from 55 tomograms and imported into RELION-5 v5.0.1^87^ at a 4.8 Å pixel size and in a 124 pixel box using WarpTools v2.0.0^88,89^. An initial round of 3D refinement in RELION was run with restricted sampling parameters (initial angular sampling = 0.1°; offset range = 1 pixel; offset step = 1 pixel; local searches from 0.1°), producing a reconstruction with clear AuNPs and pre-and postsynaptic membranes and with weak density corresponding to the 15F1 Fab and receptor ATD region (**Extended Data Fig. 4d**). A subsequent round of 3D refinement with restricted sampling and with a 100 Å diameter spherical mask that excluded the AuNPs and membranes resulted in improved density for the receptor ATD and LBD. Iterative subsequent rounds of 3D refinement incrementally masking smaller regions of the receptor led to modest further improvements in receptor density. The particles were then aligned to the receptor ∼2-fold axis and refined with C2 symmetry through two more rounds of 3D refinement masking the full receptor and then the ATD region. This map and alignment were further refined with an ATD-focused mask in M with MCore refinement iterations that included stage angle refinement, magnification/Cs/Zernike3, per-tilt series weight, per-tilt weight, and temporal pose resolution estimations. The resulting ATD-focused reconstruction had an ATD-masked Fourier Shell Correlation (FSC)-estimated resolution of 13.9 Å at an FSC cutoff of 0.143 (**Extended Data Fig. 4e**).

To push the resolution of receptor LBDs and TMDs, regions that are distal to the AuNPs and separated from the AuNPs by the ATD domains, an additional round of 3D refinement in RELION was run with a mask on the LBD:TMD regions. A round of 3D classification without alignment was then run with the same LBD:TMD mask, which identified a subset of 605 particles (48% of total) with better-resolved LBD:TMD density. This subset was run through another round of masked refinement, resulting in an LBD:TMD-focused reconstruction with an LBD:TMD-masked FSC-estimated resolution of 22.0 Å (**Extended Data Fig. 4d-e**). Subsequent attempts at refinement of the LBD:TMD region in M did not improve this LBD:TMD-focused reconstruction. The structure of native mouse hippocampal AMPAR (PDBID 7LDD)^30^ was fit to a composite map of the ATD-and LBD:TMD-focused refinements, which aligned well in the 15F1 Fab, ATD, LBD, and TMD regions and even included visible density for expected TARP auxiliary subunits that surround the receptor TMDs (**Extended Data Fig. 4f-g**). Map resolutions estimates were determined by RELION-5 FSC analysis and by local resolution estimation with RESOLVE^90^ (**Extended Data Table 2**). Receptor poses were visualized alongside membrane segmentations and AuNPs within tomograms using the ArtiaX^91^ toolbox in ChimeraX^92^. In figures, a surface representation of the previously determined hippocampal AMPAR structure (PDBID 7LDD) was used to illustrate the receptor poses.

### AMPA receptor interaction geometry

To determine whether AMPA receptors formed discrete interactions, we used the 15F1 “H4K-2Cys” Fab-AuNP labeled tomograms and analyzed them using two different approaches. First, AuNPs were assigned to AMPA receptors based on the subtomogram averaging analysis described above, with each receptor bound by two AuNPs. We considered two AMPA receptors as potentially interacting if the minimum distance between any AuNP of one receptor and any AuNP of the other was less than 12 nm. For all potential pairs, we measured two angles, θ_rot_ and θ_cc_. For each pair, the receptor whose two AuNPs were closer together was designated the reference. θ_rot_ is the angle between the reference receptor’s AuNP–AuNP vector and that of the other receptor, and θ_cc_ is the angle between the reference receptor’s AuNP–AuNP vector and the vector connecting the two receptors’ centroids (**Fig. 5g**).

As an independent approach that does not depend on the AMPA-receptor assignments from subtomogram averaging, we also treated each AuNP as a node in a proximity graph, connecting two AuNPs by an edge if their centers were closer than 12 nm. From this graph we extracted all sets of four AuNPs that formed a connected subgraph. Because the pairing of the four AuNPs into two receptor-like pairs is not known for this approach, each subgraph was partitioned into two pairs using the assignment that minimized the total intra-pair distance. As in the paired analysis, the pair with the shorter intra-pair AuNP–AuNP distance was designated the reference, and θ_rot_ and θ_cc_ were computed identically.

To generate reference distributions for fully ordered versus fully disordered AMPA packing, we simulated two 2D arrangements of AMPA receptors at matched density (∼5,500 receptors/µm², corresponding to 90 nm²/AuNP). The crystalline pattern was built by tiling a two-dimensional lattice (unit-cell vectors of 149.3 Å and 121.8 Å) with every receptor given the same in-plane orientation. The liquid-like pattern was built by placing receptors via random sequential adsorption of hard disks (exclusion diameter 100 Å) at the same density, with each receptor’s in-plane orientation drawn independently at random; residual steric clashes between receptor Cα atoms (AMPA receptor structure, PDB 7LDD) were then removed by iteratively displacing clashing receptor pairs apart within the membrane plane until no clashes remained. For both patterns, the two AuNP label positions expected per receptor were computed from the center of the AuNP density in the single particle analysis reconstruction of the AMPAR:15F1 “H4K-2Cys” Fab-AuNP complex^31^. To mimic positional uncertainty of AuNP localization relative to AMPA receptors, isotropic Gaussian noise (σ = 1 nm) was added to all simulated AuNP positions before they were passed through the same graph-construction, subgraph-extraction, and θ_rot/θ_cc calculation used for the experimental data.

### Annotation of molecular features in tomograms

In total, 260 tomograms containing 287 synaptic regions were determined to have sufficient quality alignment, synaptic membrane segmentations, and AuNP picks for subsequent analyses, as collectively judged by a team of seven curators (**Extended Data Table 1**). During team curation, tomograms were also placed into three categories based on the visible quality of tissue in the tomogram field of view: (1) tissue-like (consistent smaller spacing between cellular compartments), (2) semi tissue-like (slightly larger spacing or large gaps between some cellular compartments), and (3) synaptosome-like (isolated synaptosome with few surrounding compartments) (**Supplementary File 1**). These categories were assigned to 48%, 30%, and 22% of the synaptic regions, respectively. During initial tilt series alignment, reconstruction, and curation, molecular features including microtubules, actin, postsynaptic density (PSD), fusing vesicles, flotillin, and ribosomes were noted when observed. To prepare figures showing molecular representations of microtubules and flotillin, template matching was run with pytom_match_pick^93^ followed by manual curation of the picks to identify clear positive hits. Actin and PSD features were segmented manually in Dragonfly version 2025.1, when shown.

Due to the small number of flotillin and ribosome particles present in tomograms, template-matching alone did not provide a reliable metric for identifying these features with high confidence. Actin was also not identified reproducibly by template-matching or machine-learning approaches. Therefore, in a representative subset of 119 tomograms containing 132 synaptic regions (encompassing the entire 15F1 “C-term” and 15F1 “H4K-2Cys” datasets), an annotator screened each tomogram for actin, flotillin and ribosomes and recorded their localization in either the presynaptic compartment, postsynaptic compartment, or “other” unassigned compartment. For the same subset of tomograms, a second annotator repeated the screening for ribosome particles. A third annotator reviewed all putative ribosome picks together with the first two annotators to agree on a consensus set of ribosome features.

### Definition and analysis of synaptic cleft membrane and presynaptic vesicle features

Synaptic cleft-adjacent regions of the presynaptic and postsynaptic membrane meshes were defined as the region within 10-40 nm of the opposite synaptic membrane. The surface area of the postsynaptic cleft region was approximated as half of the surface area of the convex hull around all postsynaptic vertices within this 10-40 nm thresholded mesh region. The region of the presynaptic and postsynaptic segmentation meshes with vertexes facing towards the opposing cleft membrane (defined by a vertex normal having a dot product < 0 with the closest opposing membrane vertex normal) were considered to make up the “outer” membrane, and all other vertices in the synaptic region were considered as the “inner” membrane. The synaptic cleft width for each synaptic region was estimated as the average of the nearest neighbor distance between each point on the synaptic cleft regions of the “outer” presynaptic and postsynaptic membranes. The center of the cleft was defined as the geometric center of all synaptic cleft-designated vertices in the presynaptic and postsynaptic membrane for a particular synaptic region.

In many cases, the inner and outer membrane of presynaptic vesicles were initially segmented as separate vesicle annotations. To include only a single “outer” membrane segmentation of each presynaptic vesicle in subsequent analyses, the annotated presynaptic vesicle meshes were each fit to a sphere and the smaller of any overlapping spheres were removed from analysis. The shortest distance between each remaining vesicle mesh and the presynaptic cleft-adjacent membrane was calculated and used for definition of the set of “adjacent” vesicles and “fusion events”. The subset of vesicles with segmentations not touching, but within 20 nm of the presynaptic synaptic cleft membrane were defined as the “adjacent” vesicle set, whereas the subset of synaptic cleft-adjacent vesicles whose membranes were continuous with (defined as within 1 nm = 1 pixel of) the presynaptic synaptic cleft membranes were considered “fusion events”. Putative fusion sites for the “adjacent” and “fusion event” sets were defined for each vesicle as the geometric center of all points on the presynaptic synaptic cleft region within 20 nm of the vesicle segmentation, which typically places the site near the center of the fusion event pore. Vesicles that did not segment sufficiently for accurate localization were excluded from these analyses. For fusion analyses, we included vesicles in the combined set of all anti-GluA2 labeled tomograms using the 15F1 “C-term”, “H4K-2Cys”, or “H1-2Cys” label variants^31^, and from the anti-GluA2 15F1 Fab-AuNP monomer and anti-GluN1 5F11 Fab-AuNP dimer dual labeled dataset (**Extended Data Table 1**).

### Spatial analyses of AuNP labels

For monomeric AuNP analyses, the center of each AuNP pick after 3D Gaussian fitting was used for all spatial analyses, whereas for dimeric AuNPs, the center position of the dimer was used, unless otherwise noted. The nearest neighbor distance for each AuNP label was defined as the single closest distance to another identical label in the same synaptic region. The distance between each AuNP and either the presynaptic or postsynaptic membrane at the synaptic cleft was calculated as the average of the distances to the closest point on the respective “outer” and “inner” membrane regions, which approximates the distance to the “center” of each membrane segmentation. For the dimeric AuNP to postsynaptic membrane analysis, distances to both AuNPs within the dimer were calculated, rather than to the dimer center. As our definition of “inner” and “outer” membranes applied only at cleft membranes, the distance to the closest point on the postsynaptic membrane segmentation was used for comparison of synaptic and peri-or extrasynaptic AuNP-postsynaptic membrane distances. AuNPs were classified as “synaptic” if they were within 30 nm of the postsynaptic cleft membrane region, and otherwise manually classified as either perisynaptic (adjacent to a synaptic cleft region) or extrasynaptic (farther from visible cleft region and closer to spine neck, if visible). Perisynaptic and extrasynaptic AuNPs were grouped for all quantitative analyses due to ambiguity in their respective designations.

AuNP clustering was implemented using a Density-Based Spatial Clustering of Applications with Noise (DBSCAN) algorithm from the scikit-learn python library^94,95^ with a min_samples parameter of 1, which essentially reduces the algorithm to single-linkage distance-based clustering. Using higher min_samples values could successfully separate larger, elongated, and potentially overlapping clusters into globular sub-clusters, but would often leave out or separate AuNPs that were part of clear intra-receptor label pairs. Therefore, we chose to use the more conservative single-linkage distance-based clustering strategy to ensure that AuNP labels would be consistently co-clustered when within a distance range where co-localization could be attributable to AuNPs bound to the same receptor or to closely packed receptors that interact either directly or through scaffolding proteins. For this clustering interpretation, we chose the eps distance threshold to be ∼ 4-7 nm larger than the single particle AMPAR: Fab-AuNP structure-based average intra-receptor AuNP pair distance estimate for each label^31^. The structure-based average AuNP label pair distances and clustering eps cutoffs used for each Fab-AuNP label were as follows: 15F1 “C-term” (expected 9 nm; eps = 16 nm), 15F1 “H4K-2Cys” (expected 7 nm; eps = 14 nm), and 5F11 “C-term” (expected 24 nm; eps = 28 nm). In this analysis, we define a “cluster” as a group of at least two closely spaced receptors. Therefore, any resulting clusters with less than 4 AuNP labels, which represent less than two receptors under the assumption that the majority of AMPA and NMDA receptors have two GluA2 or GluN1 subunits, respectively, were considered “isolated”. For the 15F1 “C-term” Fab-AuNP paired analysis, AuNPs were considered “paired” if at least one other AuNP was within 16 nm (otherwise considered “single”).

Cluster area was estimated as the area of a concave hull around all cluster points projected to a plane of best fit, after expanding the hull by 5 nm to account for the receptor footprint outside of the AuNP points. Cluster AuNP density was reported as the total number of AuNPs in a cluster divided by the cluster area and typically reported as the inverse of this density (nm^2^/AuNP). To visually represent the local density of AuNP labels across the synaptic cleft, a cylinder-shaped sliding-probe was used to calculate local AuNP density across the cleft. Two percent of the outer-facing vertices of the postsynaptic membrane were selected to sample the AuNP density, chosen by greedily maximizing the distance between them so that they were as evenly spread as possible. At every sampled vertex, the number of AuNP labels within a cylinder with radius 25 nm and extending 50 nm outward along the locally averaged membrane normal vectors was divided by the area 625π (πr^2^) to calculate the AuNP density. The coordinates of sampled vertices were transformed using **M_cleft_** and densities were overlaid as a heat map on the synaptic cleft MIP visualizations, showing only values within a mask made up of the concave hull around sampled vertices (**Supplementary Fig. 9f**).

We employ an edge-corrected bivariate Ripley’s K function, as has been done in related studies^16,96^, to evaluate the spatial relationship between AuNPs and features of interest for pairwise analyses between (1) AuNP labels and the synaptic cleft center (2) AuNP monomer and AuNP dimer labels, and (3) AuNP labels and fusion event sites, where these sets define points in two classes. The bivariate Ripley’s K function was defined as a function of distance r from class 1 points as:

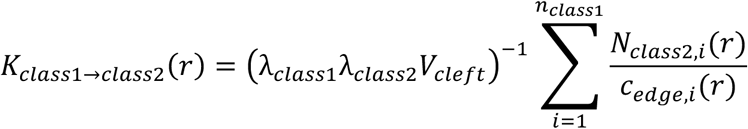

where *λ*_class1_ and *λ*_class2_ are the density of class 1 and class 2 points within the synaptic cleft, respectively, V_cleft_ is the volume of the synaptic cleft, defined as the partial volume of the convex hull around synaptic cleft presynaptic and postsynaptic membrane regions that sits between the two cleft membranes, N_class2,i_(r) is the number of class 2 points within distance r from each class 1 point, and c_edge,i_(r) is an edge correction factor for each class 1 point estimated as the fraction of a sphere centered at the class 1 point and of radius r located within the synaptic cleft volume. AuNPs outside of the synaptic cleft convex hull volume were excluded from Ripley’s analyses.

Because the feature assigned as class 1 or class 2 is meaningless but can lead to different results, we averaged both directions^97^:

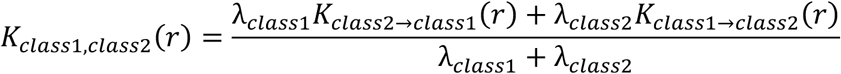

A 3D generalized, variance normalized version of this Ripley’s function is described as a Ripley’s H function with the following definition:

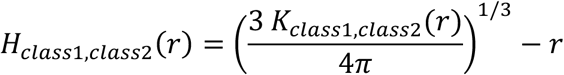

Under complete spatial randomness, H≈0 for all r, whereas distances where H > 0 and H < 0 are indicative of more or less class 2 points within r of class 1 points than expected by random chance, respectively. When pooling data from multiple synaptic regions, Ripley’s K functions from each region were averaged and then converted to a pooled Ripley’s H function.

To further test for enrichment of receptors at the center or edge of synapses, we calculated the pair correlation function (PCF)^97^ that quantifies enrichment of points in shells with thickness Δr (5 nm) around the cleft center.

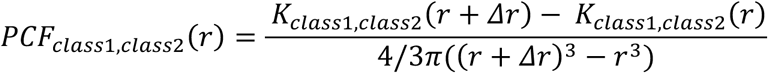

Under complete spatial randomness PCF≈1 for all r, with PCF > 1 and PCF < 1 indicative of more or less AuNPs observed than expected by chance within this distance shell, respectively.

To relate the observed bivariate Ripley’s function for monomer and dimer positions to modeled “extremes” that represent complete colocalization or independent clustering, two control setups were simulated using the full set of label positions for each synaptic region: (1) a “label permutation” strategy where the entire set of positions were randomly reassigned, keeping the same relative number of monomers and dimers, and (2) a “greedy clustering” strategy where a monomer or dimer was randomly assigned first, the remainder of matching labels were assigned to the next closest positions, after which the remainder were assigned to the alternative label (**Extended Data Fig. 7a-c**). For each control setup, 1000 simulations per synaptic region were run. A maximum absolute difference (MAD) test calculated from r = 30-50 nm (a range chosen to exclude the contribution of AuNP intra-receptor pairing but remain within the captured synaptic region) was used to assess difference from the “null” control models by comparison of T_observed_ and T_critical_ values^96^. These MAD test T values are defined as follows:

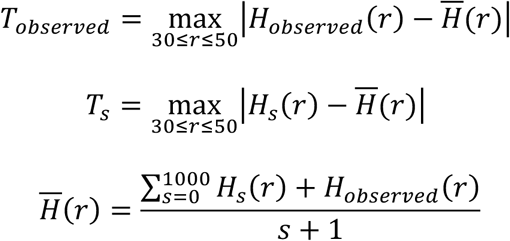

where s is the number of simulated null sets (s=1000). The critical T value (T_critical_) is defined as the 99^th^ percentile of the simulated T_s_ values, where the null hypothesis is rejected when T_observed_ > T_critical_.

To determine the spatial dependence of the observed Ripley’s functions between fusion events and monomer or dimer AuNP labels, three control vesicle sets were compared: (1) non-fusing “adjacent” vesicles within 20 nm of the presynaptic cleft membrane, (2) fusion event sites shifted 40 nm in a random direction along the presynaptic membrane, and (3) label permutated fusion event and AuNP label sites. The “40 nm shifted” sites were generated in 100 replicates per synaptic region, and the Ripley’s K function was averaged over all replicates. The label permutated sites were attained by randomly reassigning the set of fusion events and AuNP label sites and the Ripley’s K function was averaged from 100 permutations per synaptic region. When a fusion point was permutated to a site originally assigned to an AuNP position, it was shifted to the closest point on the presynaptic membrane to ensure consistency with the observed fusion sites. In addition to Ripley’s analyses, the cumulative number AuNP labels within r distance of fusion event, adjacent vesicle, 40 nm shifted, and label permutation fusion sites were calculated and averaged over all synaptic regions.

Significance of AuNP distance comparisons were determined by Welch’s t test, using GraphPad Prism version 11.0.1. Significance of differences between Ripley’s H functions comparing AuNP label and cleft center or fusion site positions were determined using a paired or unpaired permutation and Monte Carlo test of the integrated squared difference of the mean Ripley’s H curves over 0-100, 0-200, and 0-500 nm distance ranges (**Supplementary Table 1**). The integrated square difference T values between two Ripley’s H analyses were defined as

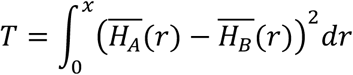

where H_A_ and H_B_ are the Ripley’s H curves from the two groups being compared (e.g. 15F1 and 5F11 labeling) and x is the analyzed distance range (100, 200, or 500 nm). Unpaired permutation tests were used to compare labels from different tomogram sets. For this test, curves from two sets were randomly reassigned into two groups for each of N=10,000 permutations, preserving the original group sizes, and T was computed for each permutation. A Monte Carlo p-value was defined as follows

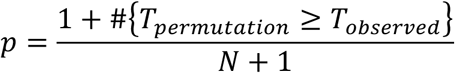

For dual labeled data that contain monomer and dimer labels within the same cleft, paired permutation tests were run using the same Monte Carlo p values as the unpaired permutation tests but instead comparing to randomly swapped monomer and dimer labels for each cleft over N=10,000 permutations. T values were calculated as follows

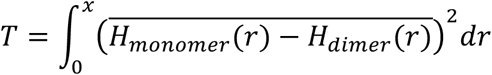

In the case of comparing observed Ripley’s curves to simulated 40 nm shifted or label permutated vesicle positions, a Diggle Monte Carlo was used, where a null reference curve was calculated as follows

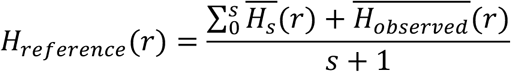

and s is the number of simulated curves. T and p values were calculated by taking the integrated square difference from the reference curve as follows

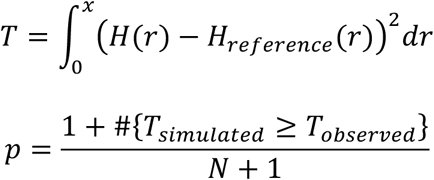

## Data Availability

The cryo-EM STA maps of 15F1 “H4K-2Cys” Fab-AuNP: AMPA receptor complex have been deposited in the Electron Microscopy Data Bank (EMDB) under accession numbers EMD-79324 and EMD-79325 (see Extended Data Table 2). All raw tilt series data, reconstructed tomograms, and associated annotations are deposited in the Biohub CryoET Data Portal.

## Supporting information

Supplementary Information Guide

## Acknowledgements

We thank Rachel Courtney for assistance with manuscript preparation, Ryan Hylton for input on ribosome annotations, and other Gouaux lab members for feedback on experiments and the manuscript. We also thank C. Jahr, G. Westbrook, E. Schnell, and B. Barad for input on the manuscript. We acknowledge the generous support and use of the HHMI Janelia Cryo-EM facility for FIB milling on an Aquilos2, which was operated by Momoko Shiozaki and Nikki Jean, and data collection on Krios microscopes, which were operated by Drs. Rui Yan, Shixin Yang, and James Jung (HHMI Janelia). A portion of this research was supported by NIH grant R24GM154185 and performed at the Pacific Northwest Center for Cryo-EM (PNCC) with assistance from Janette Myers and Nancy Meyer. We also acknowledge support for this work from the Chan Zuckerberg Imaging Institute, where a portion of the data was collected with assistance from Reza Paran, Elizabeth Montabana, Garrett Greenan and Nikki Jean. The research reported in this publication used computational infrastructure supported by the OHSU Office of Research Infrastructure Programs, Office of the Director, of the National Institutes of Health under Award Number S10OD034224. The content is solely the responsibility of the authors and does not necessarily represent the official views of the National Institutes of Health. We acknowledge the use of AI agents including Cursor, ChatGPT, and Claude Code in preparation and testing of data analysis code, which has all been made publicly available on GitHub. E.G. is an investigator of the Howard Hughes Medical Institute and acknowledges generous support from Bernard and Jennifer LaCroute.

## Author contributions

C.J.S., H.K., and G.D. prepared and characterized AuNPs and Fab-AuNP conjugates. A.M. and C.J.S. carried out cryo-ET tissue sample optimization and preparation. C.J.S., A.M., and H.K. collected and processed cryo-ET data. J.E. and E.V.S. assisted with cryo-ET data analysis. D.W., R.A., and Y.S. performed slice electrophysiology experiments under the guidance of S.J. C.J.S., J.E., A.M., and E.G. wrote the manuscript with input from all authors.

## Materials & Correspondence

Correspondence and material requests should be made to Eric Gouaux and Johannes Elferich.

## List of Figures, Tables, Other Items

**Ext. Fig. 1** Fab-AuNP conjugates enable precise AMPA and NMDA receptor labeling

**Ext. Fig. 2** Synaptic AuNP-focused cryo-ET data processing pipeline

**Ext. Fig. 3** AuNP labels identify synaptic and non-synaptic receptor positions

**Ext. Fig. 4** Subtomogram averaging supports AuNP-guided receptor placements

**Ext. Fig. 5** NMDA receptors cluster throughout the synaptic region

**Ext. Fig. 6** AuNP monomers and dimers are distinguishable in tomograms

**Ext. Fig. 7** AMPA and NMDA receptors are intermixed at synapses

**Ext. Fig. 8** Putative synaptic vesicle fusion events and vesicle-associated densities are observed at synapses

**Ext. Fig. 9** AMPA receptors are marginally enriched near fusion events

**Ext. Fig. 10** NMDA receptors are modestly more enriched at observed fusion events than AMPA receptors

**Ext. Table 1** Cryo-ET data collection

**Ext. Table 2** Cryo-ET subtomogram averaging data processing

**Sup. Fig. 1** Fab-AuNP conjugates enable precise AMPA and NMDA receptor labeling

**Sup. Fig. 2** AuNP-guided tomogram re-alignment improves local resolution for confident AuNP identification

**Sup. Fig. 3** FIB-mill redeposition on lamella surfaces is excluded from AuNP picking

**Sup. Fig. 4** Fab-AuNP labeling does not alter synaptic cleft width

**Sup. Fig. 5** Receptors and actin localize near postsynaptic density features

**Sup. Fig. 6** Flotillin cages are observed adjacent to synapses

**Sup. Fig. 7** Ribosomes are predominantly observed in postsynaptic compartments

**Sup. Fig. 8** GluA1-containing AMPA receptors are identified throughout the synaptic cleft

**Sup. Fig. 9** GluA2 labeling with 15F1 Fab “C-term” and “H4K-2Cys” conjugates provides reproducible clustering metrics

**Sup. Fig. 10** Pairwise geometrical analysis reveals liquid-like AMPAR packing

**Sup. Fig. 11** AuNP monomer and dimer classifications are validated by multiple metrics

**Sup. Table 1** Statistical comparisons for Ripley’s H analyses

**Sup. File 1** Additional tomogram slice, membrane segmentation, and MIP examples

**Sup Video 1** AuNP labels reveal AMPA receptor distribution in mouse hippocampal synapses

**Sup Video 2** Rigidly attached AuNP labels reveal multiple poses of AMPAR-AMPAR interactions

**Sup Video 3** Multiplexed AuNP labels reveal how AMPA and NMDA receptors intermingle

**Sup Video 4** Multiplexed AuNP labels defines the proximity of AMPA and NMDA receptors to vesicle fusion events

