## Extended Data for "Mapping AMPA and NMDA receptors at mouse brain hippocampal synapses by cryo-ET"

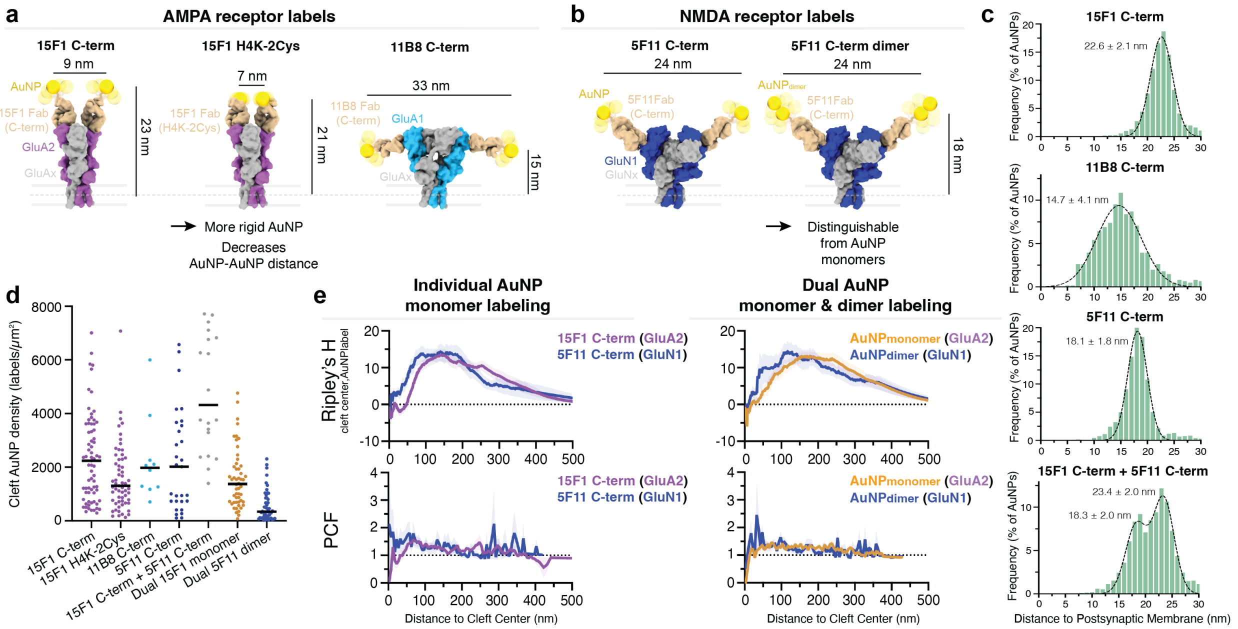


**Extended Data Fig. 1 Fab-AuNP conjugates enable precise AMPA and NMDA receptor labeling.** Molecular models of GluA2 or GluA1-containing AMPA receptors (**a**), or GluN1-containing NMDA receptors (**b**), bound to the indicated Fab-AuNP cryo-ET labels. **c**, Plot of AuNP distances to the center of the postsynaptic membrane for specified Fab-AuNP labels. Histograms are overlaid with single Gaussian or sum of two Gaussian fits and fit-derived mean and standard deviation values shown. **d**, Median density of AuNP labels within synaptic cleft for each labeling condition, per synaptic region. **e**, Bivariate Ripley's H and pair correlation functions between the synaptic cleft center and AuNP label positions. Mean and standard error of the mean shown.

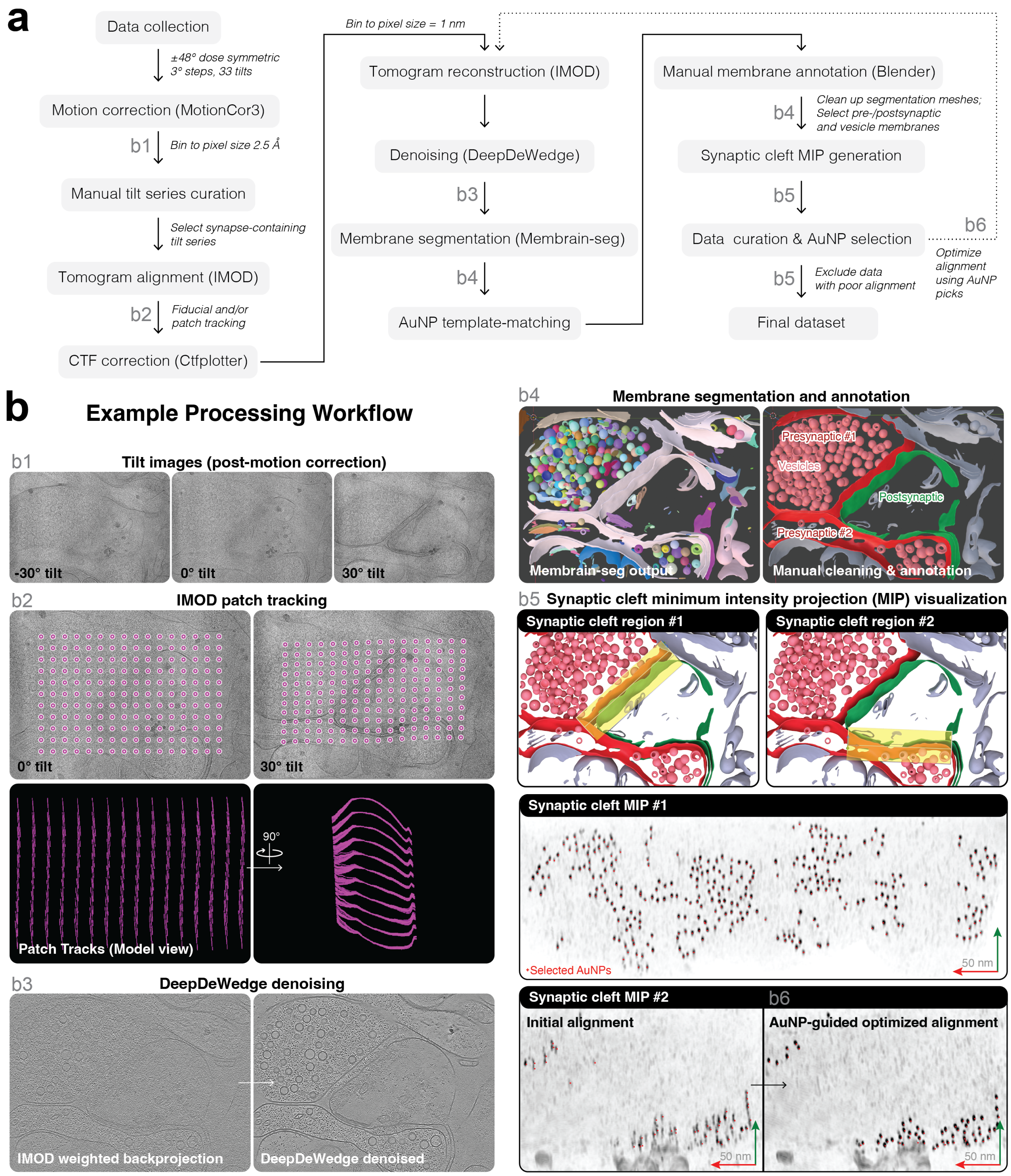


**Extended Data Fig. 2 Synaptic AuNP-focused cryo-ET data processing pipeline. a**, Workflow for cryo-ET data processing. **b**, Representative snapshots of processing steps b1-b6 noted in panel **a** for example anti-GluA2 15F1 "C-term" Fab-AuNP labeled tomogram.

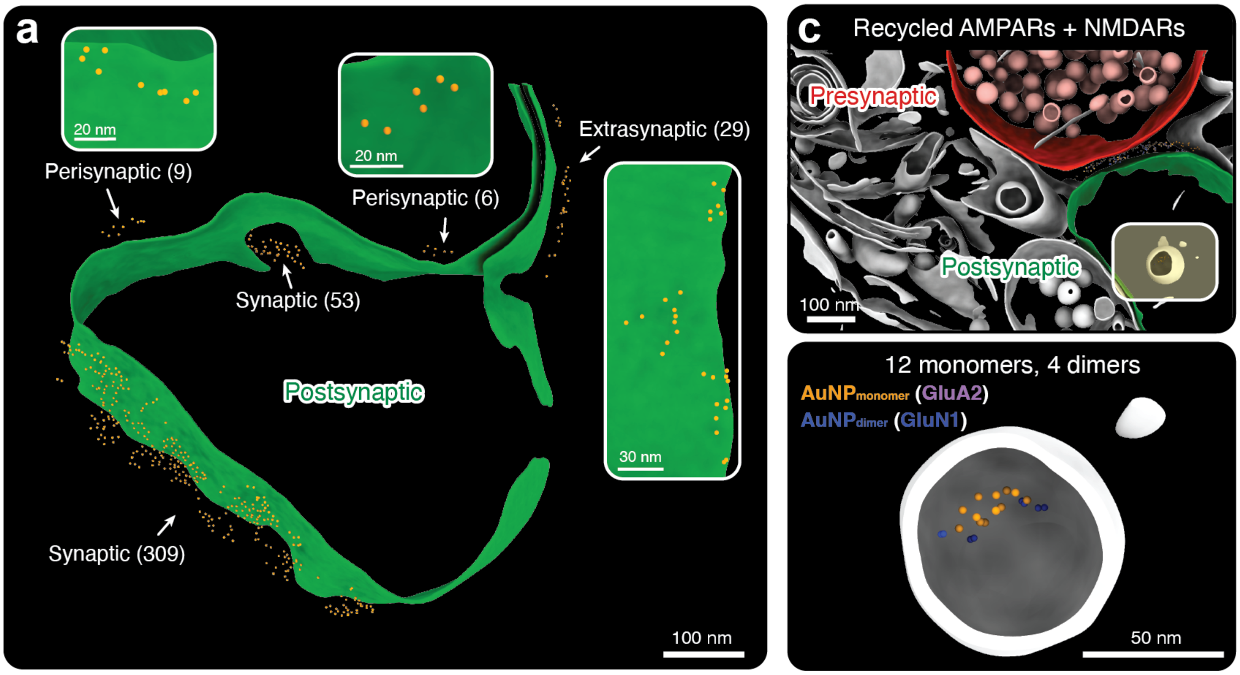


**Extended Data Fig. 3 AuNP labels identify synaptic and non-synaptic receptor positions. a**, Anti-GluA2 targeted AuNPs surrounding the postsynaptic membrane in synaptic tomogram from Figure 1 panel **e**. The number of AuNPs in each region are noted, along with insets showing local views normal to the membrane. **b**, Anti-GluA2 15F1 Fab-AuNP$_{monomer}$ and anti-GluN1 5F11 Fab-AuNP$_{dimer}$ dual labeled synaptic tomogram (top). Clipped view of boxed region showing intracellular postsynaptic compartment that contains co-internalized AMPA and NMDA receptors (bottom).

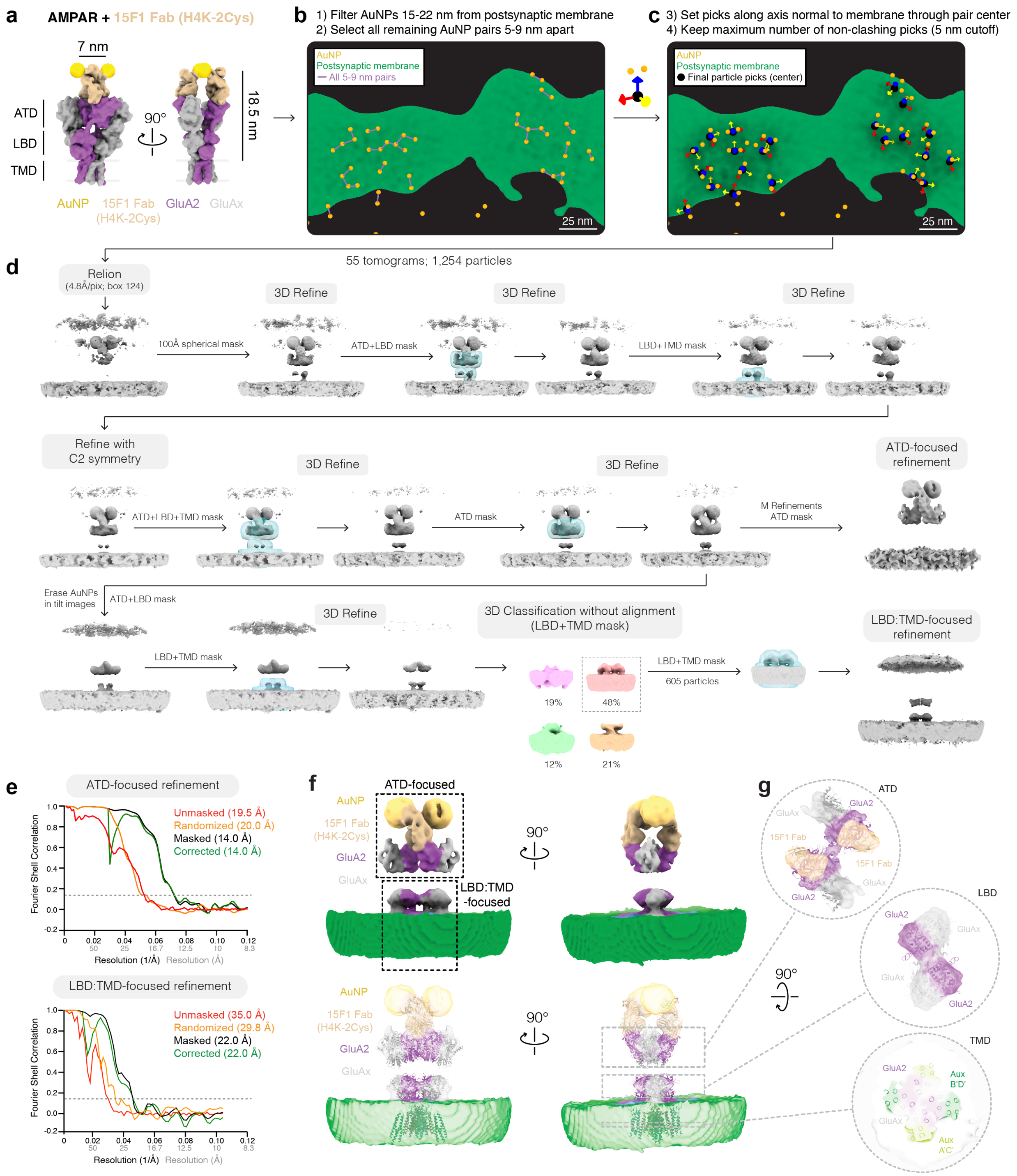


**Extended Data Fig. 4 Subtomogram averaging supports AuNP-guided receptor placements. a,** Molecular model of AMPA receptor bound to GluA2-specific 15F1 "H4K-2Cys" Fab-AuNP cryo-ET label. The amino terminal domains (ATD), ligand binding domains (LBD), and transmembrane domains (TMD) of the receptor are denoted. The experimentally determined average intra-receptor AuNP-AuNP and AuNP-postsynaptic membrane edge distances are 7 nm and 18.5 nm, respectively. **b-c,** AuNP-guided particle picking and pose assignment strategy (see Methods). **d**, Subtomogram averaging workflow resulting in ATD- and LBD:TMD-focused reconstructions at 14 and 22 Å resolutions, respectively. **e**, Relion-5 Fourier Shell Correlation plots for ATD-focused and LBD:TMD-focused reconstructions. **f**, Composite map of 15F1 “H4K-2Cys” Fab-AuNP: AMPA receptor complex (top) with fit model of native hippocampal 15F1 Fab: AMPA receptor complex (PDB 7LDD). **g**, Clipped views of map-model overlays at ATD, LBD, and TMD regions of the receptor.

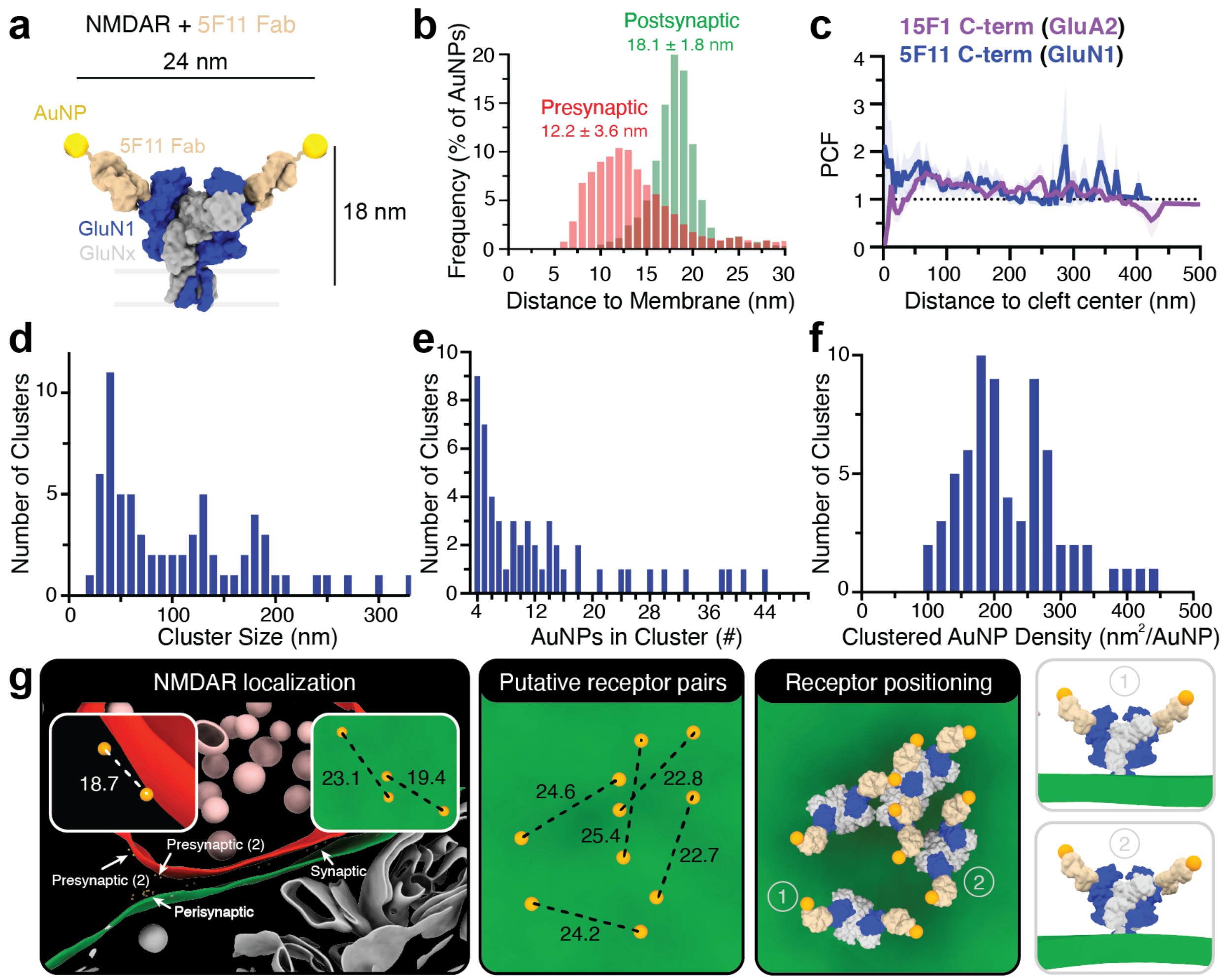


**Extended Data Fig. 5 NMDA receptors cluster throughout the synaptic region. a**, Molecular model of GluN1-containing NMDA receptor bound to 5F11 "C-term" Fab-AuNP cryo-ET label (based on PDBID 9PZV). **b**, Histogram of AuNP-membrane distances in 5F11 "C-term" Fab-AuNP labeled data (n=2,248 AuNPs). **c**, Pair correlation function (PCF) between synaptic cleft center and AuNP positions (bin size = 5). Mean and standard error of the mean shown (n=29 synaptic regions), overlaid with equivalent function for 15F1 "C-term" Fab-AuNP data (see Fig.2 panel **f**). Histograms of AuNP cluster sizes (bin size 10) (**d**), AuNPs per cluster (bin size 1) (**e**), and clustered AuNP inverse density (bin size 10) (**f**). **g**, Example tomogram containing synaptic, perisynaptic, and presynaptic localized 5F11 "C-term" Fab-AuNPs (left). Putative receptor pairs (middle) and NMDA receptor positions (right) are shown for the perisynaptic AuNP cluster, along with side views of two docked receptor examples that show stereotyped distance between AuNPs and postsynaptic membrane. 3D distances between putative receptor AuNP pairs are noted in nm.

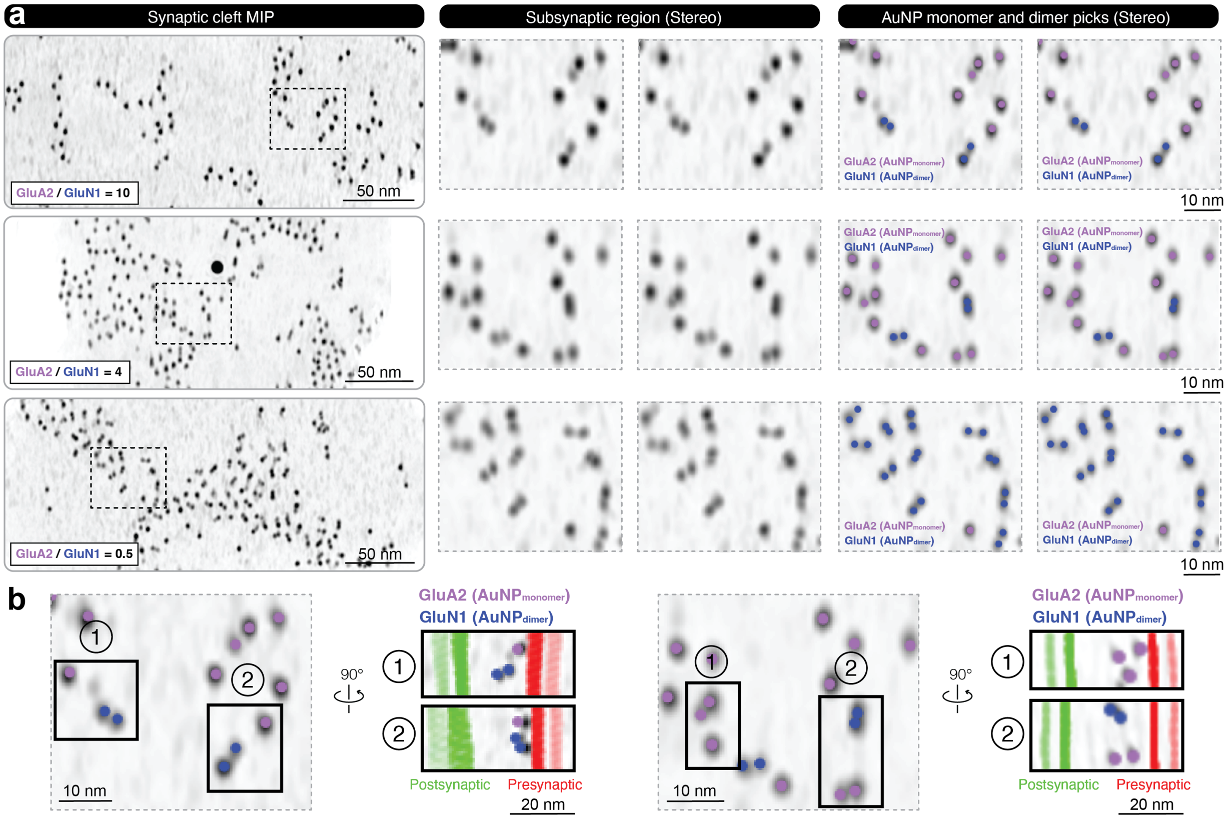


**Extended Data Fig. 6 AuNP monomers and dimers are distinguishable in tomograms. a**, Synaptic cleft minimum intensity projections (MIPs) from three representative anti-GluA2 15F1 Fab-AuNP$_{monomer}$ and anti-GluN1 5F11 Fab-AuNP$_{dimer}$ dual labeled tomograms (left), stereo views of boxed subsynaptic regions (middle), and overlaid monomer and dimer fits (right). **b**, Orthogonal side-on synaptic views of subregions from panel **a**.

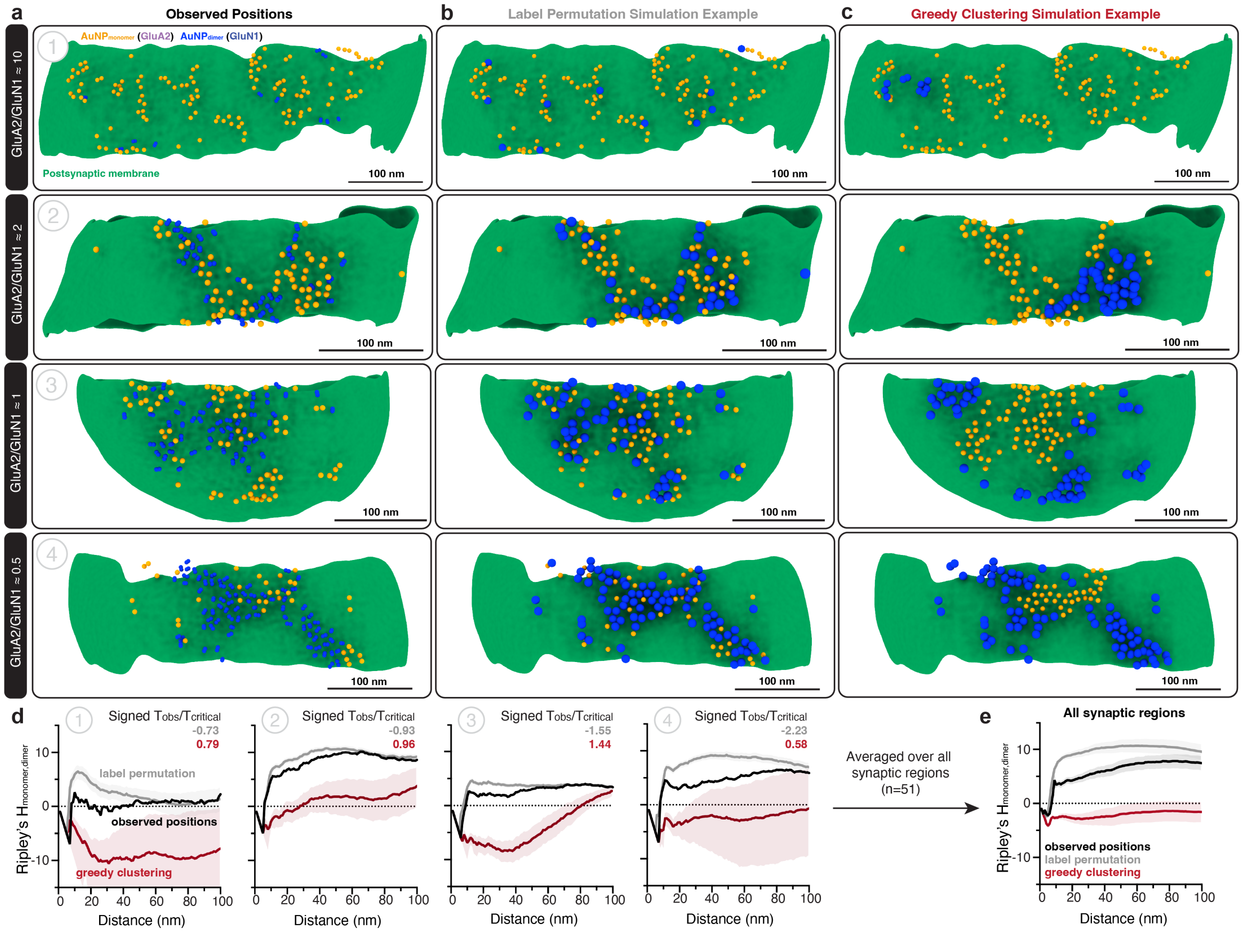


**Extended Data Fig. 7 AMPA and NMDA receptors are intermixed at synapses. a**, Four example AMPA and NMDA receptor dual labeled tomograms viewed normal to the postysynaptic membrane, with observed $AuNP_{monomer}$ and $AuNP_{dimer}$ positions denoted. AuNP spheres are shown at approximately ~2x scale for visibility. **b,c**, Single example of simulated $AuNP_{monomer}$ and $AuNP_{dimer}$ positions in label permutation or greedy clustering null models for equivalent tomograms from panel **a**. For simulated examples, the dimers are represented by a single blue sphere at the center of the simulated dimer position. **d**, Bivariate Ripley's $H_{monomer,dimer}$ function for each synaptic region from panels **a-c**. The mean and standard deviation (n=1000 simulations each) are shown. The maximum absolute difference test $T_{observed}/T_{critical}$ values, multiplied by the direction of change (Signed $T_{obs}/T_{critical}$) comparing the observed function with label permutation or greedy clustering simulated controls are shown (see Methods). **e**, Averaged Ripley's functions from all analyzed tomograms (n=51 synaptic regions). The mean and standard error of the mean (shaded region) are shown.

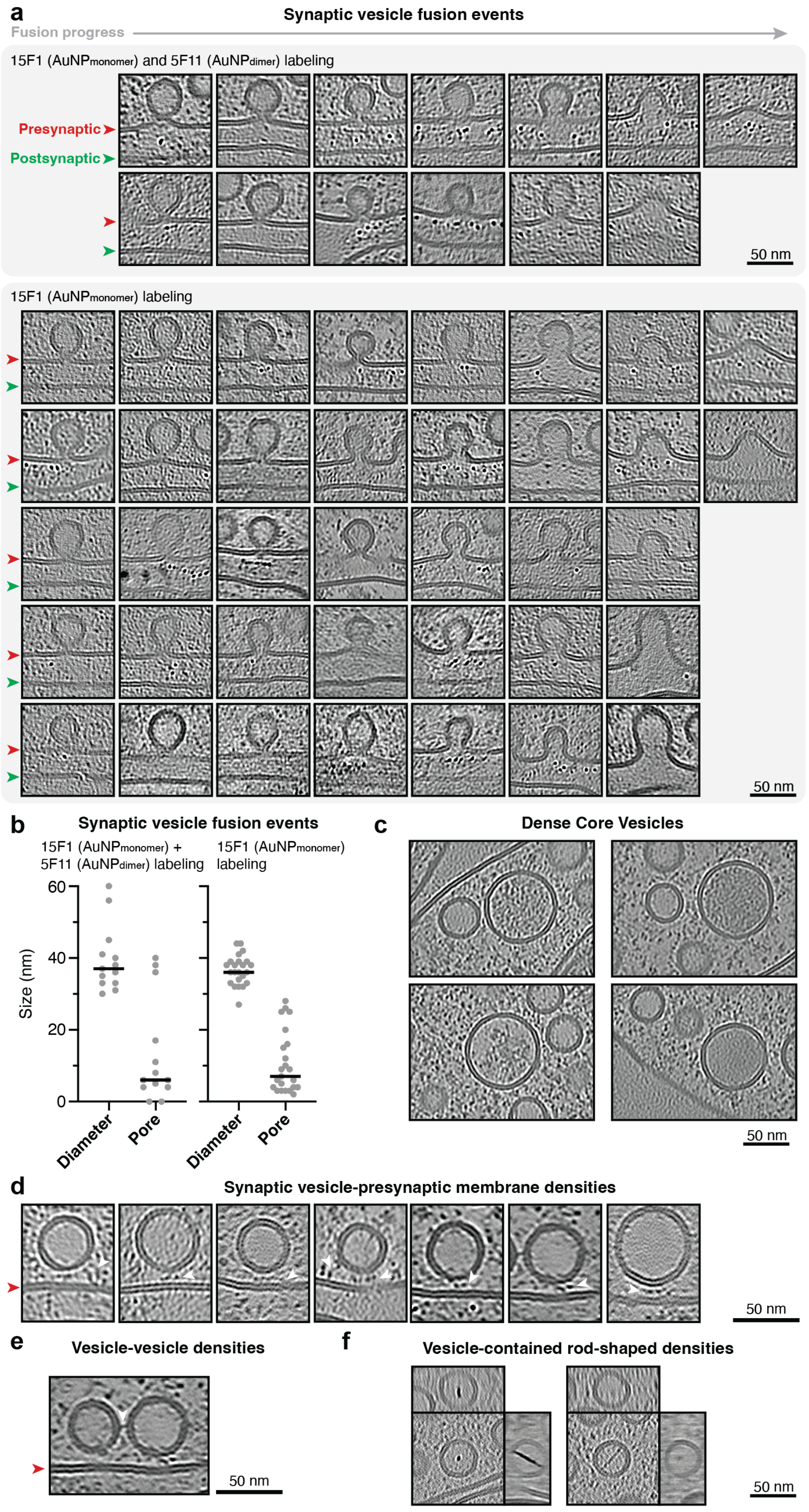

a
Synaptic vesicle fusion events
Fusion progress
15F1 (AuNPmonomer) and 5F11 (AuNPdimer) labeling
Presynaptic
Postsynaptic
50 nm
15F1 (AuNPmonomer) labeling
50 nm
b
Synaptic vesicle fusion events
15F1 (AuNPmonomer) + 5F11 (AuNPdimer) labeling
15F1 (AuNPmonomer) labeling
Size (nm)
60
40
20
0
Diameter
Pore
Diameter
Pore
c
Dense Core Vesicles
50 nm
d
Synaptic vesicle-presynaptic membrane densities
50 nm
e
Vesicle-vesicle densities
50 nm
f
Vesicle-contained rod-shaped densities
50 nm

**Extended Data Fig. 8 Putative synaptic vesicle fusion events and vesicle-associated densities are observed at synapses**. **a**, Denoised tomogram slices including putative synaptic vesicle fusion events from anti-GluA2 15F1-AuNP$_{monomer}$ and anti-GluN1 5F11-AuNP$_{dimer}$ dual labeled dataset (top) or anti-GluA2 15F1-AuNP labeled datasets (bottom). **b**, Fusing vesicle diameters and pore widths from examples shown in panel **a**. **c**, Four examples of vesicles with > 50 nm diameter and darker relative interior density, highlighted as putative dense core vesicles. **d**, Examples of densities connecting vesicles to synaptic cleft presynaptic membranes. **e**, Example of a density tethering two presynaptic vesicles together. **f**, Two examples of a rod-shaped density bridging across the inside of presynaptic vesicles.

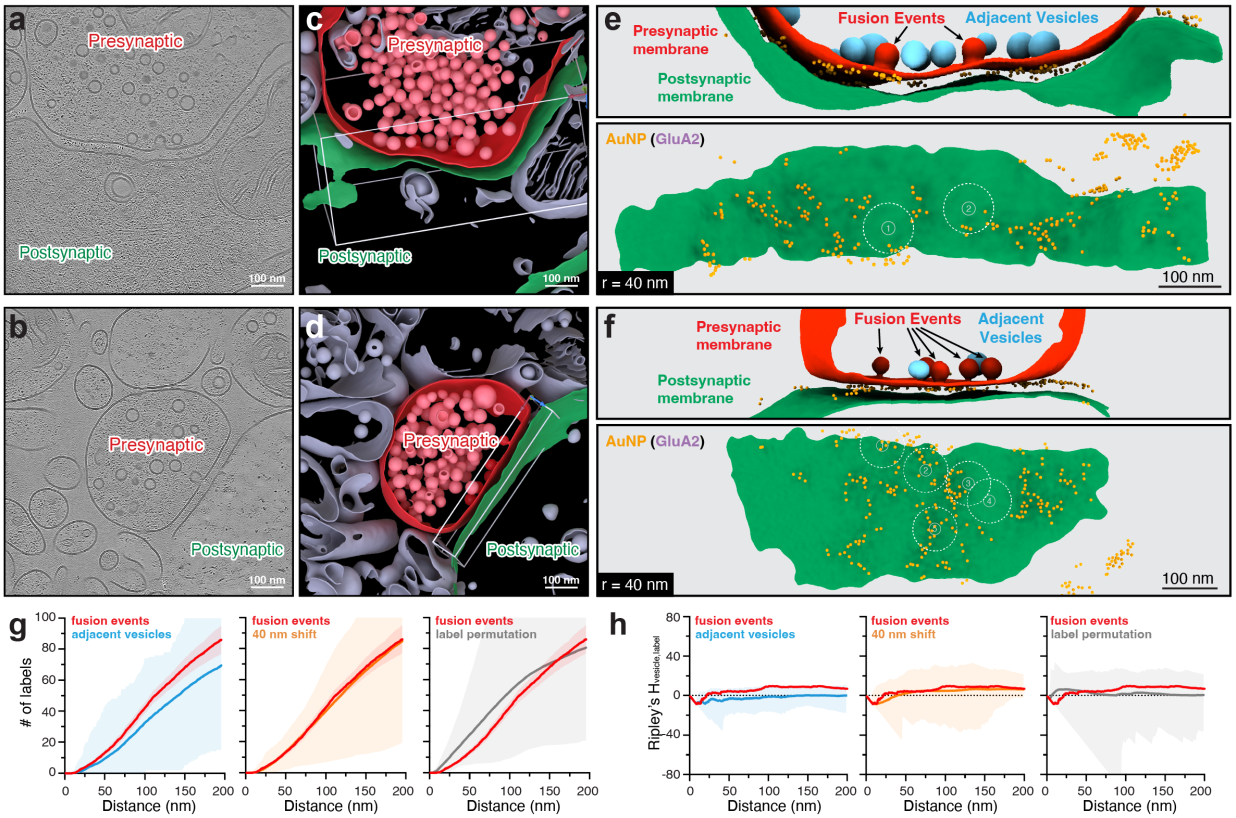


**Extended Data Fig. 9 AMPA receptors are marginally enriched near fusion events.** Tomogram slices (**a,b**) and segmentations (**c,d**) containing fusion events from 15F1 "C-term" Fab-AuNP labeled dataset. **e,f**, Synaptic cleft membranes and AuNP positions from tomograms shown in panels a-d, with fusion events and adjacent vesicles denoted. AuNPs are shown at twice their actual size for clarity. **g**, Cumulative frequency plots for 15F1-labeled data showing the average number of labels within a given distance from fusion events and adjacent vesicles (n=14 tomograms, 23 fusion events, 56 adjacent vesicles), fusion sites shifted tangentially 40 nm in a random direction ("40 nm shift"; n=100 samplings per vesicle), or a randomly permutated position ("label permutated"; n=100 samplings per cleft) (see Methods). Mean and standard error of the mean (fusion events), range per vesicle (adjacent vesicles and 40 nm shift) or range per tomogram (label permutation) shown by bold line and shaded regions, respectively. **h**, Bivariate Ripley's $H_{vesicle,label}$ between AuNP label positions and vesicle sites for fusing, adjacent, 40 nm shift, or label permutated sets. Mean and 95% envelopes are represented by bold line and shaded region.

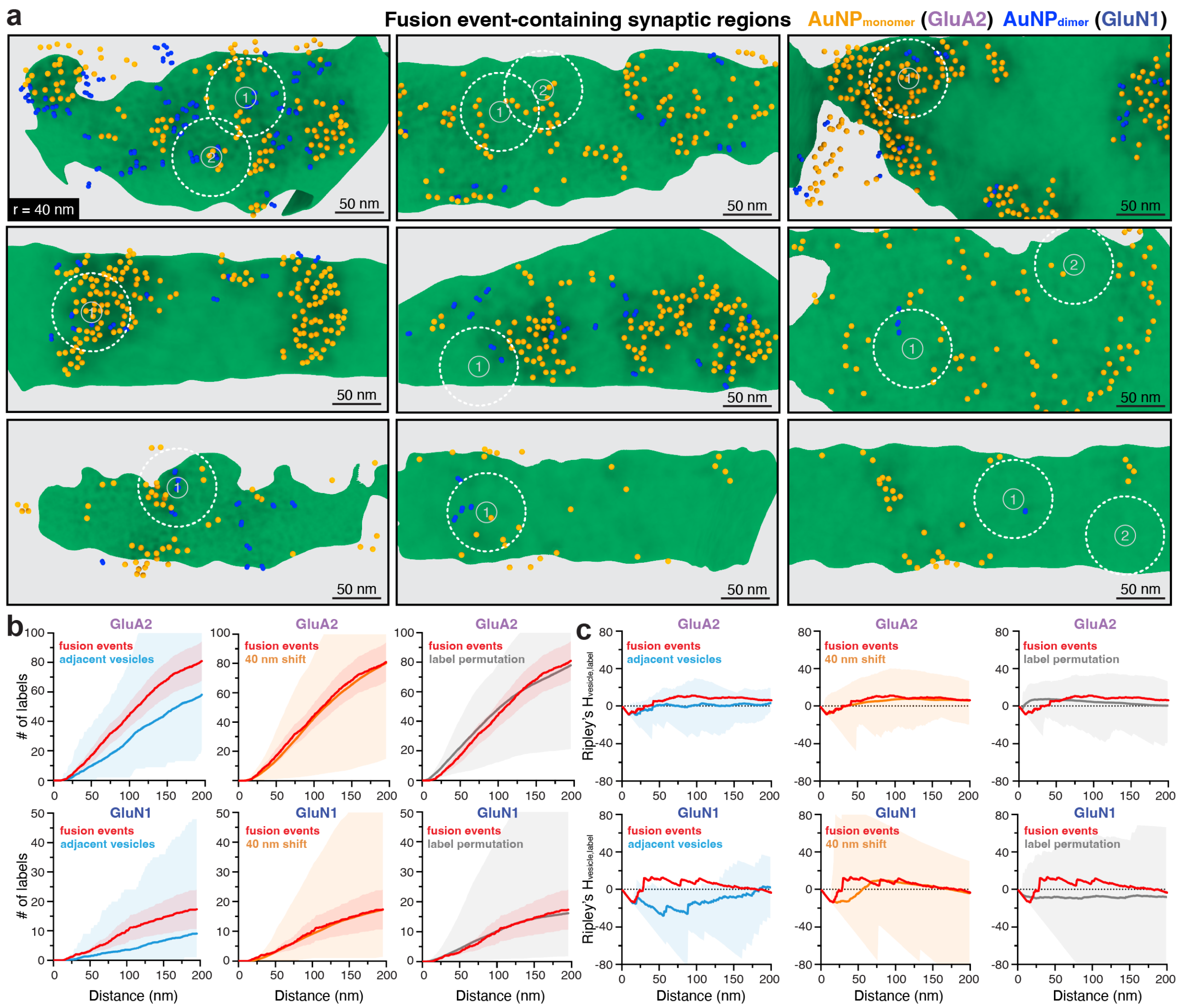


**Extended Data Fig. 10 NMDA receptors are modestly more enriched at observed fusion events than AMPA receptors. a**, Synaptic cleft regions viewed normal to the postsynaptic membrane showing all fusion events from anti-GluA2 15F1-AuNP$_{monomer}$ and anti-GluN1 5F11-AuNP$_{dimer}$ dual labeled dataset. AuNPs are displayed ~2x scale for clarity. **b**, Cumulative frequency plots for dual labeled data showing the average number of labels within a given distance from fusion events and adjacent vesicles (n=8 tomograms, 11 fusion events, 16 adjacent vesicles), fusion sites shifted tangentially 40 nm in a random direction ("40 nm shift"; n=100 samplings per vesicle), or a randomly permutated position ("label permutated"; n=100 samplings per cleft) (see Methods). Mean and standard error of the mean (fusion events), range per vesicle (adjacent vesicles and 40 nm shift) or range per tomogram (label permutation) shown. **h**, Bivariate Ripley's $H_{vesicle,label}$ between AuNP label positions and vesicle sites for fusing, adjacent, 40 nm shift, or label permutated sets. Mean and 95% envelopes are shown.

### Ext. Table 1 Cryo-ET data collection

| | 15F1 (C-term) | 11B8 (C-term) | 15F1 (H4K-2Cys) | 15F1 (H1-2Cys) | 5F11 (C-term) | 15F1+5F11 (C-term) | 15F1monomer +5F11dimer | Unlabeled |
|---|---|---|---|---|---|---|---|---|
| **Data collection** | | | | | | | | |
| Grids (no.) | 19 | 2 | 7 | 2 | 7 | 3 | 12 | 5 |
| Lamella (no.) | 188 | 31 | 161 | 32 | 99 | 34 | 249 | 71 |
| Tilt series (no.) | 1805 | 187 | 681 | 347 | 465 | 311 | 2339 | 1109 |
| Synapse-containing tomograms (no.) | 756 | 106 | 233 | 206 | 159 | 145 | 1010 | 136 |
| Final tomograms (no.) | 64 | 8 | 55 | 23 | 29 | 19 | 46 | 39 |
| Synaptic regions (no.) | 71 | 10 | 61 | 23 | 29 | 20 | 54 | 42 |
| Tissue-like (no.) | 39 | 0 | 36 | 4 | 1 | 5 | 36 | 20 |
| Semi tissue-like (no.) | 23 | 6 | 13 | 9 | 12 | 5 | 12 | 14 |
| Synaptosome-like (no.) | 9 | 4 | 12 | 10 | 16 | 10 | 6 | 8 |
| Magnification | 42-53k | | | | | | | |
| Voltage (kV) | 300 | | | | | | | |
| Electron exposure (e–/Å$^2$) | 120-150 | | | | | | | |
| Defocus range (μm) | -2.5 to -7 | | | | | | | |
| Pixel size (Å) | 1.9-2.3 | | | | | | | |

### Ext. Table 2 Cryo-ET subtomogram averaging data processing

| | H4K-2Cys 15F1 Fab-AuNP: AMPAR (ATD-focused) (EMD-72324) | H4K-2Cys 15F1 Fab-AuNP: AMPAR (LBD:TMD-focused) (EMD-72325) |
|---|---|---|
| **Data collection** | | |
| Magnification | 53k | |
| Voltage (kV) | 300 | |
| Electron exposure (e–/Å$^2$) | 120 | |
| Defocus range (μm) | -3 to -5 | |
| Pixel size (Å) | 2.4 | |
| **Processing** | | |
| Symmetry imposed | C2 | C2 |
| Initial particle images (no.) | 1,254 | 1,254 |
| Final particle images (no.) | 1,254 | 605 |
| Map resolution (Å) | 14.0 (Relion) | 22.0 (Relion) |
| FSC threshold | 0.143 | 0.143 |
| Resolution range (Å) | 14.0-40 (RESOLVE) | 24.0-37.0 (RESOLVE) |
