## Supplementary Information Guide for "Mapping AMPA and NMDA receptors at mouse brain hippocampal synapses by cryo-ET"

**Supplementary Data**
This file contains Supplementary Figs.1-11 and Supplementary Table 1.

**Supplementary File 1**
This file contains denoised tomogram slice, membrane segmentation, and cleft minimum intensity projection (MIP) examples from all datasets.

**Supplementary Video 1: AuNP labels reveal AMPA receptor distribution in mouse hippocampal synapses.** This video shows the denoised density of a representative tomogram of a synapse labeled with the 15F1 "C-term" Fab-AuNP, also depicted in Figs. 1–4, first as slices through the density and then as a tilting projection of the tomogram. After superimposing the membrane annotations, a minimum-intensity projection of the density within the largest synaptic cleft illustrates clustering of AMPA receptors. Annotation of AuNP distances from the postsynaptic membrane reveals potential noncanonical receptor states. A minimum-intensity projection of the density within a second synaptic cleft then illustrates how realignment of the tilt series guided by the AuNP densities improves interpretability of AuNPs that define AMPA receptor locations. Finally, a minimum-intensity projection outside a synaptic cleft shows AuNPs associated with extrasynaptic AMPA receptors.

**Supplementary Video 2: Rigidly attached AuNP labels reveal multiple poses of AMPAR-AMPAR interactions.** This video shows the denoised density of a representative tomogram of a synapse labeled with the 15F1 "H4K-2Cys" Fab-AuNP, also depicted in Fig. 5, first as slices through the density and then as a projection of the tomogram. After annotation of membranes, a minimum intensity projection of the density within the synaptic cleft reveals AuNPs that can be assigned to pairs corresponding to single receptors. The video then illustrates how the position of the AuNP pairs and membrane can be used to estimate AMPA receptor position and orientation within the synaptic cleft. This reveals how AMPA receptors are directly adjacent to one another in multiple distinct poses.

**Supplementary Video 3: Multiplexed AuNP labels reveal how AMPA and NMDA receptors intermingle.** This video shows the denoised density of a representative tomogram of a synapse dual labeled with 15F1 Fab-$AuNP_{monomer}$ and 5F11 Fab-$AuNP_{dimer}$, one of the examples depicted in Fig. 6, first as slices through the density and then as a projection of the tomogram. After annotation of membranes, a minimum intensity projection of the density within the synaptic cleft shows monomeric and dimeric AuNPs that can be clearly distinguished and reveal intermingling of AMPA and NMDA receptors.

**Supplementary Video 4: Multiplexed AuNP labels defines the proximity of AMPA and NMDA receptors to vesicle fusion events.** This video shows the denoised density of a representative tomogram of a synapse dual labeled with 15F1 Fab-$AuNP_{monomer}$ and 5F11 Fab-$AuNP_{dimer}$, one of the examples depicted in Fig. 7, first as slices through the density that reveal vesicle fusion events and then as a projection of the tomogram. After annotation of membranes, fusion events and membrane adjacent vesicles are highlighted and overlaid on a minimum intensity projection of the density within the synaptic cleft. Monomeric and dimeric AuNPs can be distinguished and reveal localization of AMPA and NMDA receptors relative to vesicle fusion events.
