## Supplementary Data for "Mapping AMPA and NMDA receptors at mouse brain hippocampal synapses by cryo-ET"

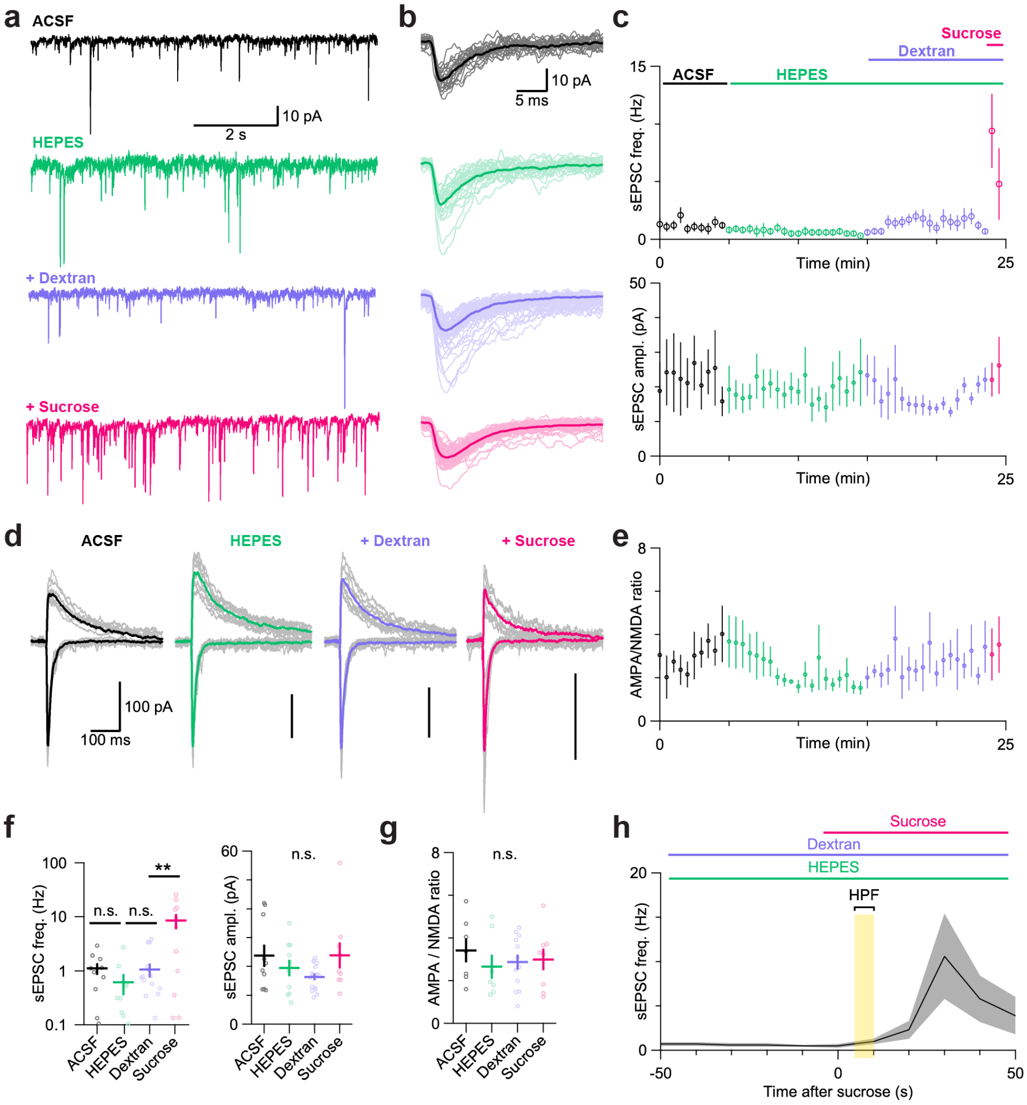


**Supplementary Fig. 1 Electrophysiological characterization of hippocampal tissue slices in cryo-ET sample preparation conditions. a,** Electrophysiological recordings of sEPSCs from CA1 neurons in various extracellular solutions. **b**, Overlay traces of sEPSC events in each condition with averaged traces shown in bold. **c**, Time course of frequency and amplitude of sEPSCs during the application of HEPES solution with dextran and sucrose cryoprotectants. **d**, Electrophysiological recordings of evoked AMPA and NMDA receptor currents from CA1 neurons in various extracellular solutions. Averaged traces are shown in bold color. **e**, Time course of AMPA/NMDA ratio throughout the indicated solution exchanges. **f**, Summary plot of frequency and amplitude of sEPSCs in each solution. **g**, Summary plot of AMPA/NMDA ratio in each solution. **h**, Expanded time course from panel **c**, highlighting the window in which high pressure freezing (HPF) of tissue slices occurs during cryo-ET tissue sample preparation.

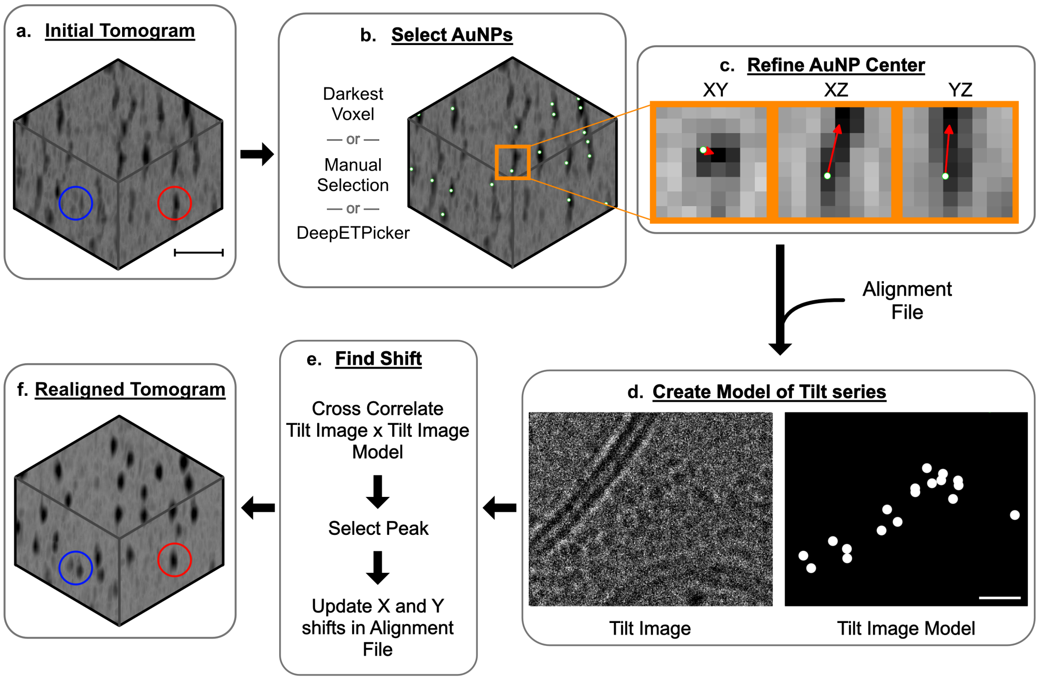


**Supplementary Fig. 2 AuNP-guided tomogram re-alignment improves local resolution for confident AuNP identification. a,** An initial denoised subvolume from representative anti-GluA2 15F1-AuNP$_{monomer}$ and anti-GluN1 5F11-AuNP$_{dimer}$ dual labeled tomogram (scale bar = 14 nm). **b,** AuNP centers (denoted as white spheres) are selected. **c,** AuNP centers are refined by 3D Gaussian fitting (red arrows indicate translation to new center). **d,** Tomogram AuNP positions are mapped back to the tilt series and a model tilt image of the AuNPs is generated (scale bar = 12.5 nm). **e,** The x and y shift between each tilt image and its corresponding alignment is found by cross correlation of the model and tilt images, followed by 2D Gaussian fitting of the cross-correlation peak and update of alignment file shifts. **f,** The same denoised subvolume as in panel **a** after AuNP-guided realignment. The red and blue circles indicate example monomeric and dimeric AuNPs, respectively, that are more clearly identifiable after realignment.

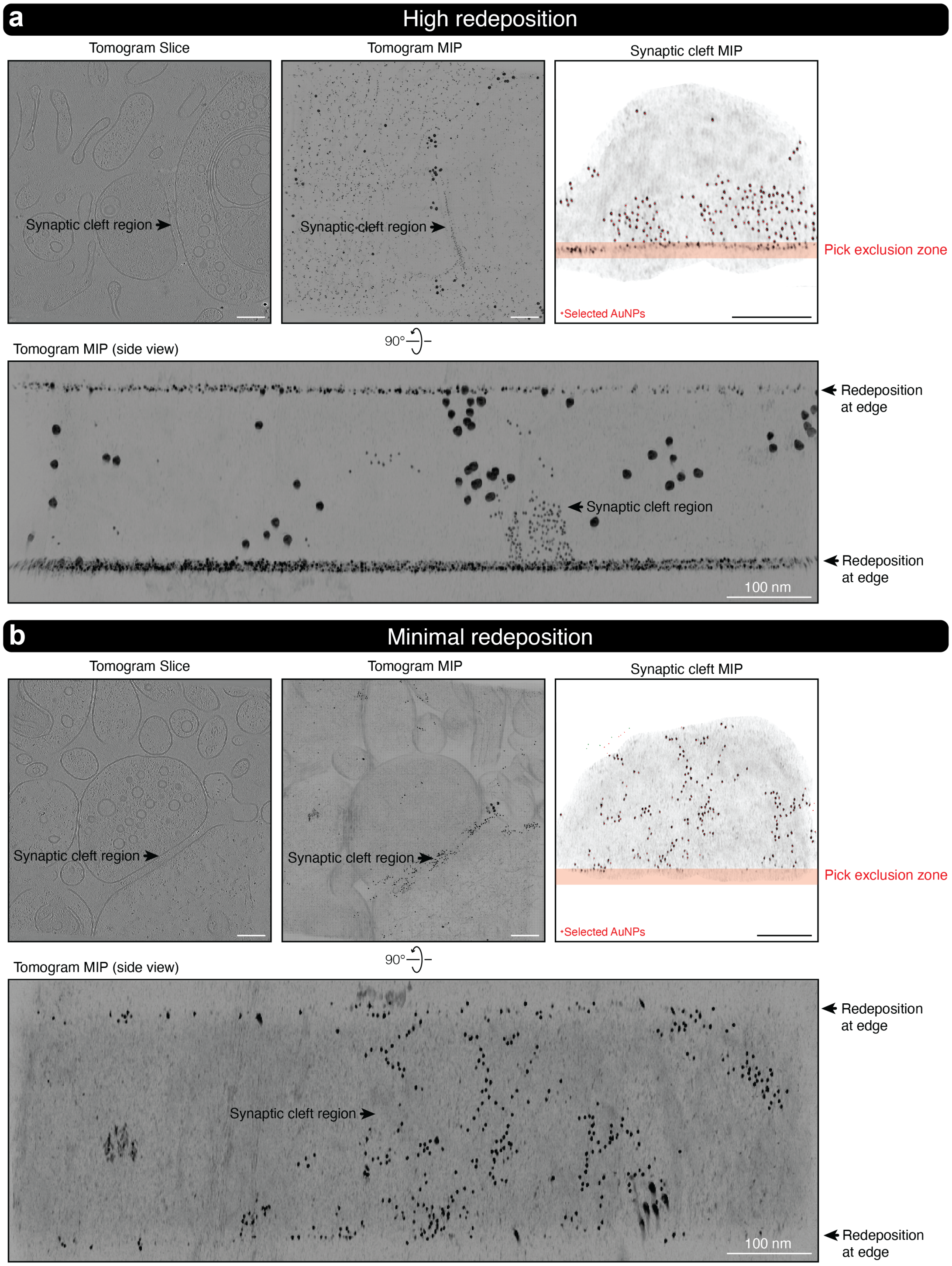


**Supplementary Fig. 3 FIB-mill redeposition on lamella surfaces is excluded from AuNP picking.** Tomogram slices and minimum intensity projections (MIP) from representative tomogram examples with high (**a**) or minimal (**b**) FIB-mill redeposition on lamella surface. To avoid picking gallium redeposition, which resembles AuNPs, we excluded any AuNP picks within ~6 nm of tomogram edges. All scale bars are 100 nm.

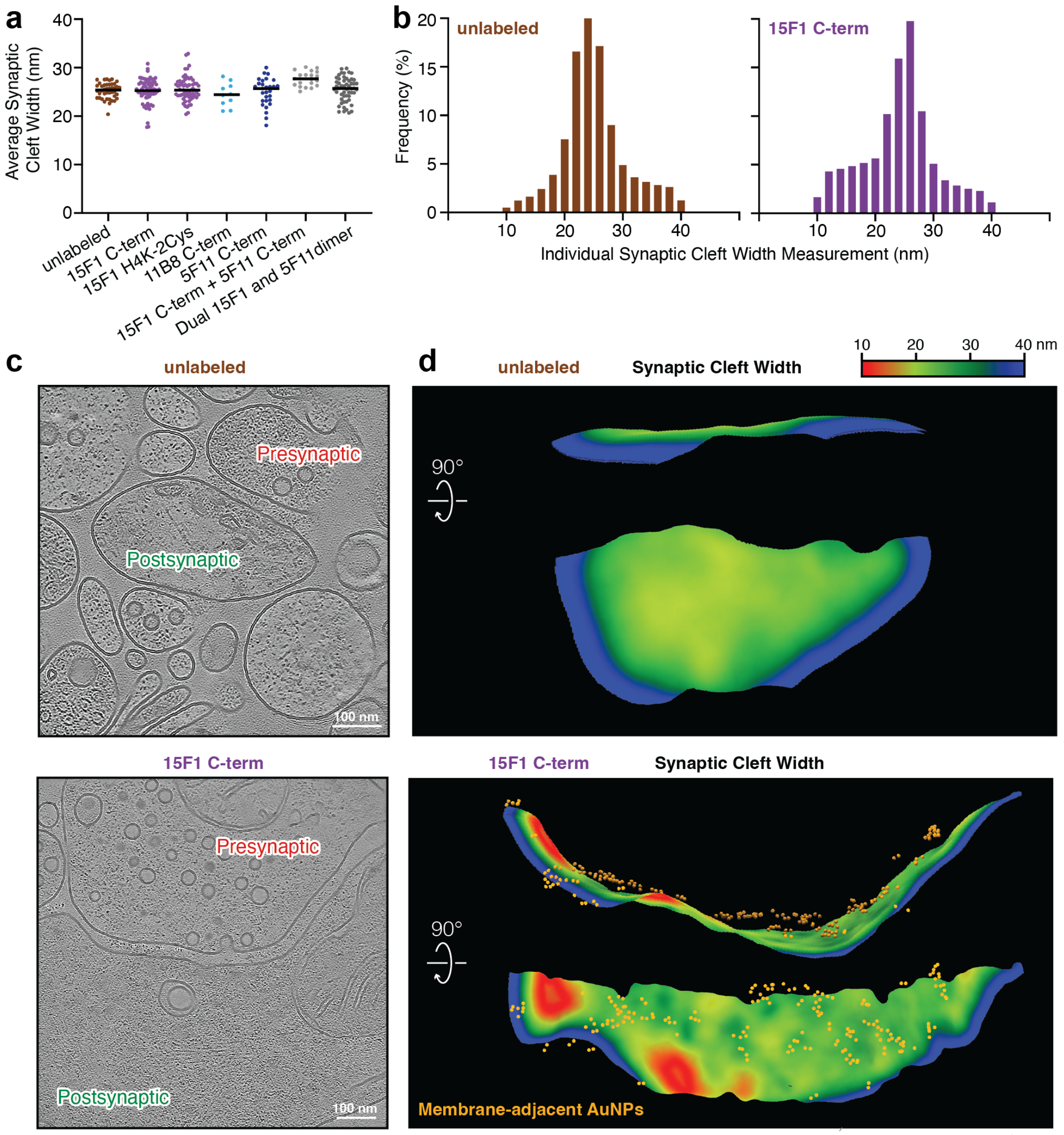


**Supplementary Fig. 4 Fab-AuNP labeling does not alter synaptic cleft width. a**, Average width of the synaptic cleft across Fab-AuNP labeling conditions, per synaptic region, calculated from cleft membrane segmentation edge-to-edge distances. **b**, Frequency histograms of synaptic cleft width measurements across entire cleft region in unlabeled and 15F1 "C-term" datasets (bin size 10 nm). **c**, Denoised tomogram slices from example unlabeled and 15F1 "C-term" Fab-AuNP labeled synaptic tomograms. **d**, Heat maps of synaptic cleft widths mapped onto postsynaptic membrane segmentations from tomograms in panel **c**.

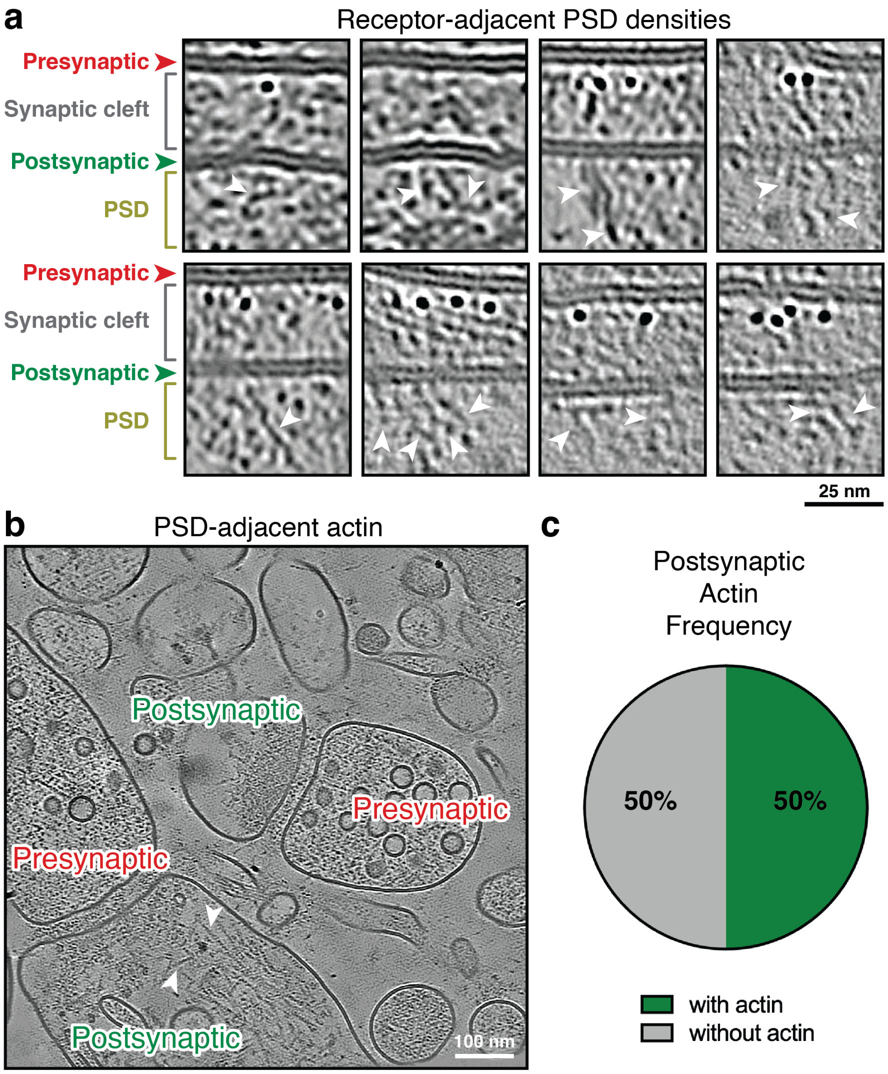


**Supplementary Fig. 5 Receptors and actin localize near postsynaptic density features. a**, Example denoised tomogram slices with flexible, filament-like postsynaptic density (PSD) features (white arrows) near AuNP-labeled AMPA or NMDA receptors. **b**, Example tomogram slice with actin features (white arrows) in postsynaptic compartment adjacent to PSD. **c**, Frequency of actin identification in postsynaptic compartments in 15F1 “C-term” and “H4K-2Cys” datasets (n=128 tomograms).

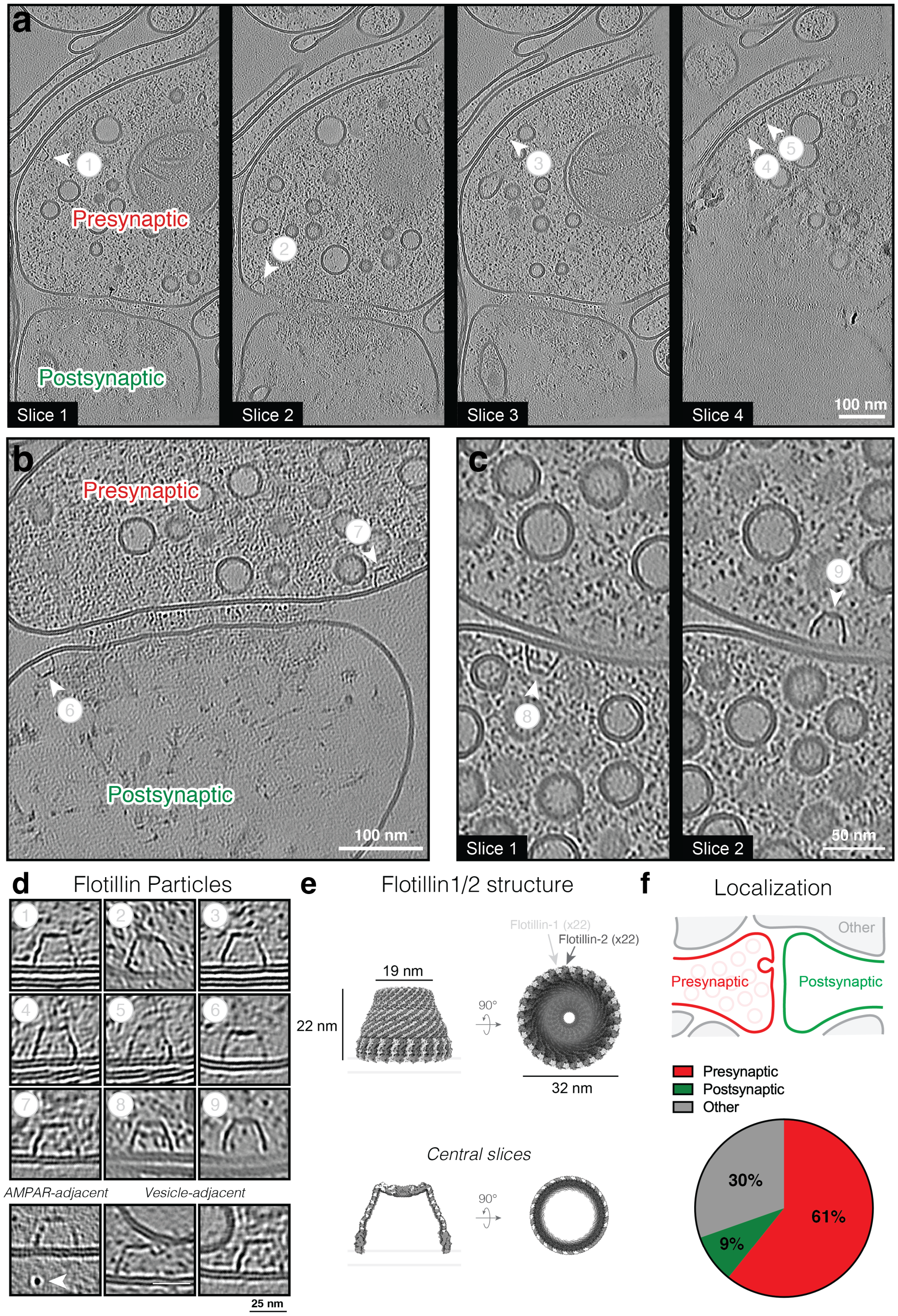


**Supplementary Fig. 6 Flotillin cages are observed adjacent to synapses.** Denoised tomogram slices from example synapses containing five flotillin cages (white arrows) in the presynaptic compartment (**a**), synaptic cleft-adjacent flotillin cages in both the presynaptic and postsynaptic compartments (**b**), or two flotillin cages in neighboring presynaptic compartments (**c**). **d**, Magnified slice views of the nine flotillin cages from panels **a**-**c** (top). Additional examples of flotillin cages adjacent to 15F1 "C-term" AuNP labels or intracellular vesicles (bottom). **e**, Molecular structure of flotillin1/2 complex and central slices (PDBID 9BQ2). **f**, Localization of flotillin-like particles in 15F1 "C-term" and "H4K-2Cys" labeled datasets (n=79 flotillin-like particles; 128 tomograms).

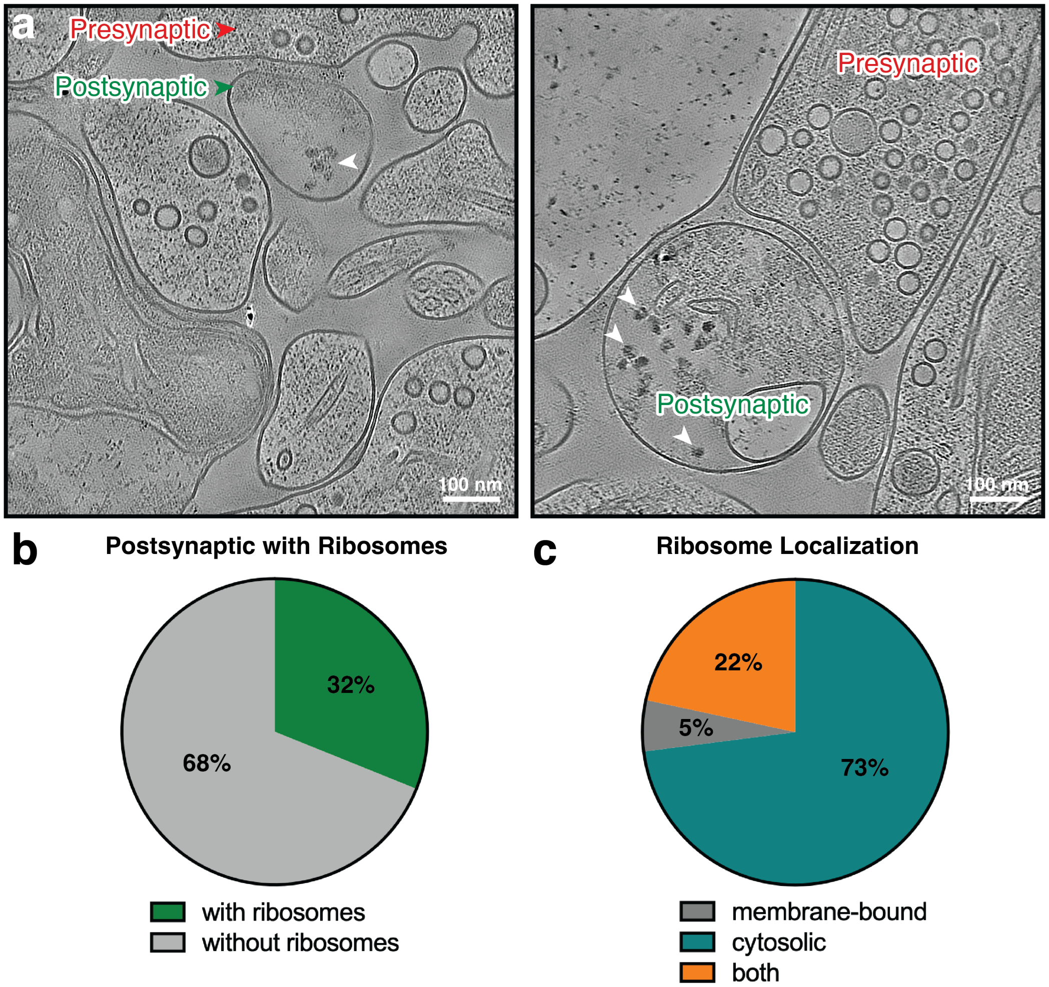


**Supplementary Fig. 7 Ribosomes are predominantly observed in postsynaptic compartments. a,** Example ribosome-like densities (white arrows) in denoised tomogram slices from two synapses. **b**, Frequency of ribosome-like particle identification in postsynaptic compartments in 15F1 "C-term" and "H4K-2Cys" Fab-AuNP labeled datasets (n=128 tomograms). **c**, Frequency of observing membrane-bound ribosomes, cytosolic ribosomes, or both, in the "with ribosomes" subset from panel **b**.

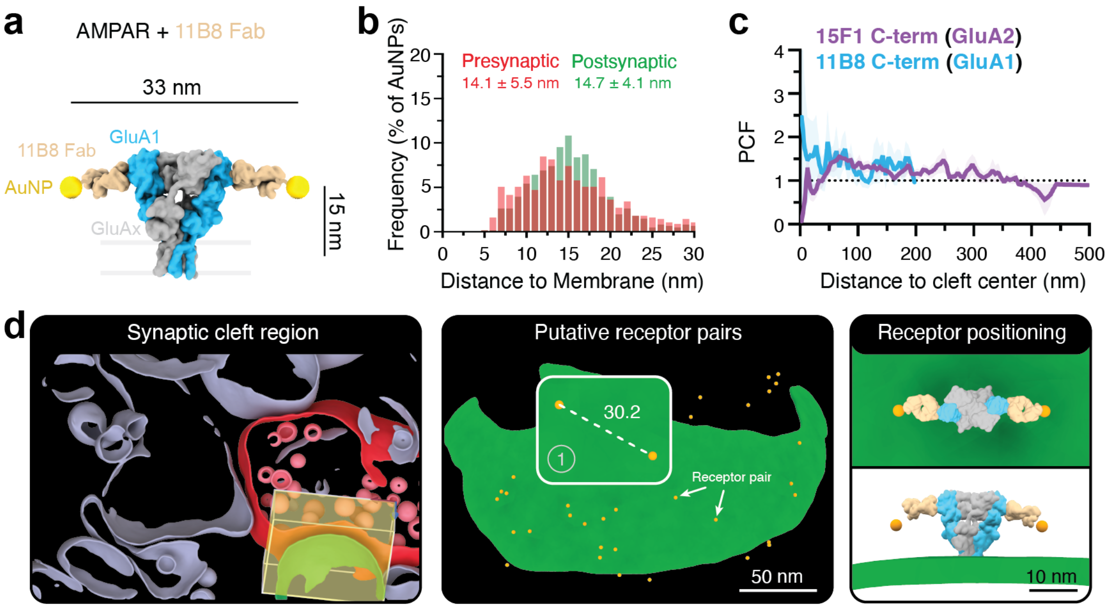


**Supplementary Fig. 8 GluA1-containing AMPA receptors are identified throughout the synaptic cleft.** **a**, Molecular model of di-heteromeric GluA2/GluA1 AMPA receptor bound to 11B8 “C-term” Fab-AuNP cryo-ET label (based on PDBID 7LDD). **b**, Histogram of AuNP-membrane distances in 11B8 “C-term” Fab-AuNP labeled data (n=610 AuNPs). **c**, Pair correlation function (PCF) between synaptic cleft center and AuNP positions (bin size = 5). Mean and standard error of the mean are shown (n=9 11B8-labeled synaptic regions), overlaid with equivalent function for 15F1 “C-term” Fab-AuNP labeled synaptic regions. **d**, Example segmentation of synapse labeled with 11B8 “C-term” Fab-AuNP (left). View normal to postsynaptic membrane (middle) with a putative receptor pair, along with top and side views of AuNP pair-guided receptor docking (right). The 3D distance between the putative receptor AuNP pair is noted in nm.

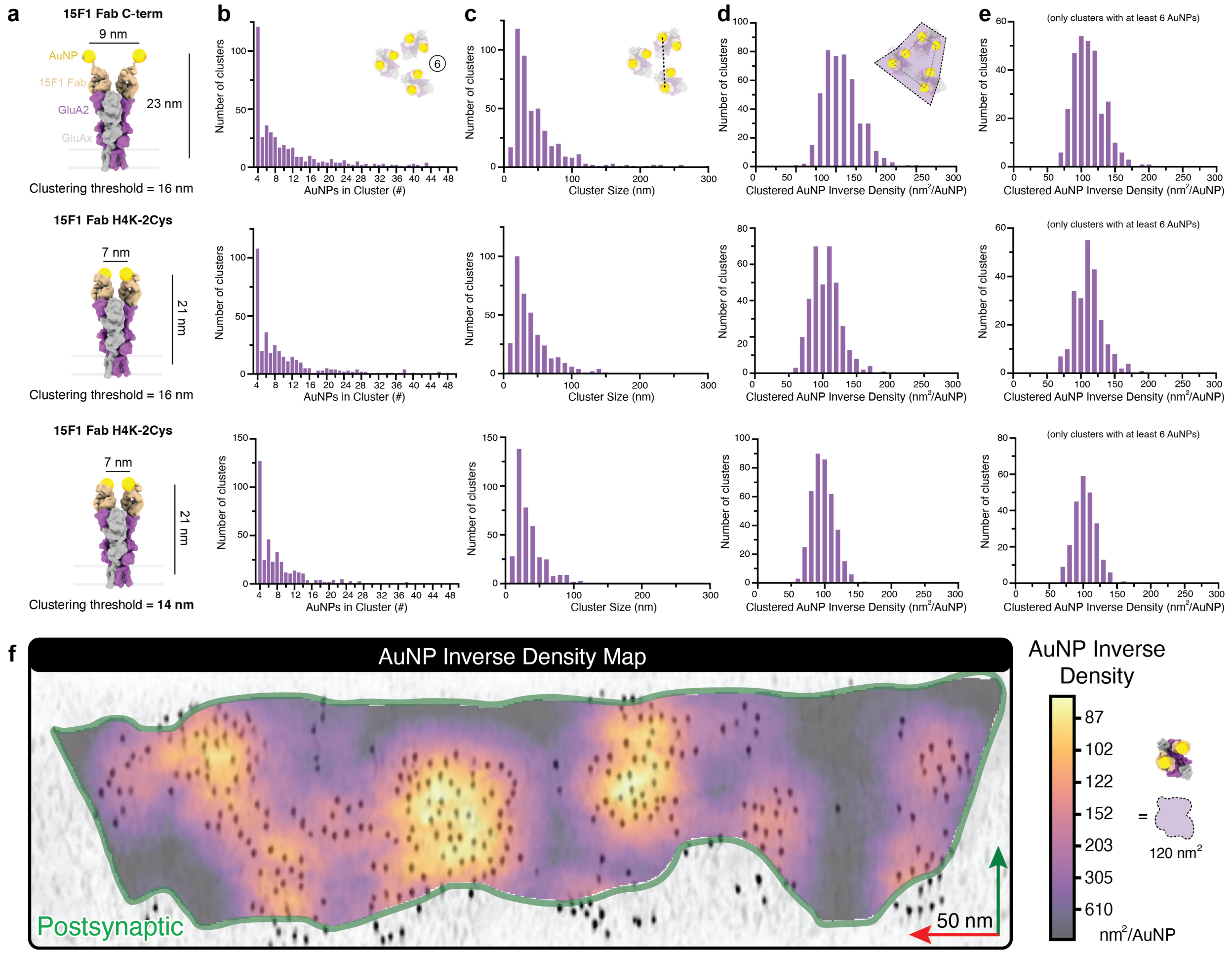


**Supplementary Fig. 9 GluA2 labeling with 15F1 Fab "C-term" and "H4K-2Cys" conjugates provides reproducible clustering metrics.** **a**, Molecular models of AMPA receptors bound to GluA2-specific 15F1 Fab "C-term" or "H4K-2Cys" labels. Distance thresholds used for the clustering metrics reported in each row are indicated. **b**, Histogram of the number of AuNPs per cluster (bin size 1), cropped to 0-50 x axis range. **c**, Histogram of AuNP cluster sizes (bin size 10). **d**, Histogram of clustered AuNP inverse density (bin size 10). **e**, Same as panel **d**, but only including clusters with at least six AuNPs that result in more accurate cluster area estimates. For panels **b-e**, values correspond to the associated Fab-AuNP label ("C-term" or "H4K-2Cys") and distance threshold (16 or 14 nm) indicated in the same row from panel **a**. **f**, AuNP inverse density map for 15F1 "C-term" Fab-AuNP labeled synapse from Fig.2 panels **d-e**, overlaid on synaptic cleft MIP (see Methods). Extent of packing, relative to the 120 nm$^2$ cross-sectional area of an AMPA receptor, can be estimated by doubling the AuNP inverse density (assuming 2:1 AuNP: AMPAR labeling).

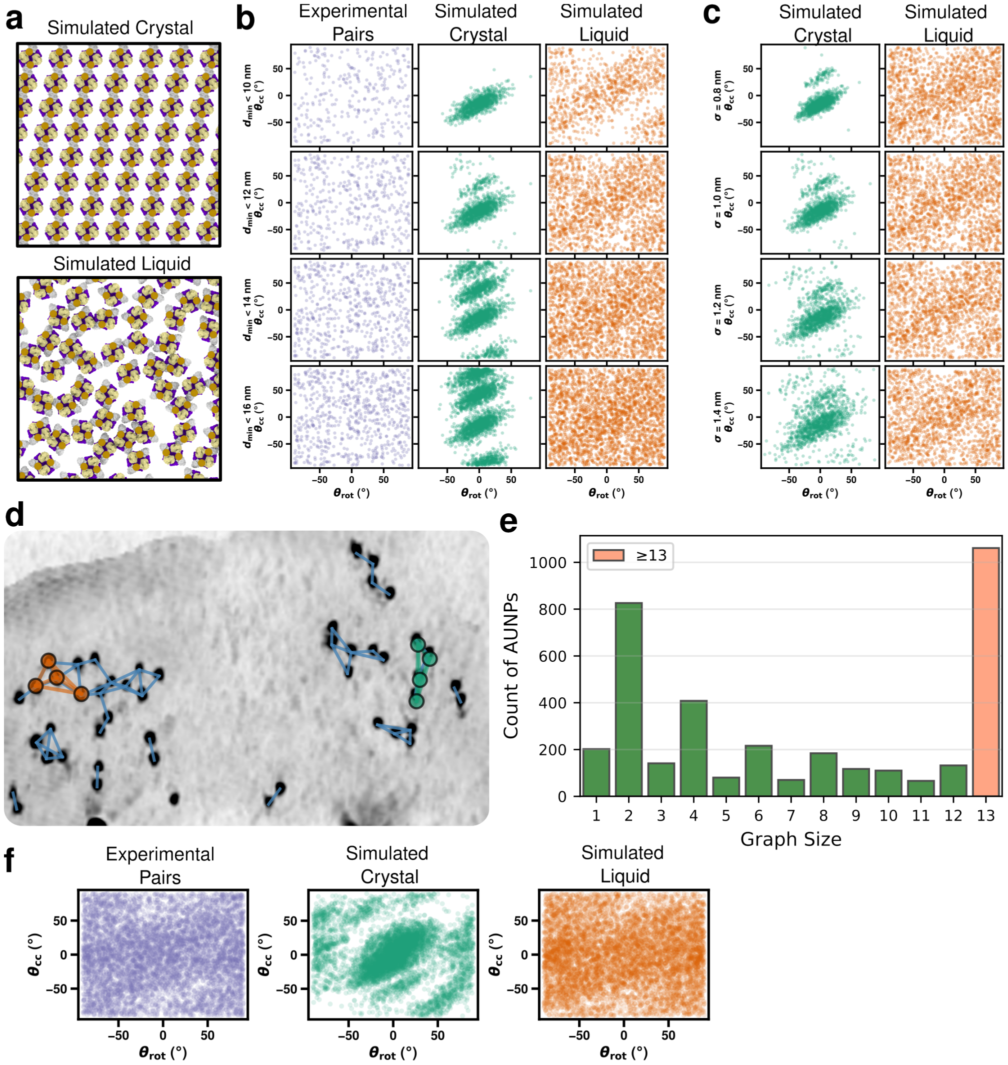


**Supplementary Fig. 10 Pairwise geometrical analysis reveals liquid-like AMPAR packing. a,** Renderings of a simulated 2D-crystalline lattice (top) and liquid arrangement (bottom) of 15F1 “H4K-2Cys” Fab-AuNP labeled AMPA receptors used as reference packing models. Box dimensions are 100 nm. **b,** Scatter plots of $\theta_{rot}$ vs. $\theta_{cc}$ for experimental receptor pairs (left), simulated crystal (middle), and simulated liquid (right) models, using increasing distance cutoffs ($d_{min}$ < 10, 12, 14, and 16 nm) to define a neighboring pair. Across all cutoffs, the experimental distribution resembles the simulated liquid more than the simulated crystal. **c,** As in panel **b**, adding increasing positional noise (σ = 0.8, 1.0, 1.2, and 1.4 nm) and confirming that the two models remain distinguishable across this range. **d,** Representative cleft MIP with detected AuNP positions and graph edges (blue) connecting AuNPs within the distance cutoff, illustrating the alternative graph-based approach for defining candidate receptor-pairs (4-sized subgraphs). Orange and teal highlight two example 4-sized subgraphs. **e,** Histogram of graph sizes across the 15F1 “H4K-2Cys” dataset. Graph sizes are predominantly even numbers, due to 2:1 AuNP: AMPAR labeling. **f,** As in Fig. 5h-j, scatter plots of $\theta_{rot}$ vs. $\theta_{cc}$ for experimental AuNP pairs (left), simulated crystal (middle), and simulated liquid (right), using all potential 4-sized subgraphs in the experimental dataset or packing simulations (rather than only those corresponding to assigned receptor poses for STA).

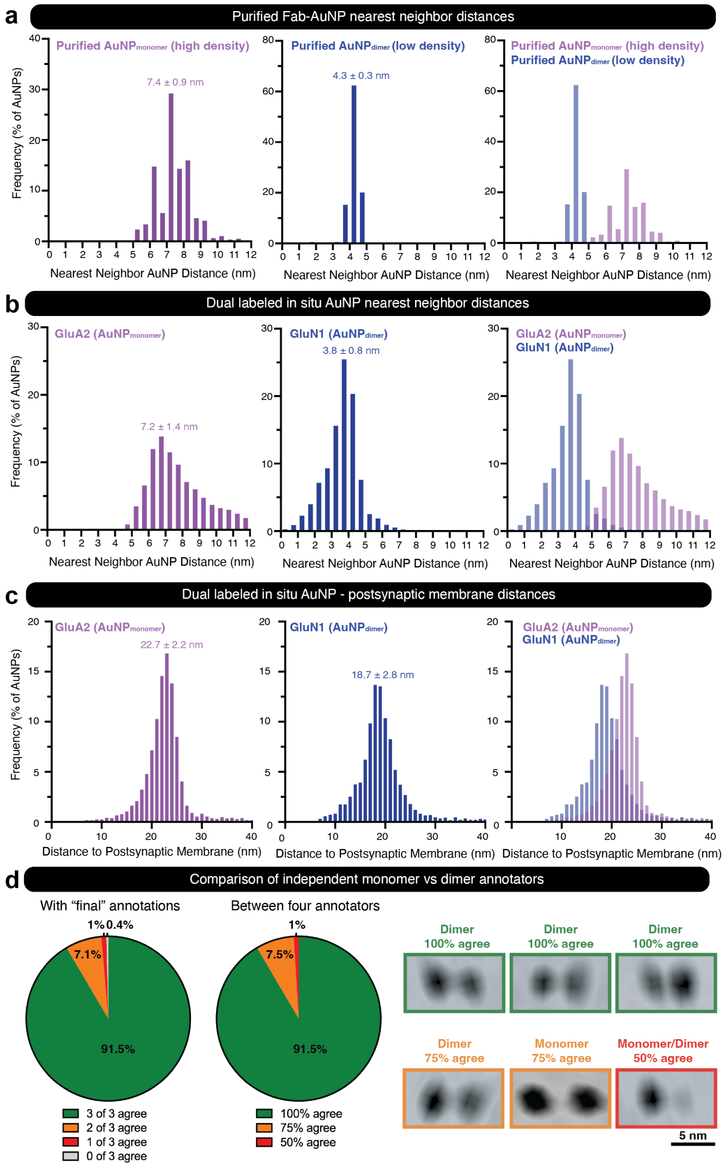


**Supplementary Fig. 11 AuNP monomer and dimer classifications are validated by multiple metrics. a,** Frequency histograms (bin size 0.5 nm) of AuNP nearest neighbor distances for purified 15F1 “C-term” Fab-AuNP$_{monomer}$ plunge frozen at high density or 5F11 “C-term” Fab-AuNP$_{dimer}$ plunge frozen at low density, plotted independently (left, middle) and overlaid (right). **b,** Frequency histograms (bin size 0.5 nm) of AuNP nearest neighbor distances for dual labeled 15F1 Fab-AuNP$_{monomer}$ and 5F11 Fab-AuNP$_{dimer}$ labeled FIB-milled tissue samples, plotted independently (left, middle) and overlaid (right). **c,** Frequency histograms (bin size 1 nm) of postsynaptic membrane to AuNP distances for dual labeled 15F1 Fab-AuNP$_{monomer}$ and 5F11 Fab-AuNP$_{dimer}$ (includes distances to both AuNPs within the dimers) labeled FIB-milled tissue samples, plotted independently (left, middle) and overlaid (right). **d**, Assessment of consistency between independent manual assignments of AuNP monomers and dimers in representative subset of five dual labeled tomograms. Comparison of three annotators with the “final” classifications (left), comparison of all four manual annotators (middle), and example particles that fall into the classes designated in the latter.

### Supplementary Table 1. Statistical comparisons for Ripley's H analyses

| p-values* | 0-100 nm | 0-200 nm | 0-500 nm |
|---|---|---|---|
| **Cleft distribution (cleft center, AuNP label)** | | | |
| *15F1 (GluA2)* | | | |
| *vs 5F11 (GluN1)* | 0.045 | 0.227 | 0.314 |
| *Dual 15F1 monomer (GluA2)* | | | |
| *vs dual 5F11 dimer (GluN1)* | 0.793 | 0.731 | 0.611 |
| **Fusion events (vesicle, AuNP label)** | | | |
| *15F1 (GluA2)* | | | |
| *Fusion events* | | | |
| *vs adjacent vesicles* | 0.131 | 0.061 | 0.051 |
| *vs 40 nm shift* | 0.129 | 0.079 | 0.069 |
| *vs label permutation* | 0.109 | 0.040 | 0.050 |
| *Dual 15F1 monomer (GluA2)* | | | |
| *Fusion events* | | | |
| *vs adjacent vesicles* | 0.710 | 0.529 | 0.841 |
| *vs 40 nm shift* | 0.475 | 0.436 | 0.485 |
| *vs label permutation* | 0.050 | 0.069 | 0.178 |
| *Dual 5F11 dimer (GluN1)* | | | |
| *Fusion events* | | | |
| *vs adjacent vesicles* | 0.086 | 0.065 | 0.127 |
| *vs 40 nm shift* | 0.257 | 0.257 | 0.446 |
| *vs label permutation* | 0.069 | 0.337 | 0.376 |

*Reported p-values are from permutation and Monte Carlo tests of the integrated squared difference of the mean Ripley's H curves over the indicated distance ranges (see Methods).
