## Supplementary File 1 for "Mapping AMPA and NMDA receptors at mouse brain hippocampal synapses by cryo-ET"

| Tomogram ID: 20231017_EGmilled24-2_68 | SET: anti-GluA2 15F1 "C-term" |
| --- | --- |
| Cleft ID: 0 | Type: tissue |

### Denoised Tomogram Slice

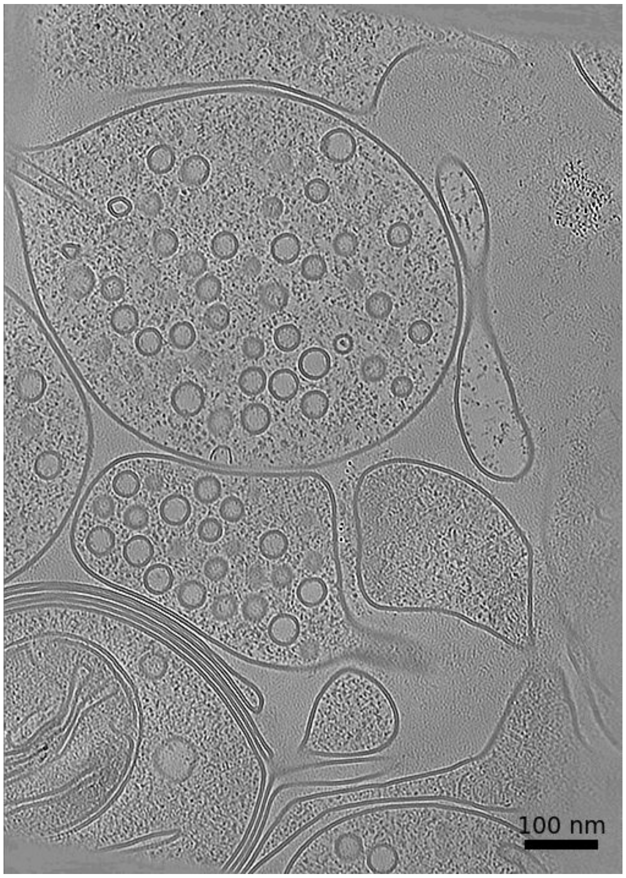


### Membrane Segmentation & Cleft Position

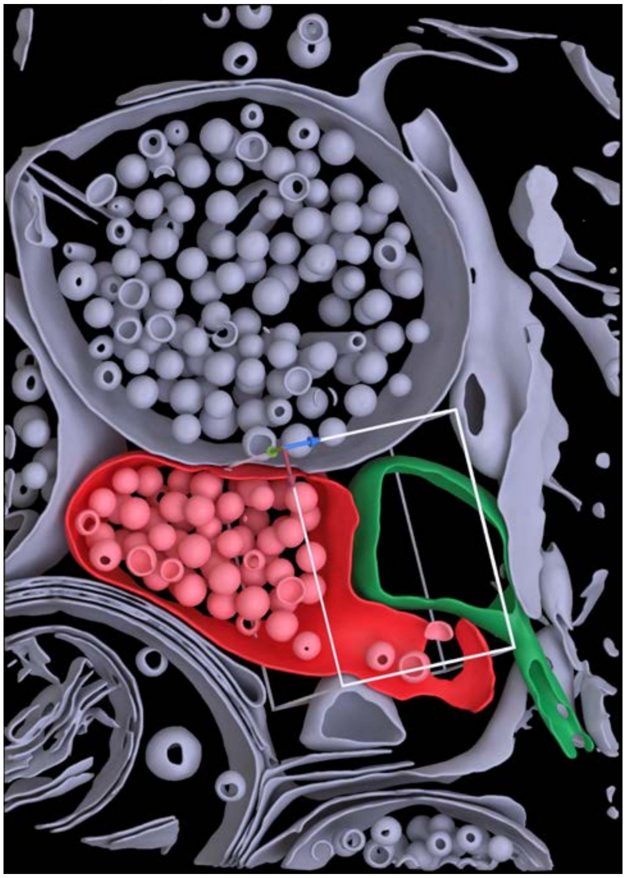

### Cleft MIP

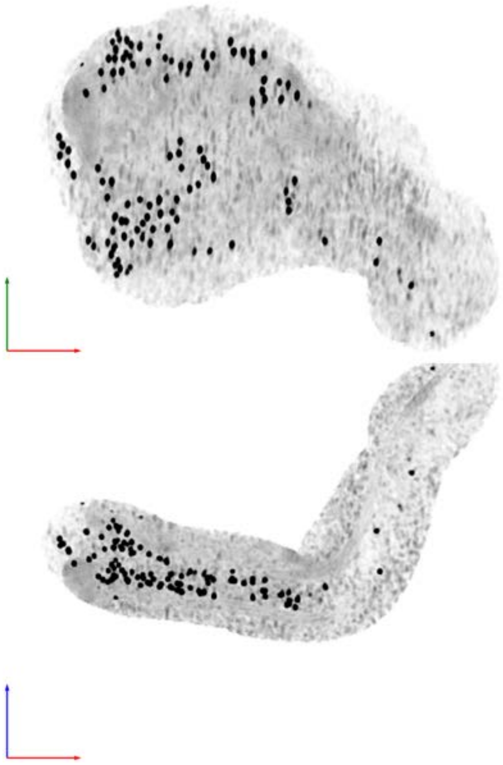

| Tomogram ID: 20231017_HippAu_159 | SET: anti-GluA2 15F1 "C-term" |
|---|---|
| Cleft ID: 0 | Type: tissue |

**Denoised Tomogram Slice**

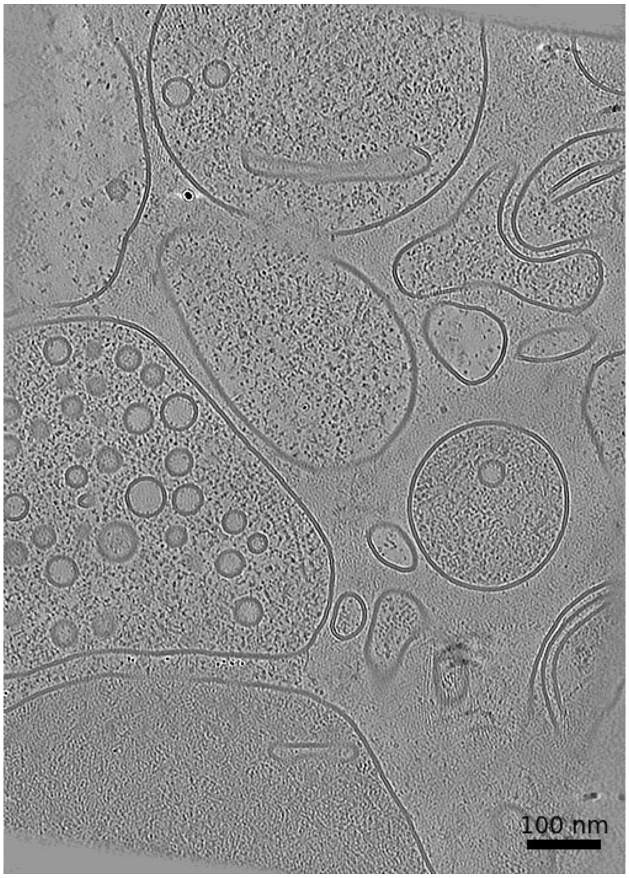


**Membrane Segmentation & Cleft Position**

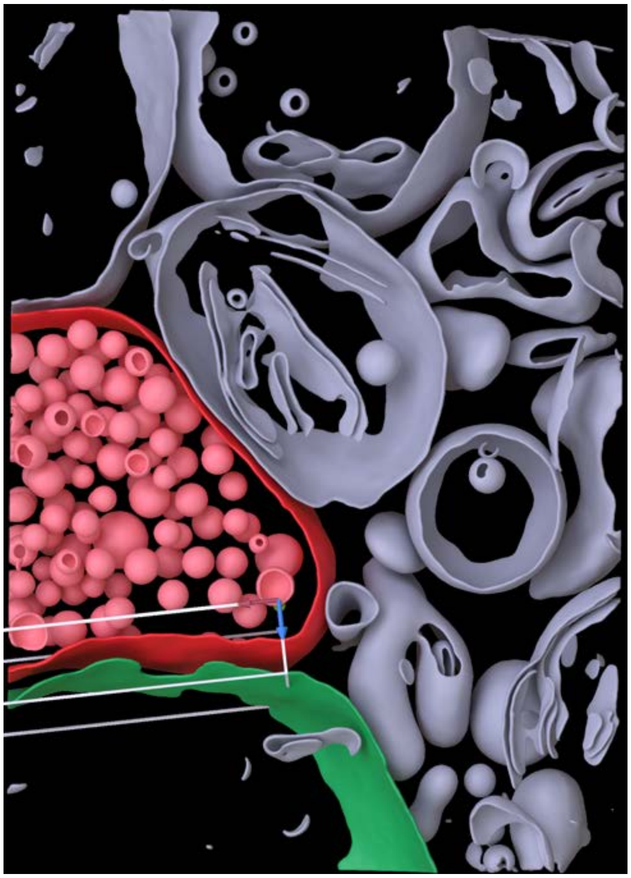

**Cleft MIP**

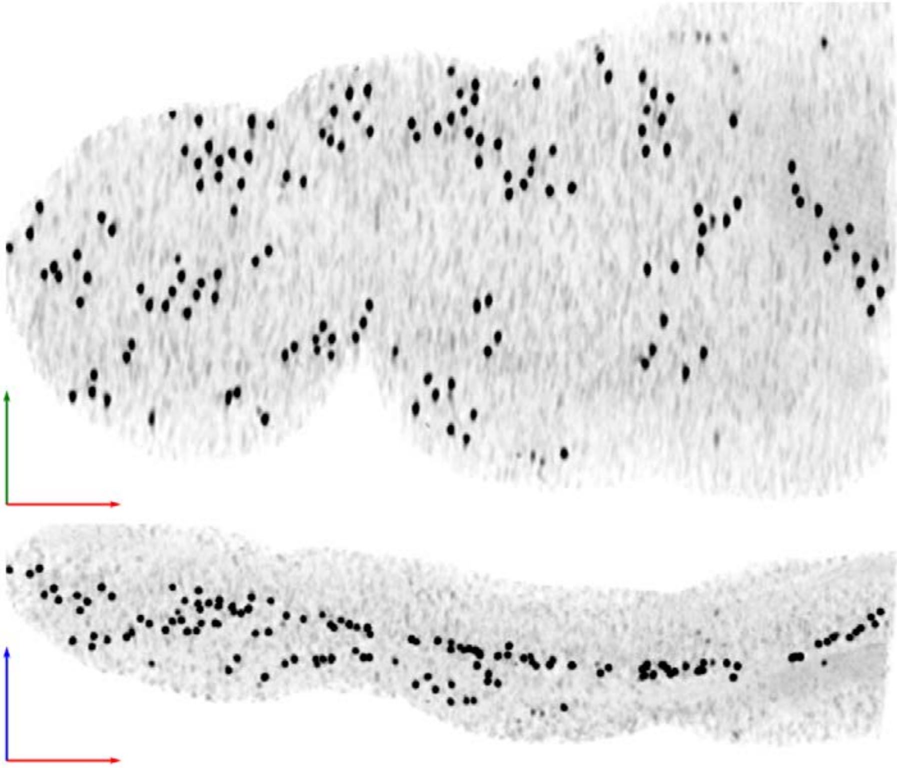

| Tomogram ID: 20231026_HippAu_26 | SET: anti-GluA2 15F1 "C-term" |
|---|---|
| Cleft ID: 0 | Type: tissue |

### Denoised Tomogram Slice

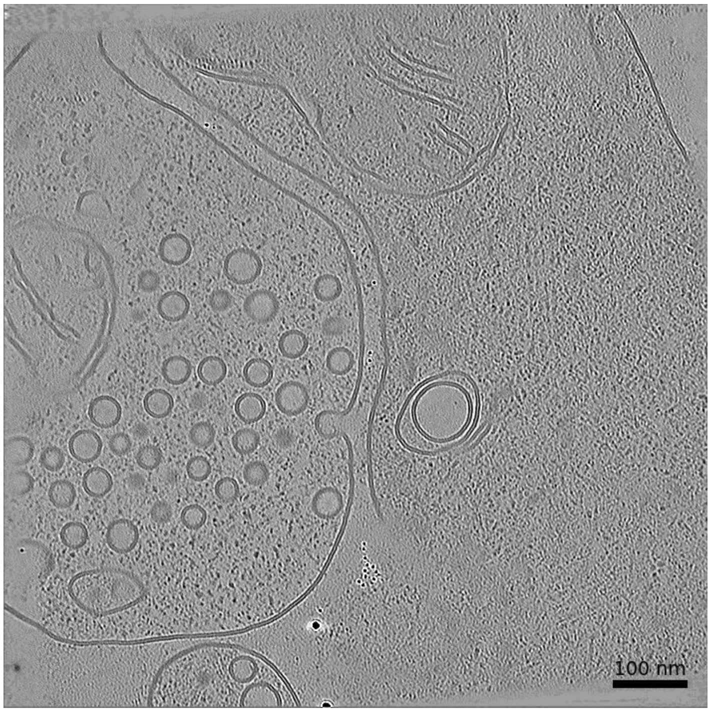


### Membrane Segmentation & Cleft Position

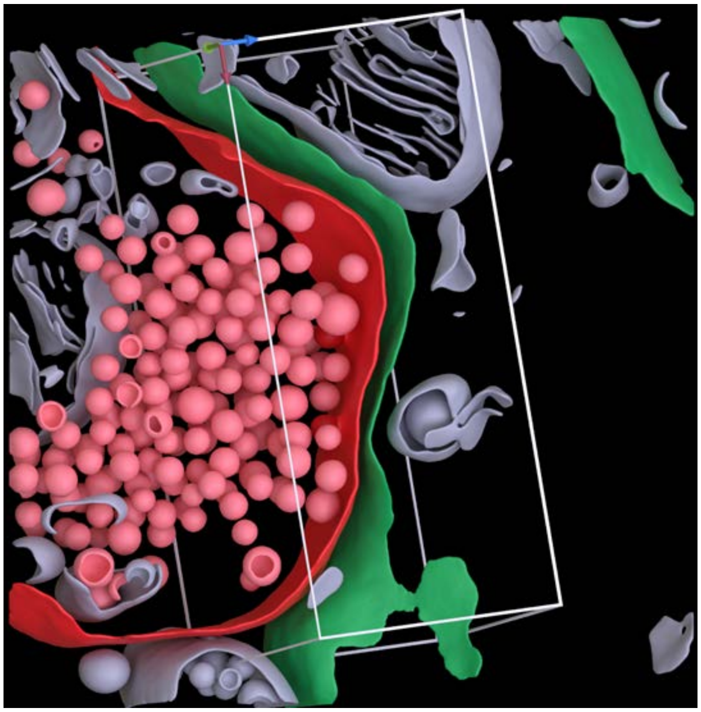

### Cleft MIP

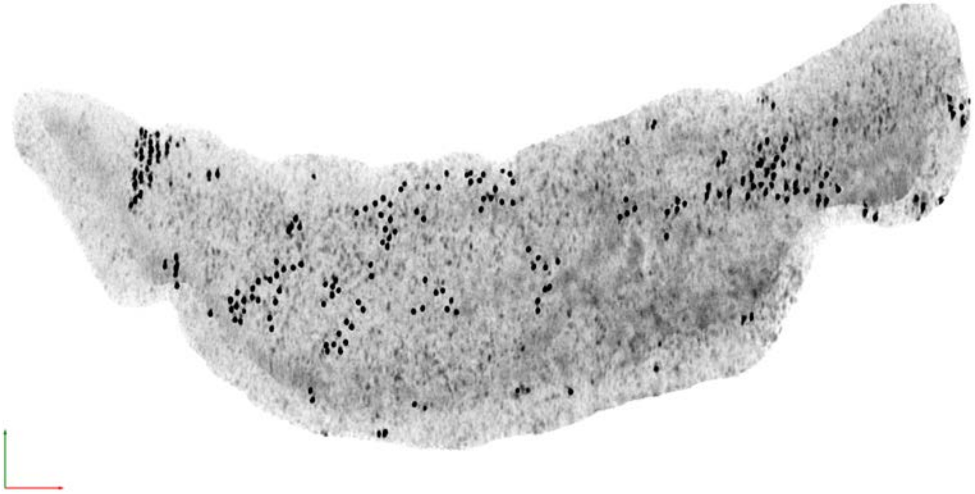

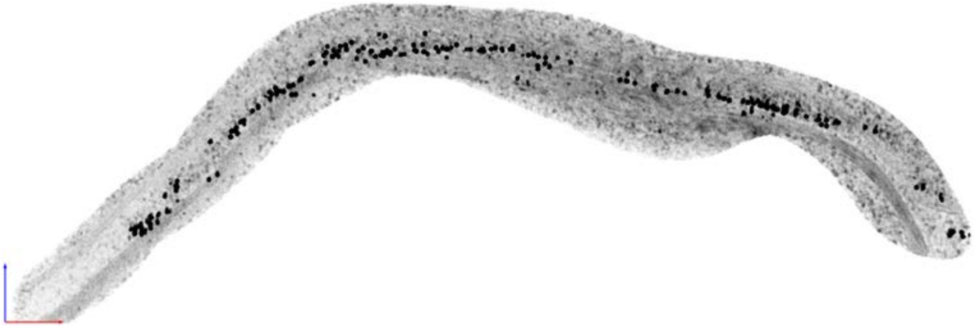

**Tomogram ID: 20240111_WaffleHipp_116** **SET: anti-GluA2 15F1 "C-term"**

Cleft ID: 0 Type: tissue

**Denoised Tomogram Slice**

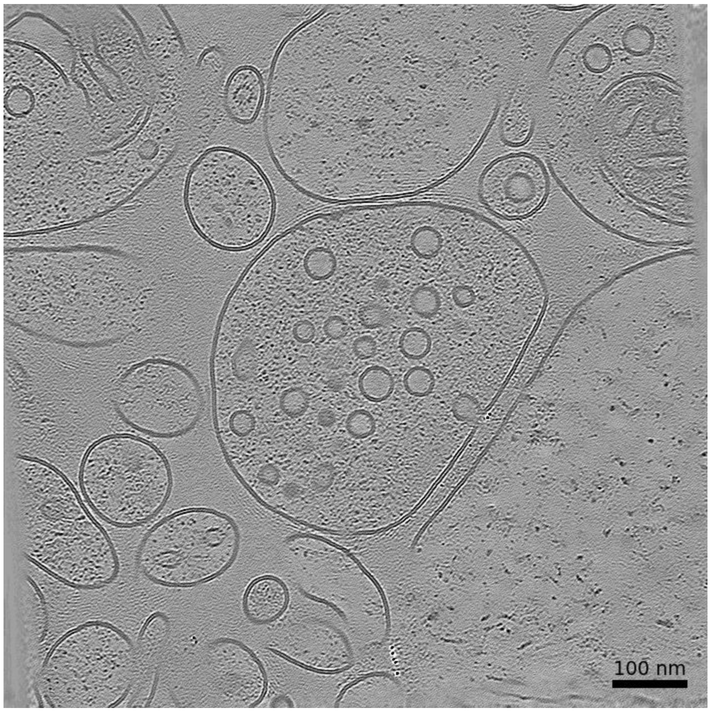


**Membrane Segmentation & Cleft Position**

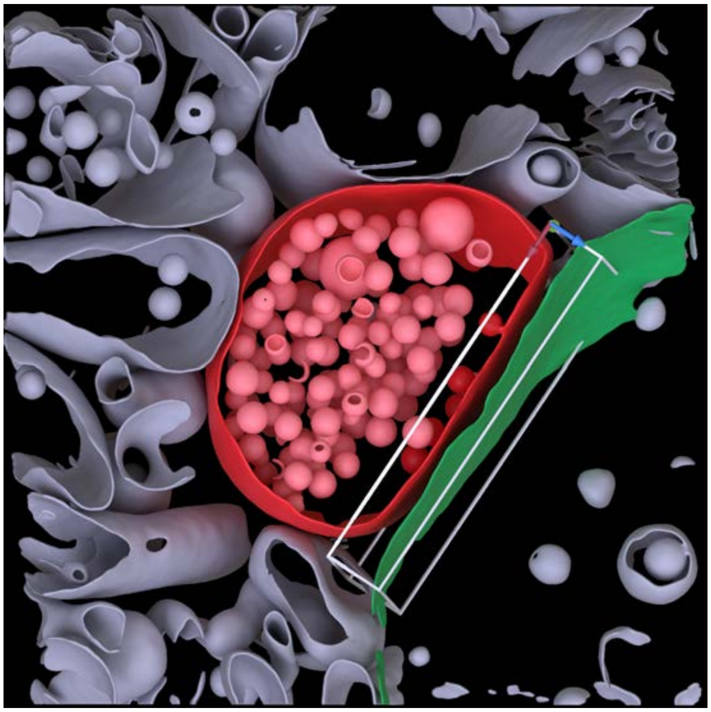

**Cleft MIP**

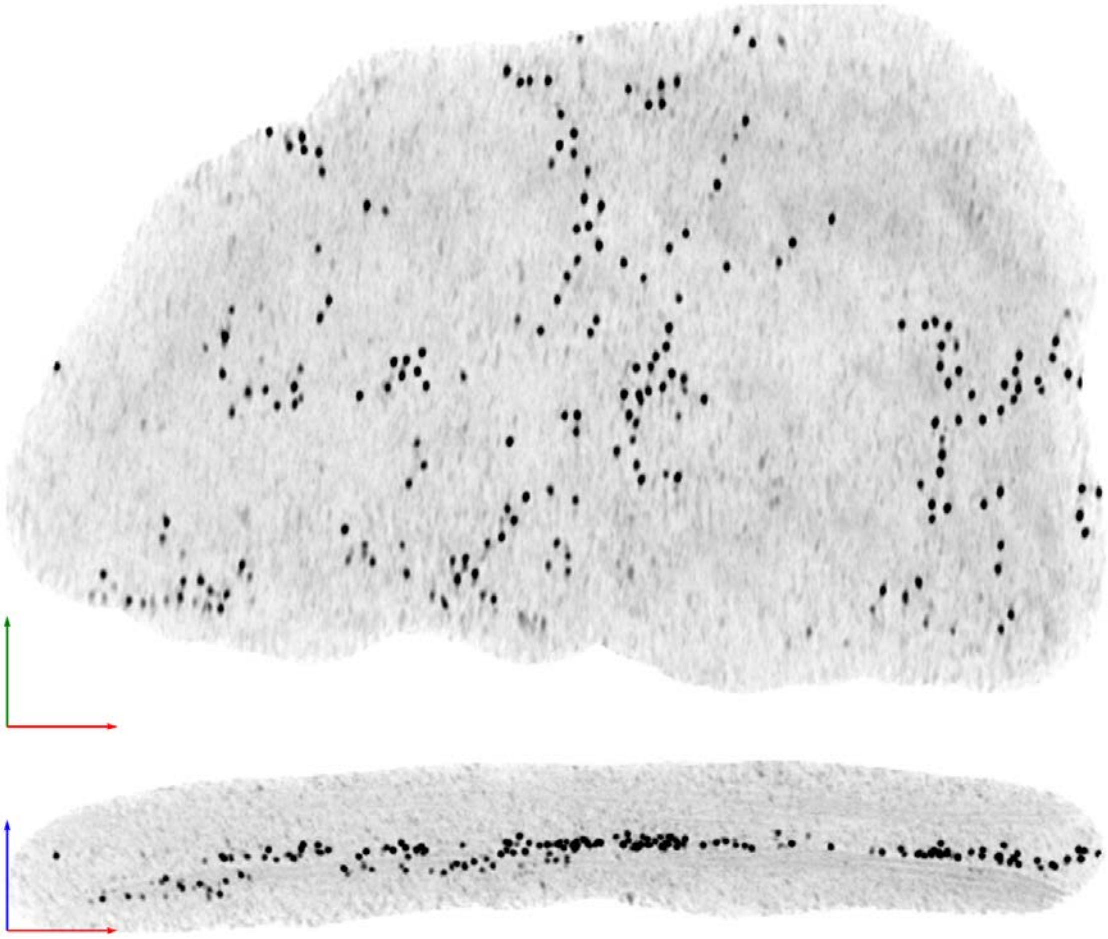

Tomogram ID: 20240111_WaffleHipp_183 | SET: anti-GluA2 15F1 "C-term"

Cleft ID: 0 | Type: tissue

**Denoised Tomogram Slice**

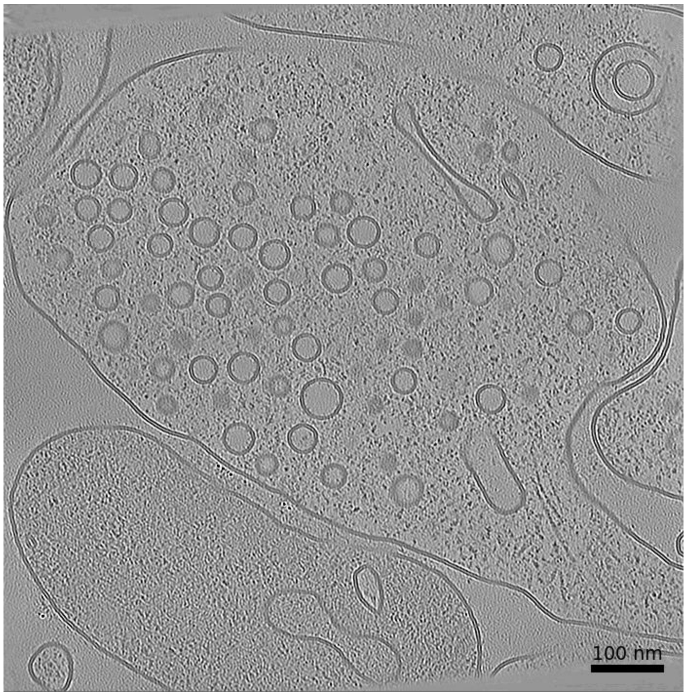


**Membrane Segmentation & Cleft Position**

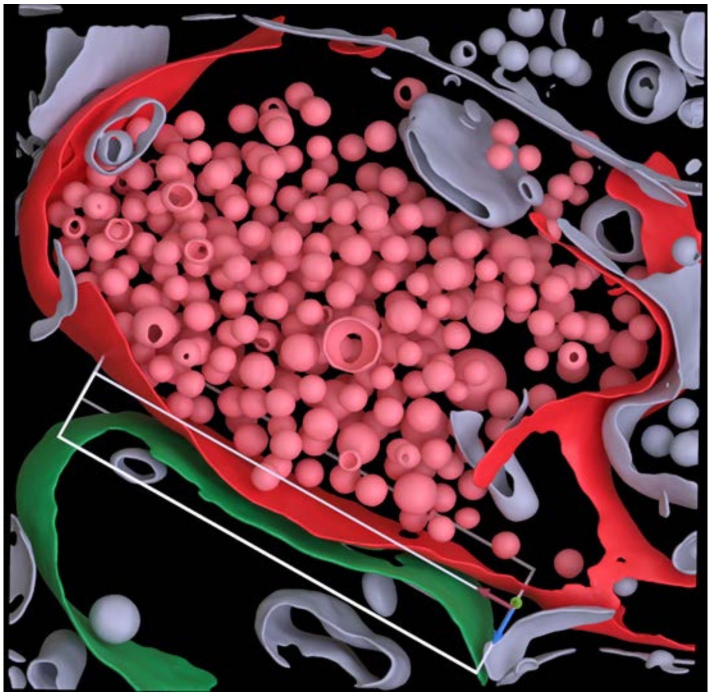

**Cleft MIP**

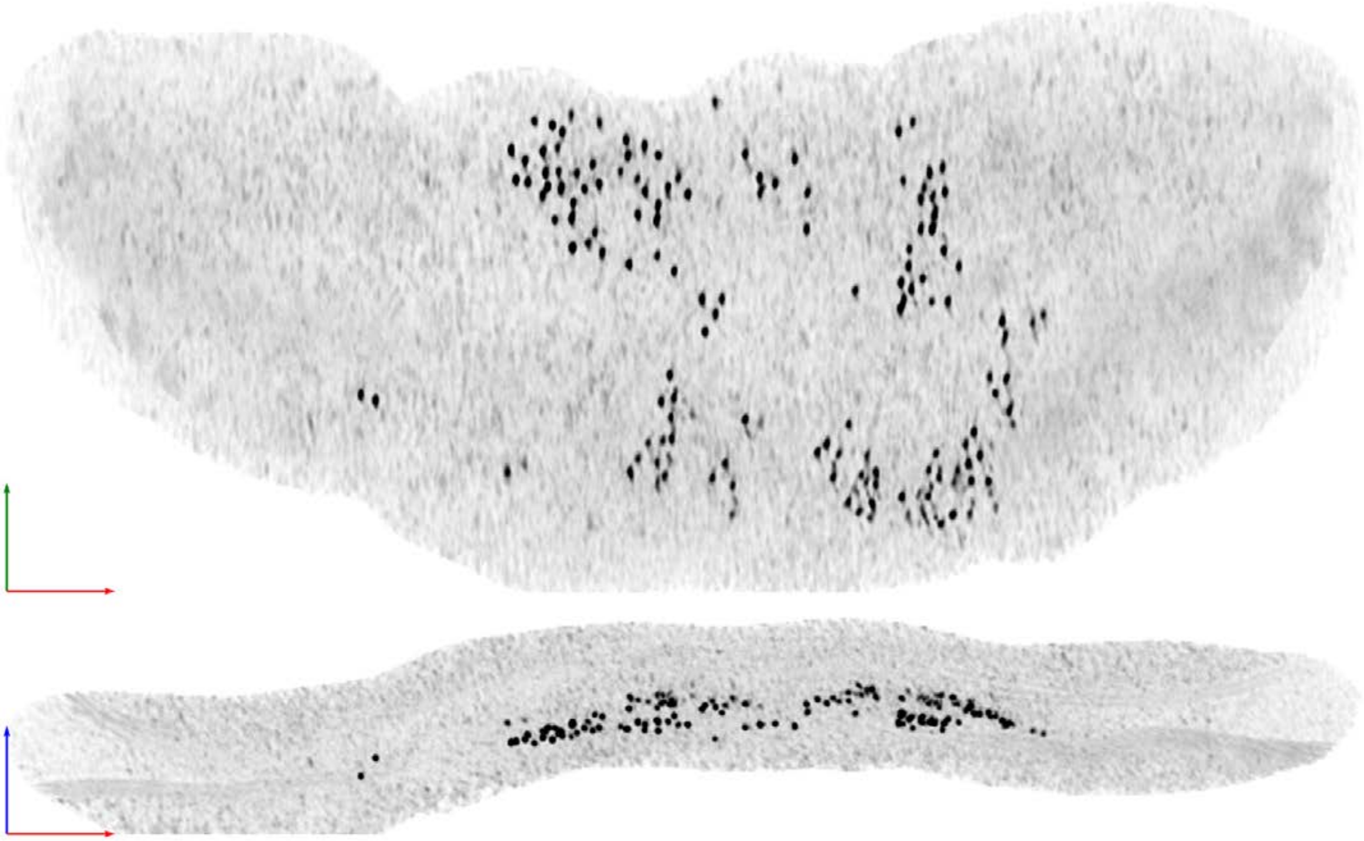

| Tomogram ID: 20240111_WaffleHipp_195 | SET: anti-GluA2 15F1 "C-term" |
|---|---|
| Cleft ID: 0 | Type: synaptosome |

**Denoised Tomogram Slice**

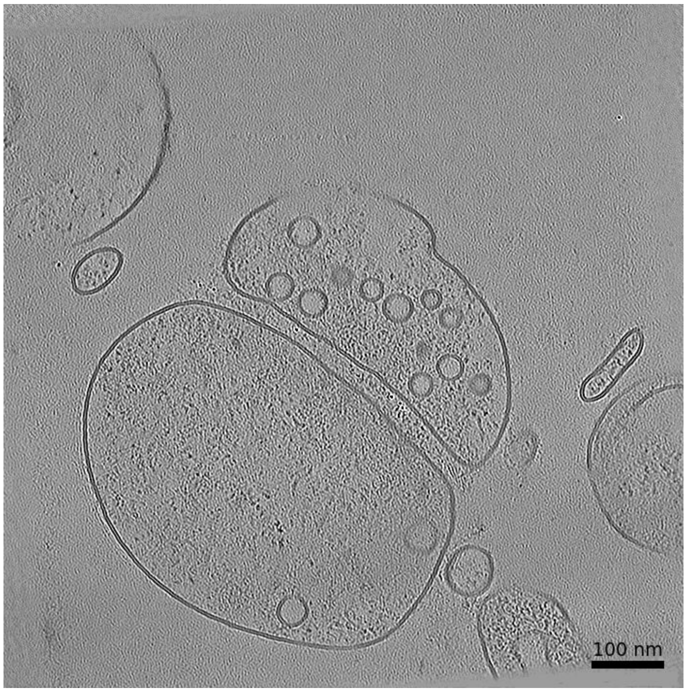


**Membrane Segmentation & Cleft Position**

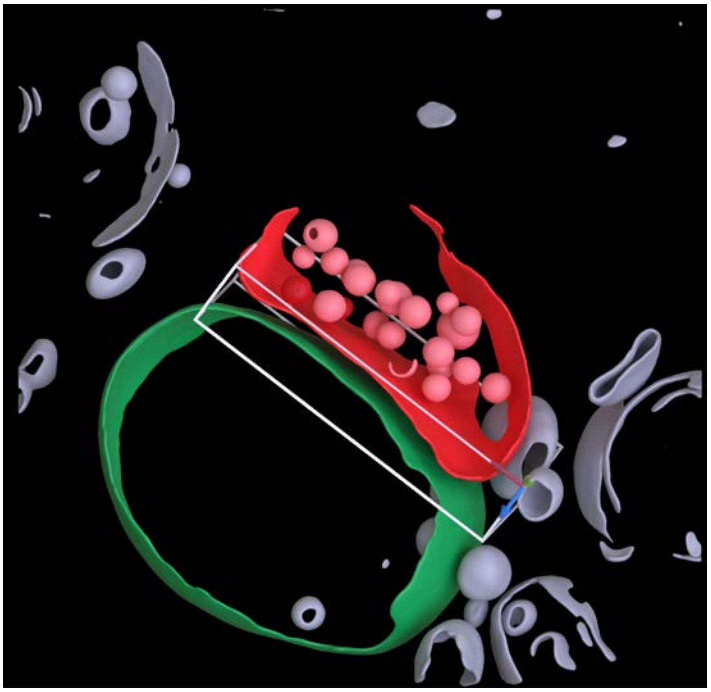

**Cleft MIP**

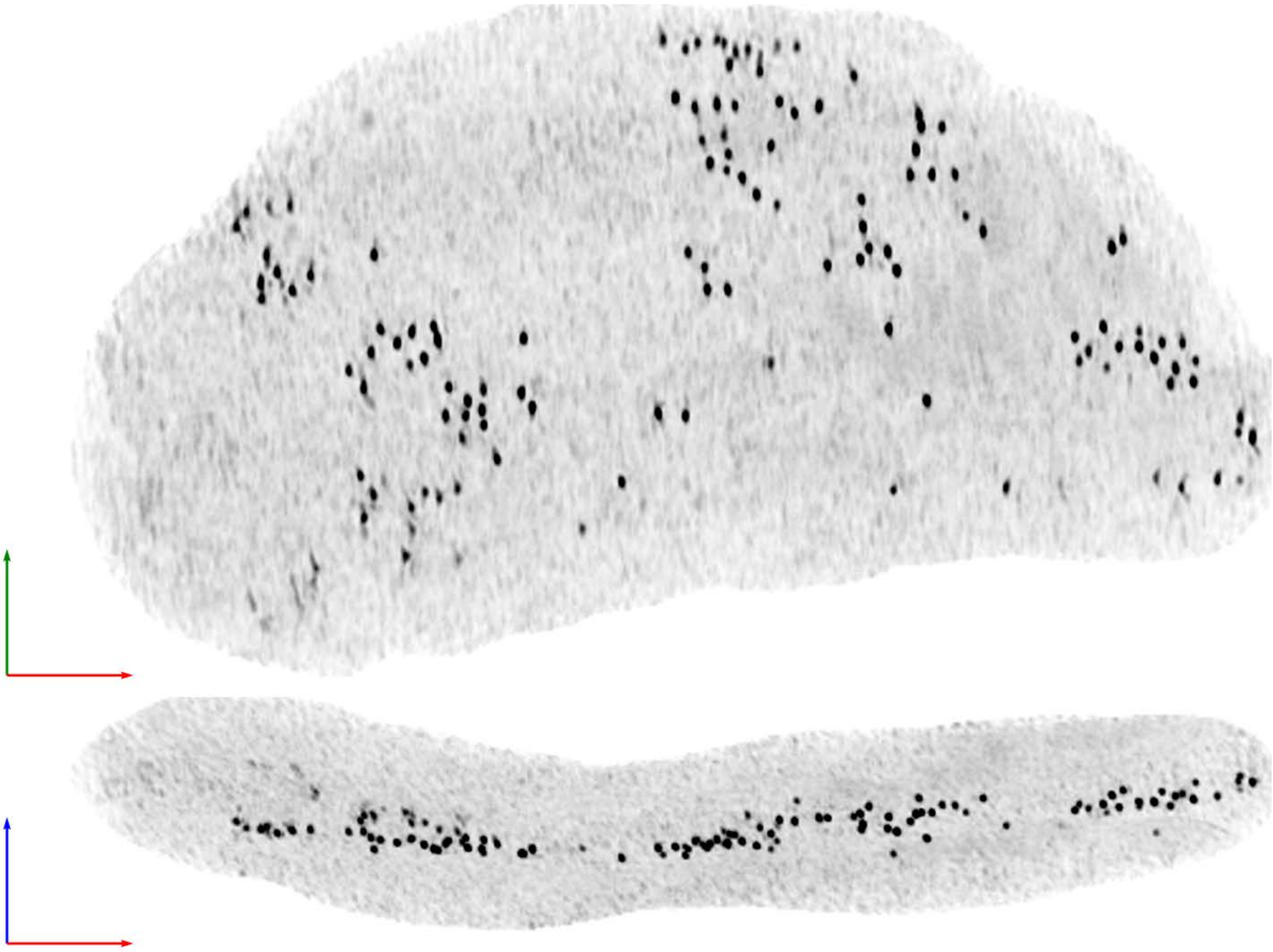

**Tomogram ID: 20240111_WaffleHipp_198** | **SET: anti-GluA2 15F1 "C-term"**

Cleft ID: 0 | Type: tissue

**Denoised Tomogram Slice**

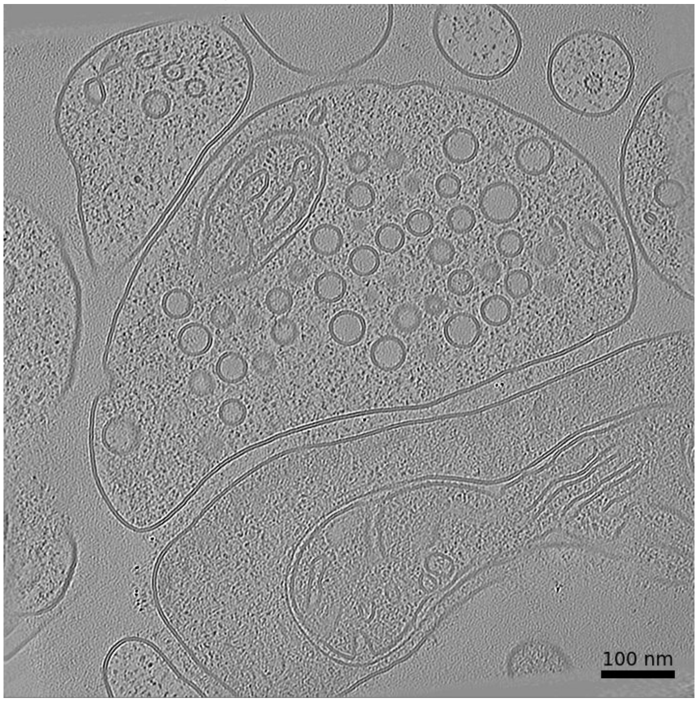


**Membrane Segmentation & Cleft Position**

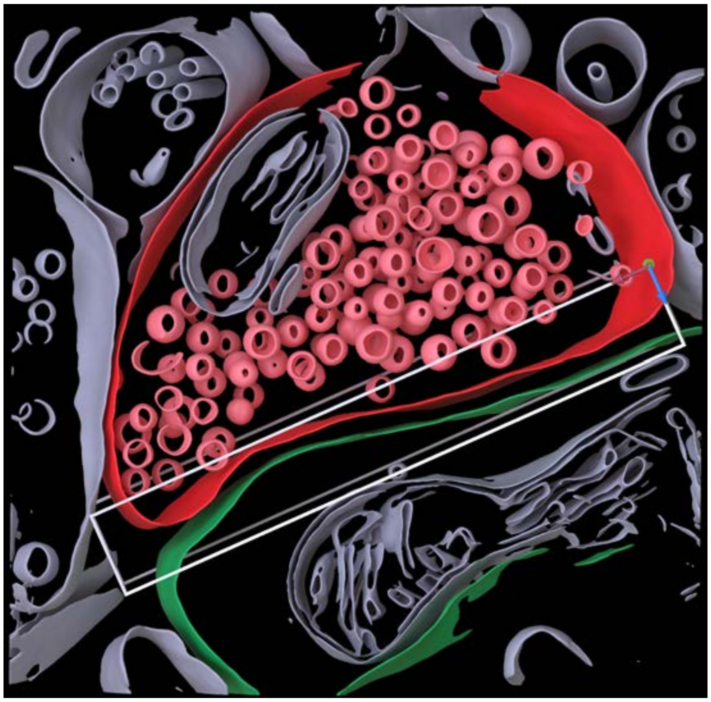

**Cleft MIP**

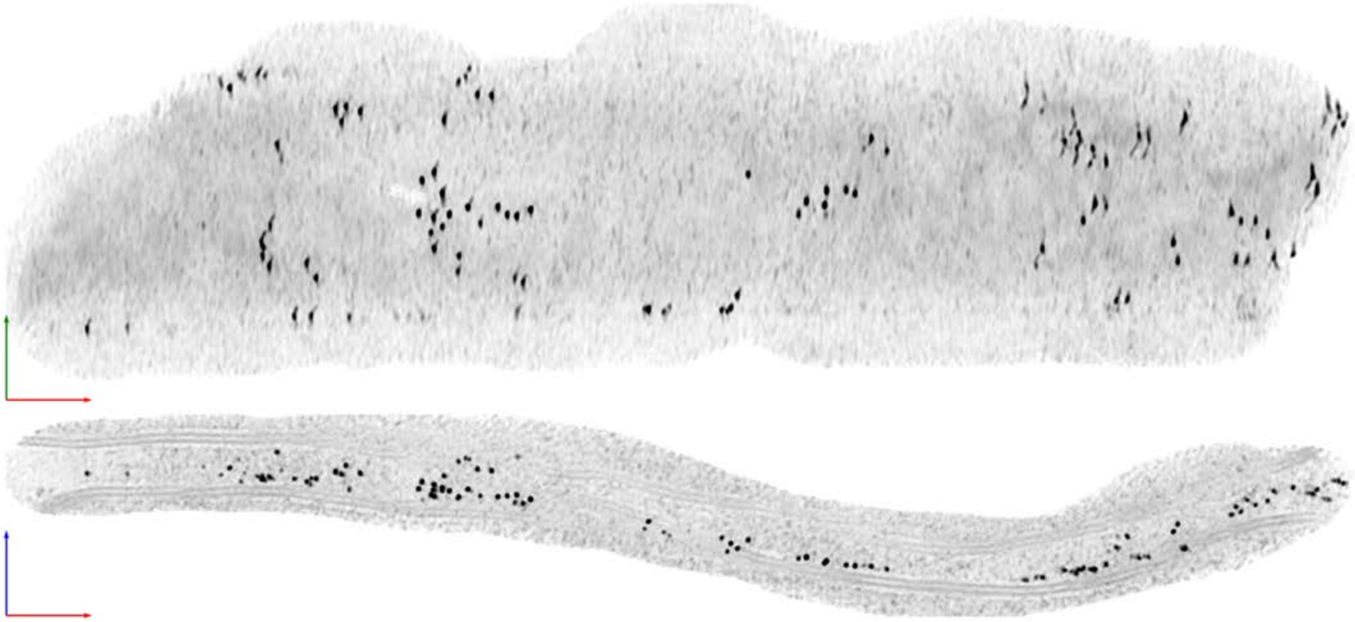

**Tomogram ID: 20240111_WaffleHipp_227** **SET: anti-GluA2 15F1 "C-term"**

Cleft ID: 0 Type: semi-tissue

### Denoised Tomogram Slice

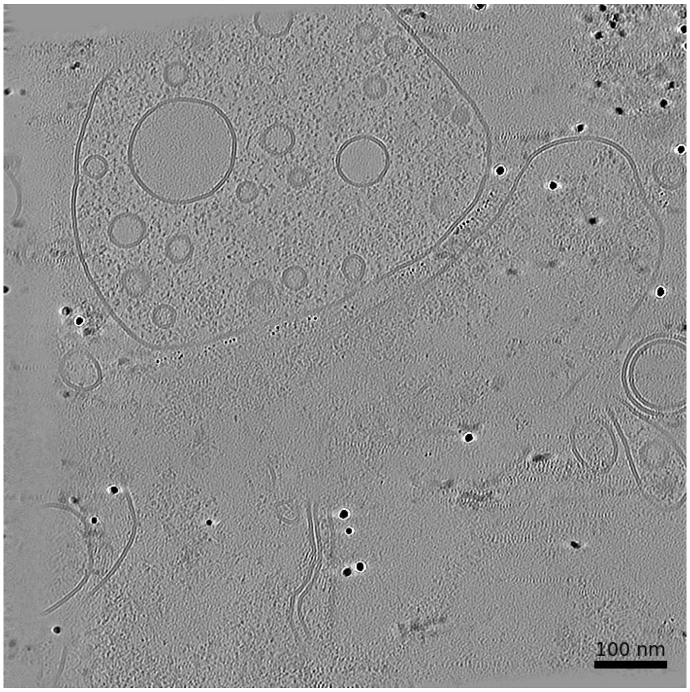


### Membrane Segmentation & Cleft Position

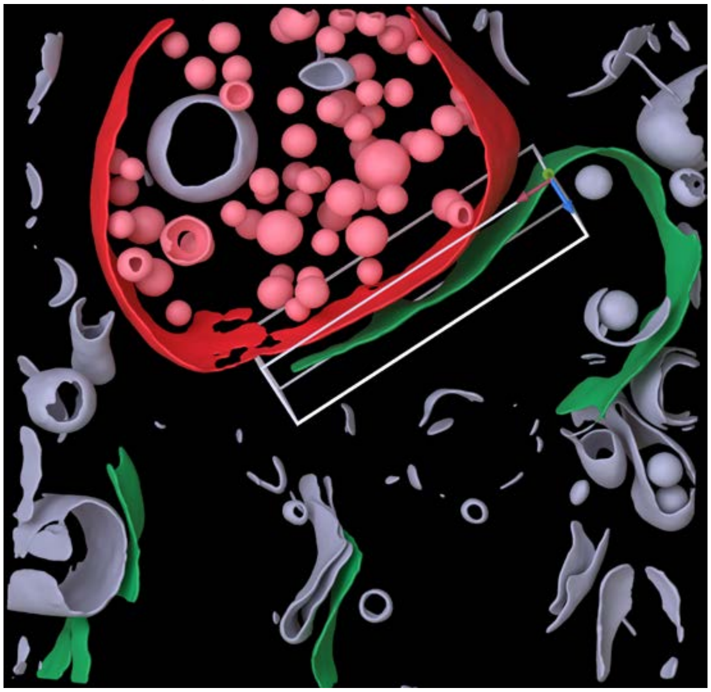

### Cleft MIP

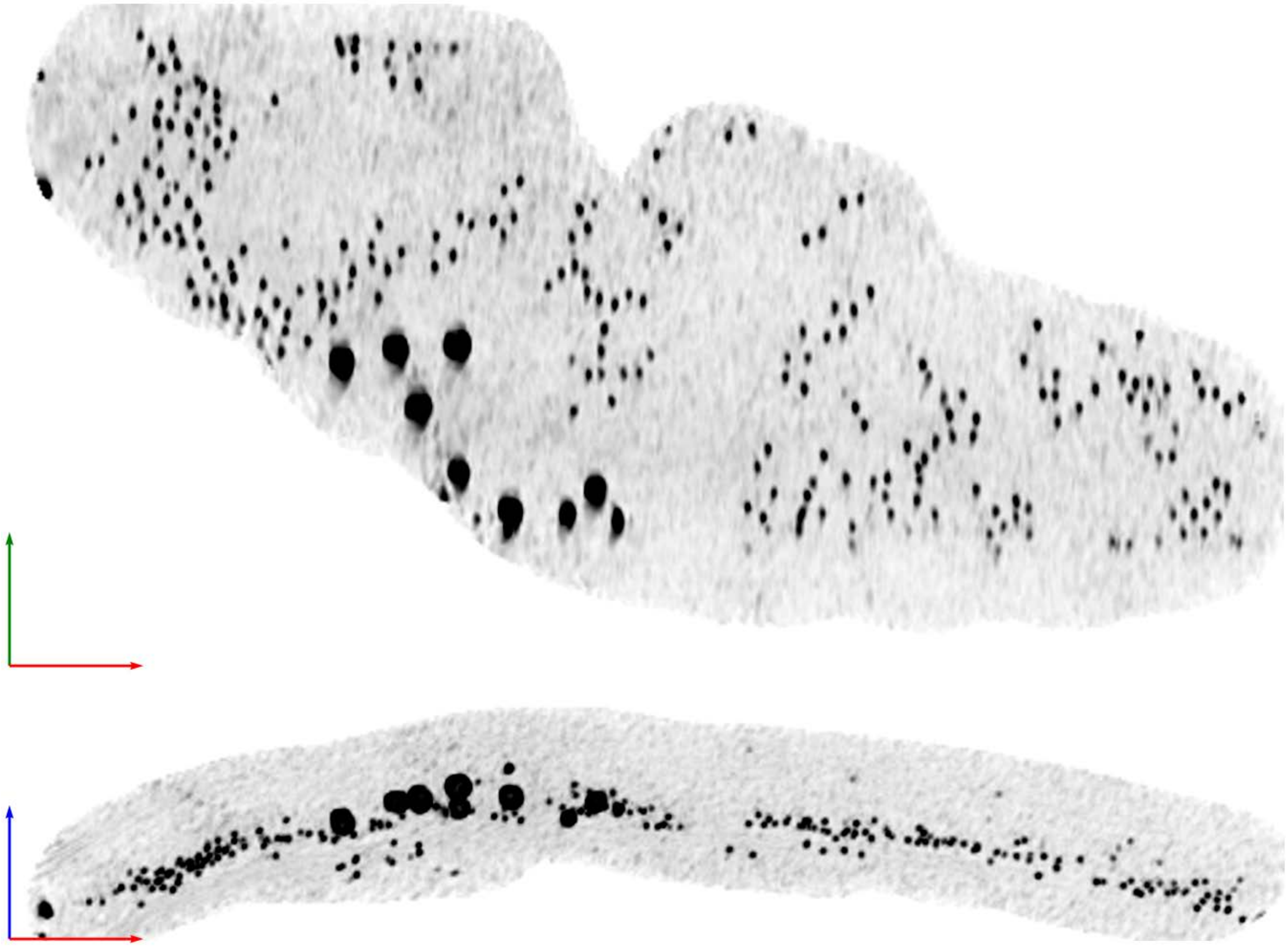

| Tomogram ID: 20240209_waffleHip_78 | SET: anti-GluA2 15F1 "C-term" |
| --- | --- |
| Cleft ID: 0 | Type: semi-tissue |

**Denoised Tomogram Slice**

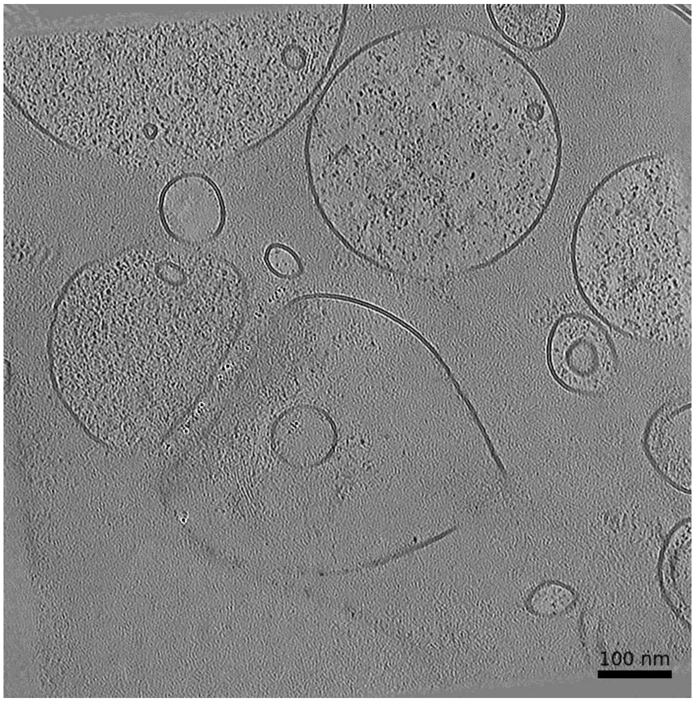


**Membrane Segmentation & Cleft Position**

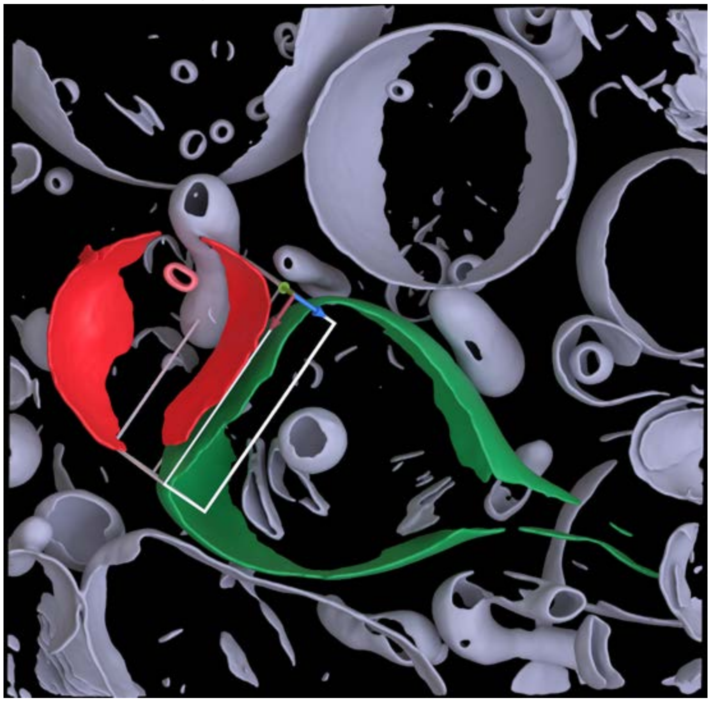

**Cleft MIP**

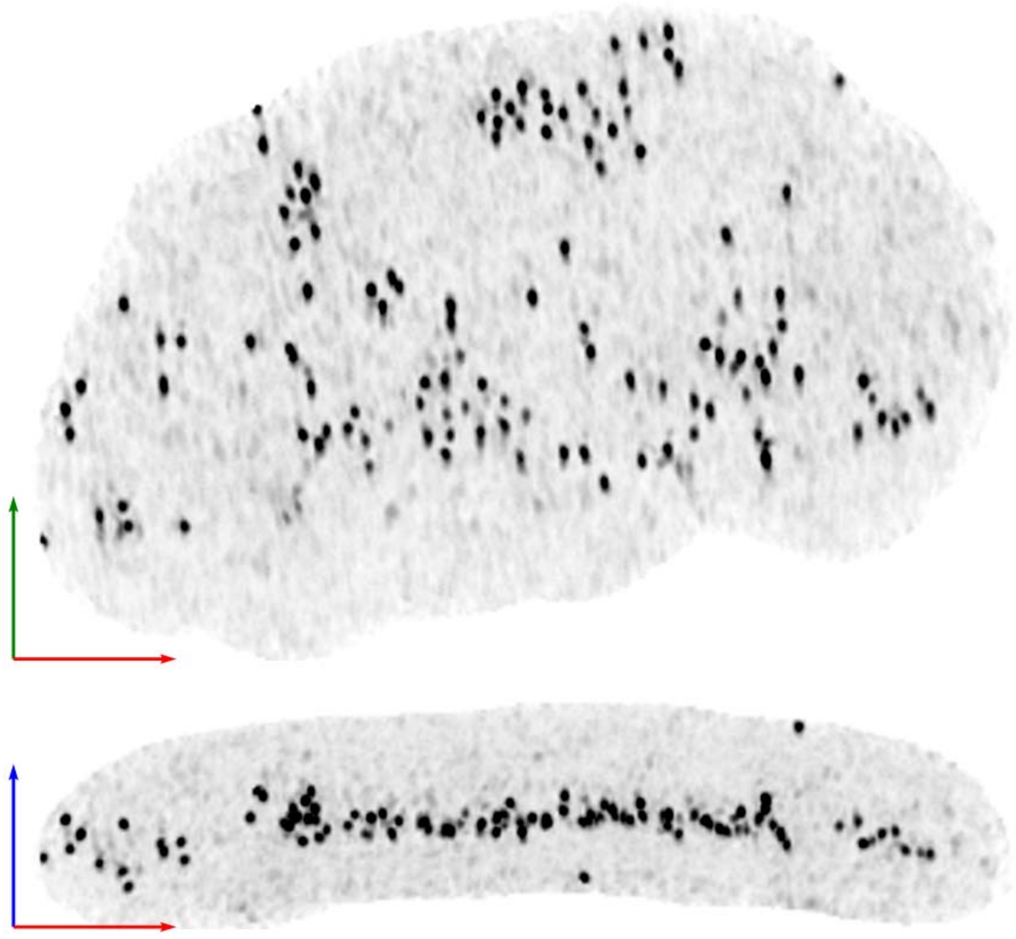

**Tomogram ID: 20240523_HippWaffle_71** **SET: anti-GluA2 15F1 "C-term"**

Cleft ID: 0 Type: tissue

### Denoised Tomogram Slice

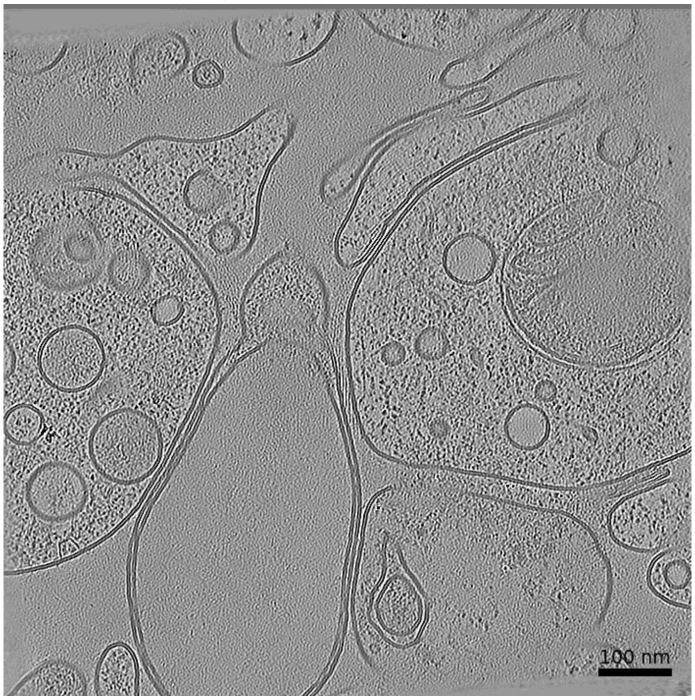


### Membrane Segmentation & Cleft Position

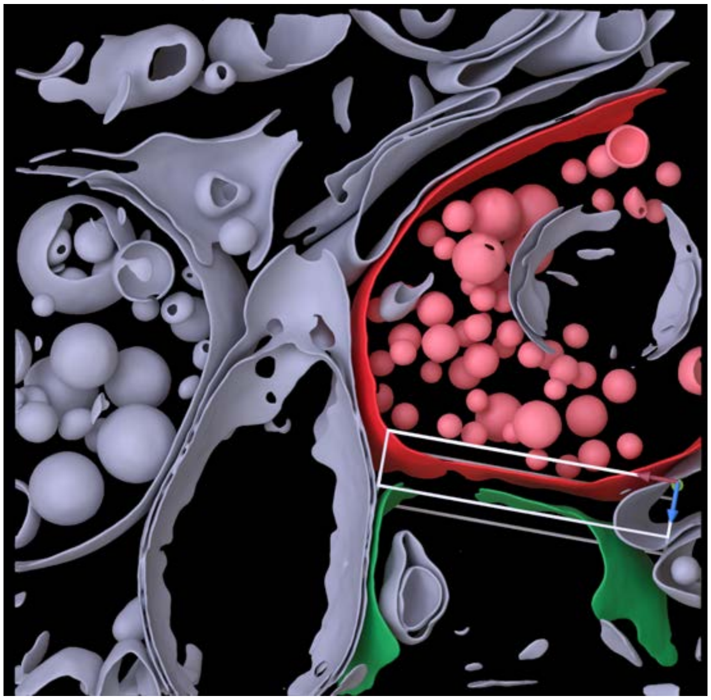

### Cleft MIP

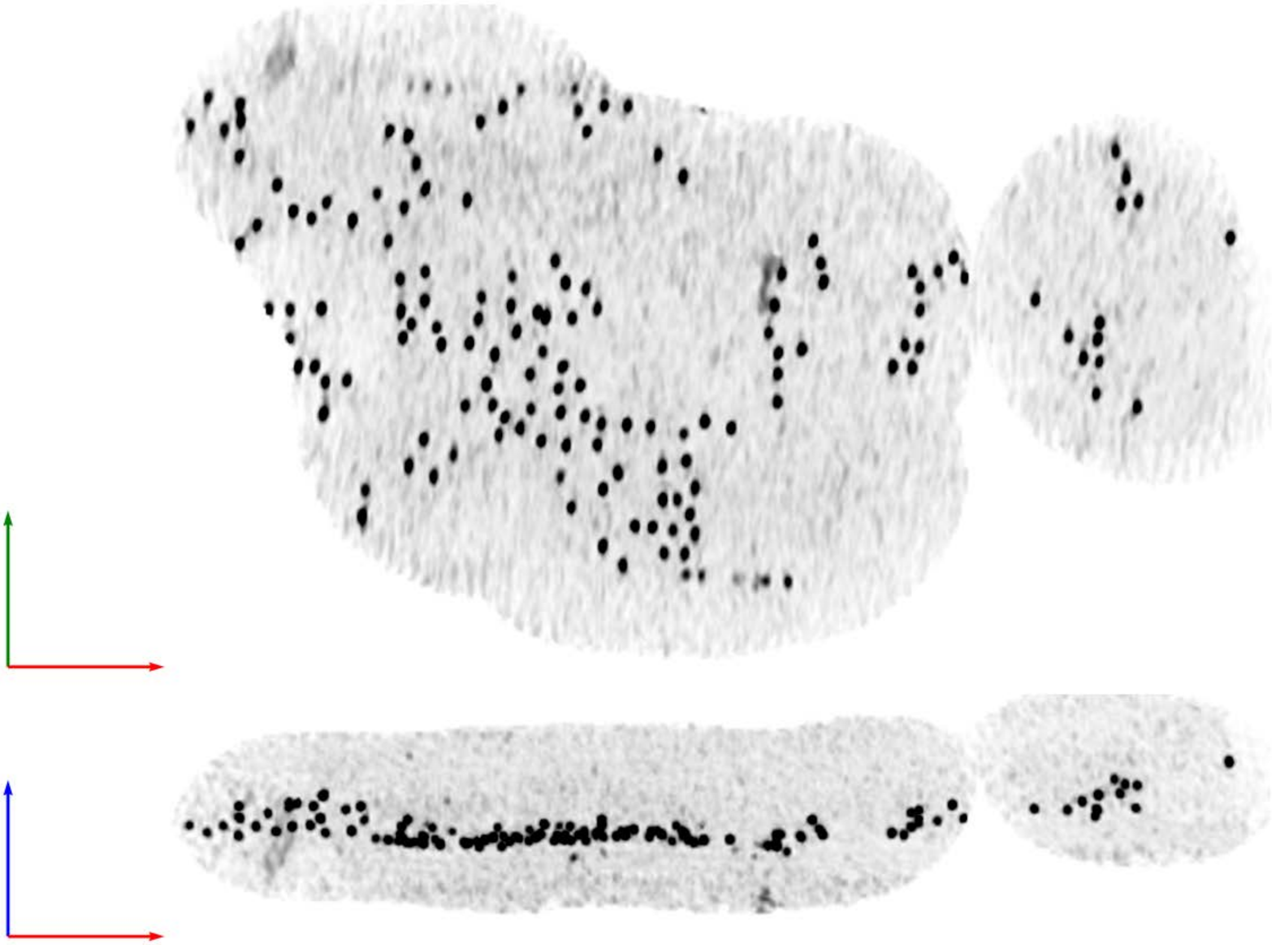

Tomogram ID: 20240624_HippWaffle_21 | SET: anti-GluA2 15F1 "C-term"

Cleft ID: 0 | Type: tissue

**Denoised Tomogram Slice**

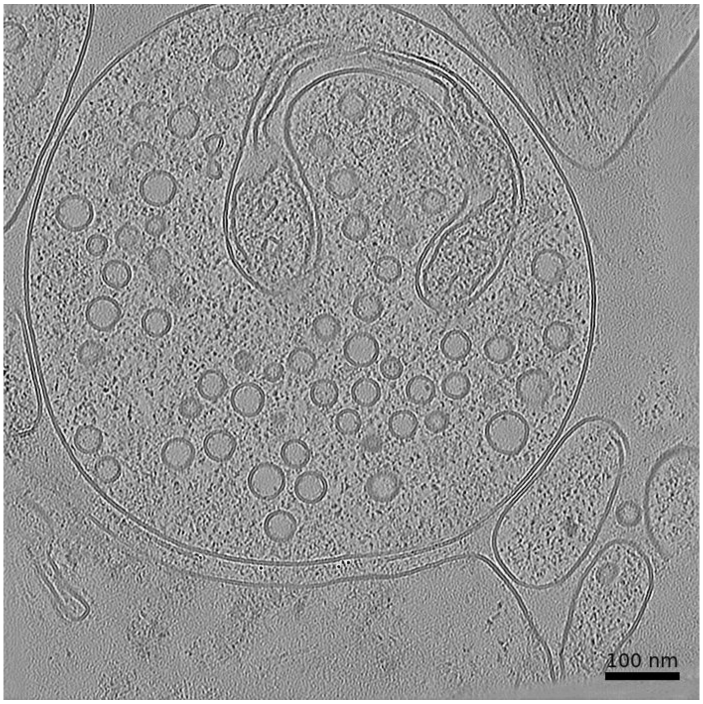


**Membrane Segmentation & Cleft Position**

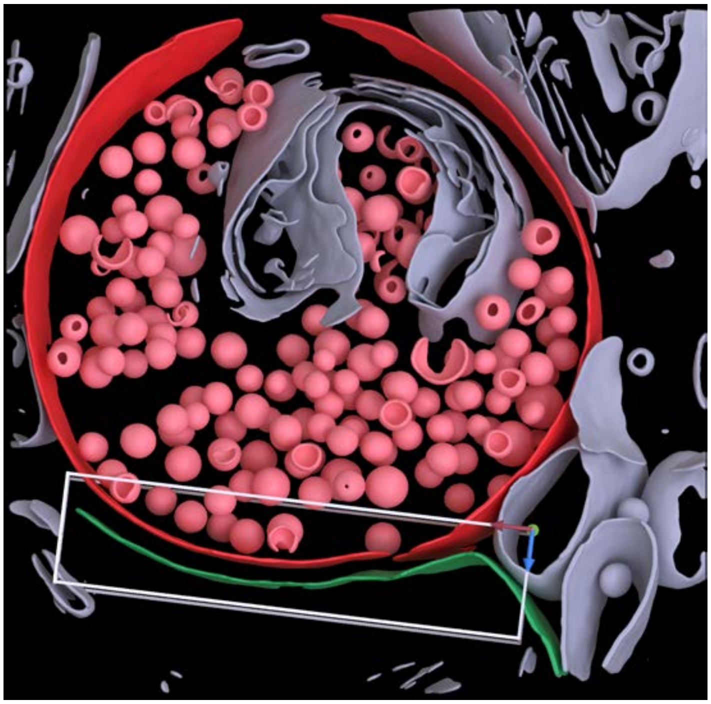

**Cleft MIP**

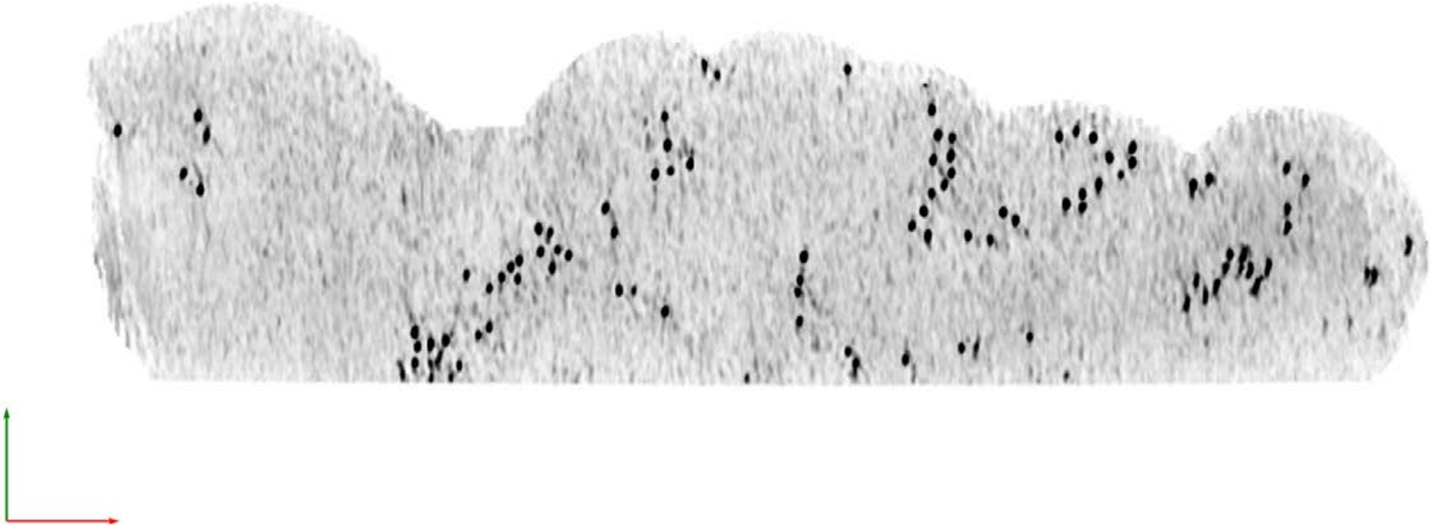

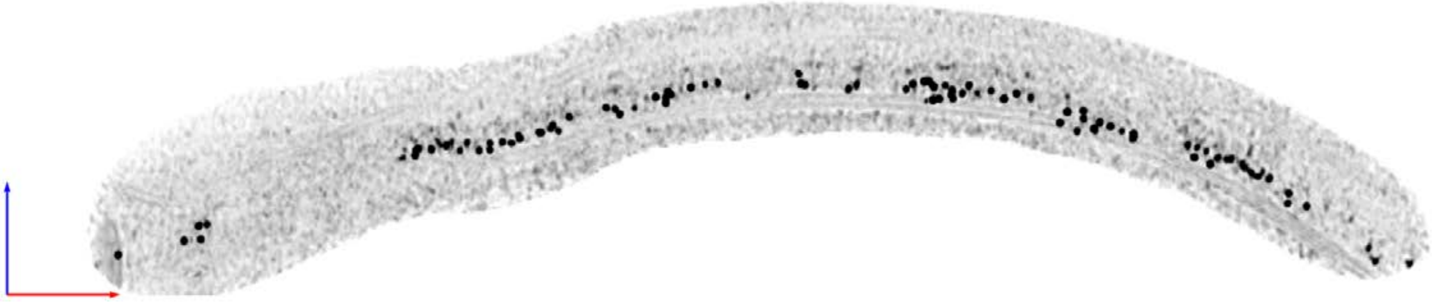

| **Tomogram ID: 20240624_HippWaffle_76** | **SET: anti-GluA2 15F1 "C-term"** |
|---|---|
| Cleft ID: 0 | Type: tissue |

**Denoised Tomogram Slice**

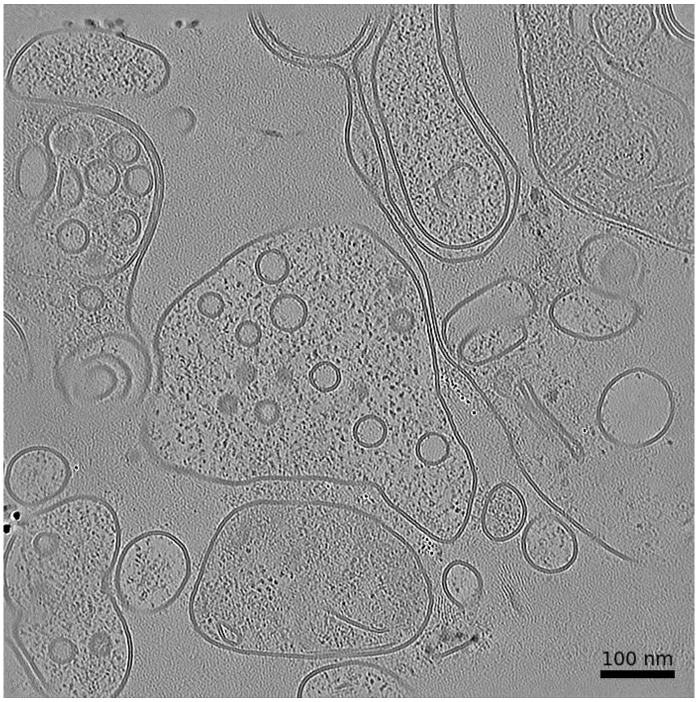


**Membrane Segmentation & Cleft Position**

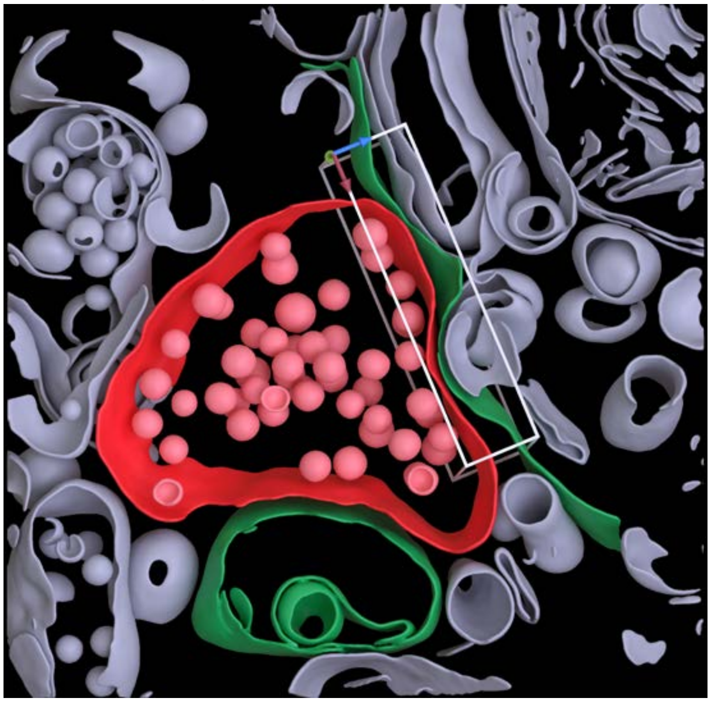

**Cleft MIP**

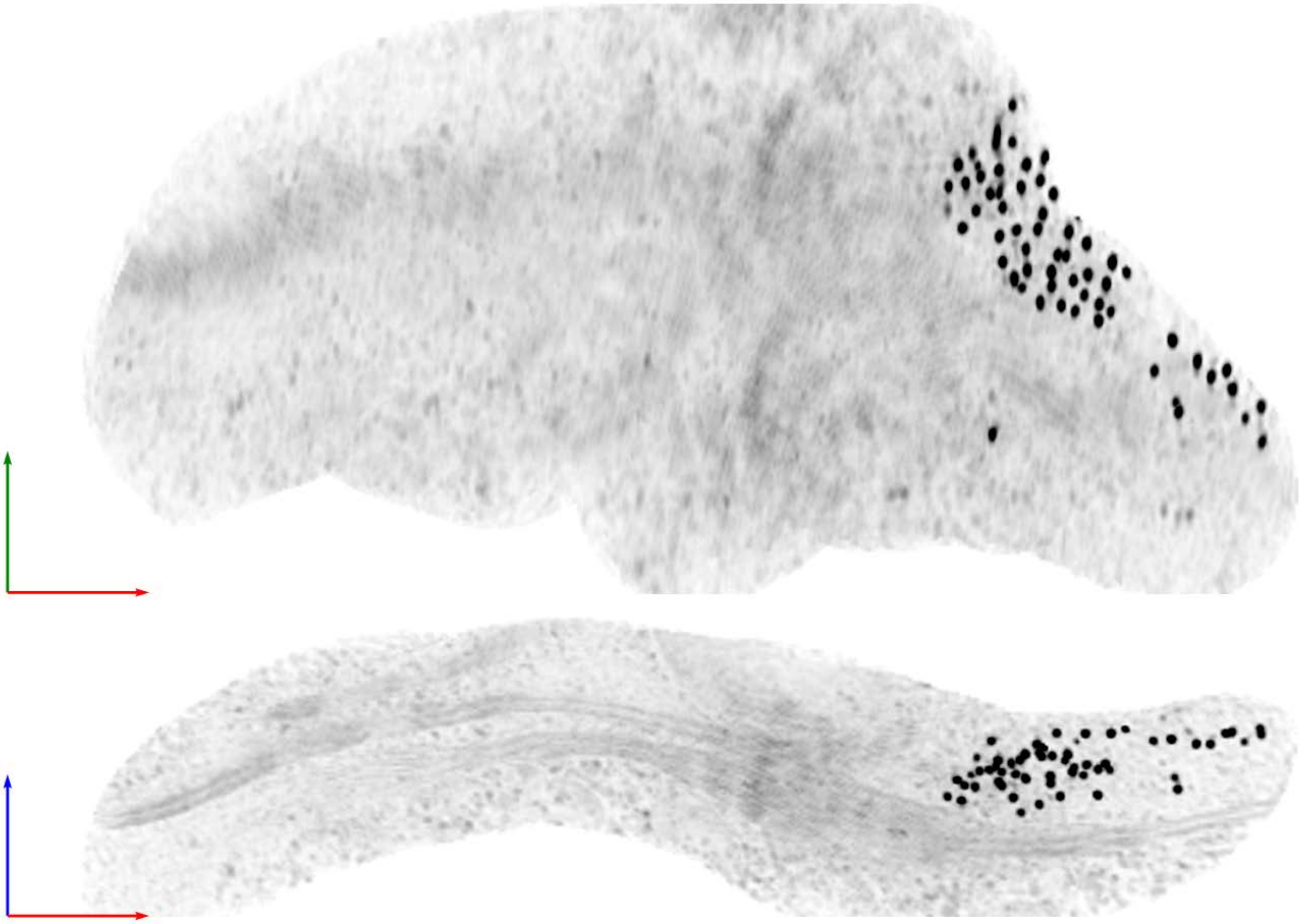

**Tomogram ID: 20240624_HippWaffle_76** | **SET: anti-GluA2 15F1 "C-term"**

Cleft ID: 1 | Type: tissue

### Denoised Tomogram Slice

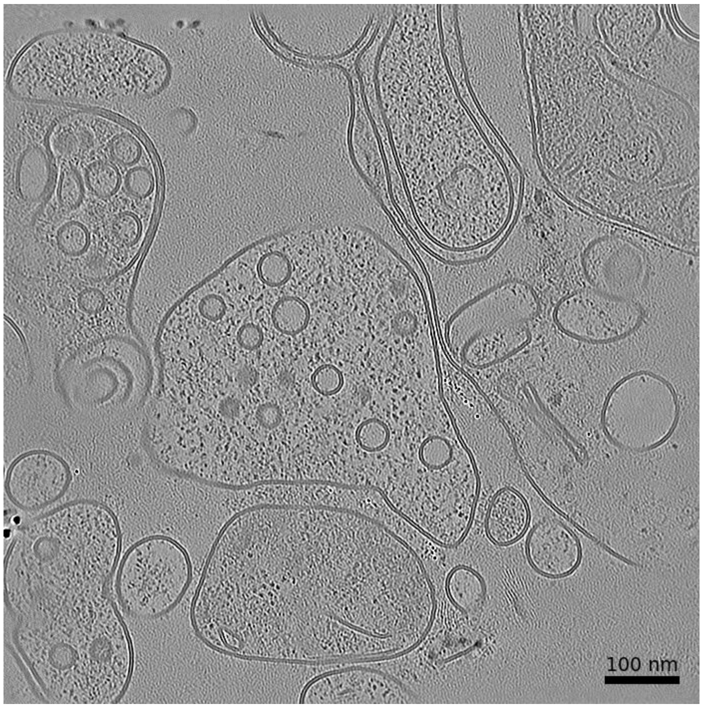


### Membrane Segmentation & Cleft Position

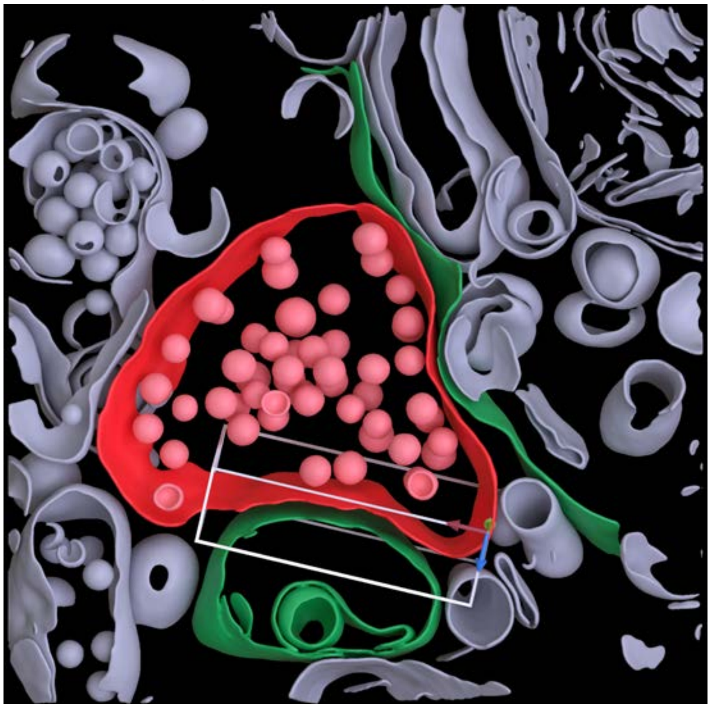

### Cleft MIP

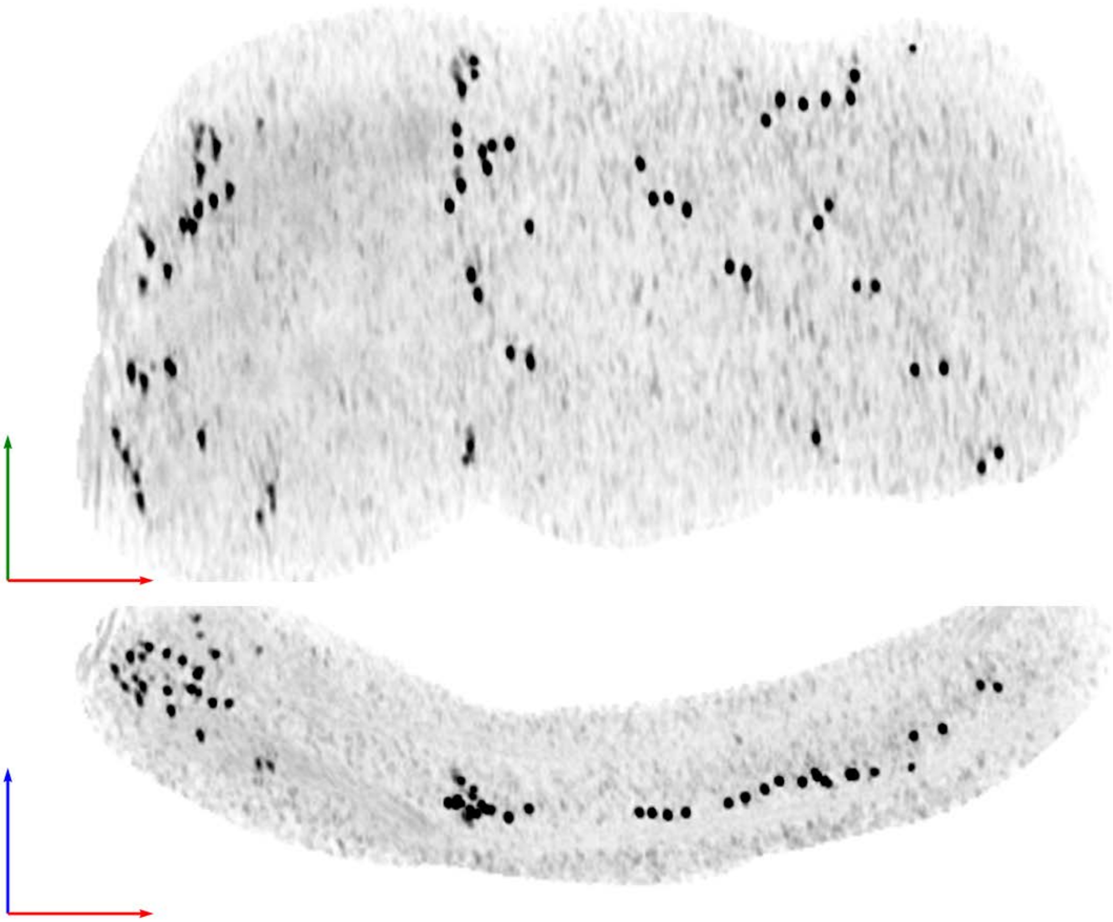

Tomogram ID: 20240624_HippWaffle_86 SET: anti-GluA2 15F1 "C-term"

Cleft ID: 0 Type: tissue

**Denoised Tomogram Slice**

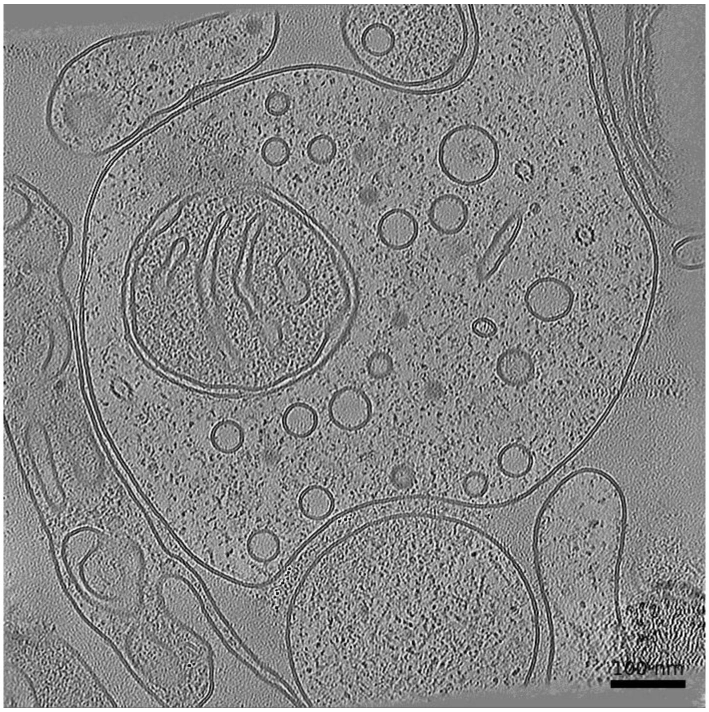


**Membrane Segmentation & Cleft Position**

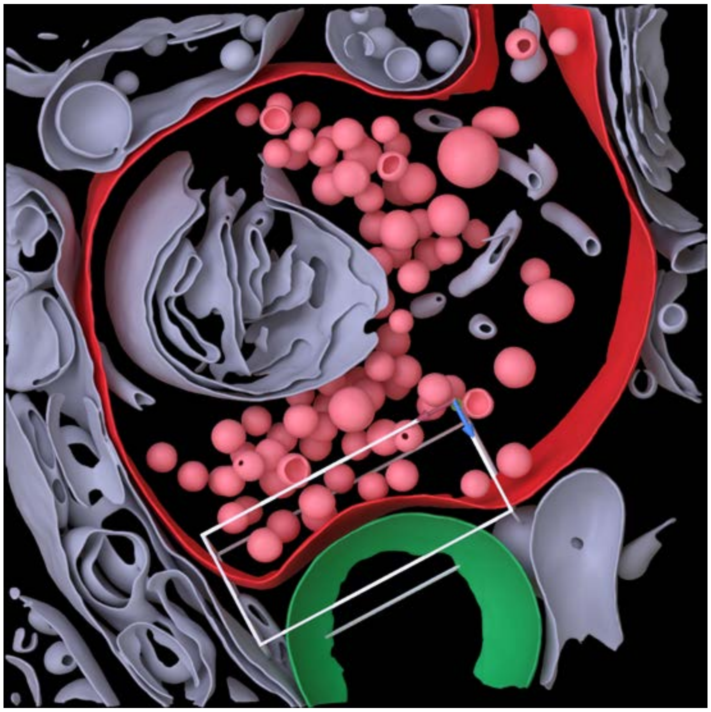

**Cleft MIP**

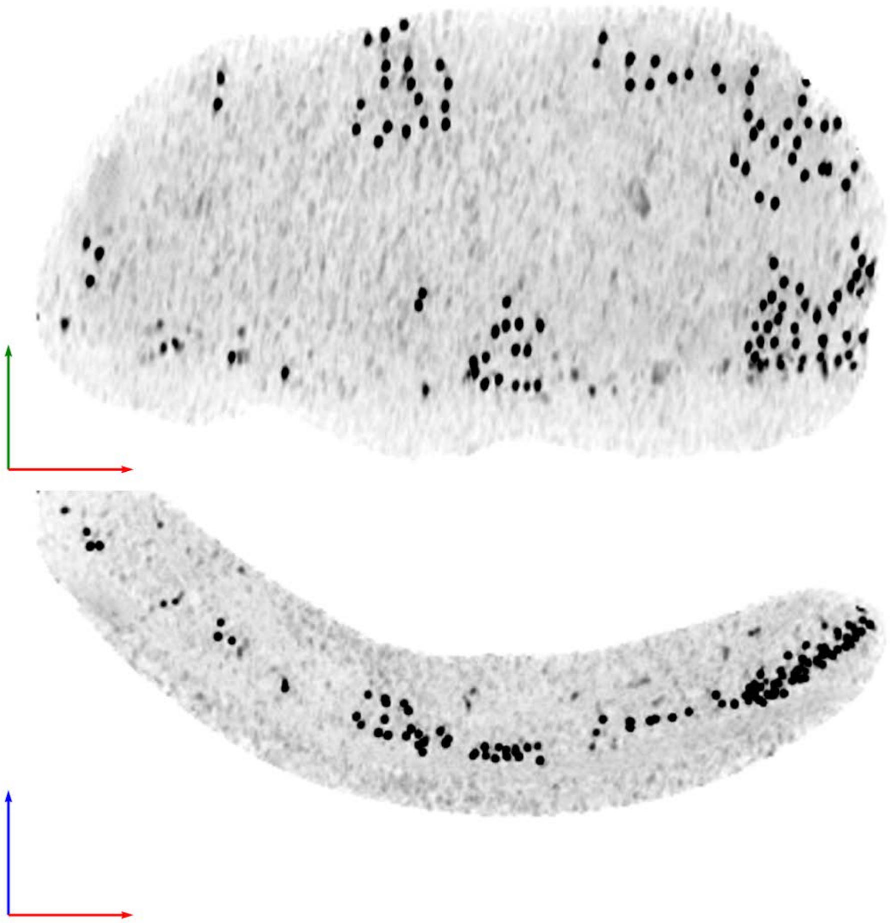

| Tomogram ID: 20241030_AMmilled12-1_12 | SET: anti-GluA2 15F1 "C-term" |
|---|---|
| Cleft ID: 0 | Type: tissue |

**Denoised Tomogram Slice**

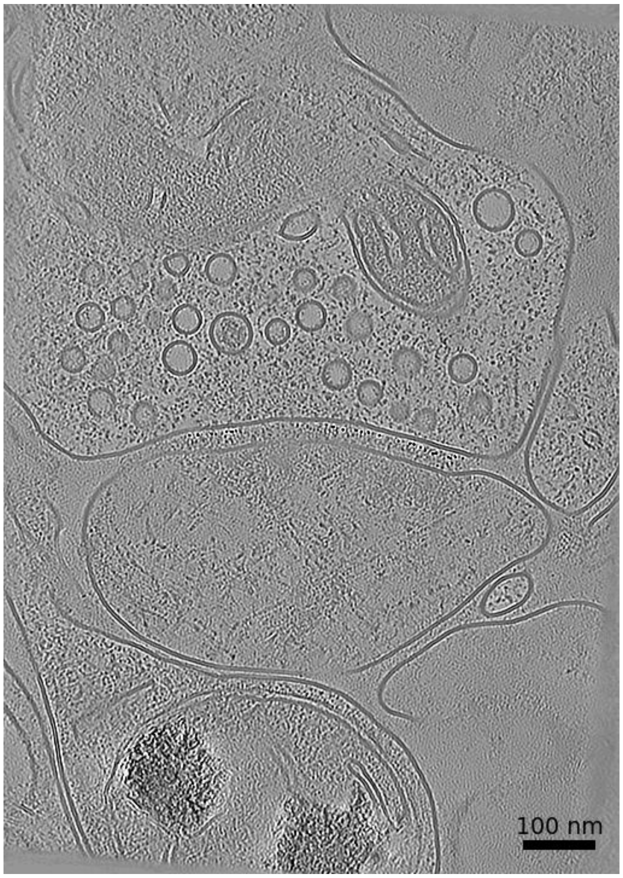


**Membrane Segmentation & Cleft Position**

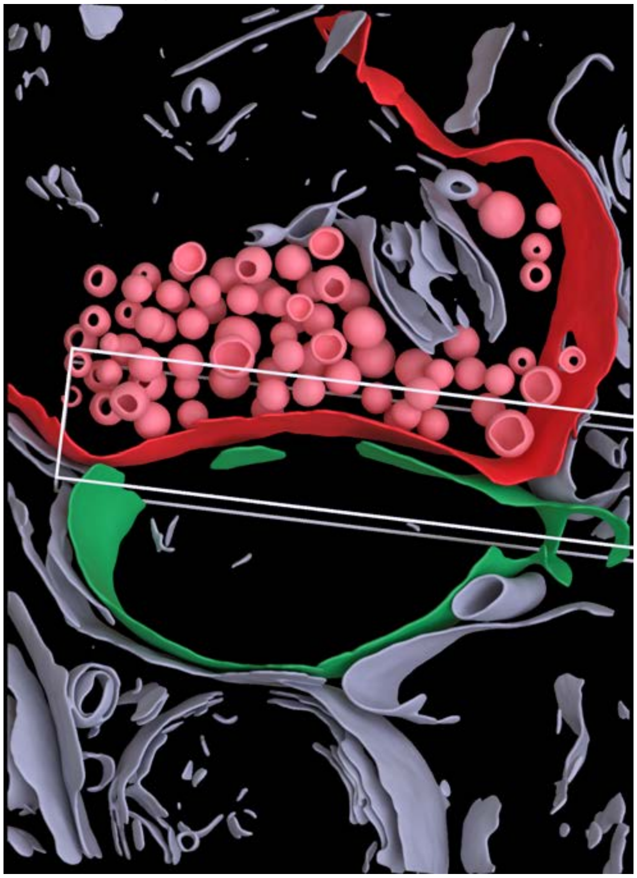

**Cleft MIP**

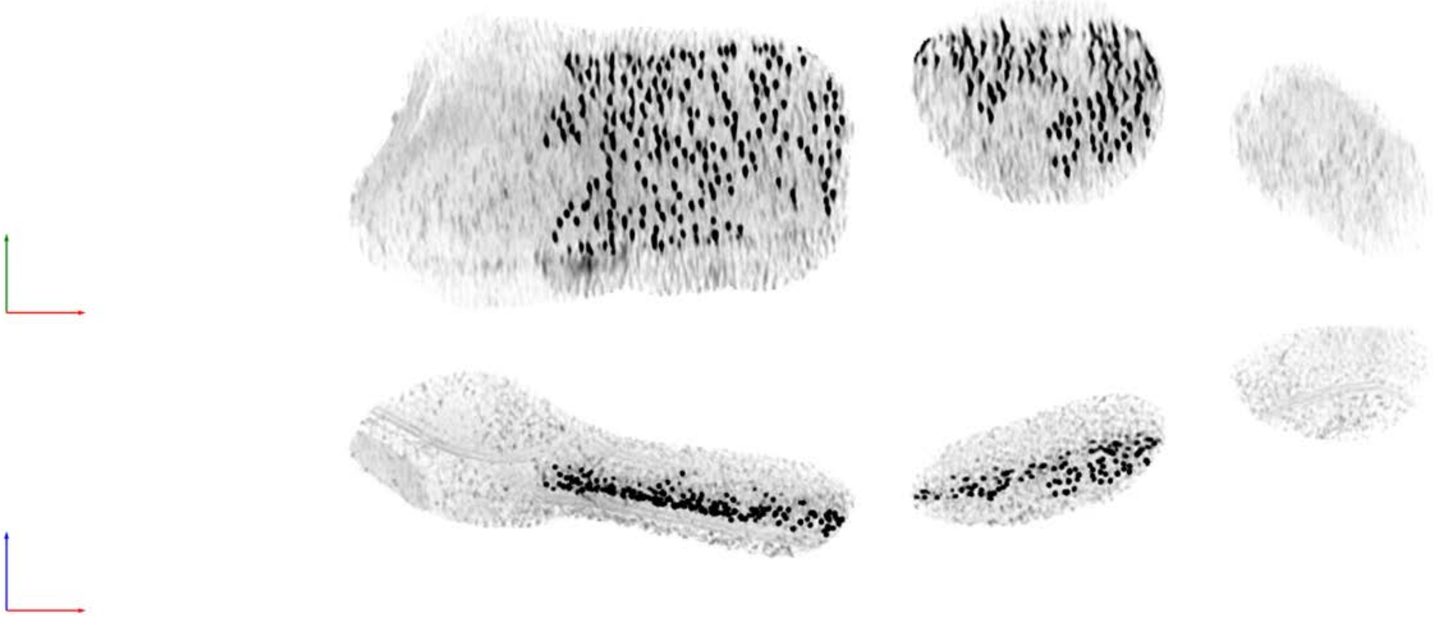

**Tomogram ID: 20241030_AMmilled12-1_15** **SET: anti-GluA2 15F1 "C-term"**

Cleft ID: 0 Type: tissue

**Denoised Tomogram Slice**

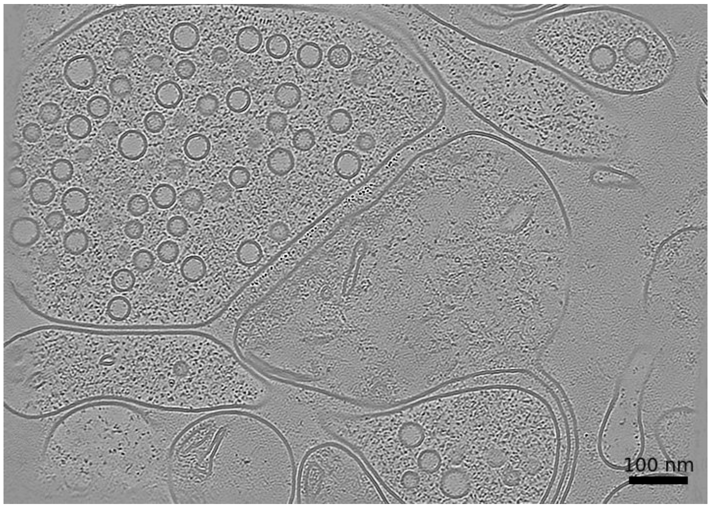


**Membrane Segmentation & Cleft Position**

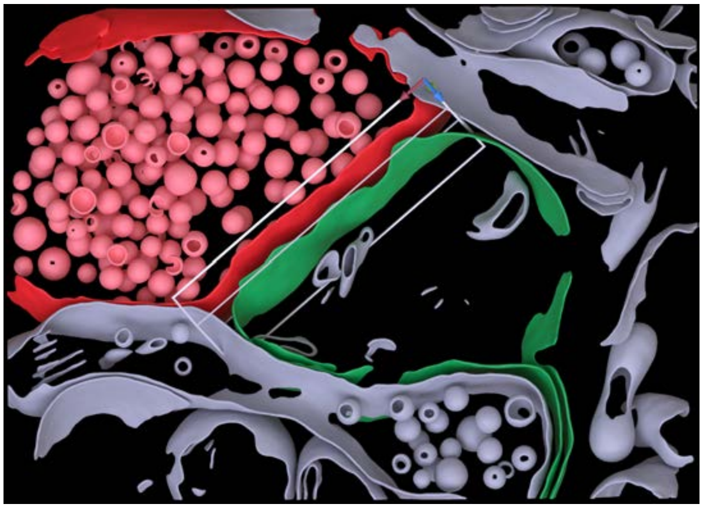

**Cleft MIP**

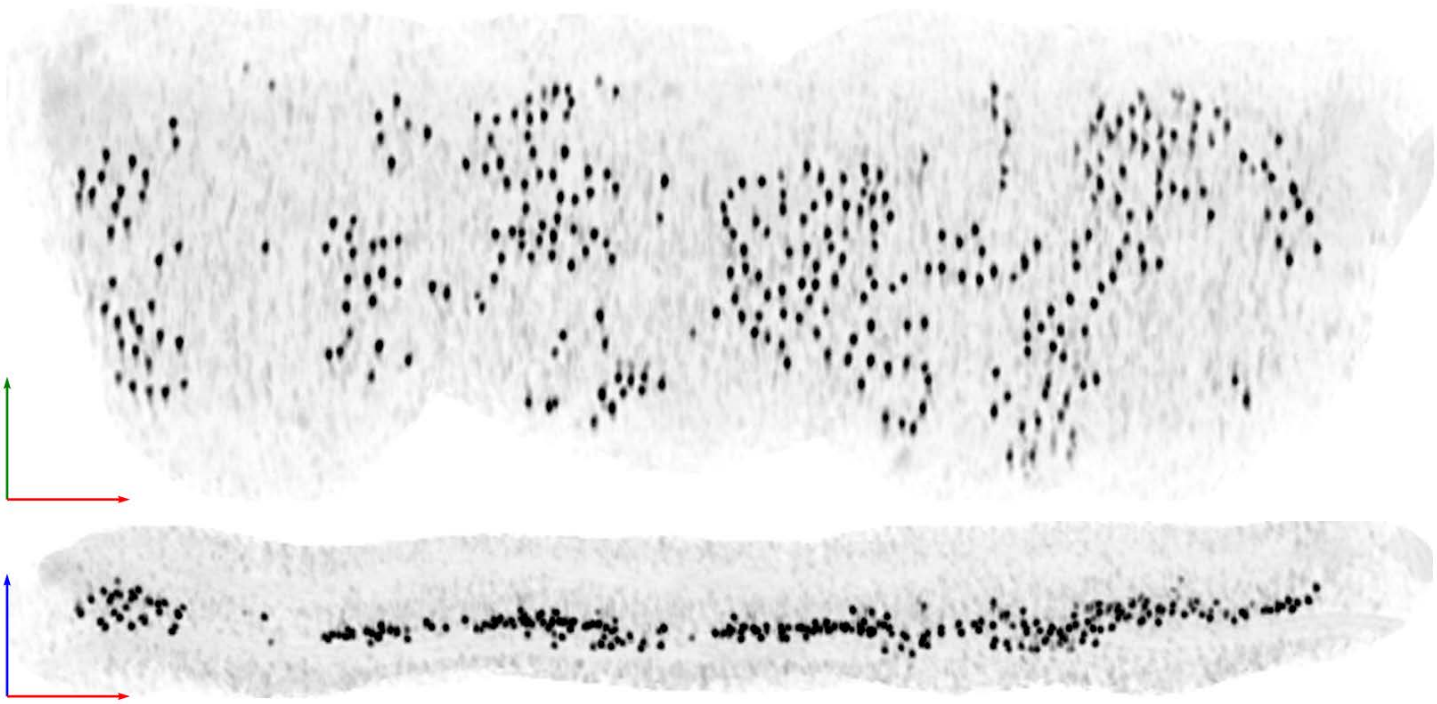

| Tomogram ID: 20241030_AMmilled12-1_15 | SET: anti-GluA2 15F1 "C-term" |
|---|---|
| Cleft ID: 2 | Type: tissue |

**Denoised Tomogram Slice**

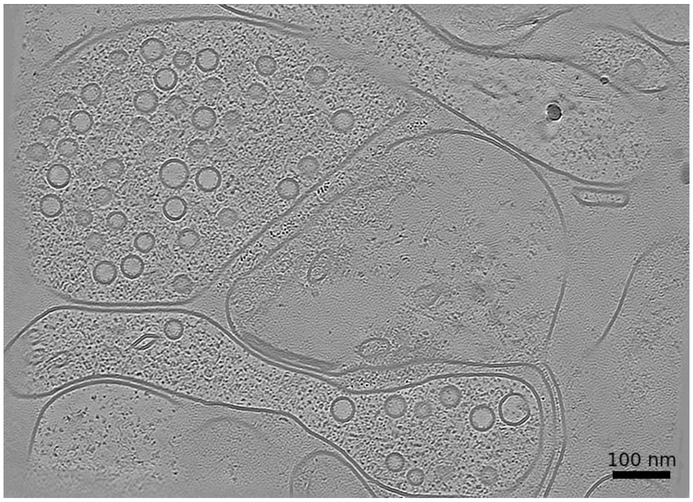


**Membrane Segmentation & Cleft Position**

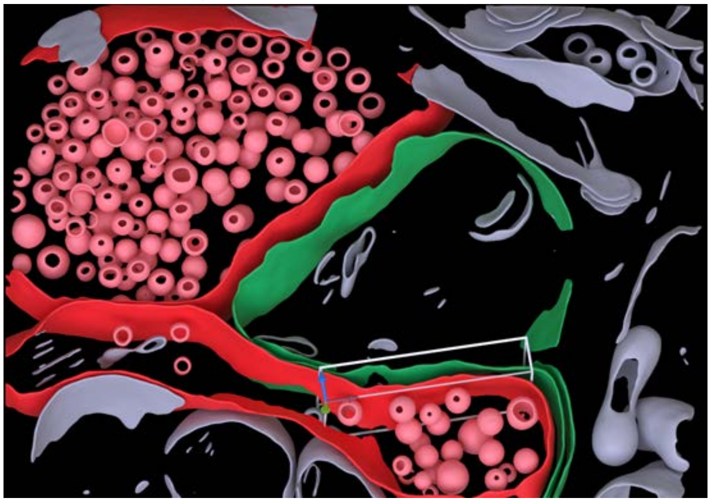

**Cleft MIP**

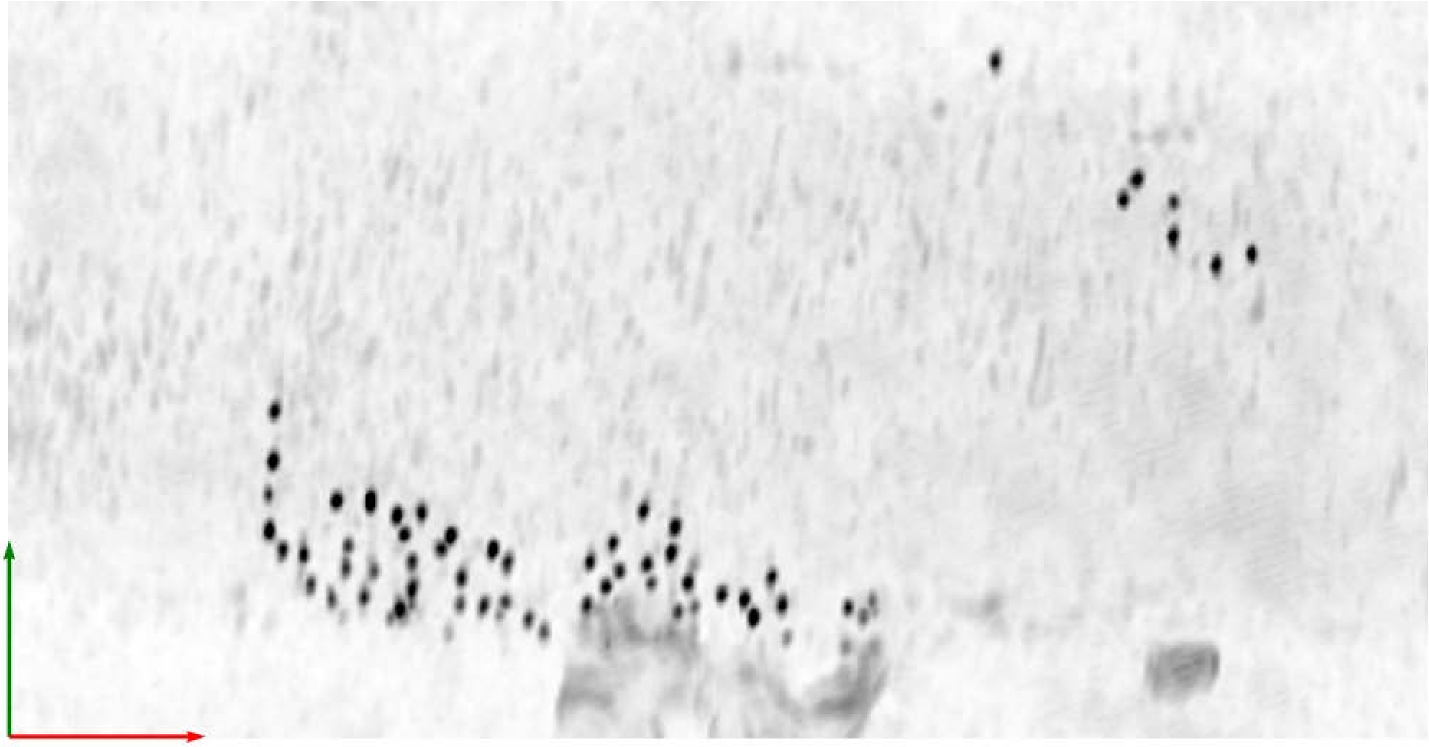

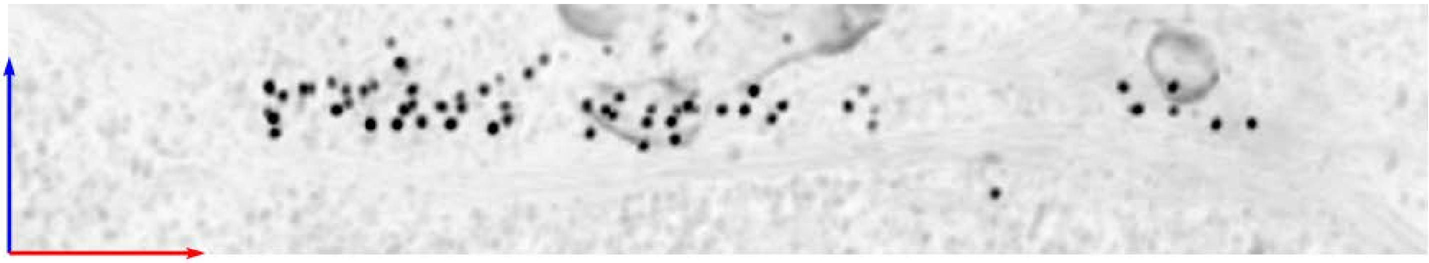

**Tomogram ID: 20241030_AMmilled12-2_78** **SET: anti-GluA2 15F1 "C-term"**

Cleft ID: 0 Type: tissue

**Denoised Tomogram Slice**

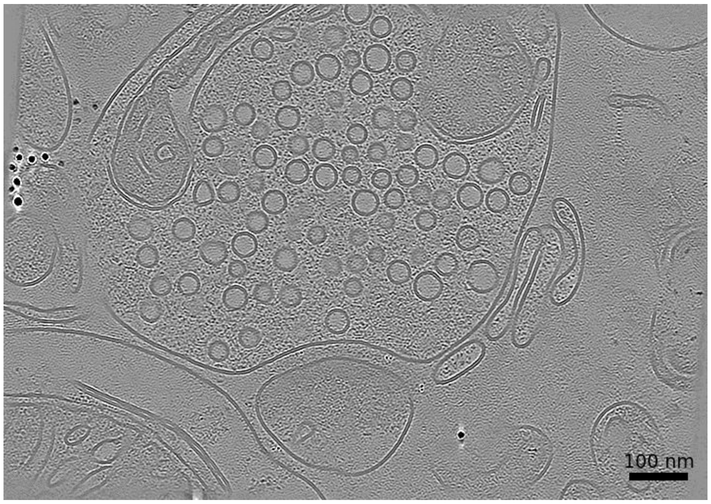


**Membrane Segmentation & Cleft Position**

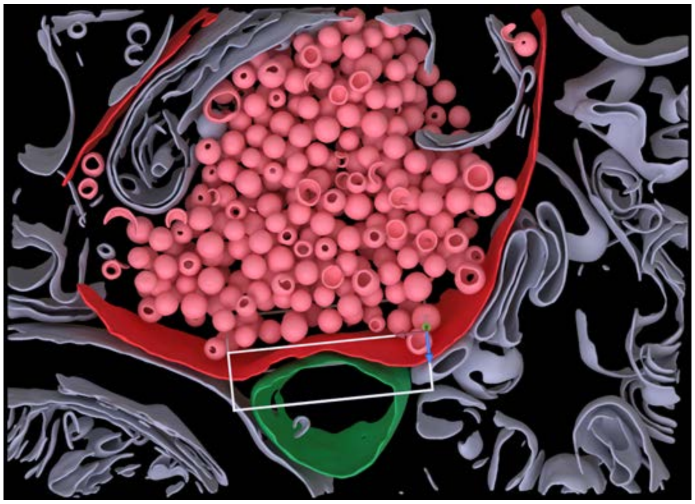

**Cleft MIP**

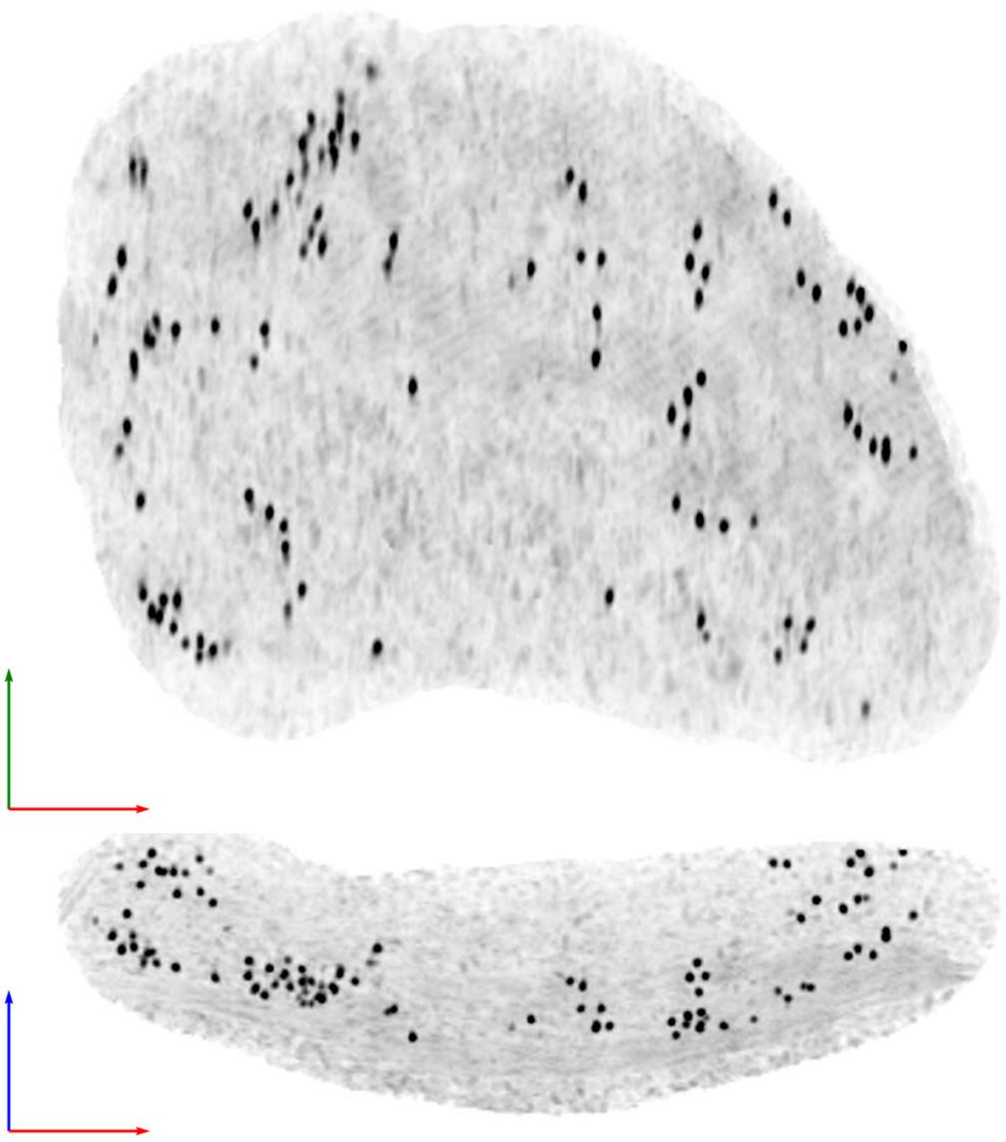

| Tomogram ID: AMmilled54-1_Position_61 | SET: anti-GluA2 15F1 "H4K-2Cys" |
|---|---|
| Cleft ID: 0 | Type: tissue |

**Denoised Tomogram Slice**

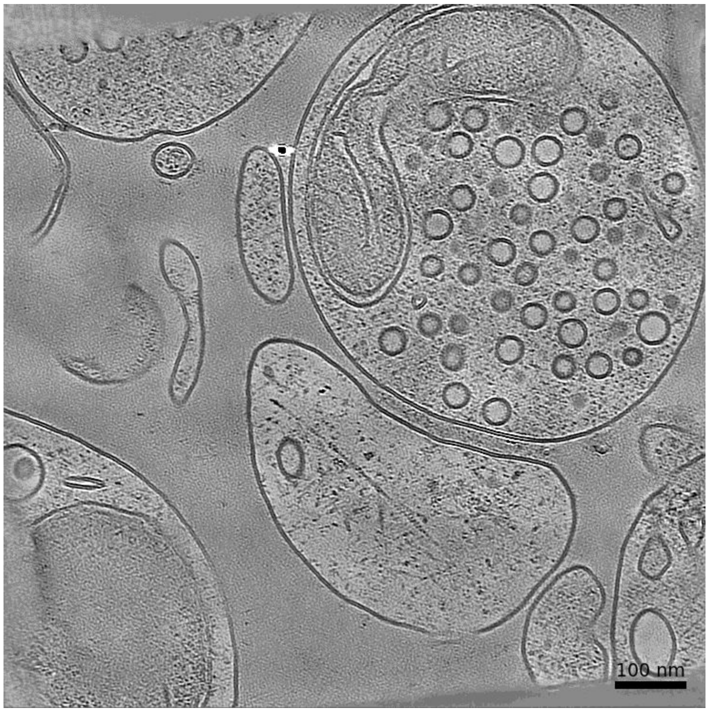


**Membrane Segmentation & Cleft Position**

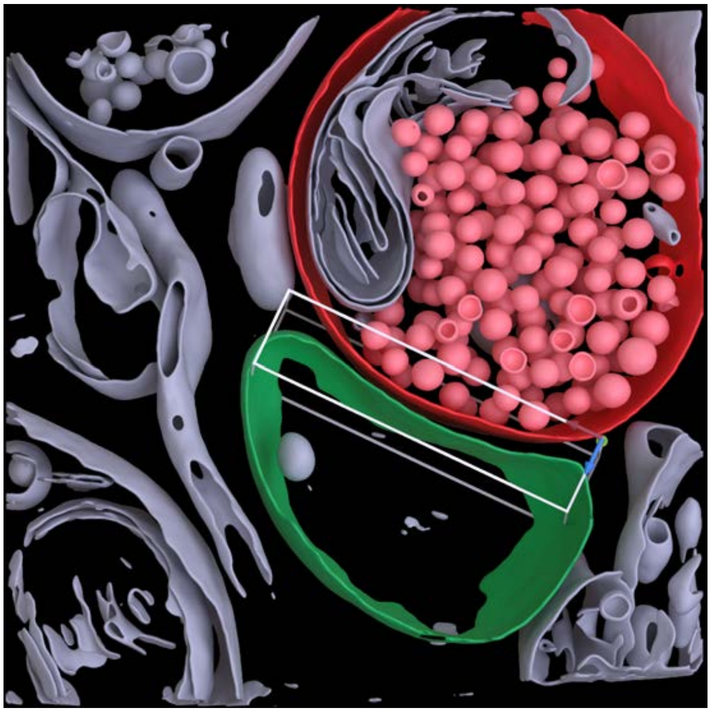

**Cleft MIP**

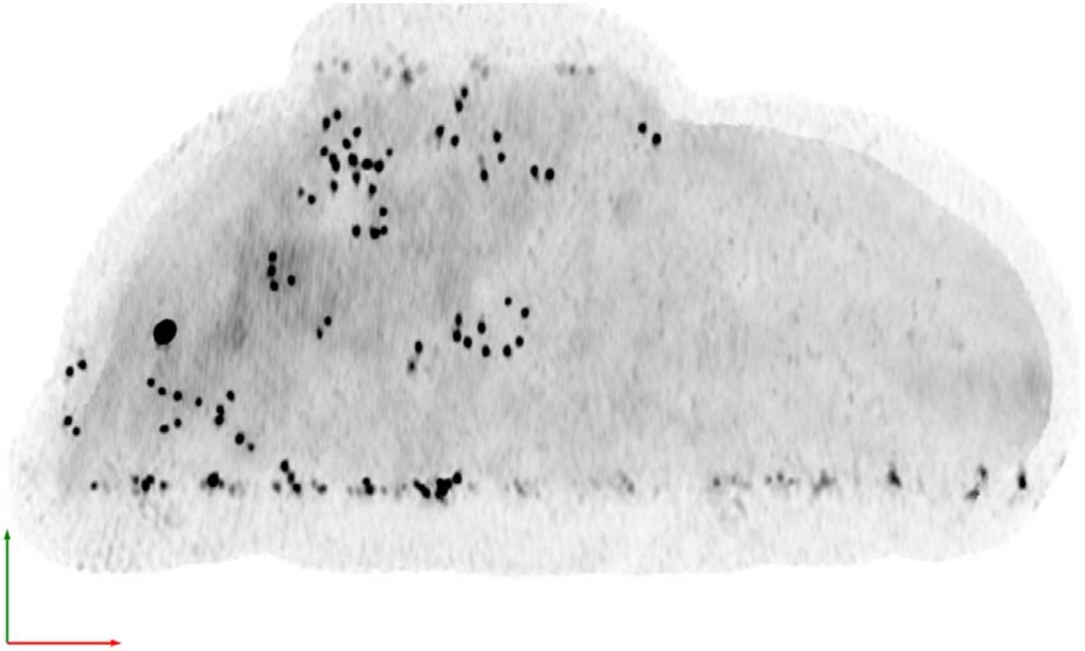

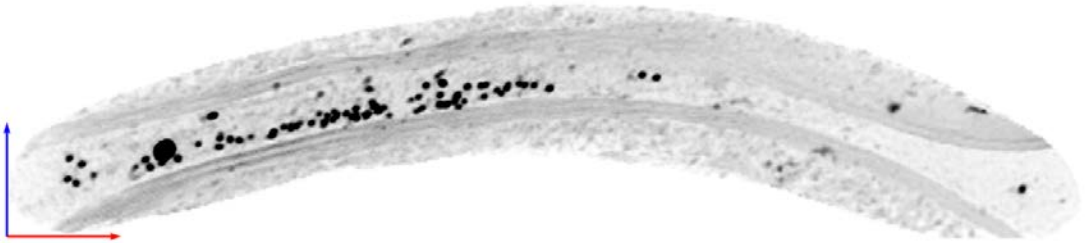

| Tomogram ID: AMmilled55-1_Position_10_3 | SET: anti-GluA2 15F1 "H4K-2Cys" |
|---|---|
| Cleft ID: 0 | Type: tissue |

**Denoised Tomogram Slice**

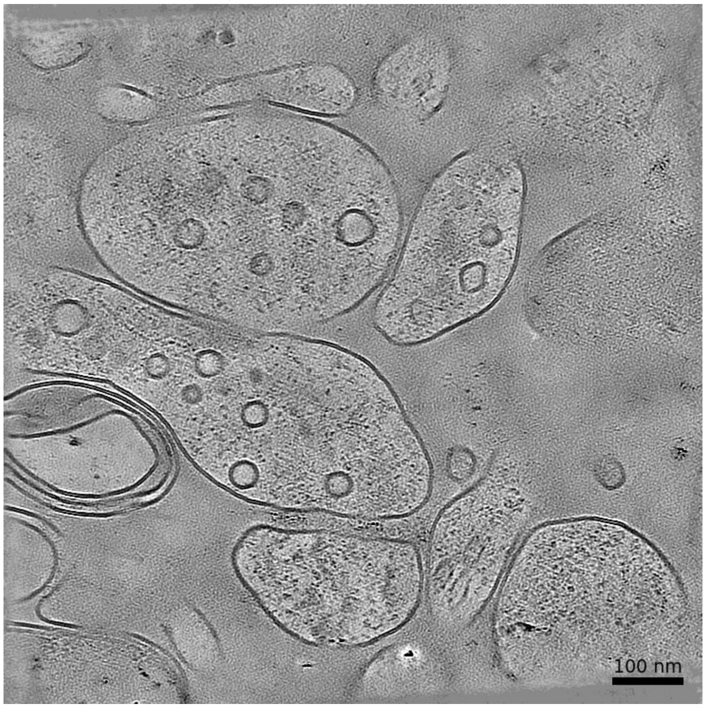


**Membrane Segmentation & Cleft Position**

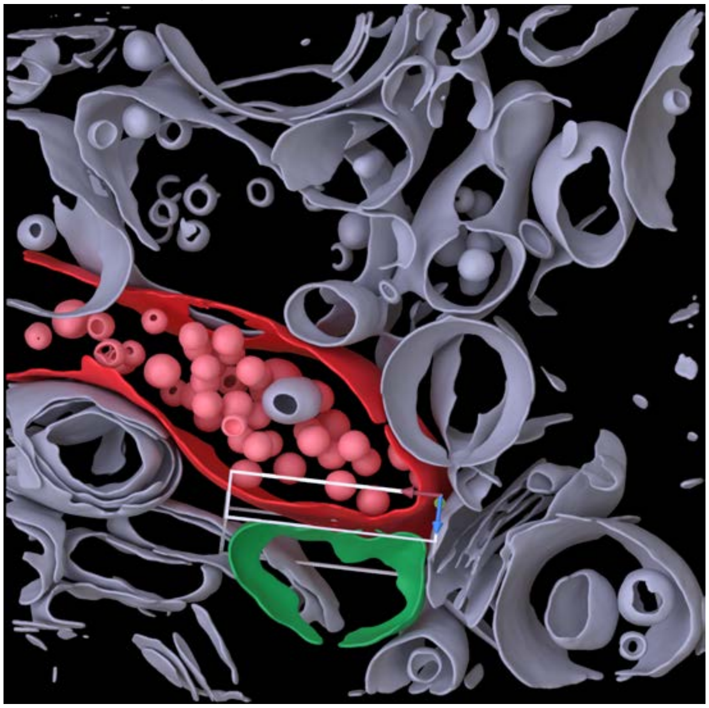

**Cleft MIP**

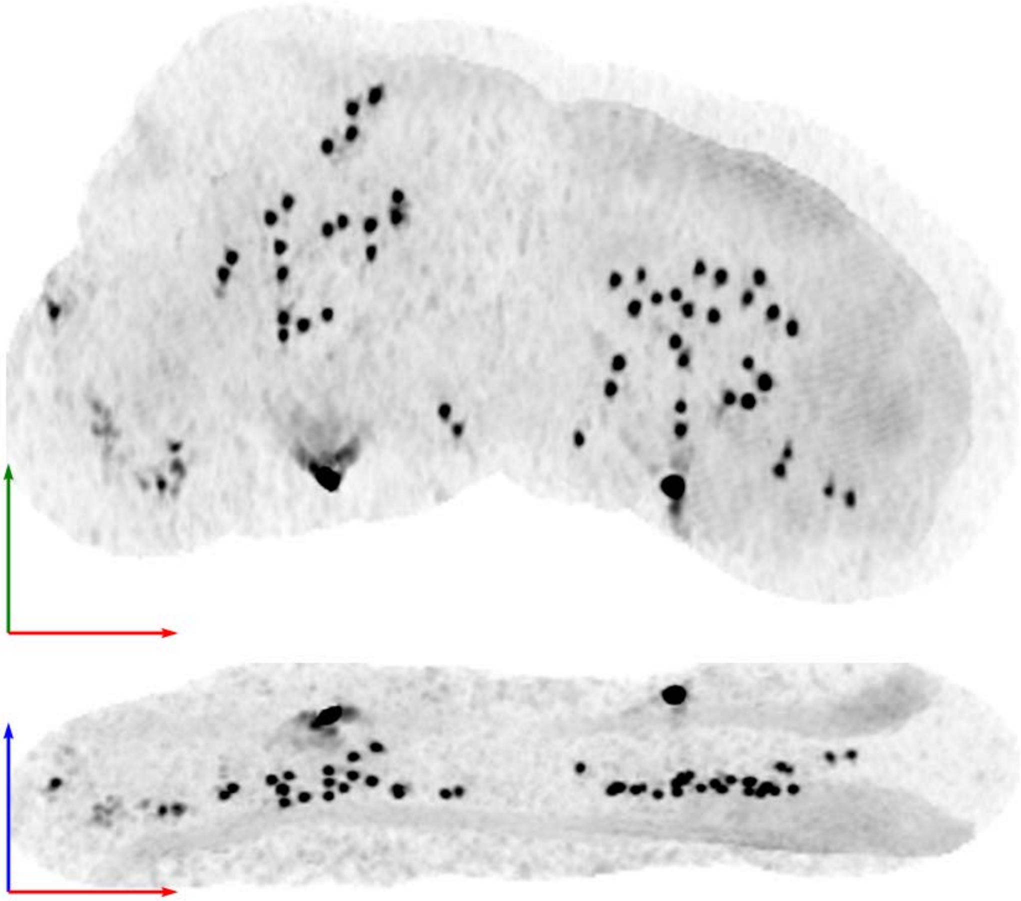

| Tomogram ID: AMmilled55-1_Position_112_2 | SET: anti-GluA2 15F1 "H4K-2Cys" |
|---|---|
| Cleft ID: 0 | Type: tissue |

**Denoised Tomogram Slice**

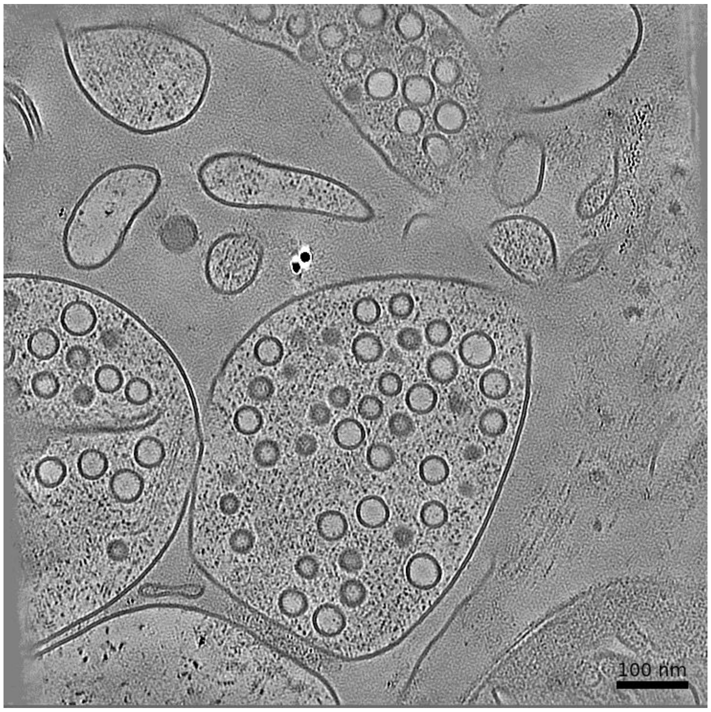


**Membrane Segmentation & Cleft Position**

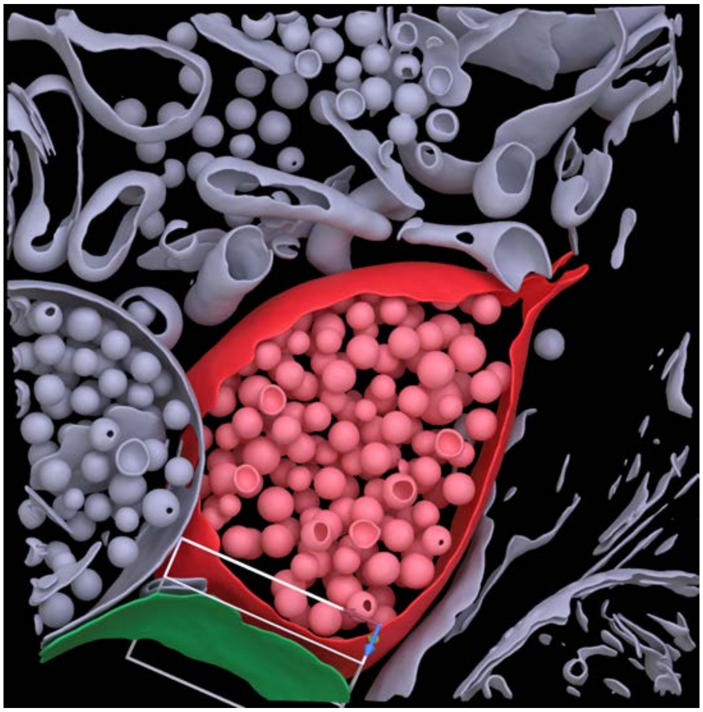

**Cleft MIP**

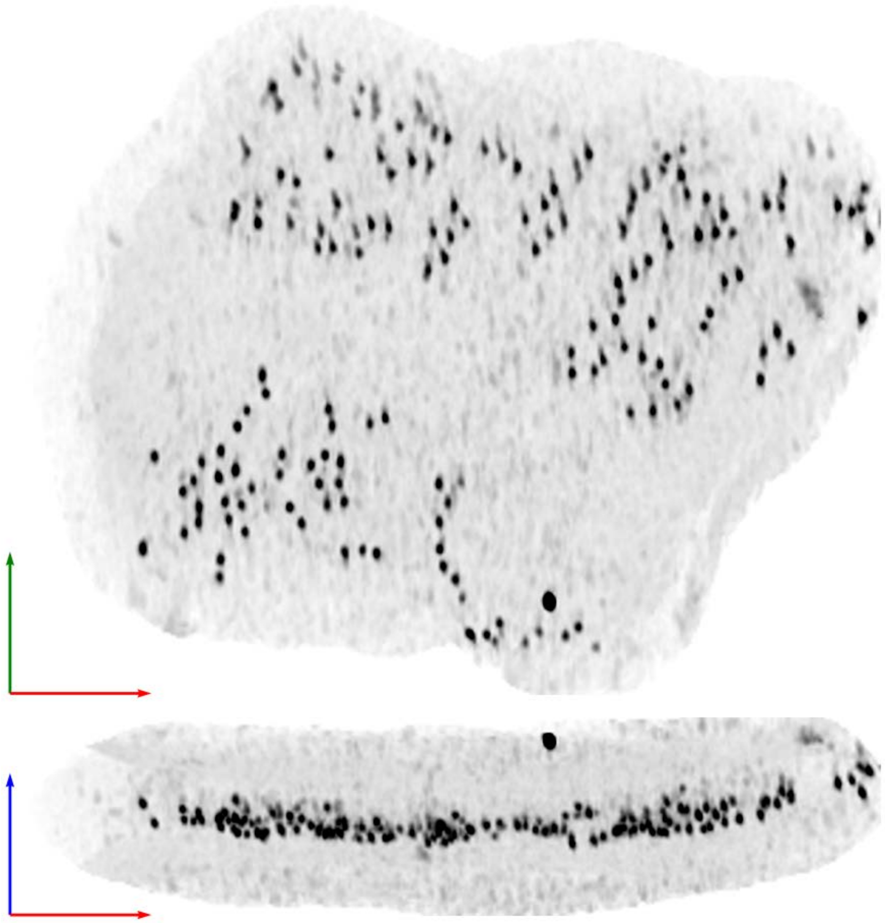

**Tomogram ID: AMmilled55-1_Position_34** **SET: anti-GluA2 15F1 "H4K-2Cys"**

Cleft ID: 0 Type: tissue

### Denoised Tomogram Slice

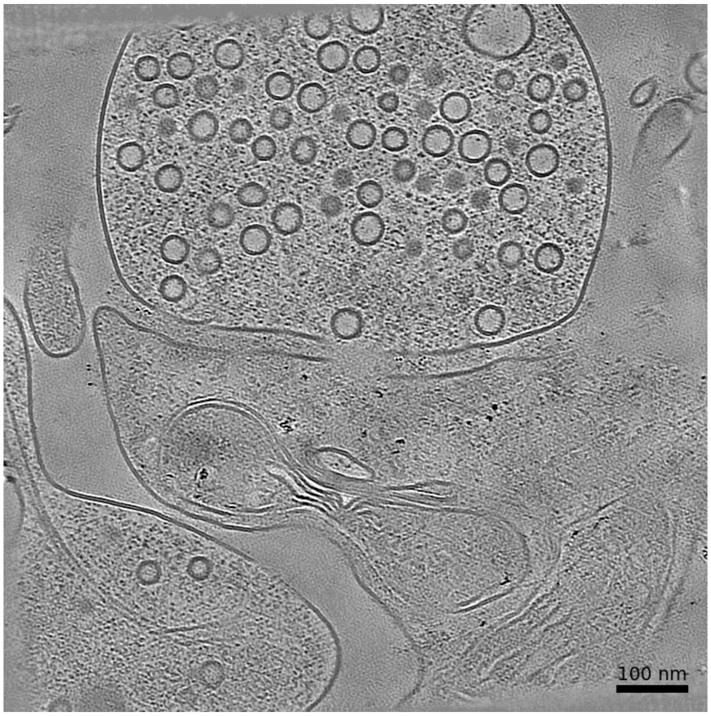


### Membrane Segmentation & Cleft Position

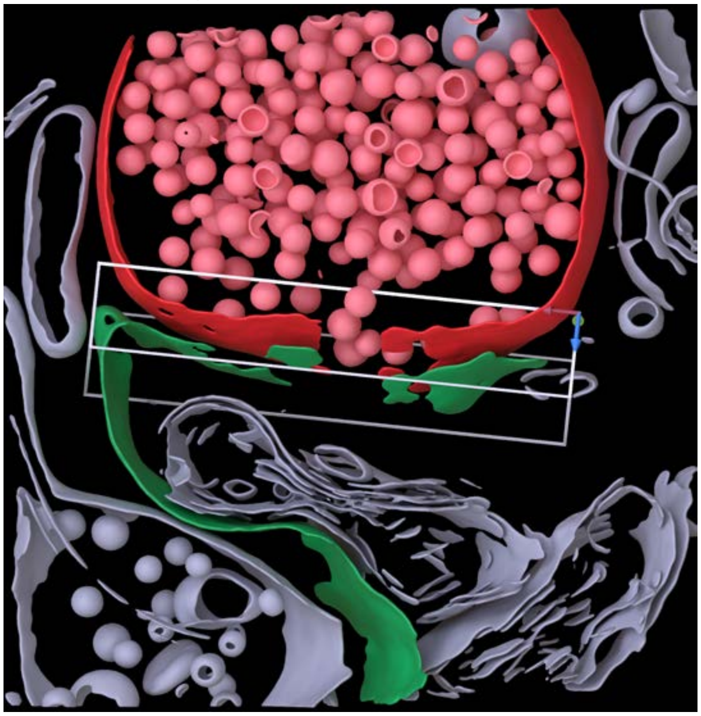

### Cleft MIP

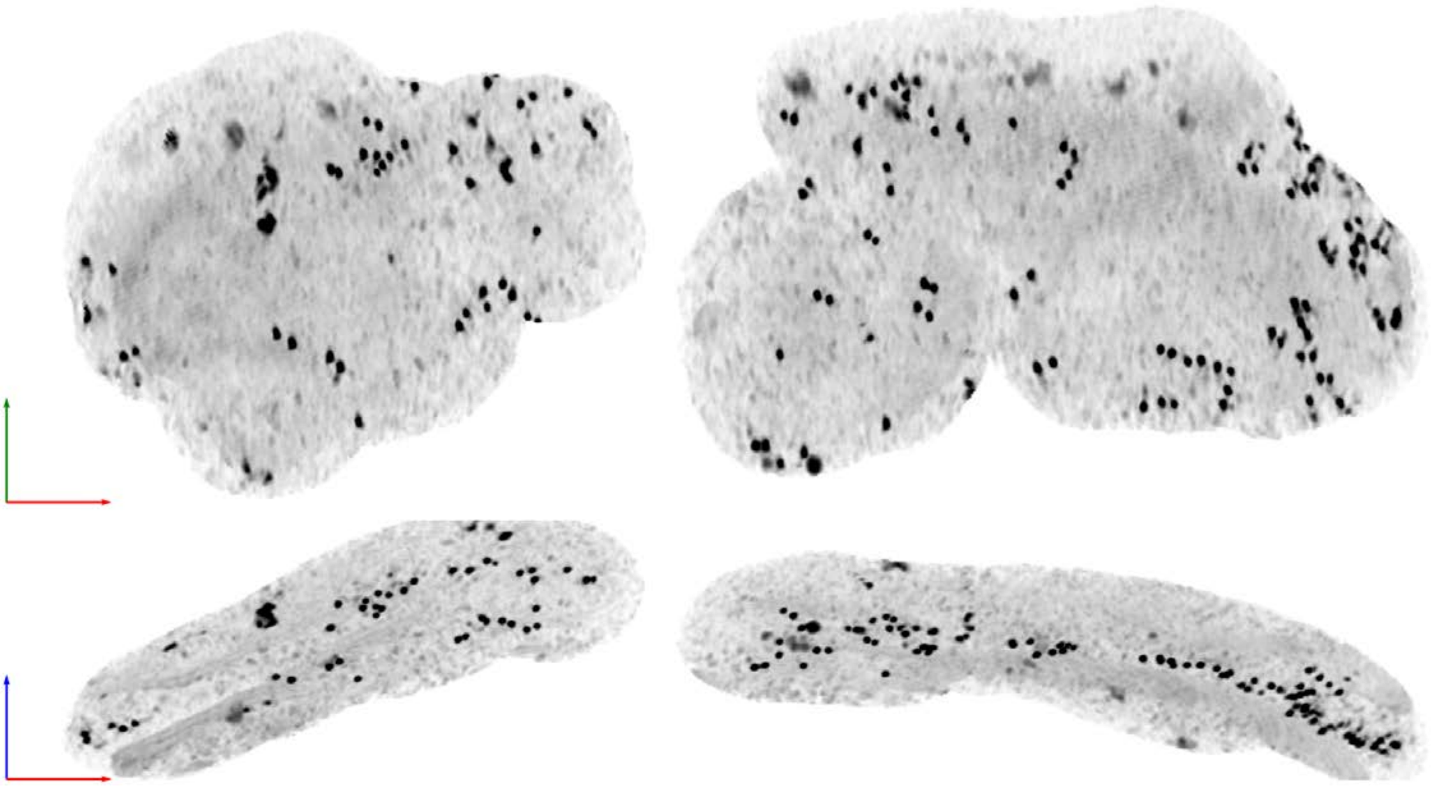

| Tomogram ID: AMmilled55-1_Position_35_2 | SET: anti-GluA2 15F1 "H4K-2Cys" |
| --- | --- |
| Cleft ID: 0 | Type: tissue |

**Denoised Tomogram Slice**

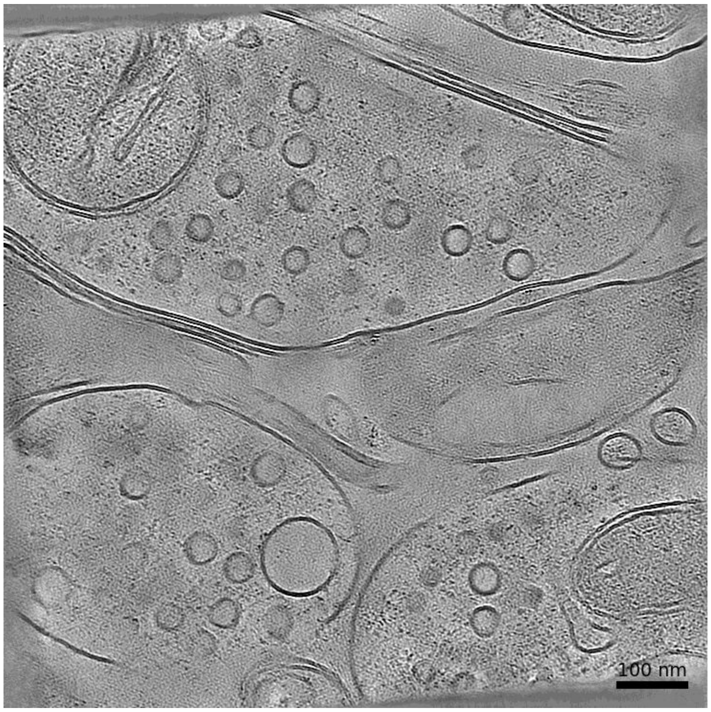


**Membrane Segmentation & Cleft Position**

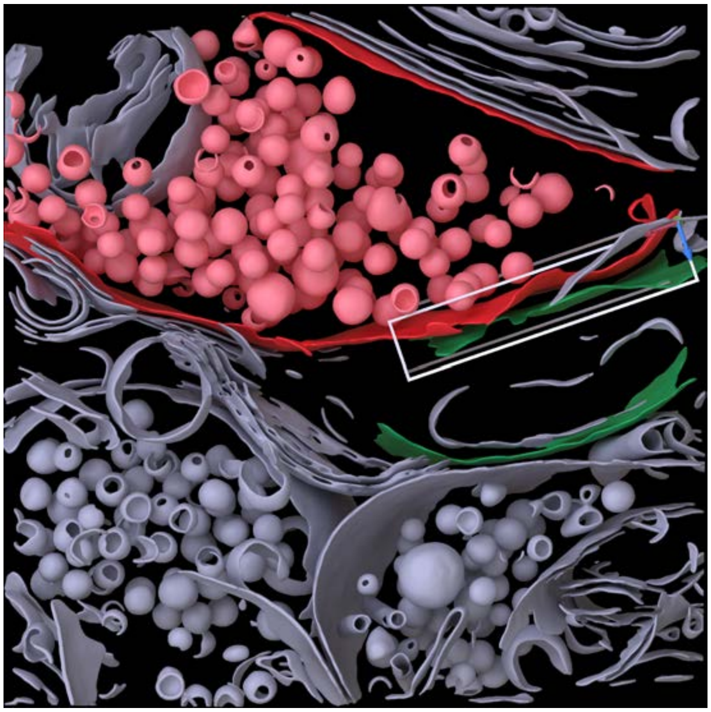

**Cleft MIP**

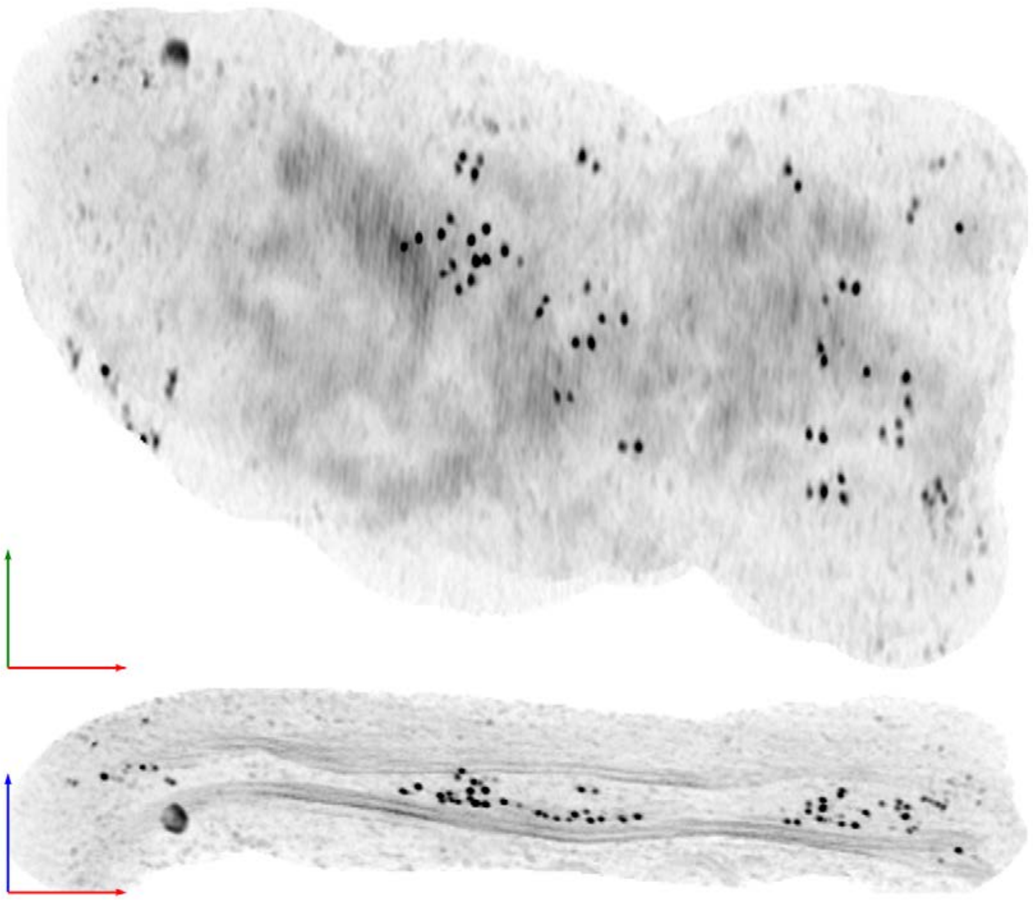

**Tomogram ID: AMmilled55-1_Position_43** **SET: anti-GluA2 15F1 "H4K-2Cys"**

Cleft ID: 0 Type: tissue

### Denoised Tomogram Slice

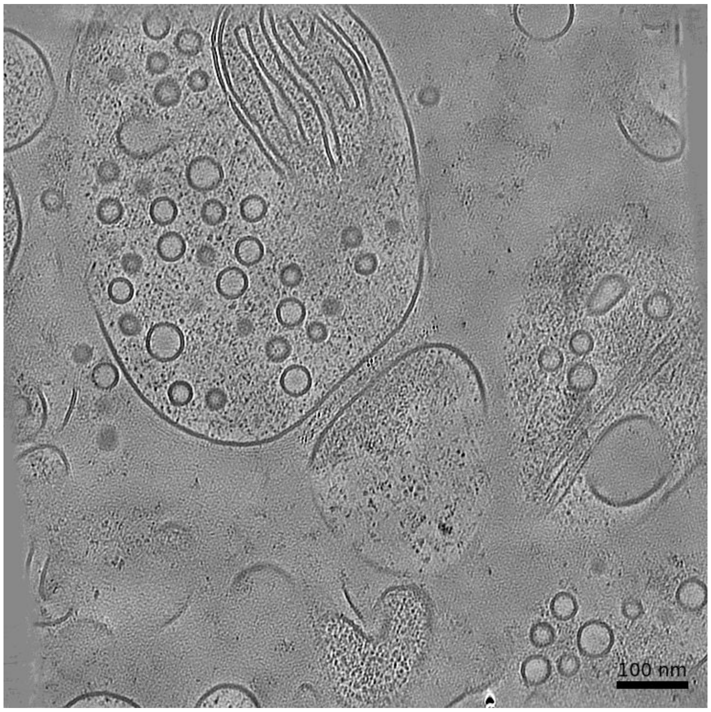


### Membrane Segmentation & Cleft Position

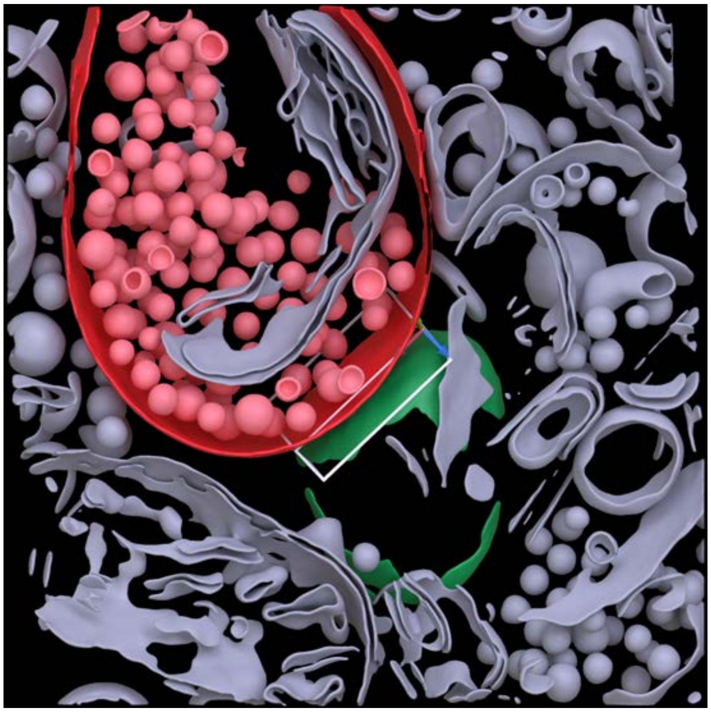

### Cleft MIP

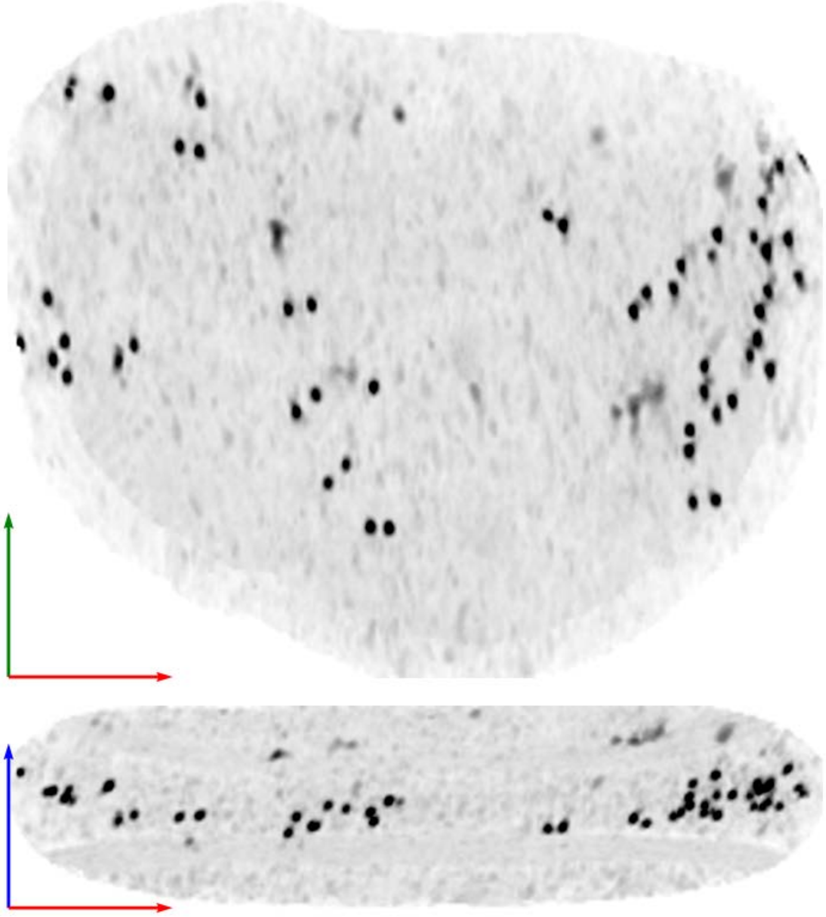

| **Tomogram ID: AMmilled55-1_Position_47** | **SET: anti-GluA2 15F1 "H4K-2Cys"** |
|---|---|
| Cleft ID: 0 | Type: tissue |

### Denoised Tomogram Slice

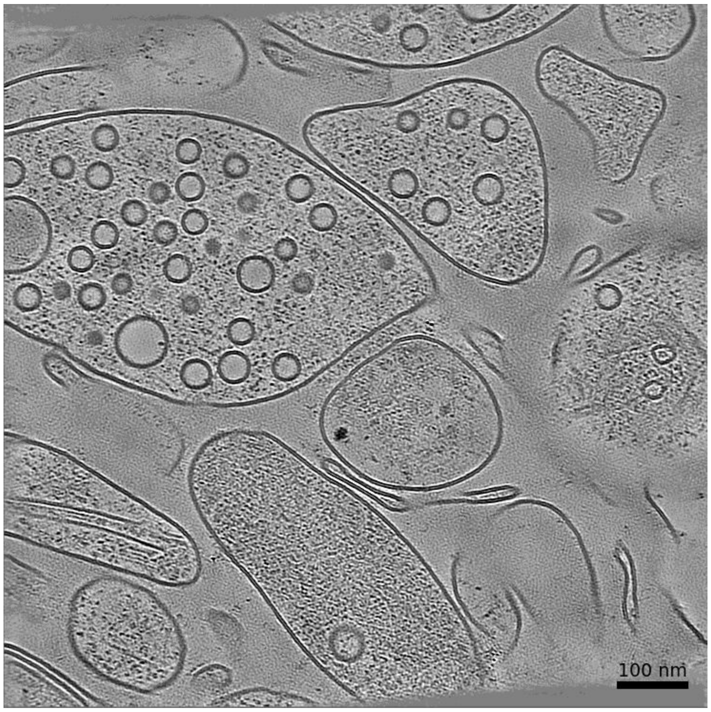


### Membrane Segmentation & Cleft Position

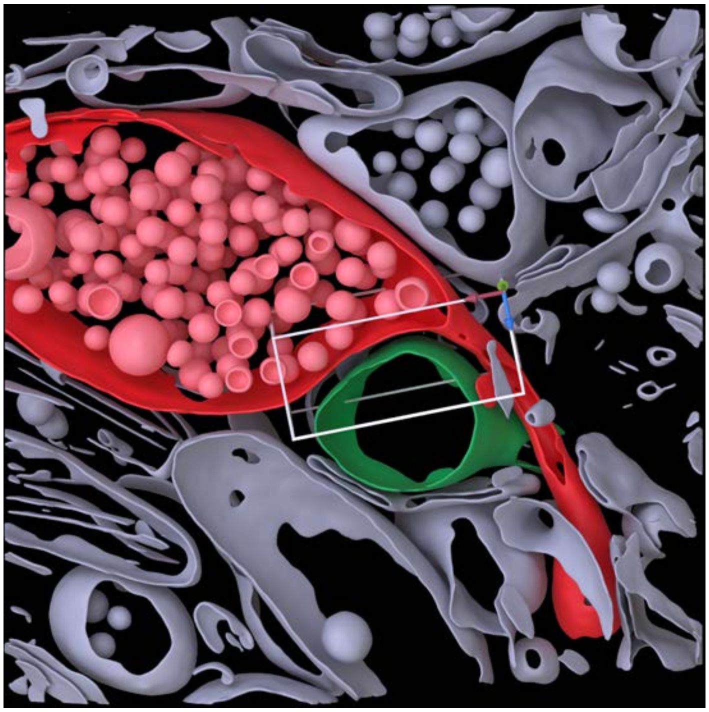

### Cleft MIP

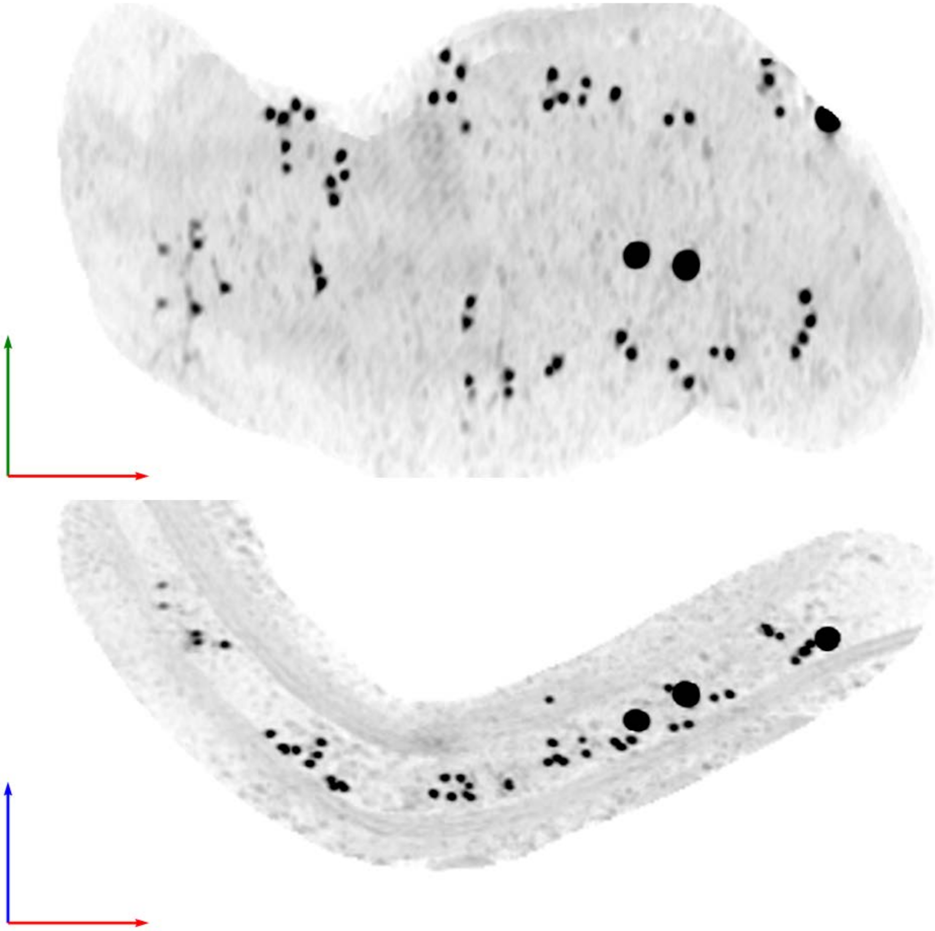

| Tomogram ID: AMmilled55-1_Position_49 | SET: anti-GluA2 15F1 "H4K-2Cys" |
|---|---|
| Cleft ID: 0 | Type: tissue |

**Denoised Tomogram Slice**

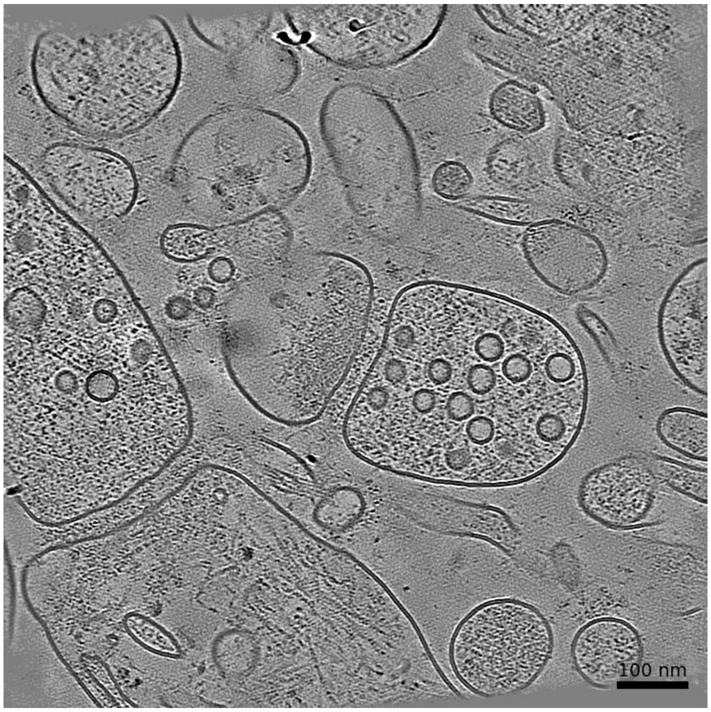


**Membrane Segmentation & Cleft Position**

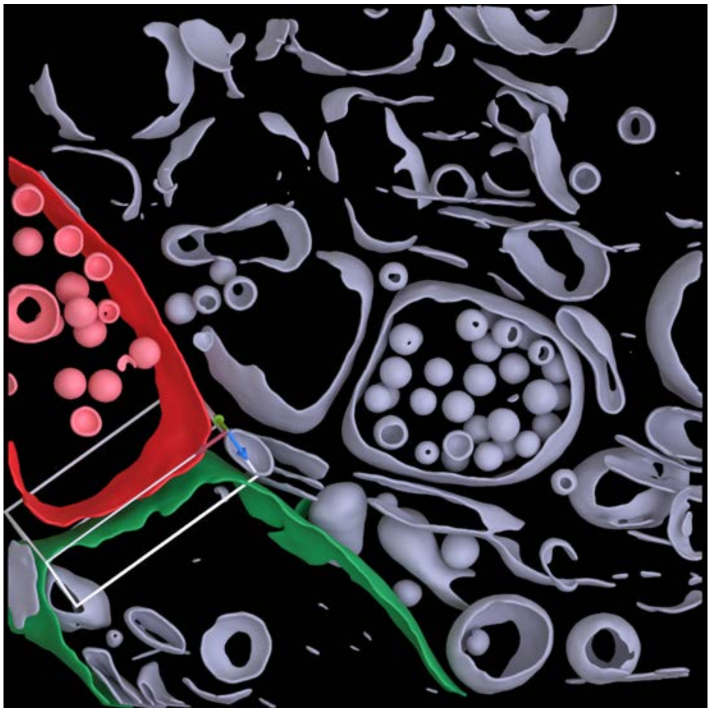

**Cleft MIP**

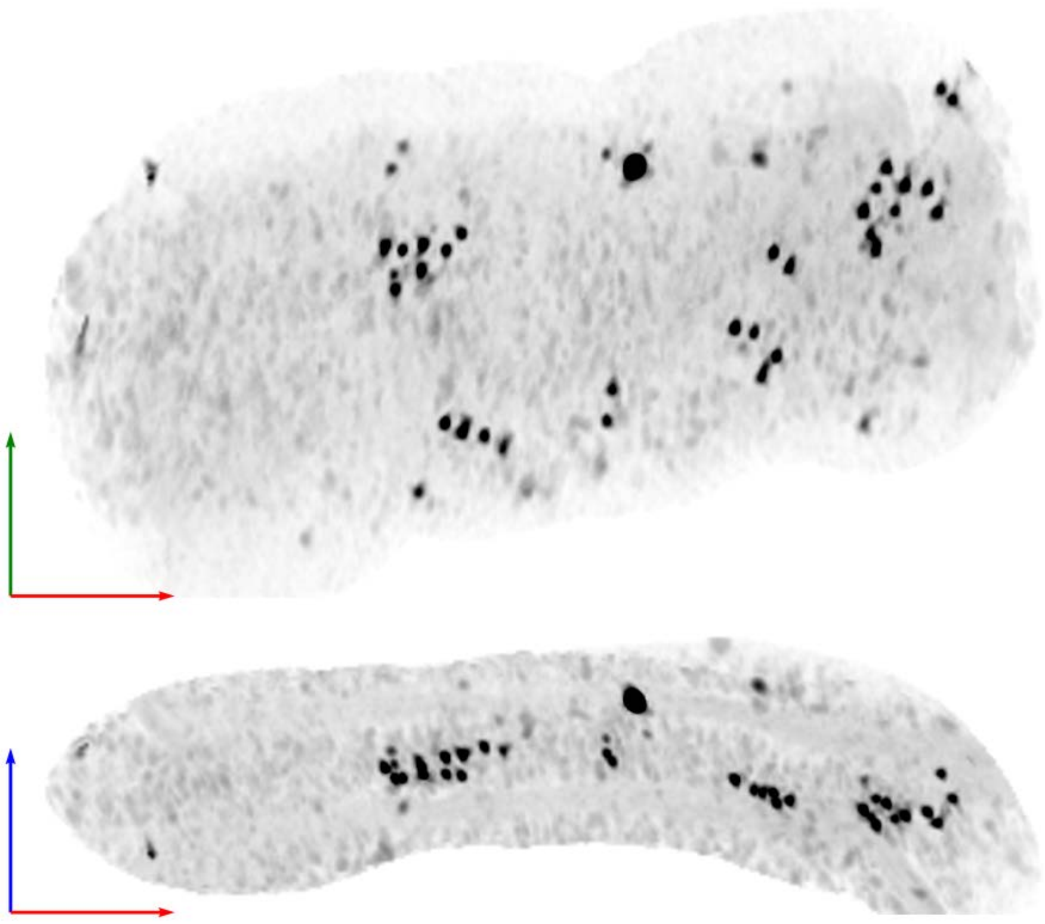

**Tomogram ID: AMmilled55-1_Position_49** | **SET: anti-GluA2 15F1 "H4K-2Cys"**

Cleft ID: 0 | Type: tissue

### Denoised Tomogram Slice

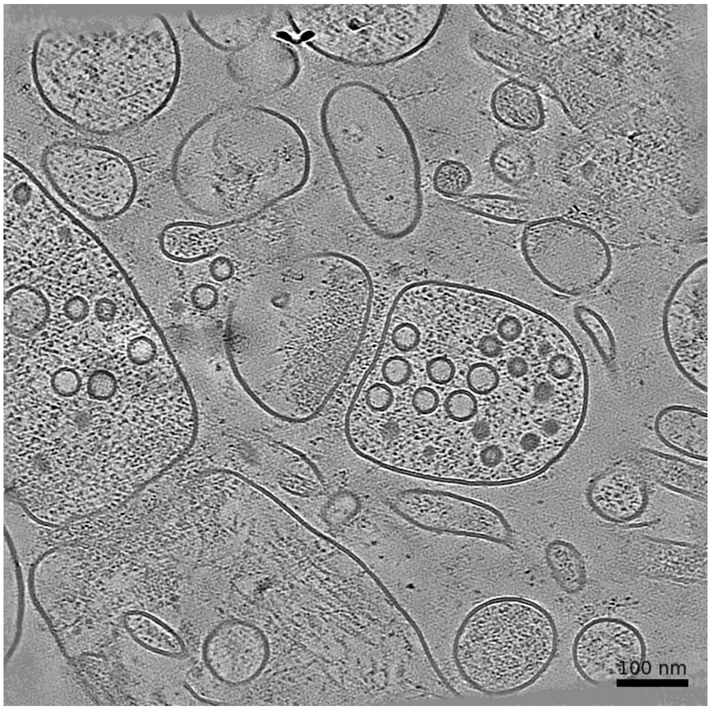


### Membrane Segmentation & Cleft Position

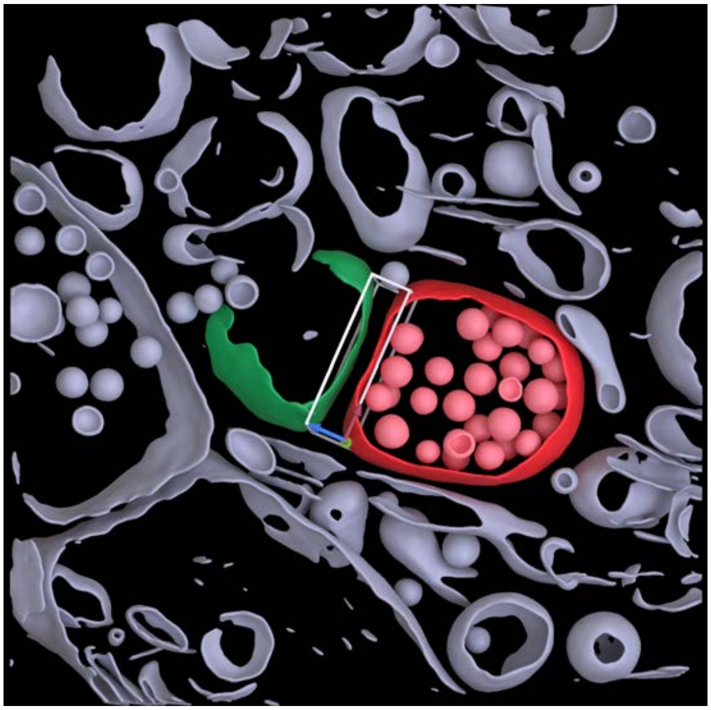

### Cleft MIP

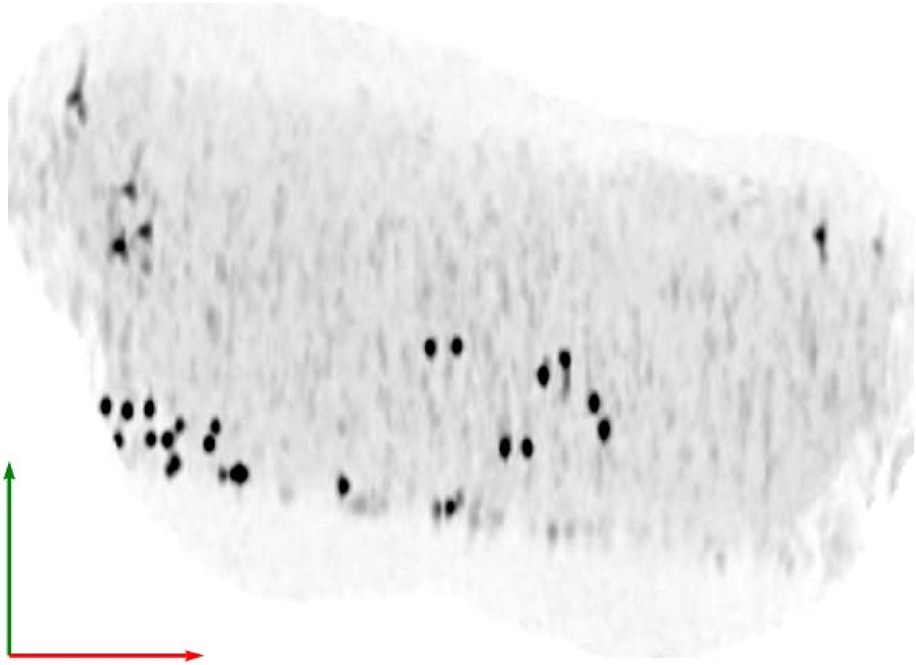

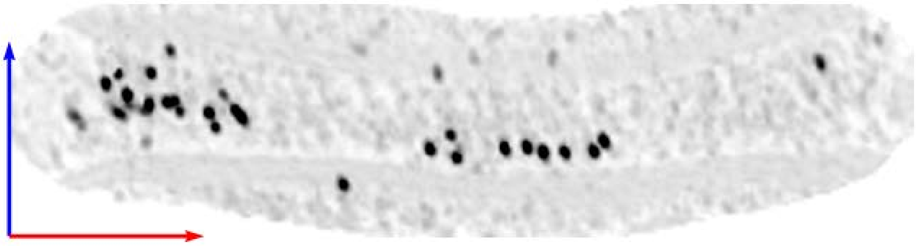

**Tomogram ID: AMmilled55-1_Position_75** **SET: anti-GluA2 15F1 "H4K-2Cys"**

Cleft ID: 0 Type: tissue

**Denoised Tomogram Slice**

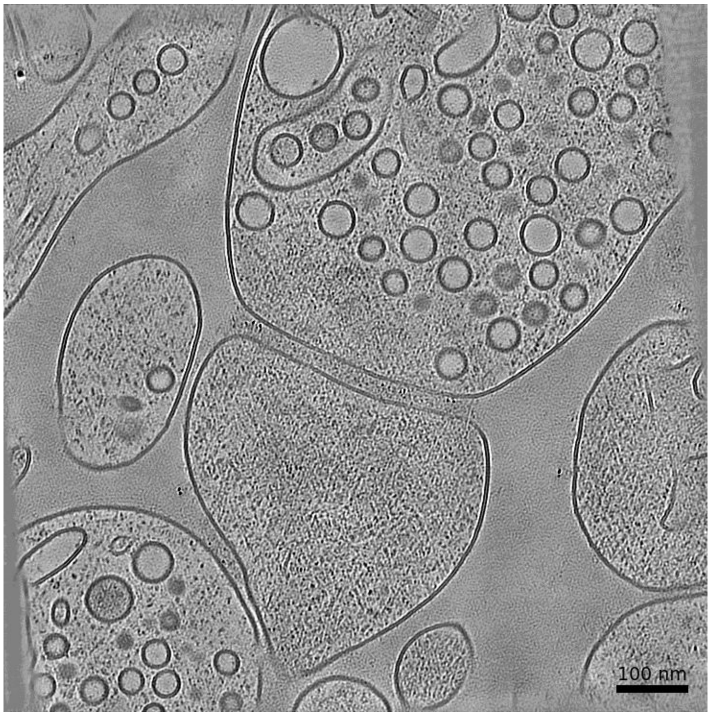


**Membrane Segmentation & Cleft Position**

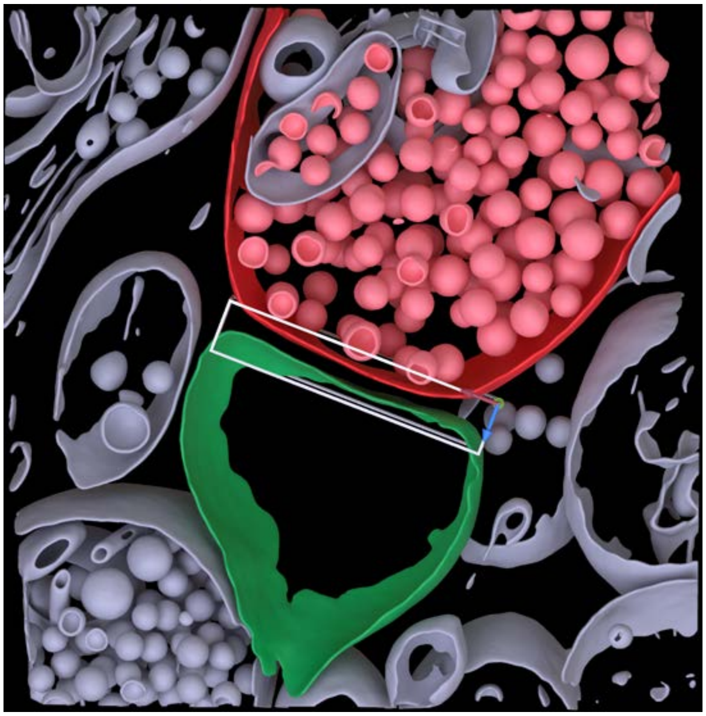

**Cleft MIP**

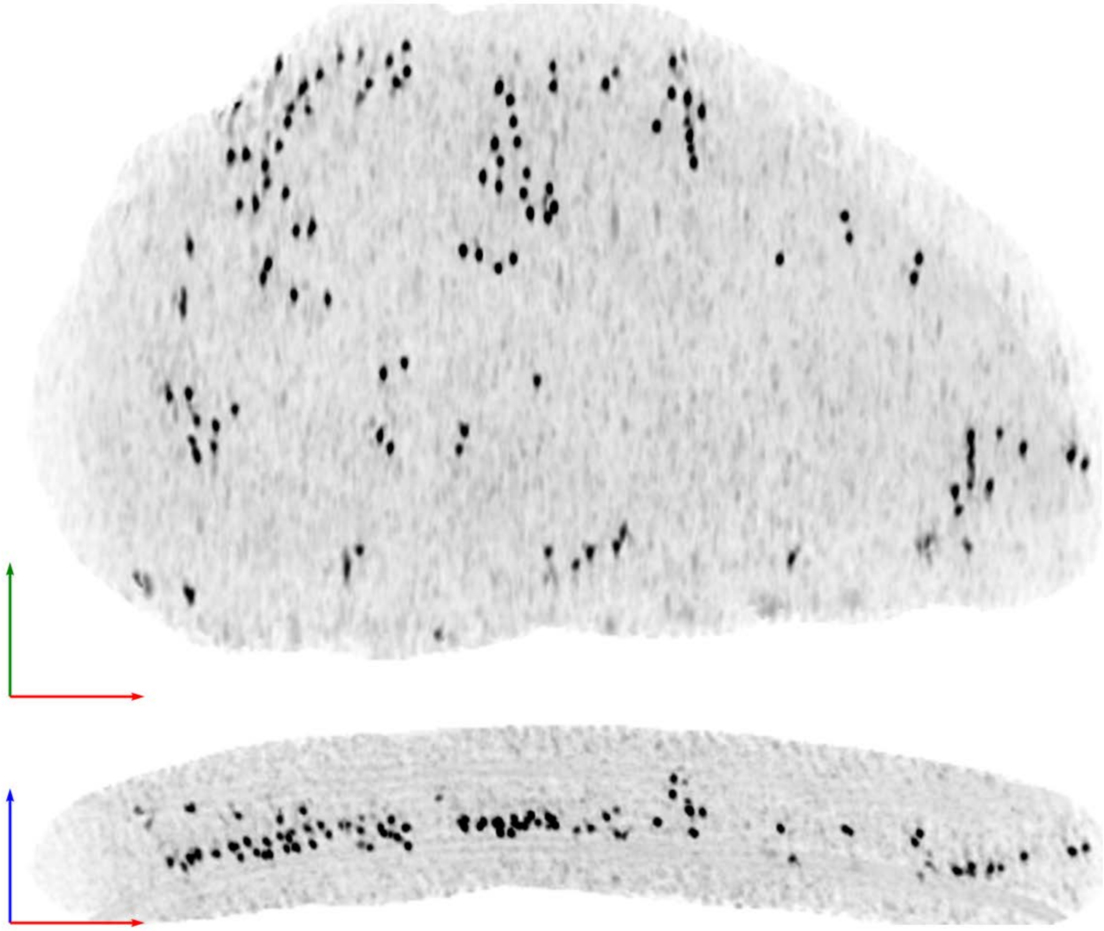

**Tomogram ID: AMmilled55-1_Position_75** | **SET: anti-GluA2 15F1 "H4K-2Cys"**

Cleft ID: 0 | Type: tissue

### Denoised Tomogram Slice

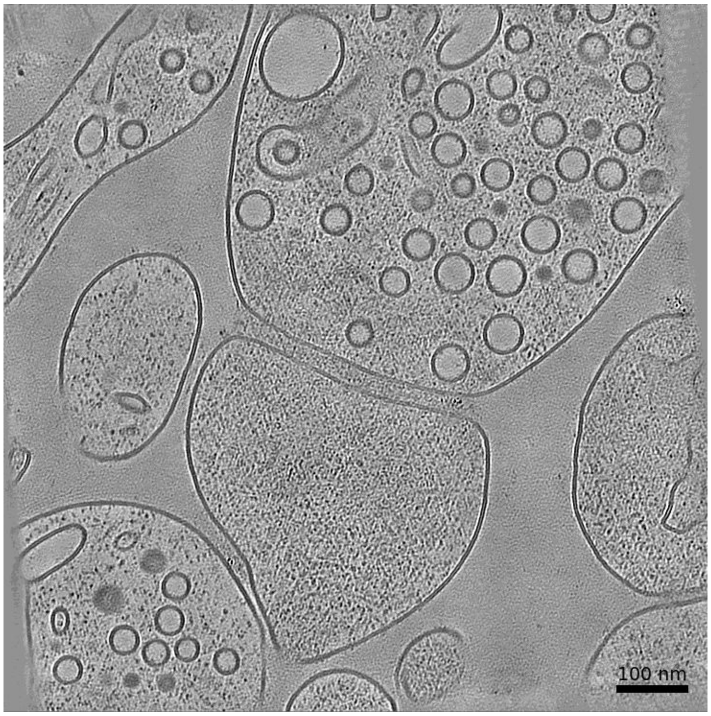


### Membrane Segmentation & Cleft Position

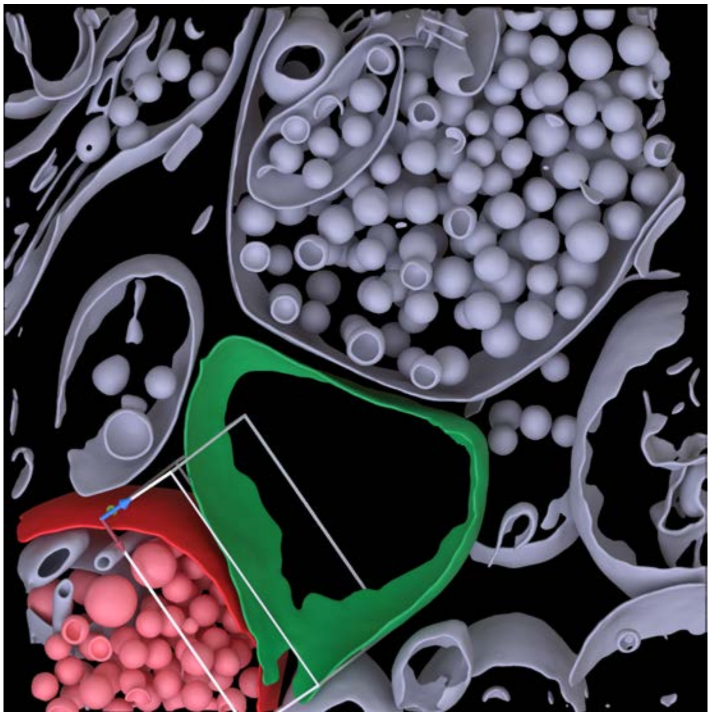

### Cleft MIP

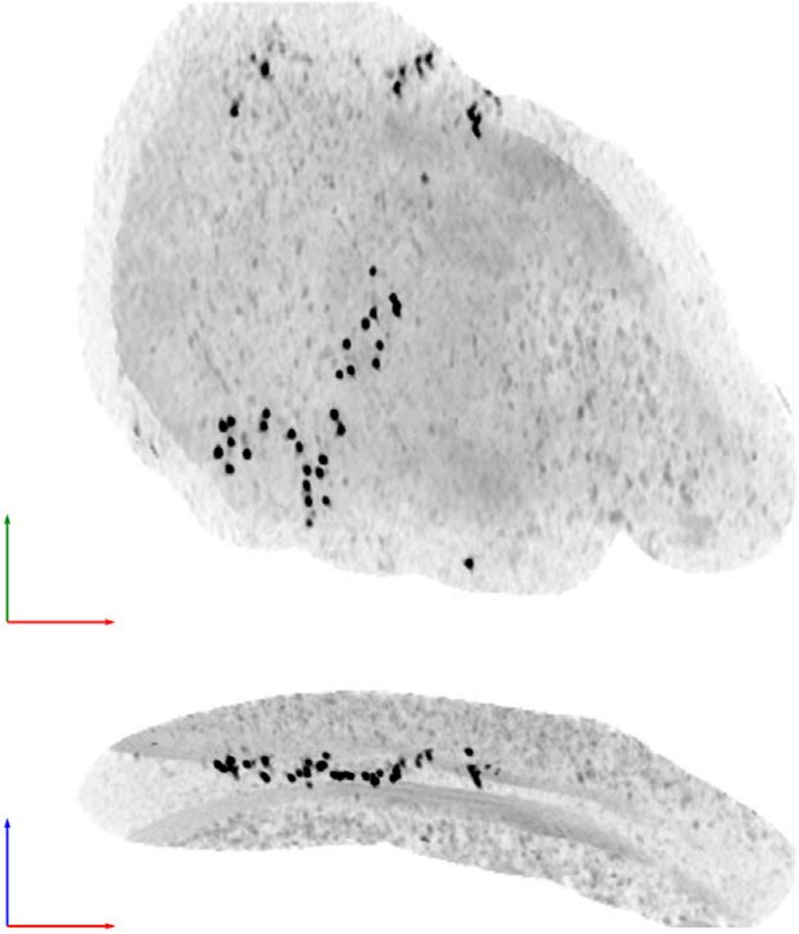

**Tomogram ID: AMmilled55-1_Position_82** **SET: anti-GluA2 15F1 "H4K-2Cys"**

Cleft ID: 0 Type: tissue

### Denoised Tomogram Slice

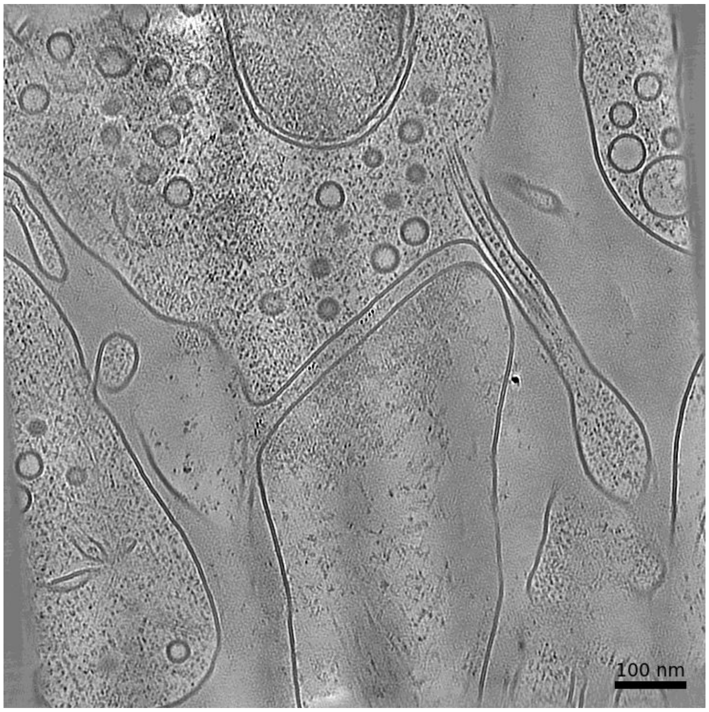


### Membrane Segmentation & Cleft Position

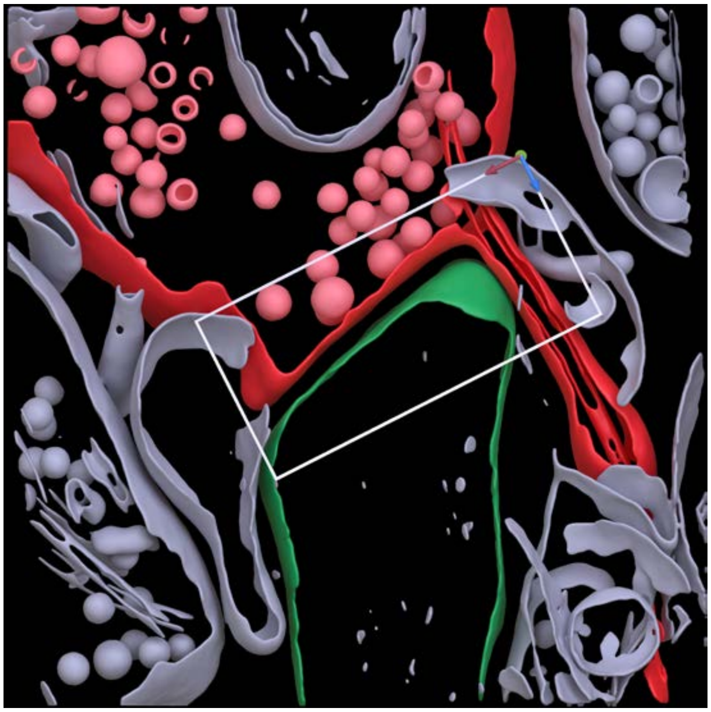

### Cleft MIP

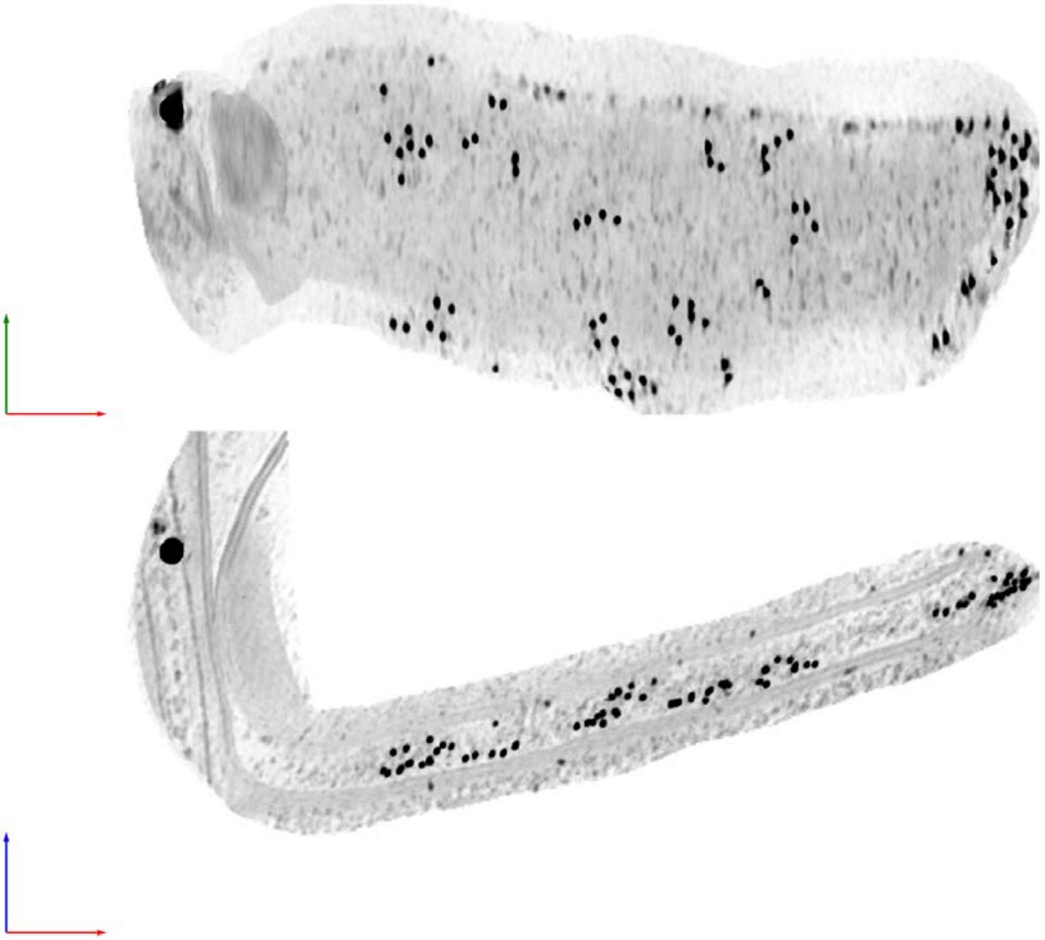

| Tomogram ID: AMmilled55-1_Position_82 | SET: anti-GluA2 15F1 "H4K-2Cys" |
| --- | --- |
| Cleft ID: 0 | Type: tissue |

**Denoised Tomogram Slice**

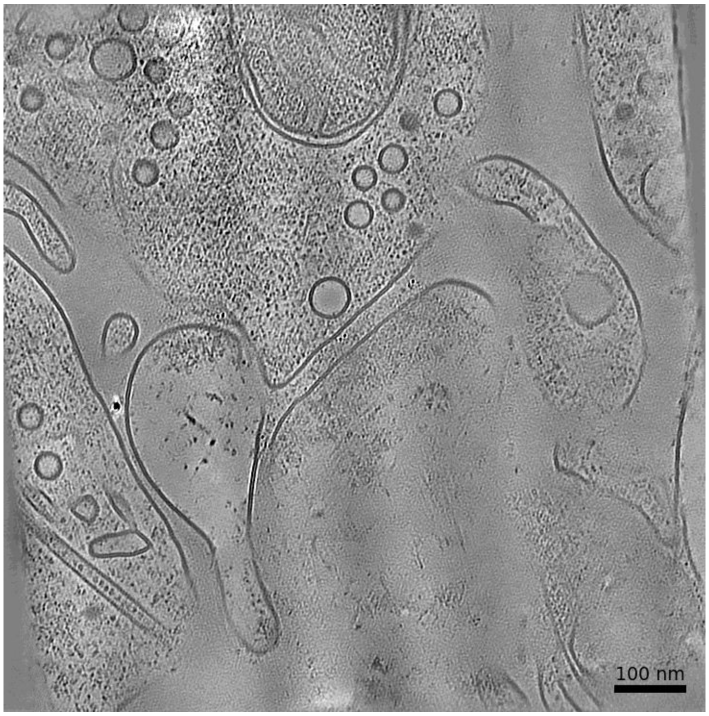


**Membrane Segmentation & Cleft Position**

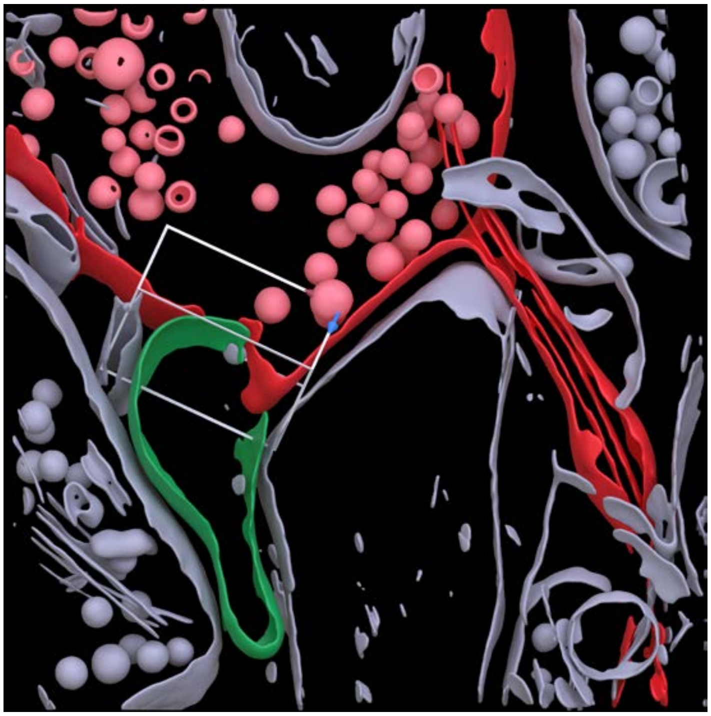

**Cleft MIP**

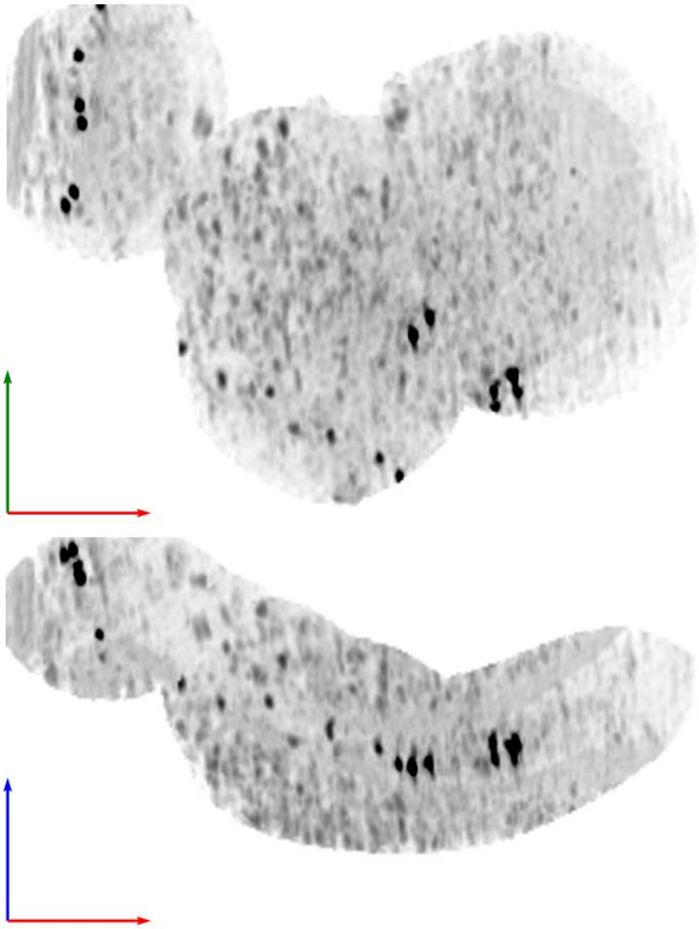

**Tomogram ID: 20250206_AMmilled27-2_100** **SET: anti-GluA1 11B8 "C-term"**

Cleft ID: 0 Type: semi-tissue

### Denoised Tomogram Slice

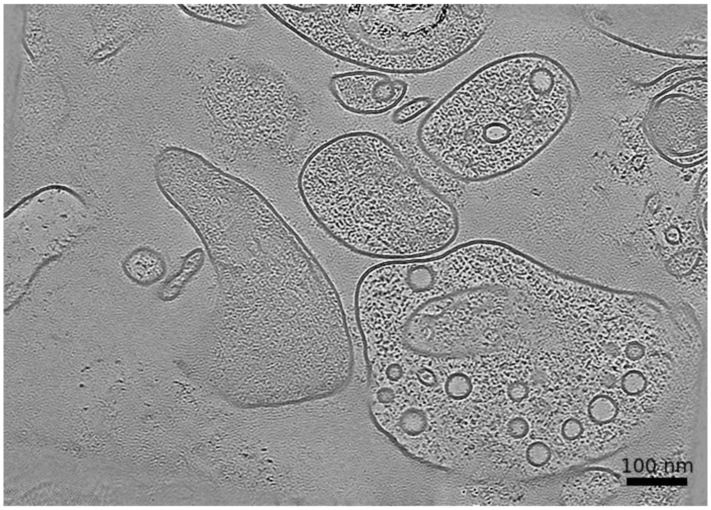


### Membrane Segmentation & Cleft Position

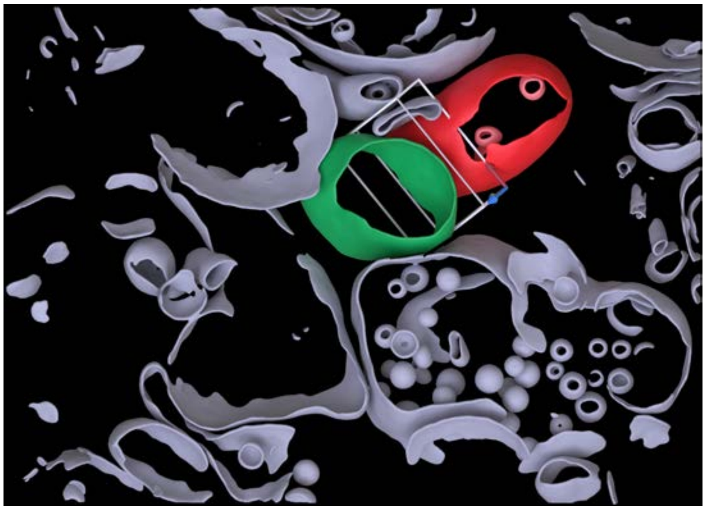

### Cleft MIP

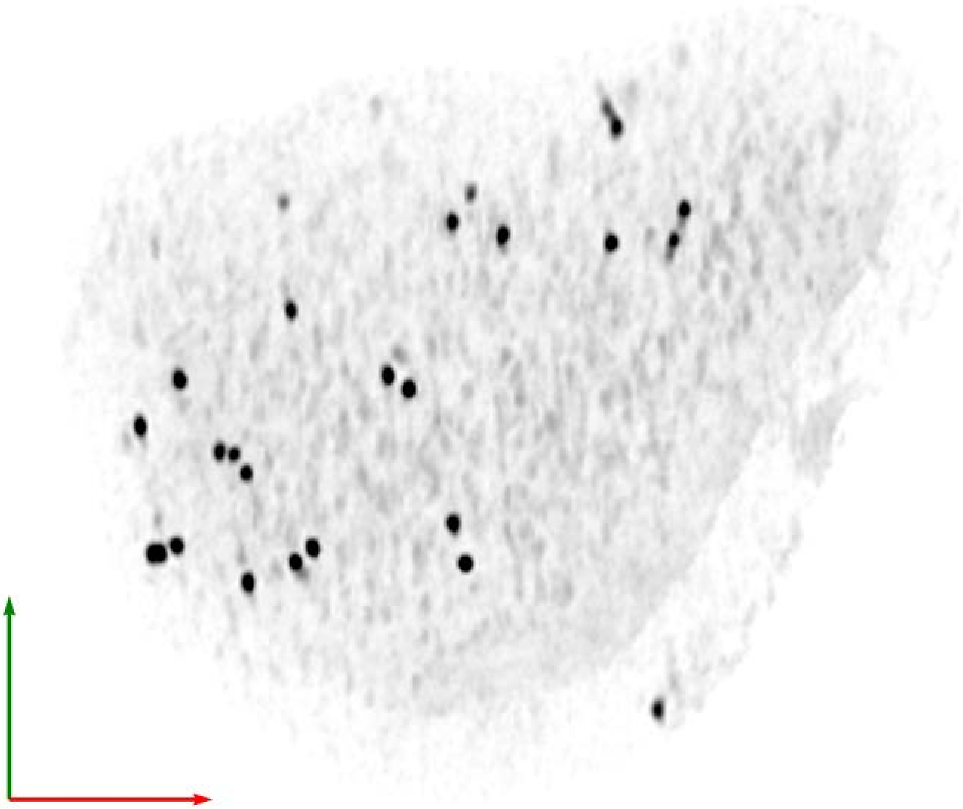

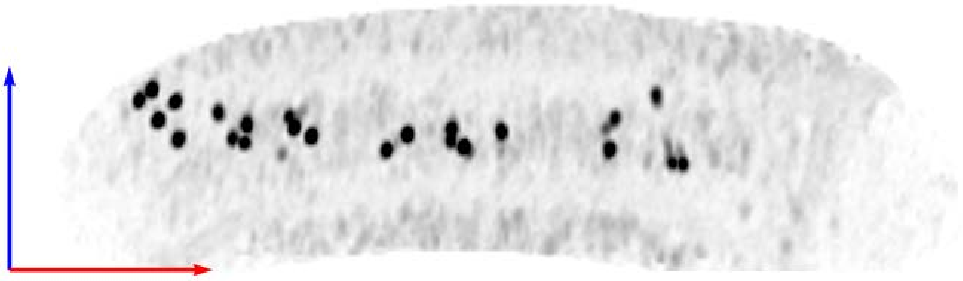

**Tomogram ID: 20250206_AMmilled27-2_100** **SET: anti-GluA1 11B8 "C-term"**

Cleft ID: 0 Type: semi-tissue

**Denoised Tomogram Slice**

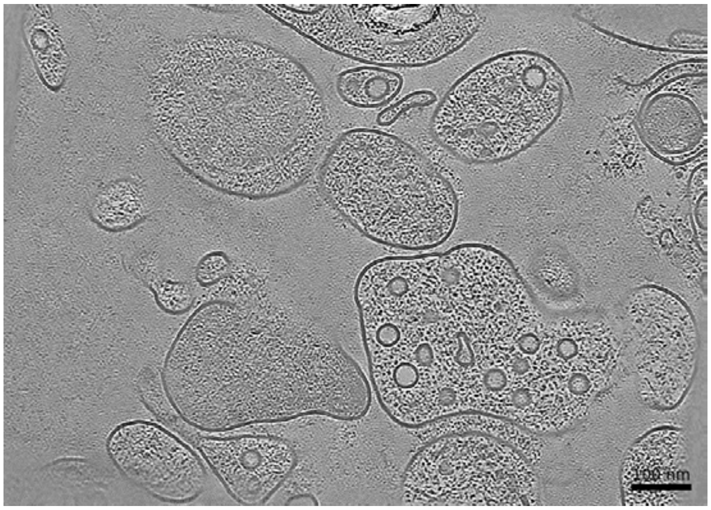


**Membrane Segmentation & Cleft Position**

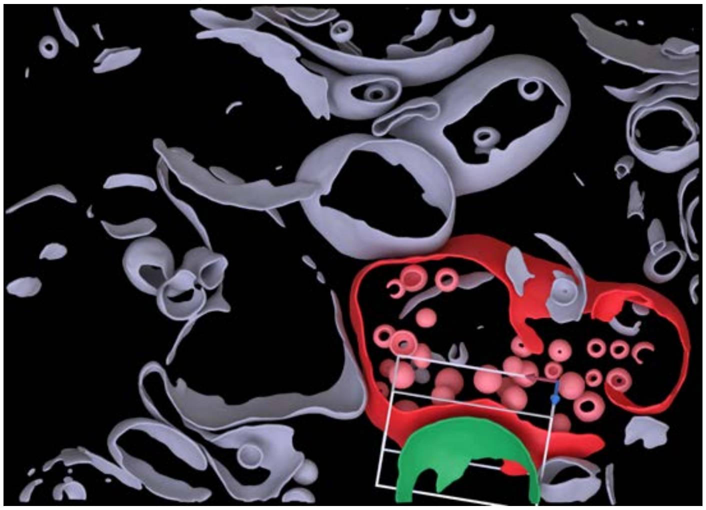

**Cleft MIP**

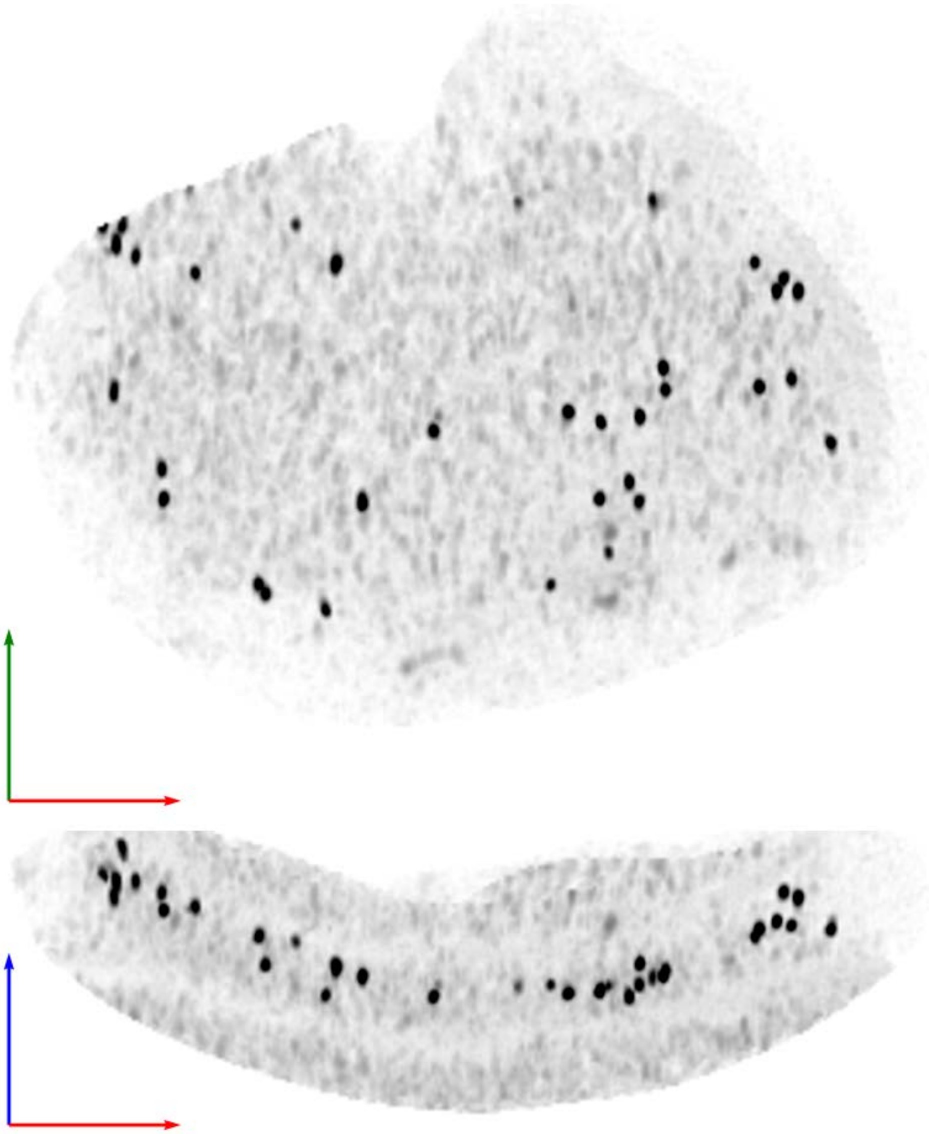

| Tomogram ID: 20250206_AMmilled27-2_101 | SET: anti-GluA1 11B8 "C-term" |
| --- | --- |
| Cleft ID: 0 | Type: synaptosome |

**Denoised Tomogram Slice**

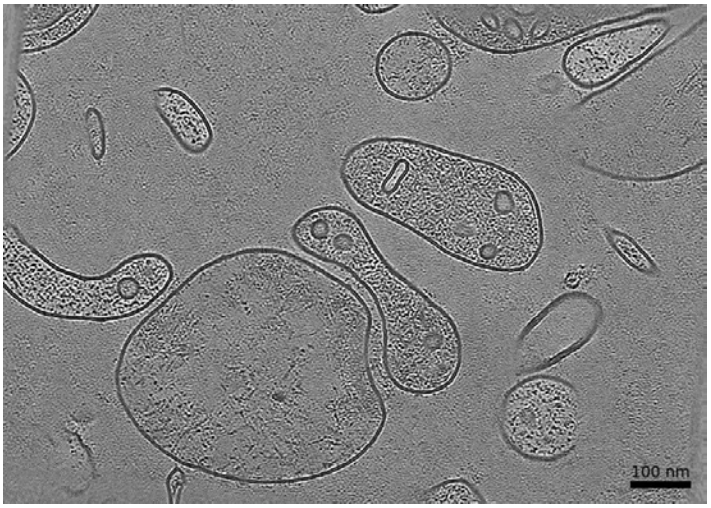


**Membrane Segmentation & Cleft Position**

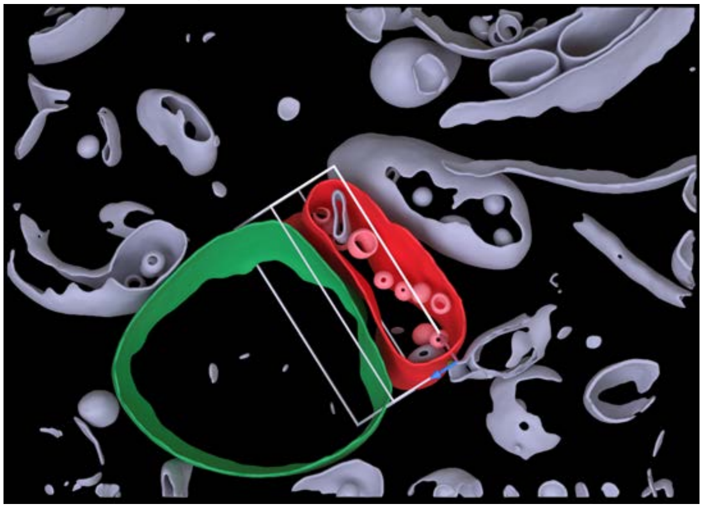

**Cleft MIP**

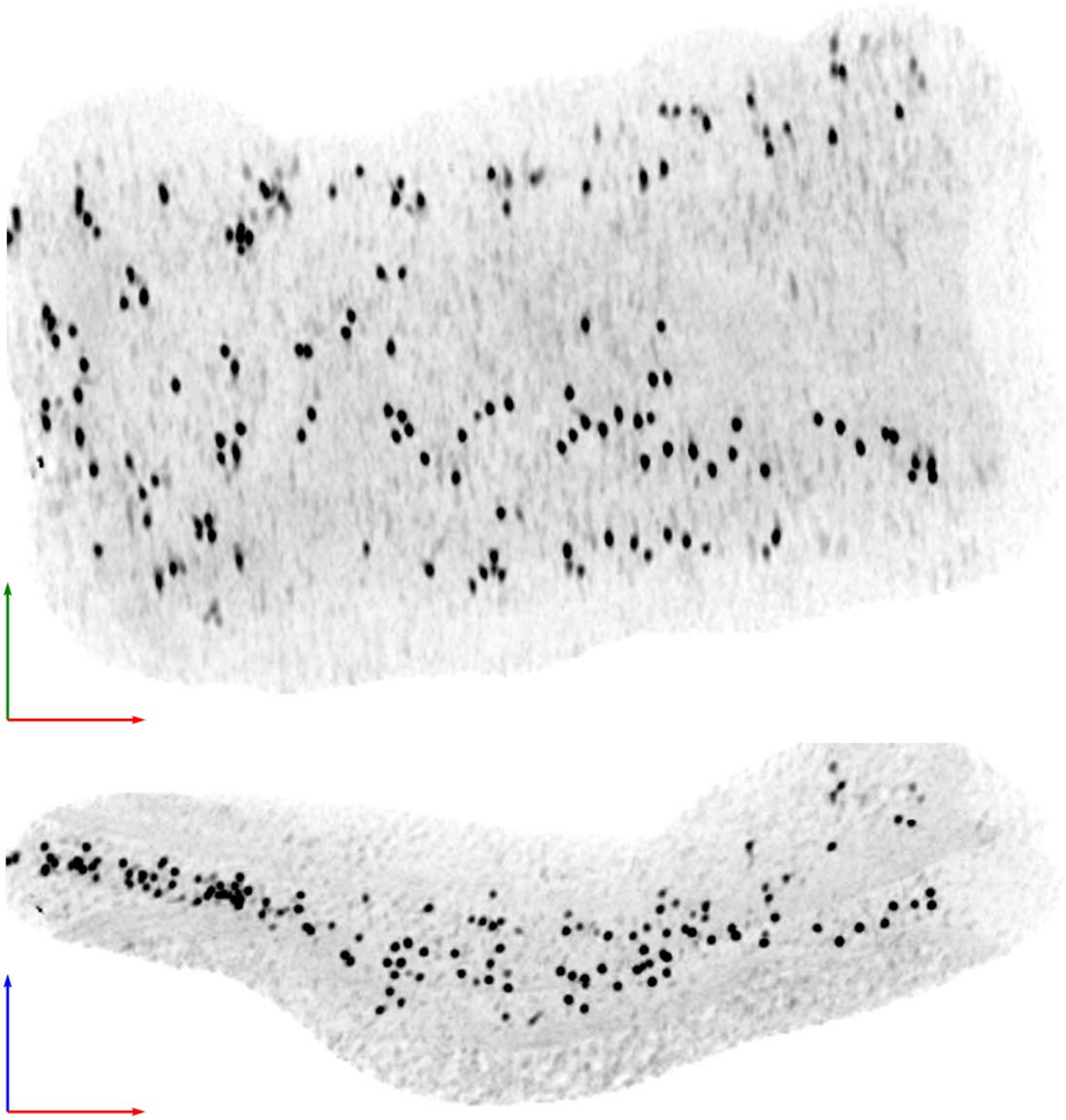

| Tomogram ID: 20250206_AMmilled27-2_289 | SET: anti-GluA1 11B8 "C-term" |
|---|---|
| Cleft ID: 0 | Type: synaptosome |

**Denoised Tomogram Slice**

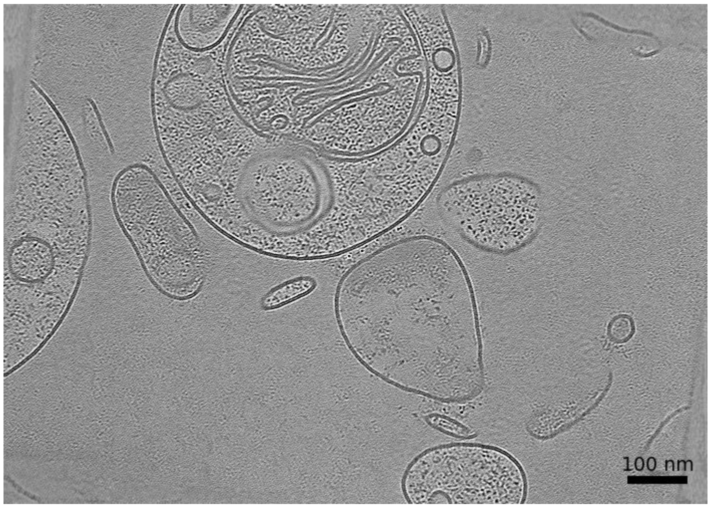


**Membrane Segmentation & Cleft Position**

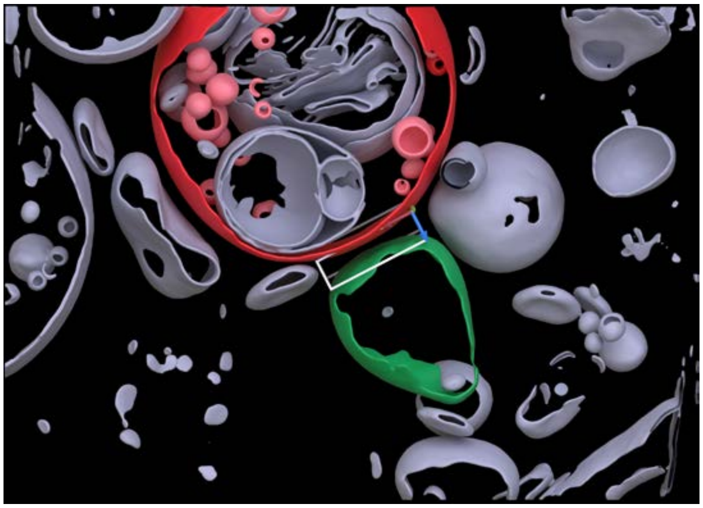

**Cleft MIP**

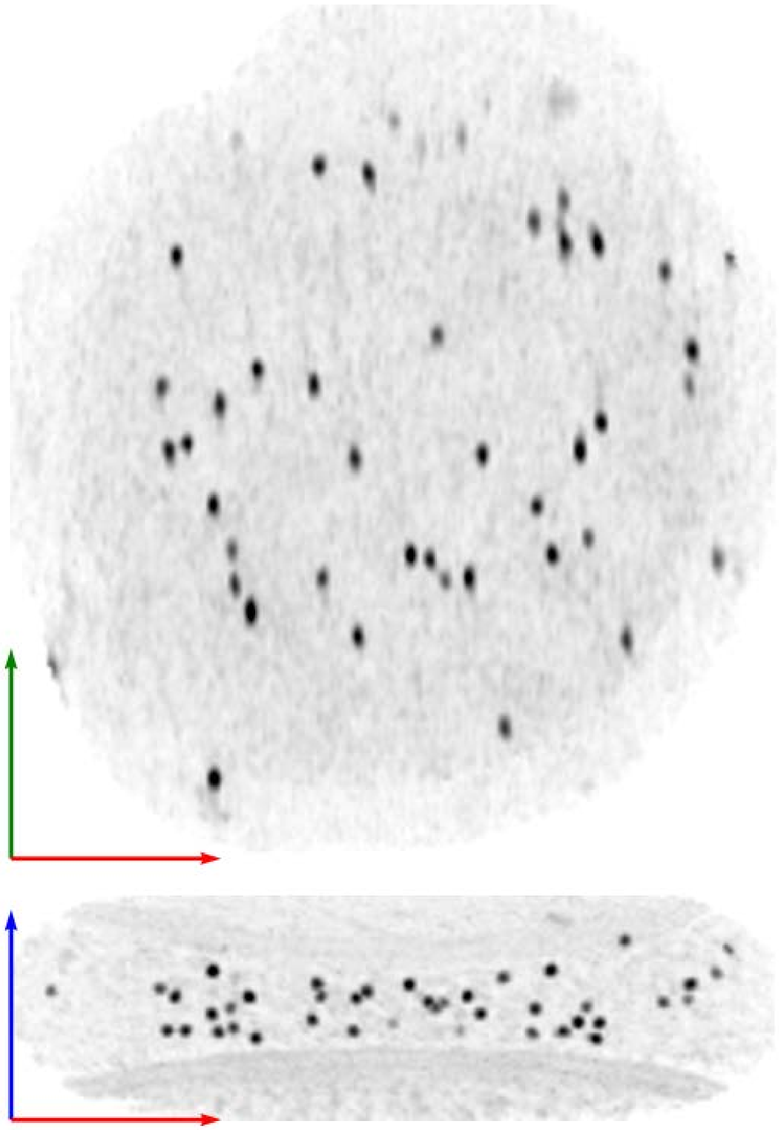

**Tomogram ID: 20240822_HippWaffle_125** **SET: anti-GluN1 5F11 "C-term"**

Cleft ID: 0 Type: synaptosome

**Denoised Tomogram Slice**

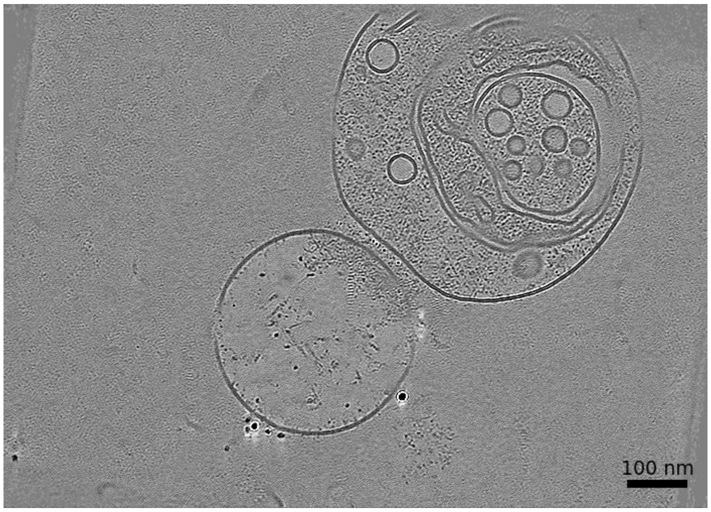


**Membrane Segmentation & Cleft Position**

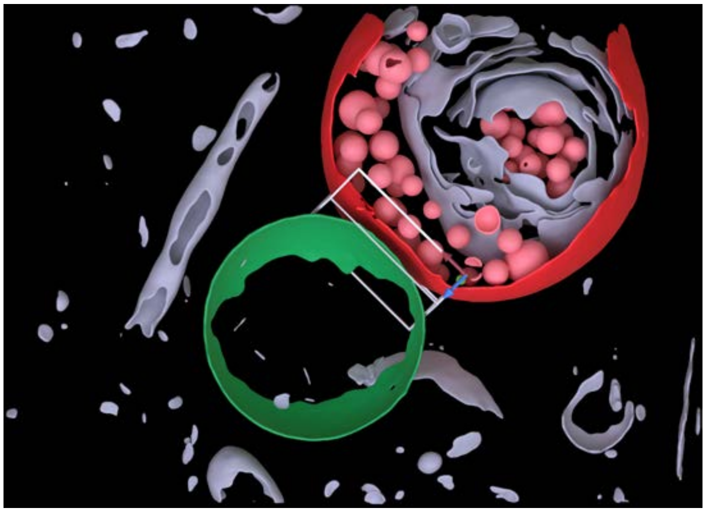

**Cleft MIP**

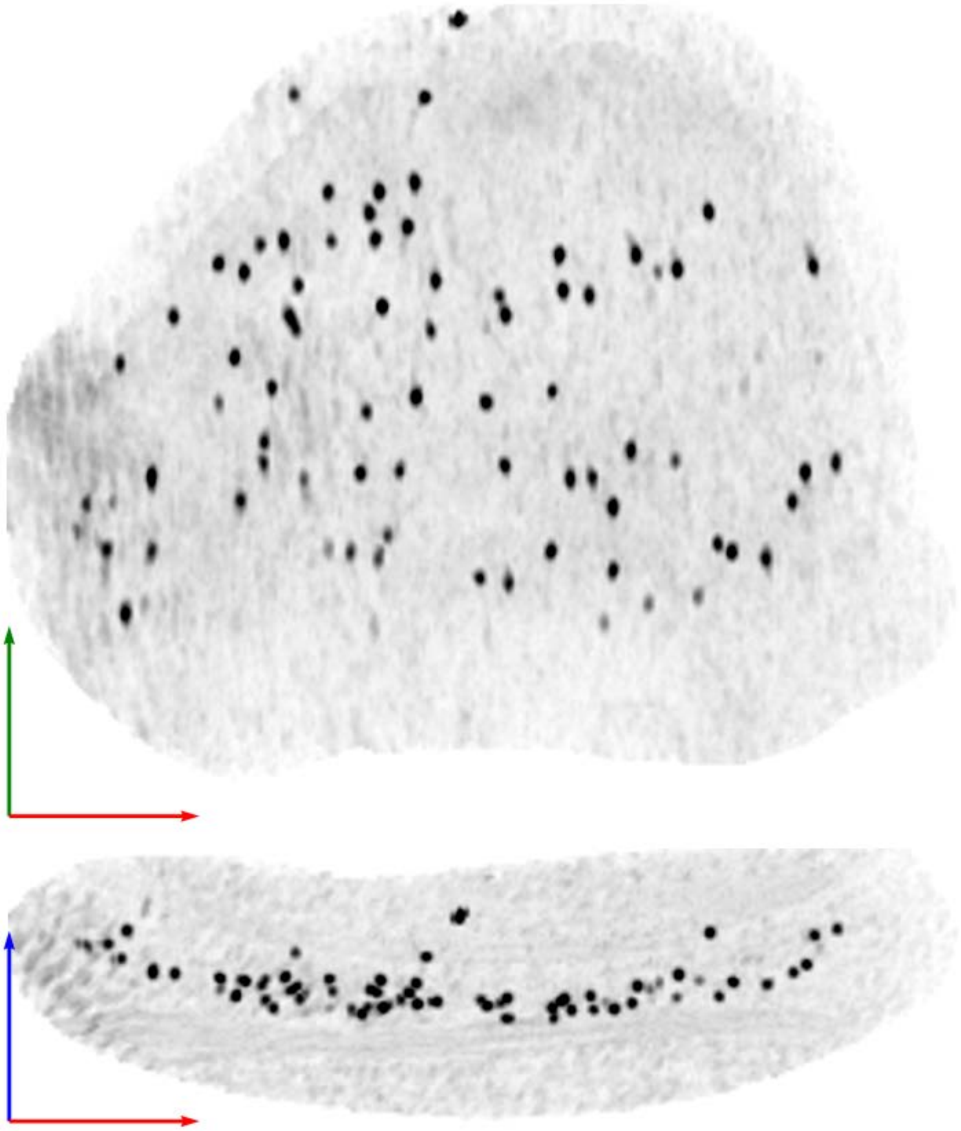

| | |
|---|---|
| **Tomogram ID: 20240822_HippWaffle_129** | **SET: anti-GluN1 5F11 "C-term"** |
| Cleft ID: 0 | Type: semi-tissue |

**Denoised Tomogram Slice**

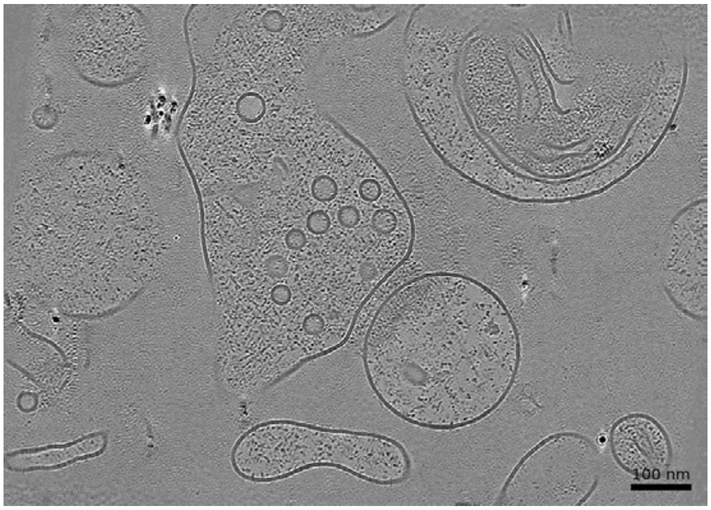


**Membrane Segmentation & Cleft Position**

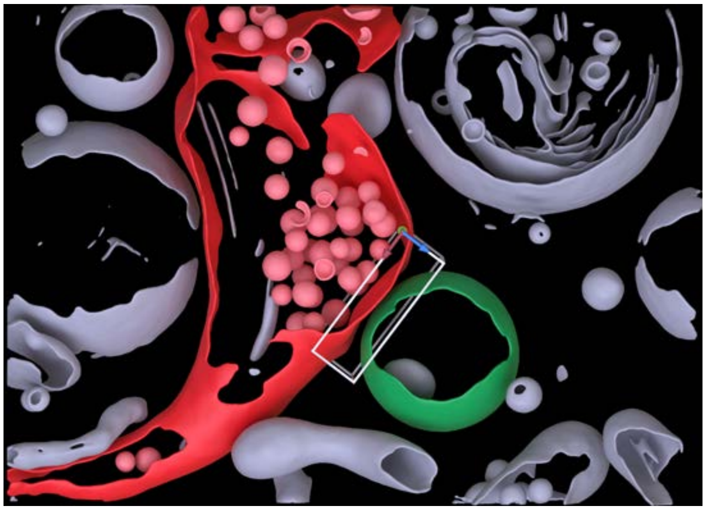

**Cleft MIP**

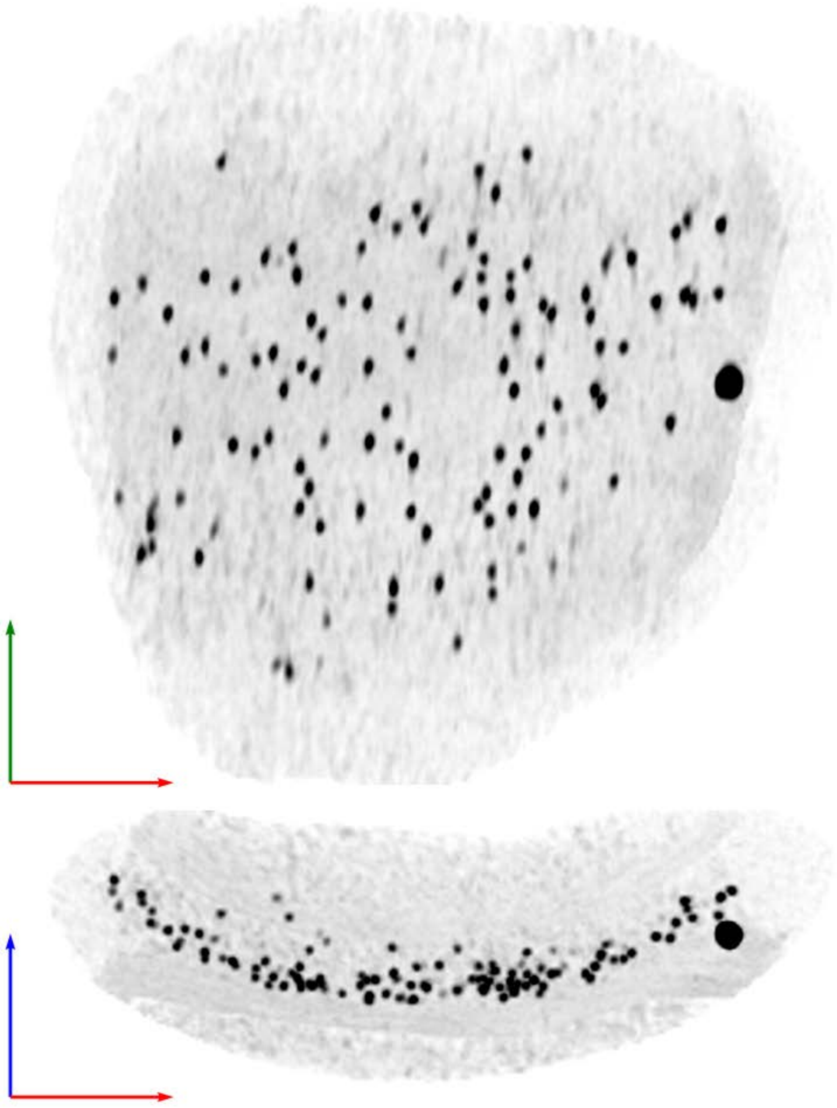

**Tomogram ID: 20240822_HippWaffle_51** **SET: anti-GluN1 5F11 "C-term"**

Cleft ID: 0 Type: synaptosome

**Denoised Tomogram Slice**

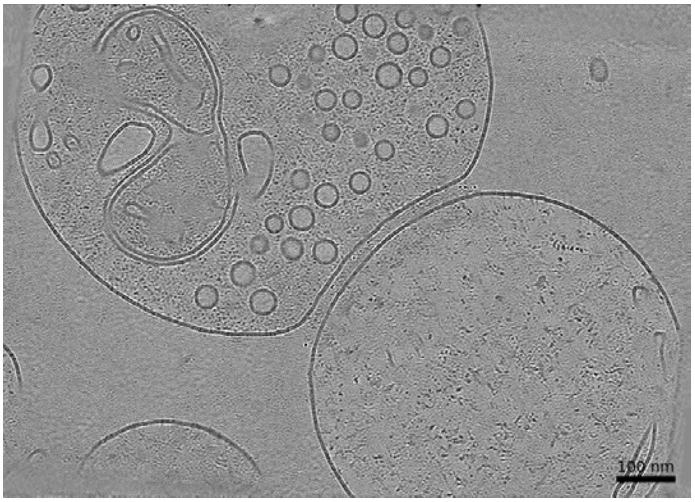


**Membrane Segmentation & Cleft Position**

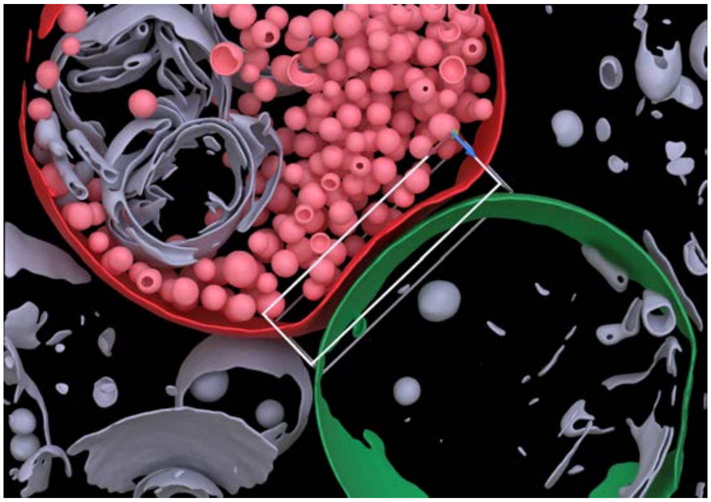

**Cleft MIP**

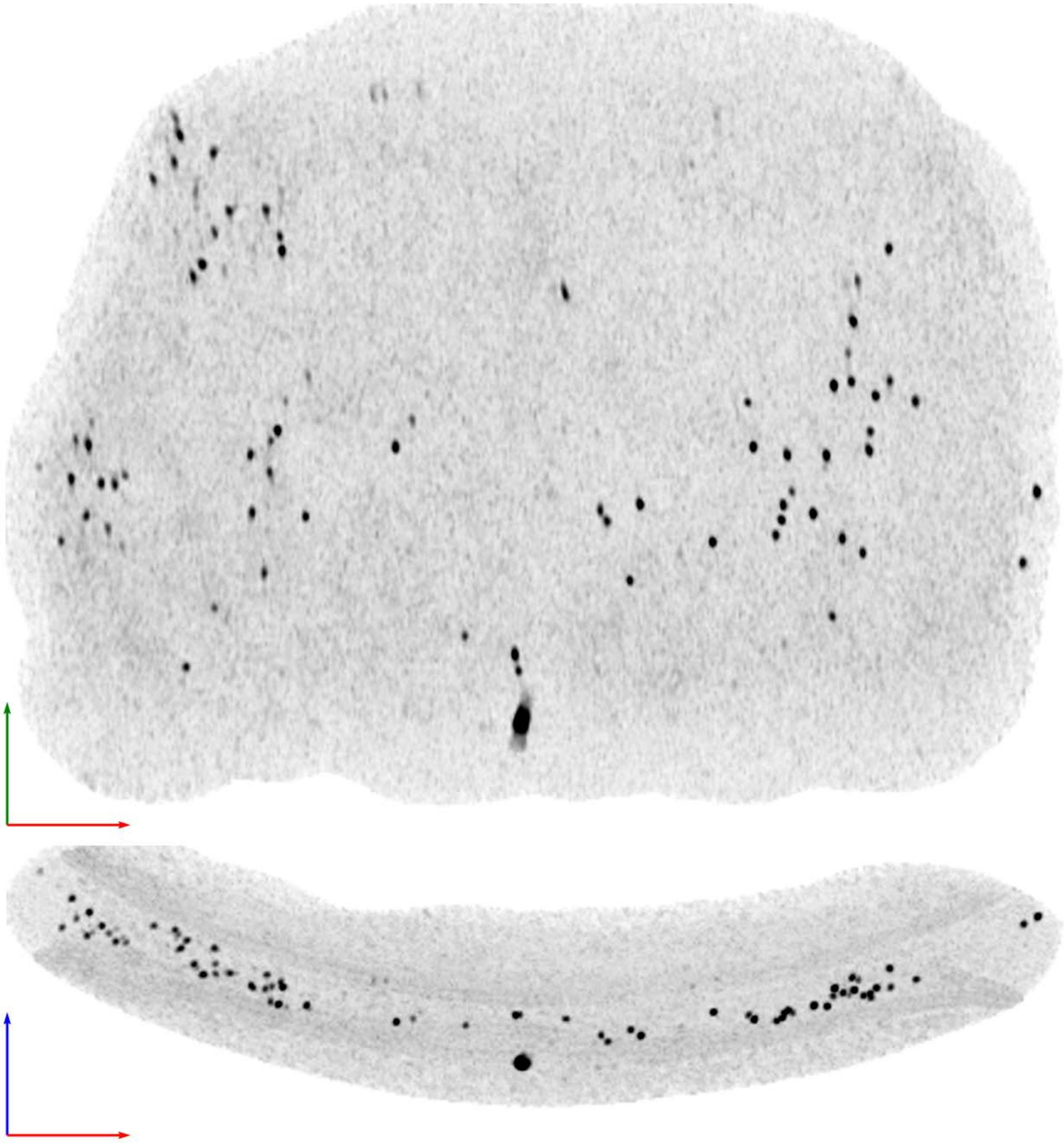

Tomogram ID: 20240920_HippWaffle_113 SET: anti-GluN1 5F11 "C-term"

Cleft ID: 0 Type: synaptosome

**Denoised Tomogram Slice**

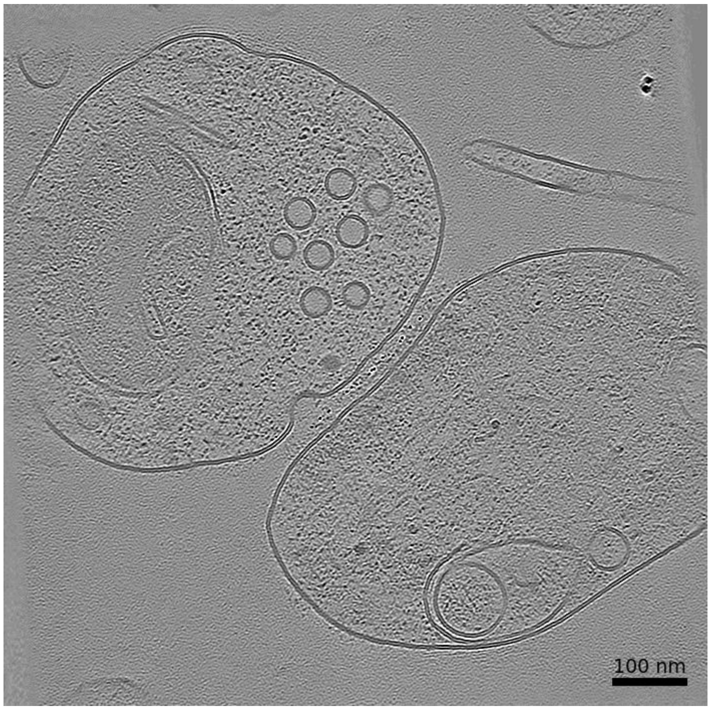


**Membrane Segmentation & Cleft Position**

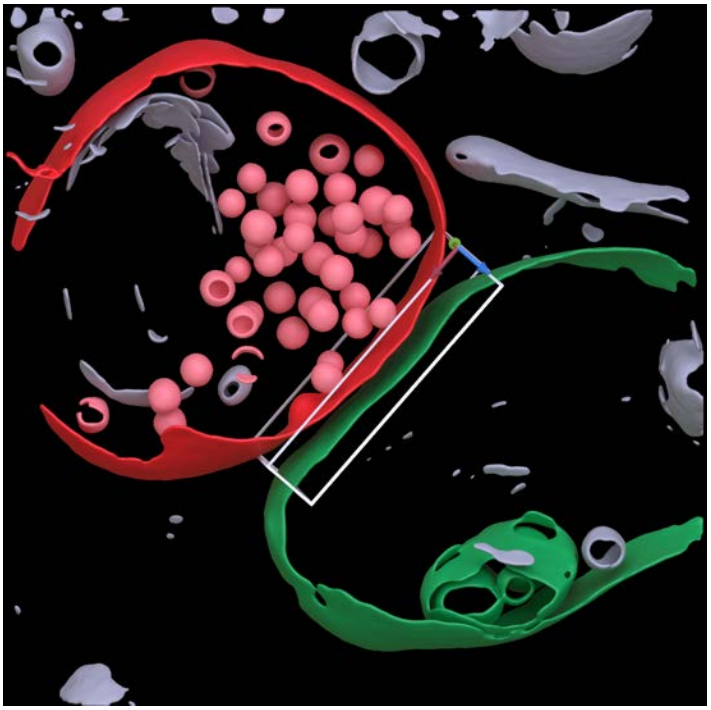

**Cleft MIP**

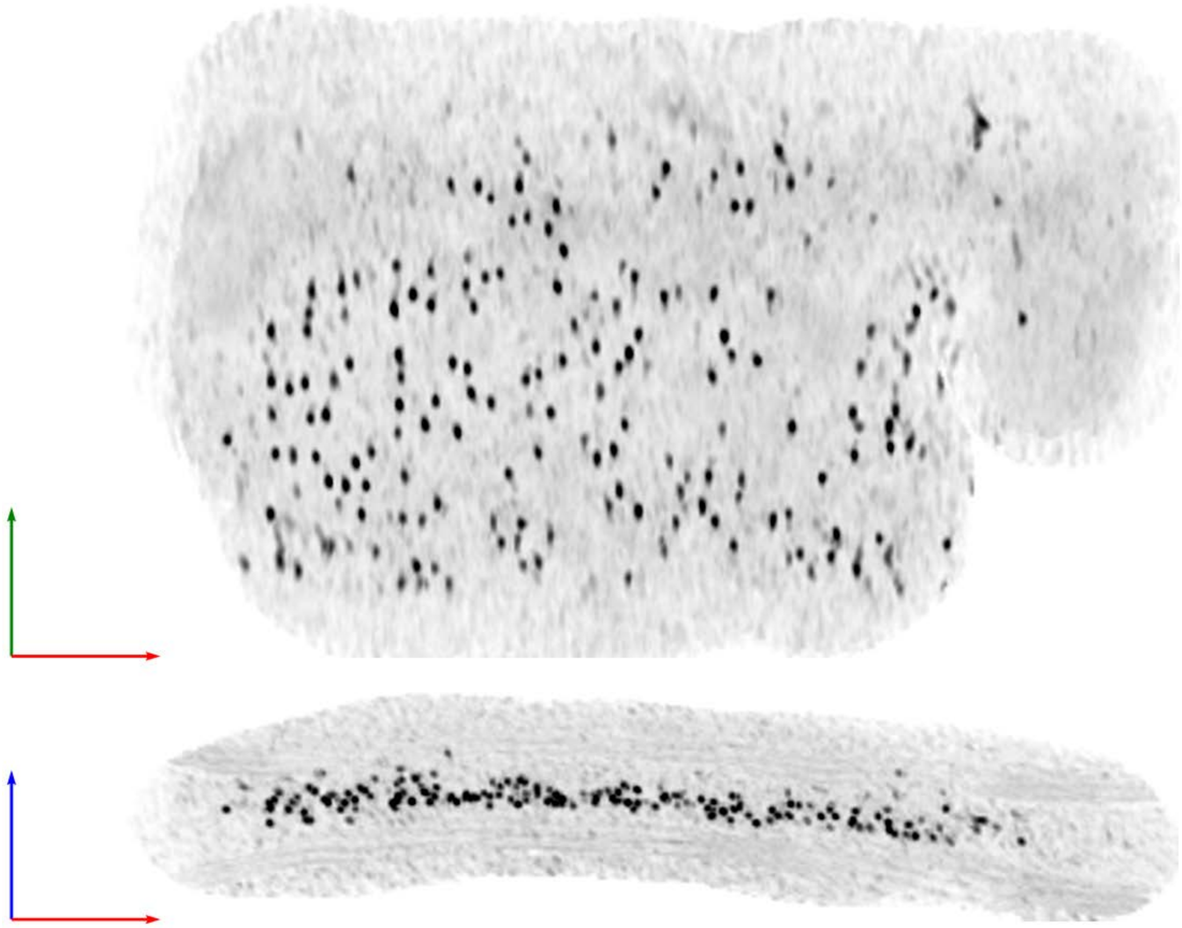

**Tomogram ID: 20240920_HippWaffle_129** **SET: anti-GluN1 5F11 "C-term"**

Cleft ID: 0 Type: semi-tissue

**Denoised Tomogram Slice**

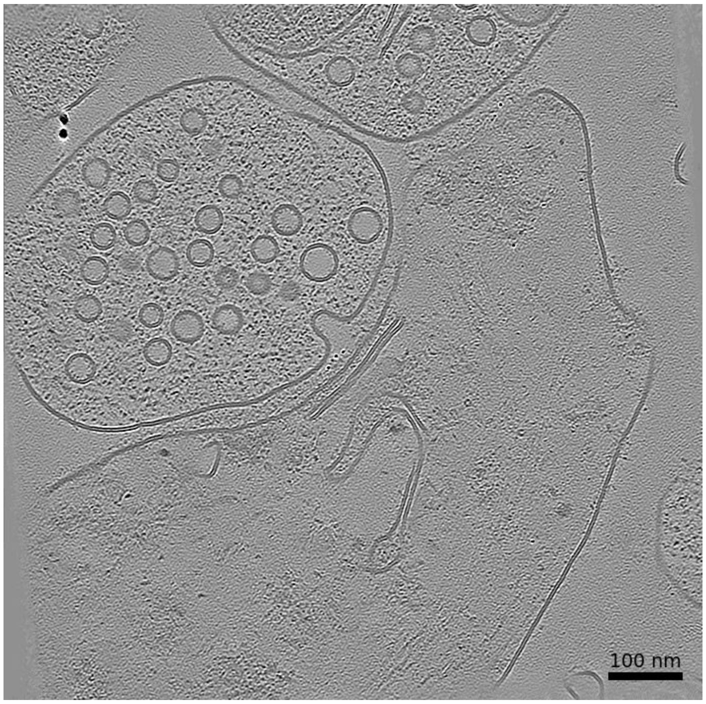


**Membrane Segmentation & Cleft Position**

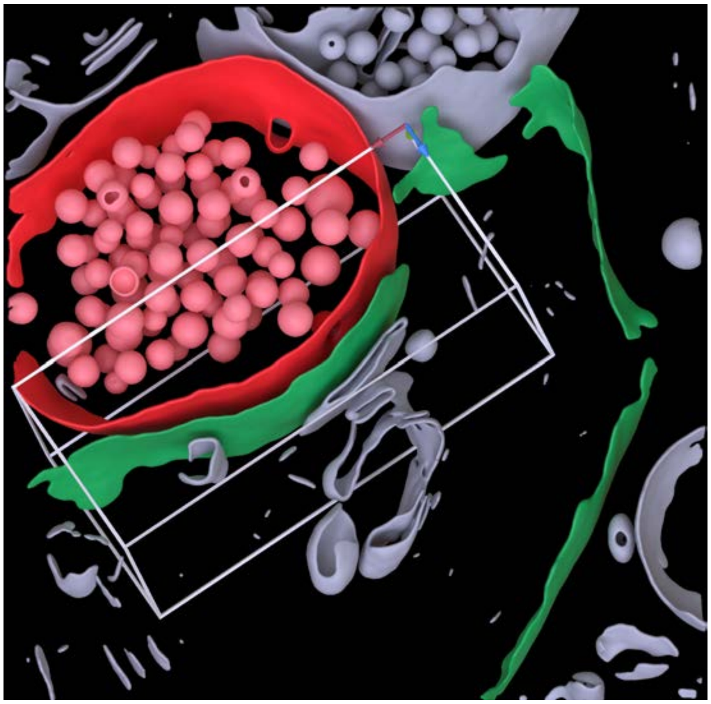

**Cleft MIP**

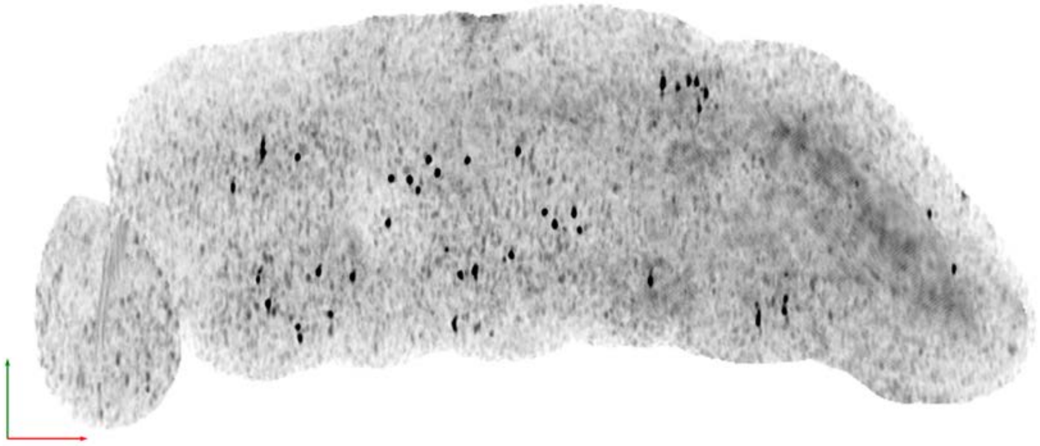

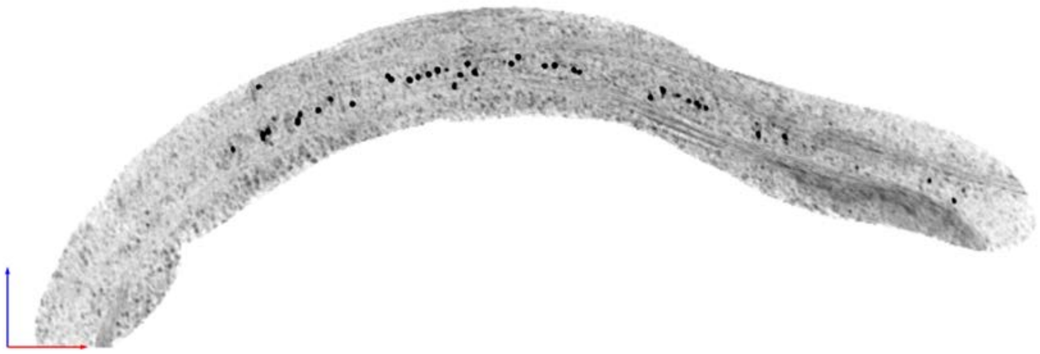

| Tomogram ID: 20241206_AMmilled24-3_58 | SET: Dual 15F1 monomer & 5F11 dimer |
| --- | --- |
| Cleft ID: 0 | Type: semi-tissue |

**Denoised Tomogram Slice**

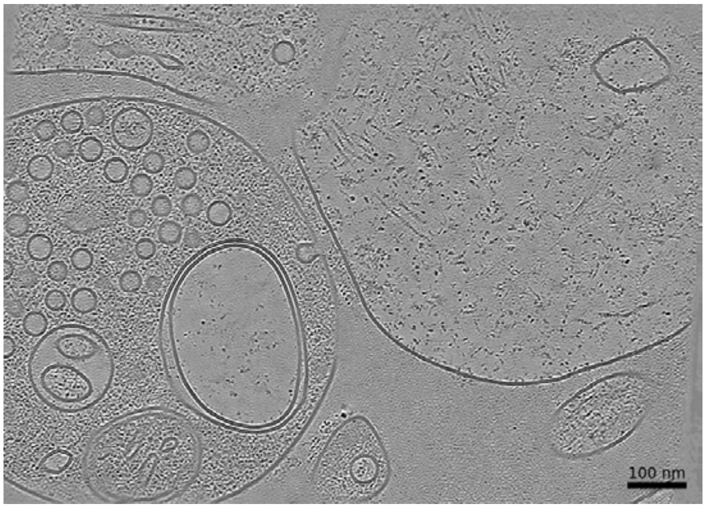


**Membrane Segmentation & Cleft Position**

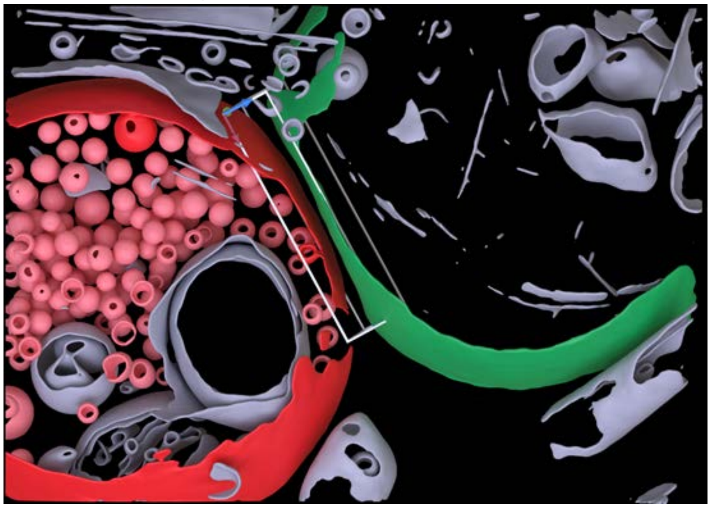

**Cleft MIP**

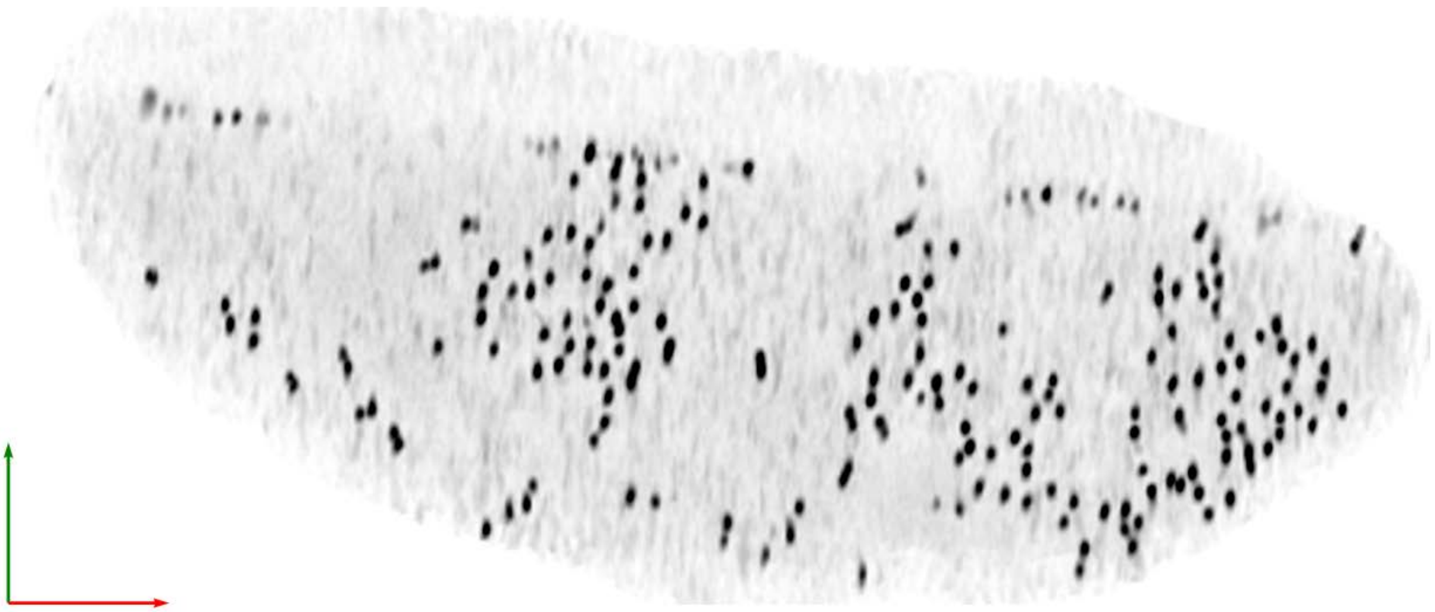

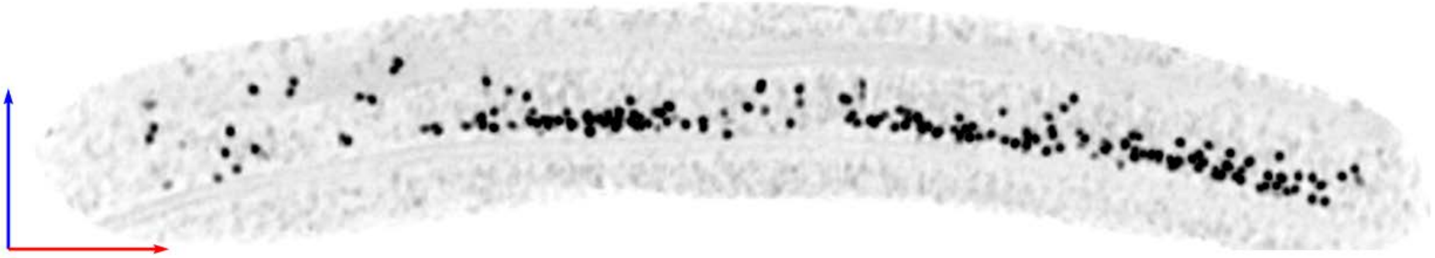

| Tomogram ID: 20241208_AMmilled24-1_64 | SET: Dual 15F1 monomer & 5F11 dimer |
|---|---|
| Cleft ID: 0 | Type: tissue |

**Denoised Tomogram Slice**

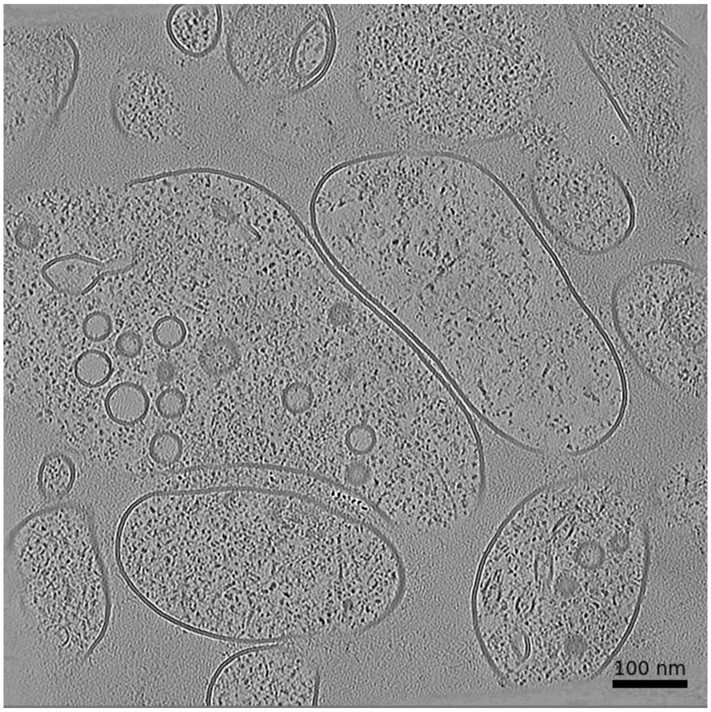


**Membrane Segmentation & Cleft Position**

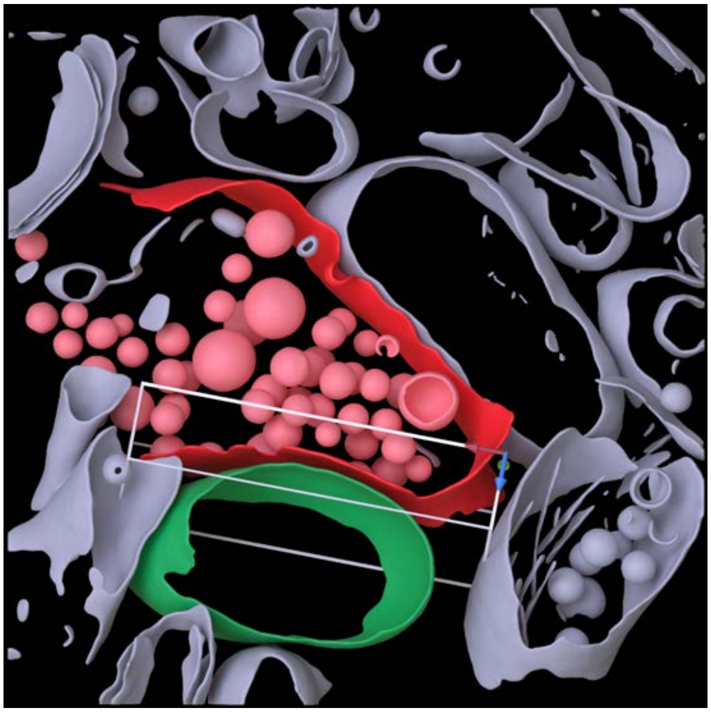

**Cleft MIP**

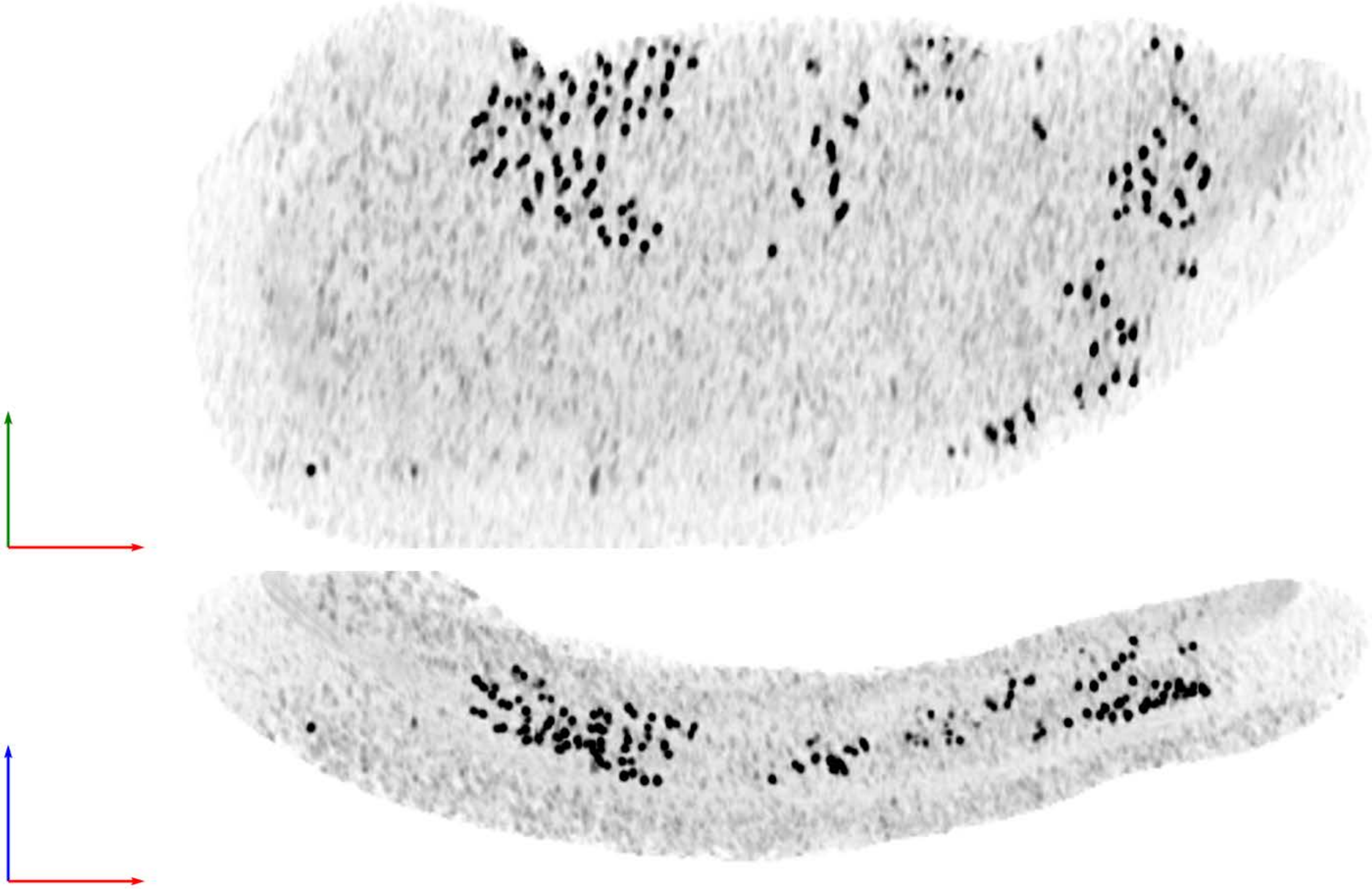

| Tomogram ID: 20241208_AMmilled24-1_65 | SET: Dual 15F1 monomer & 5F11 dimer |
| --- | --- |
| Cleft ID: 0 | Type: semi-tissue |

**Denoised Tomogram Slice**

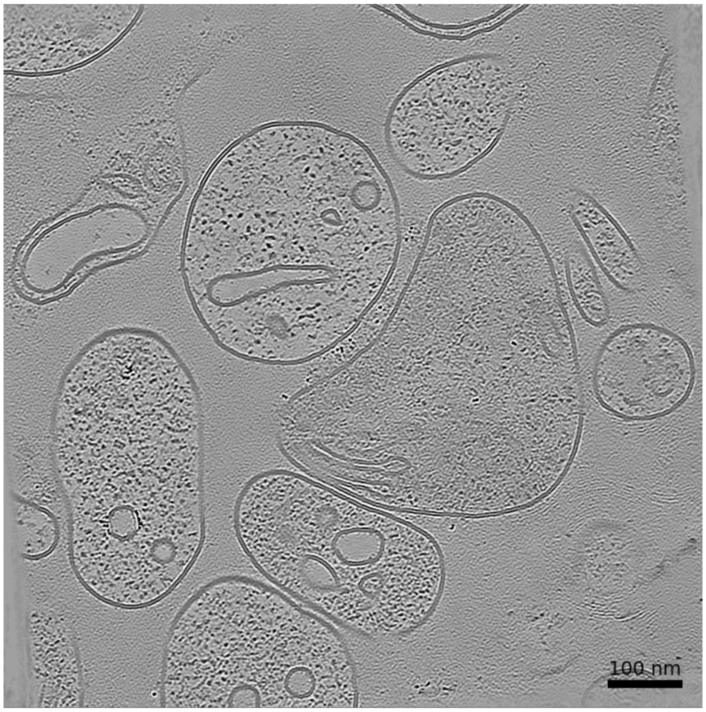


**Membrane Segmentation & Cleft Position**

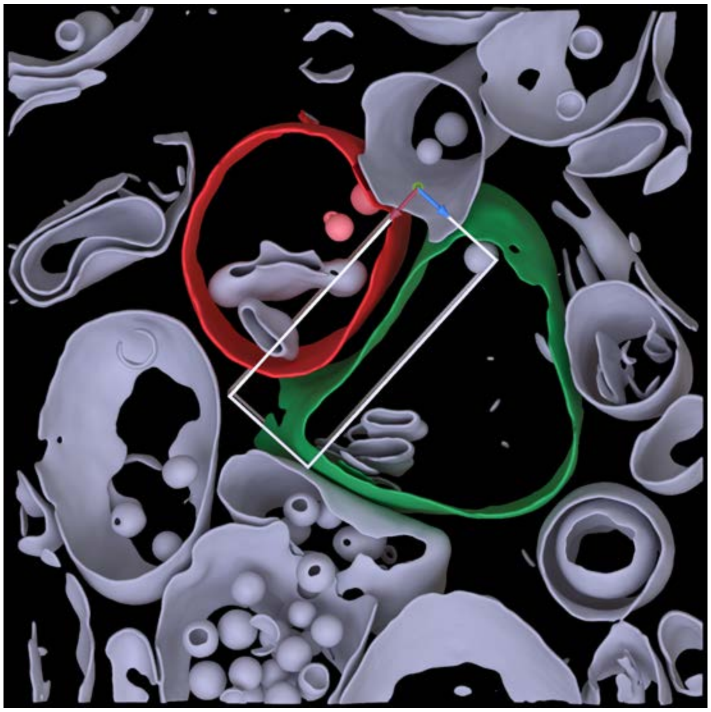

**Cleft MIP**

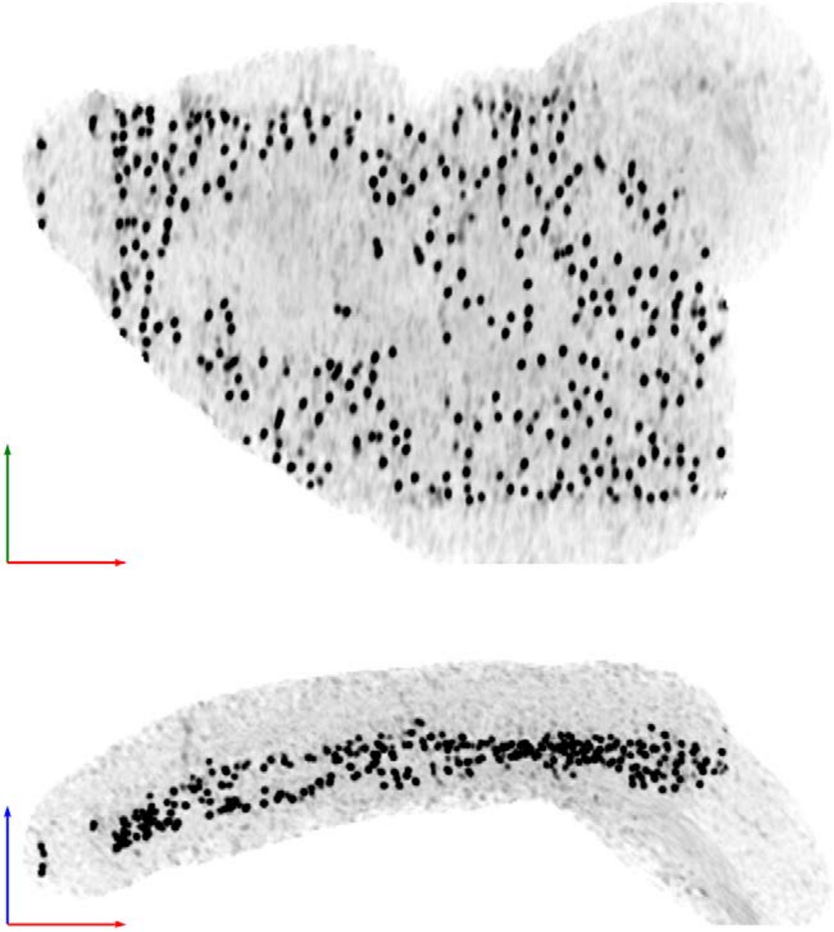

**Tomogram ID: 20241211_HippWaffle_49** | **SET: Dual 15F1 monomer & 5F11 dimer**

Cleft ID: 0 | Type: tissue

### Denoised Tomogram Slice

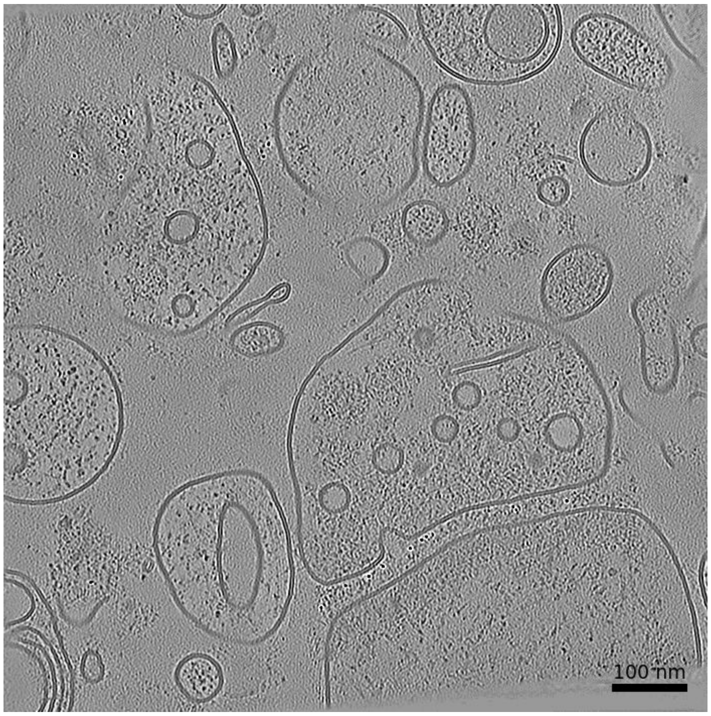


### Membrane Segmentation & Cleft Position

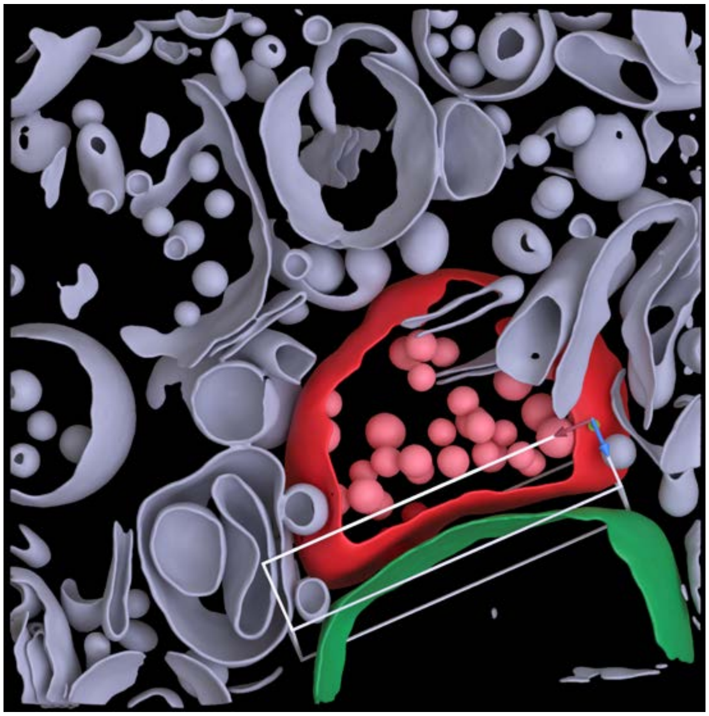

### Cleft MIP

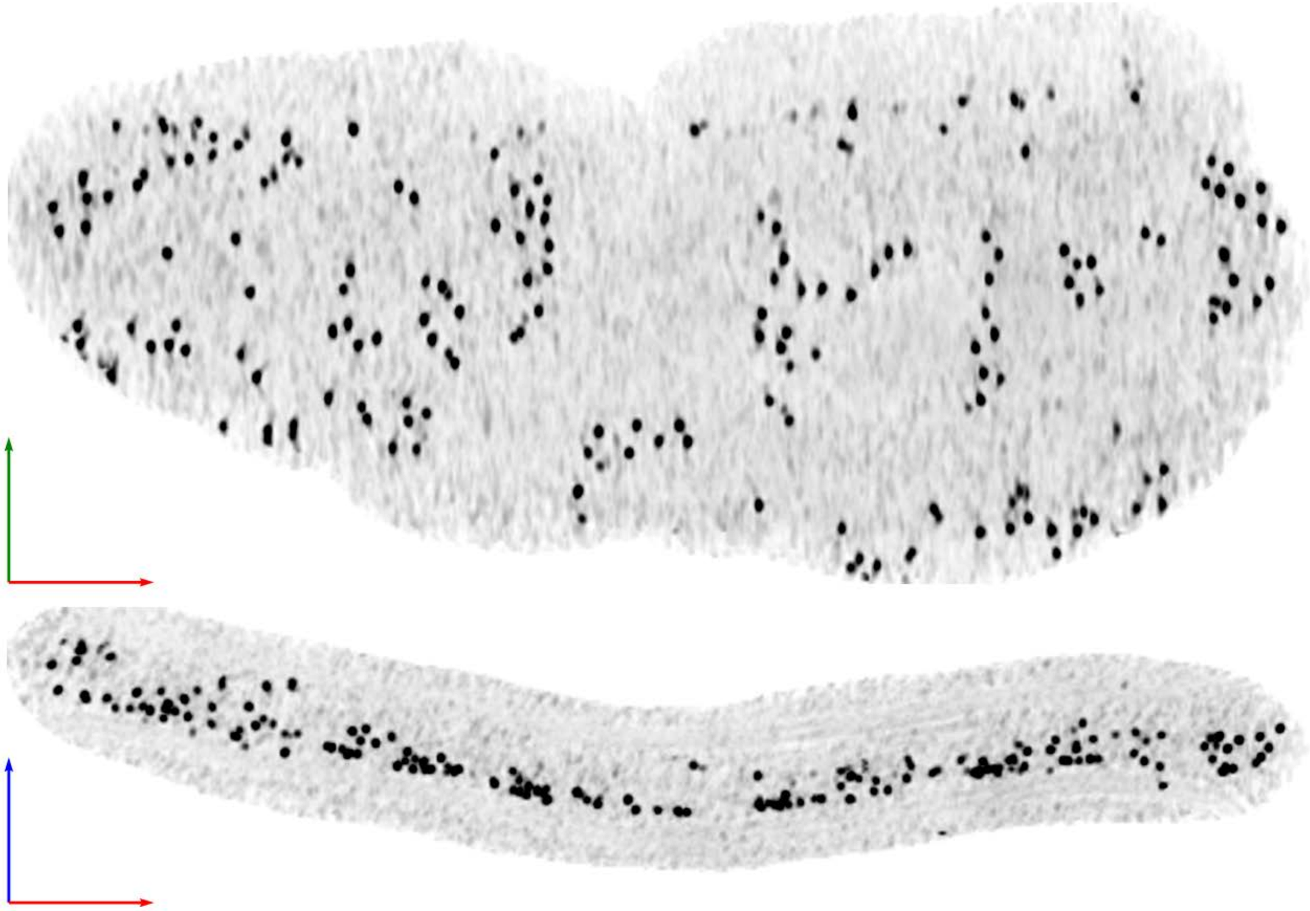

| Tomogram ID: 20241211_HippWaffle_50 | SET: Dual 15F1 monomer & 5F11 dimer |
|---|---|
| Cleft ID: 0 | Type: semi-tissue |

**Denoised Tomogram Slice**

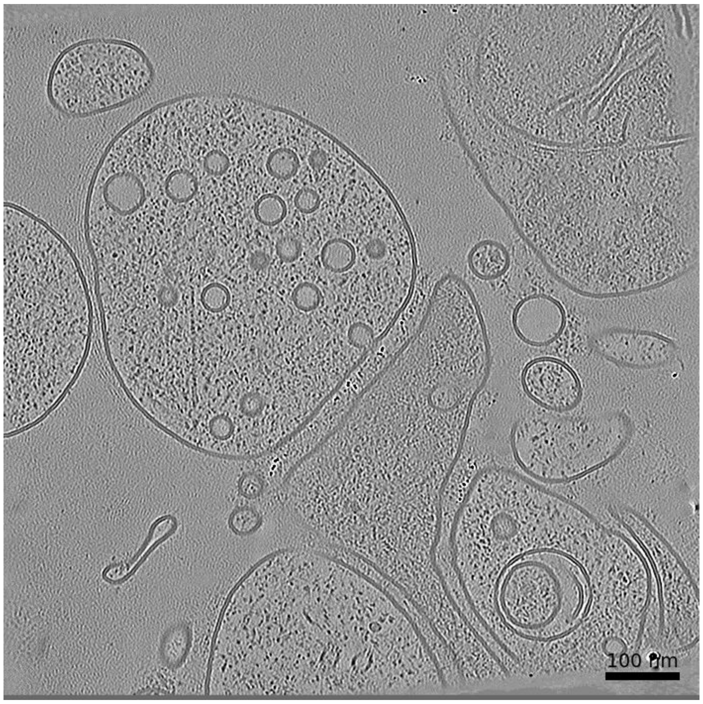


**Membrane Segmentation & Cleft Position**

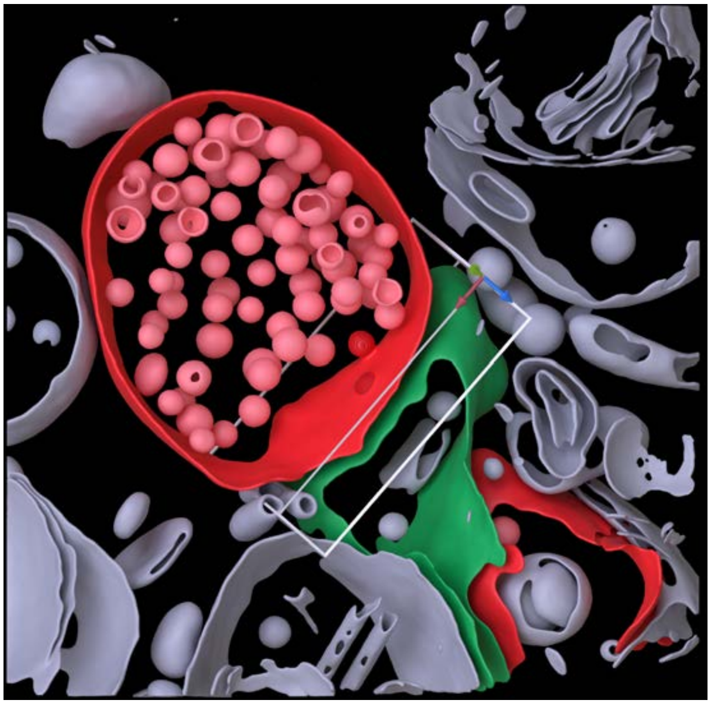

**Cleft MIP**

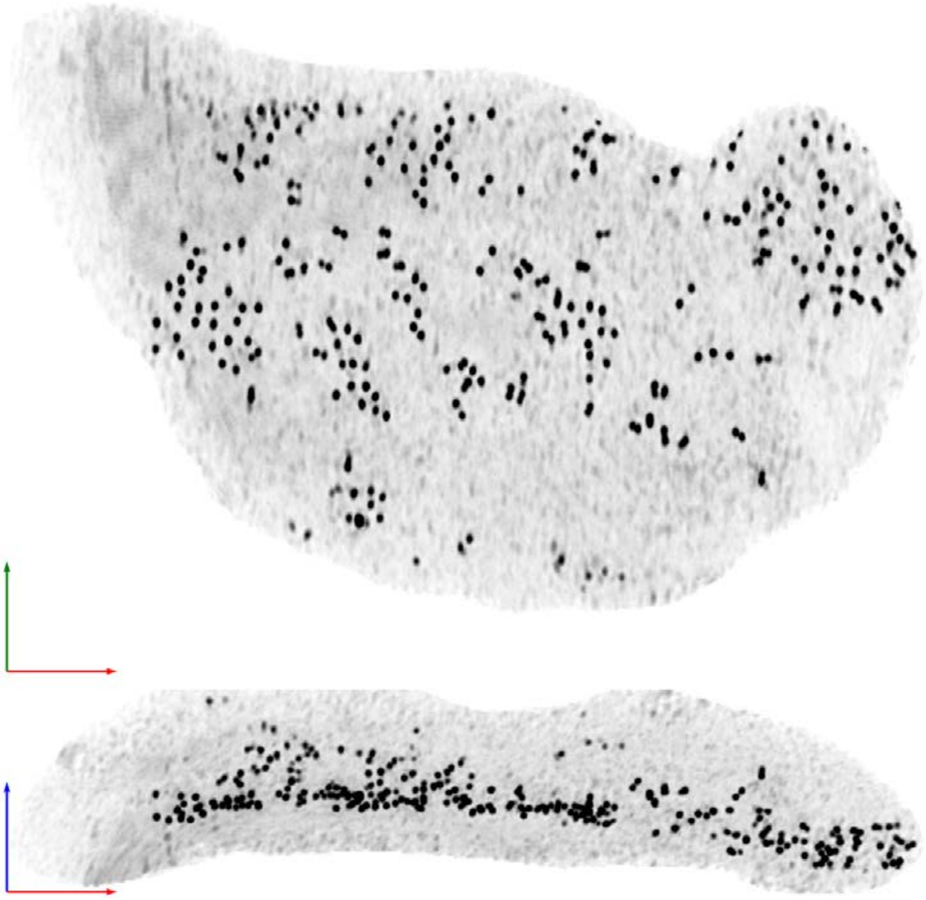

| Tomogram ID: 20241211_HippWaffle_50 | SET: Dual 15F1 monomer & 5F11 dimer |
| --- | --- |
| Cleft ID: 1 | Type: semi-tissue |

**Denoised Tomogram Slice**

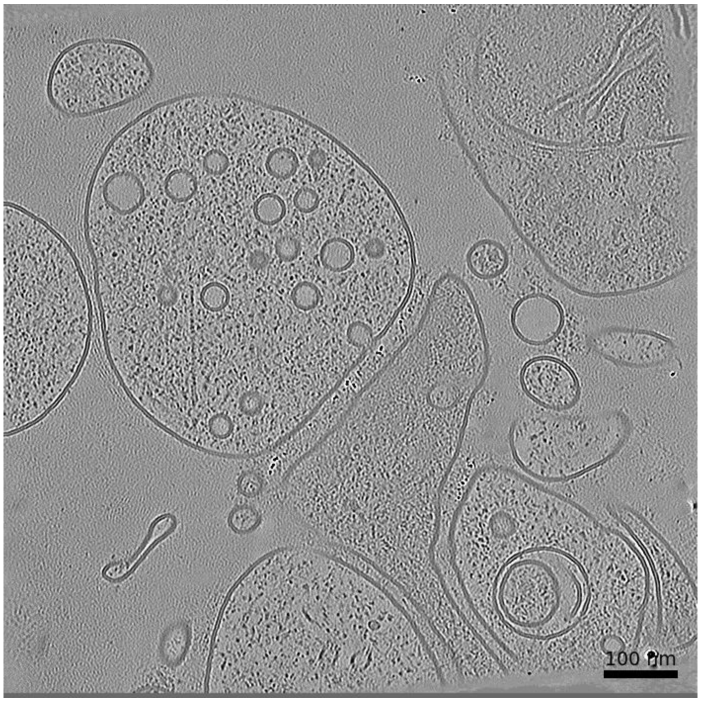


**Membrane Segmentation & Cleft Position**

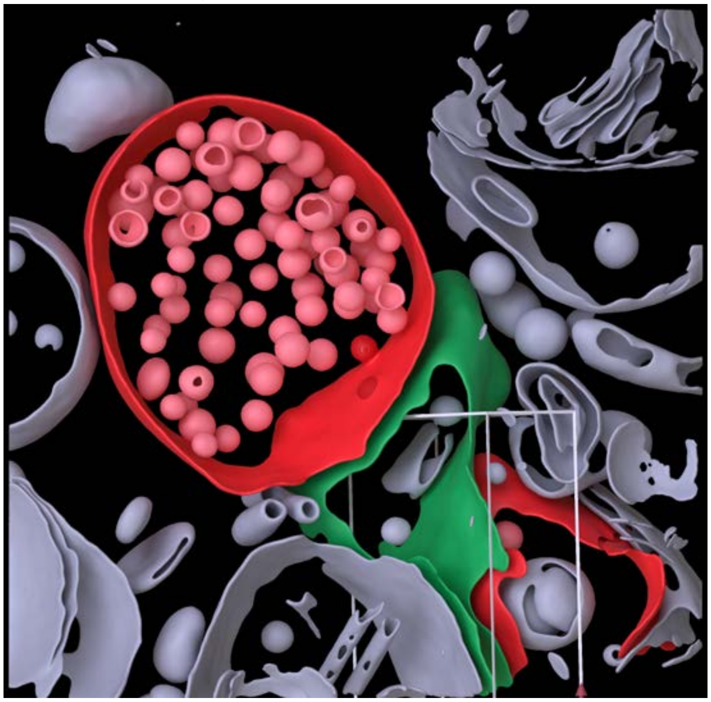

**Cleft MIP**

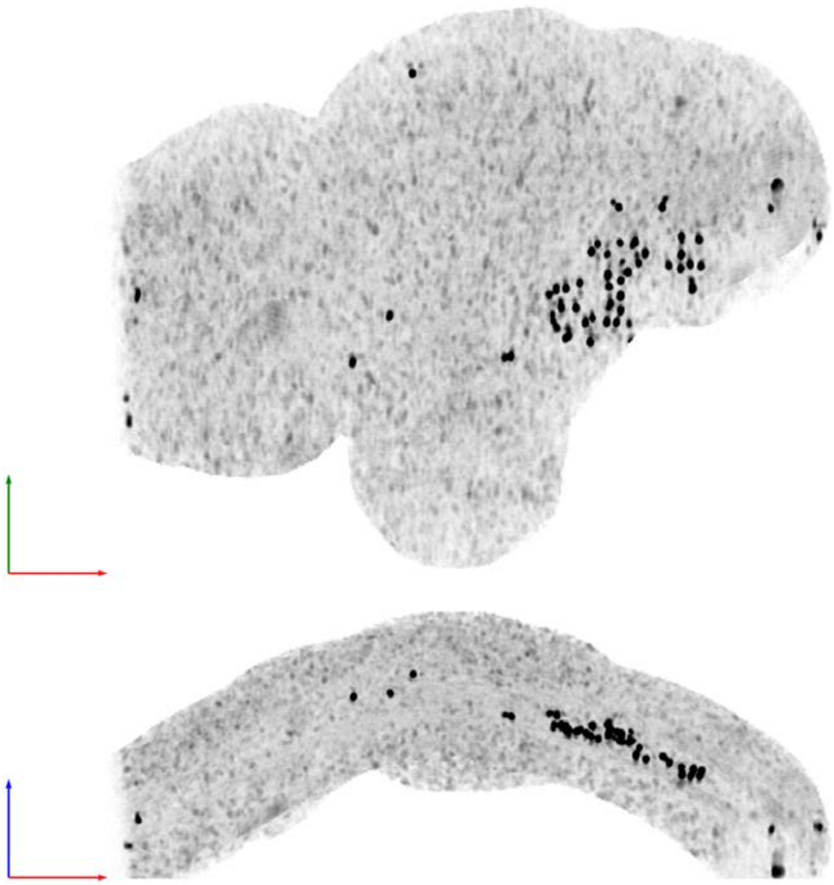

| Tomogram ID: 20250418_AMmilled29-2_Position_87 | SET: Dual 15F1 monomer & 5F11 dimer |
|---|---|
| Cleft ID: 0 | Type: semi-tissue |

**Denoised Tomogram Slice**

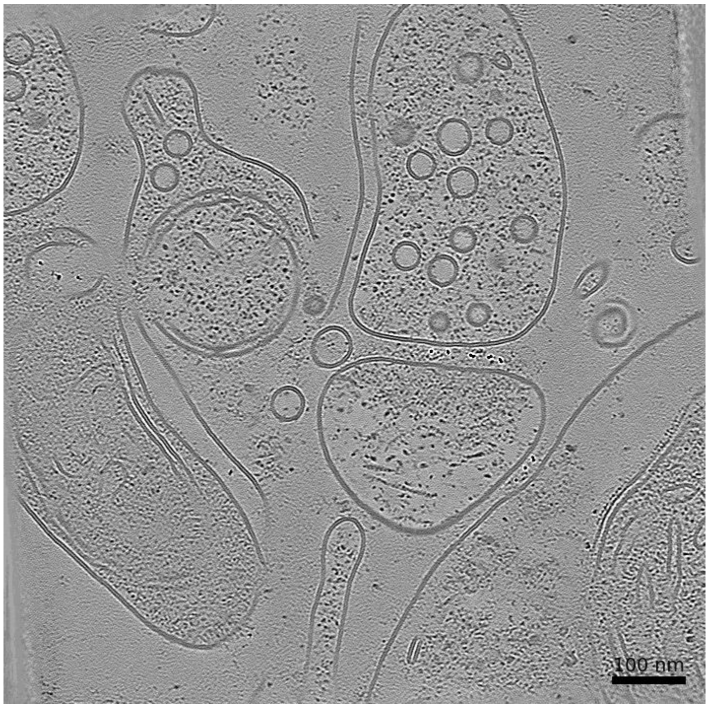


**Membrane Segmentation & Cleft Position**

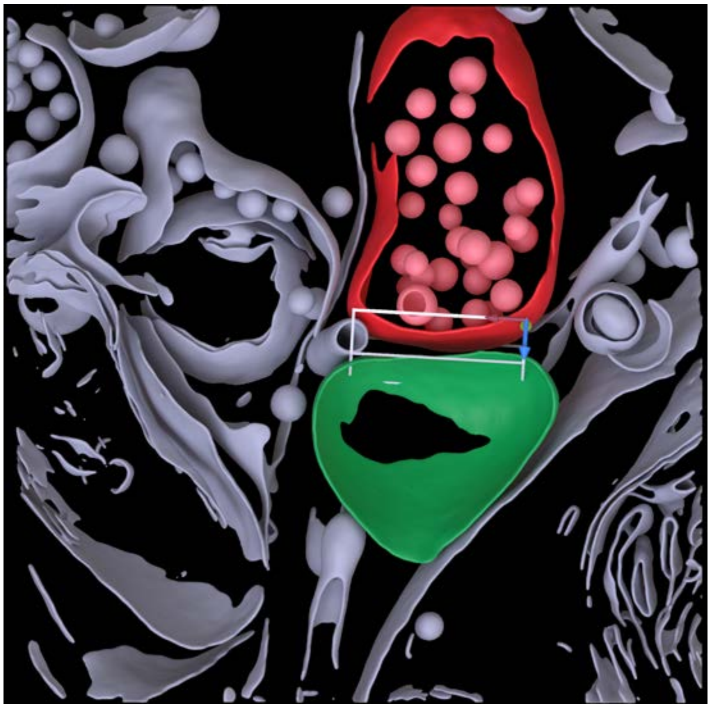

**Cleft MIP**

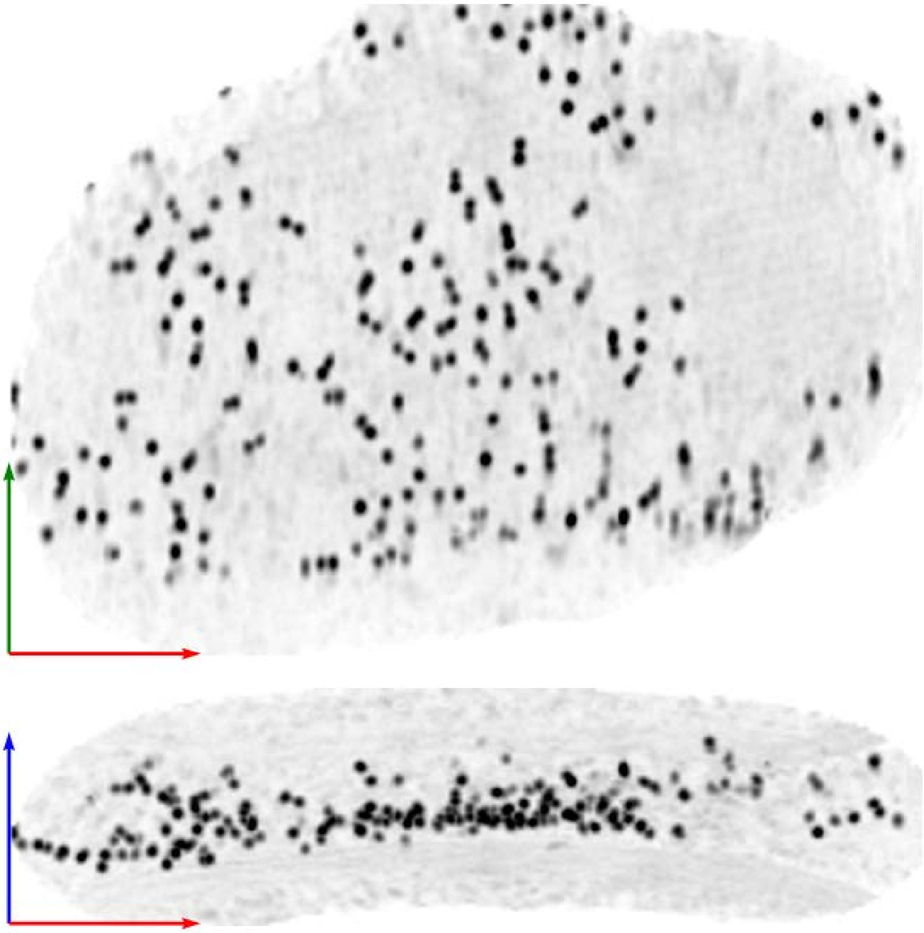

| Tomogram ID: 20251117_AMmilled40-2_Position_148 | SET: Dual 15F1 monomer & 5F11 dimer |
|---|---|
| Cleft ID: 0 | Type: tissue |

**Denoised Tomogram Slice**

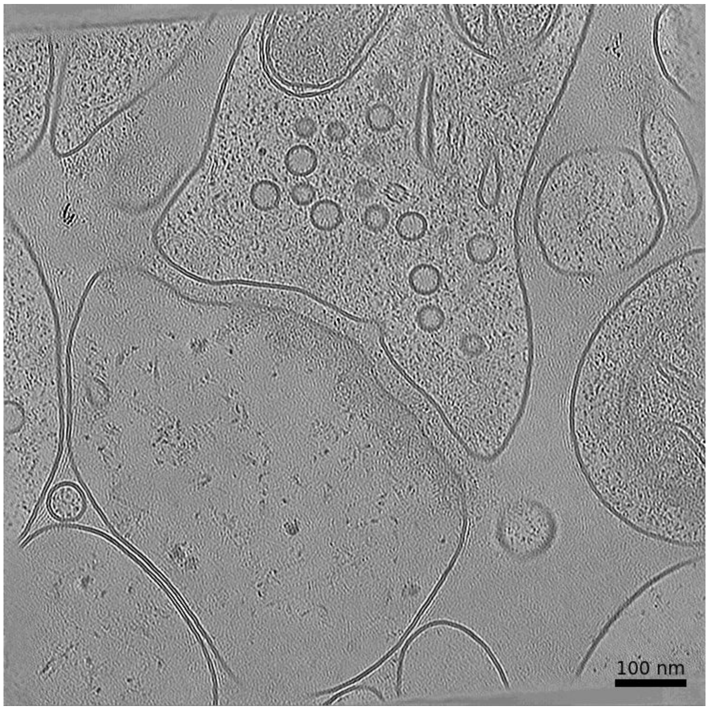


**Membrane Segmentation & Cleft Position**

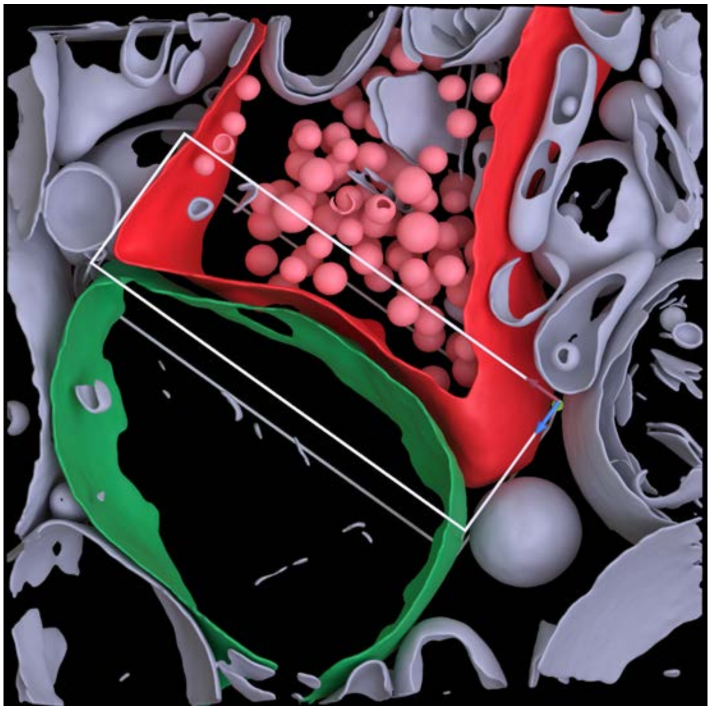

**Cleft MIP**

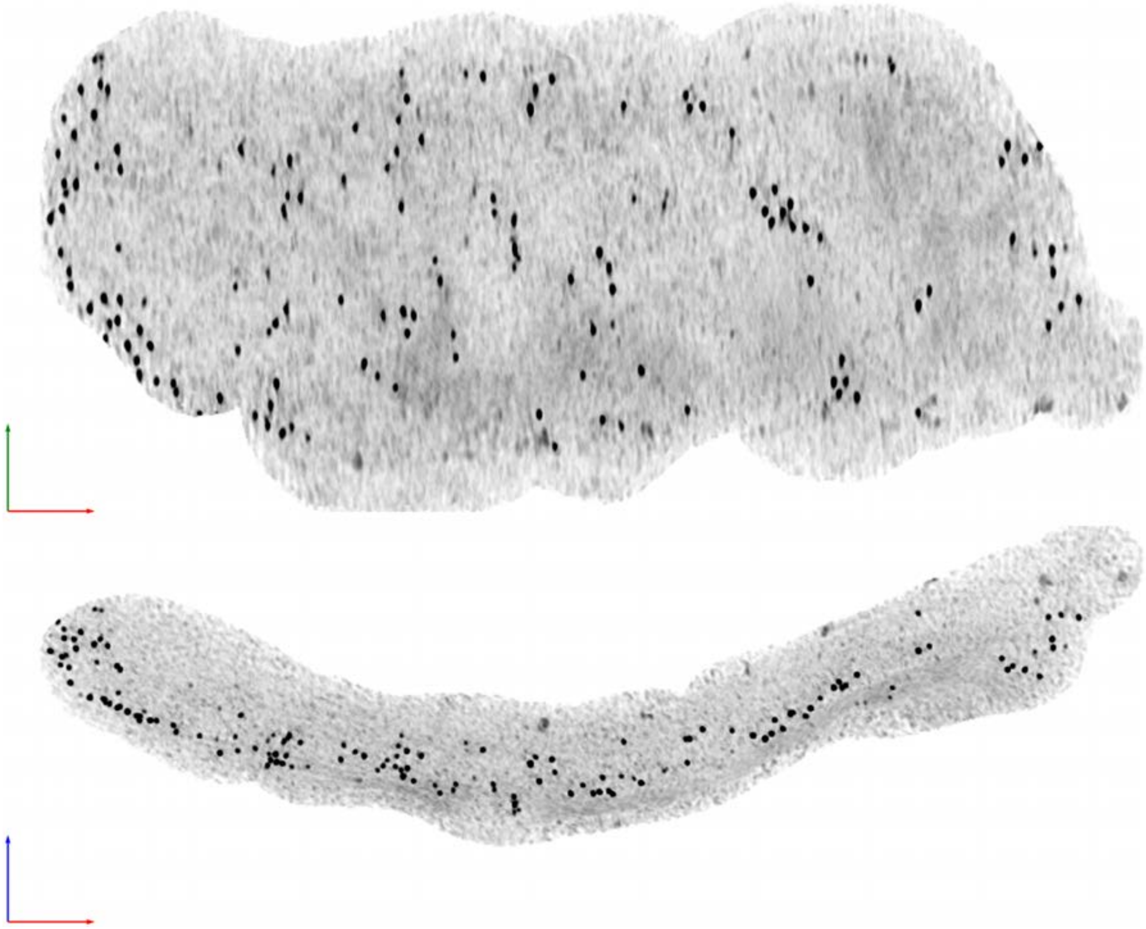

**Tomogram ID: 20251117_AMmilled40-2_Position_148** | **SET: Dual 15F1 monomer & 5F11 dimer**

Cleft ID: 0 | Type: tissue

### Denoised Tomogram Slice

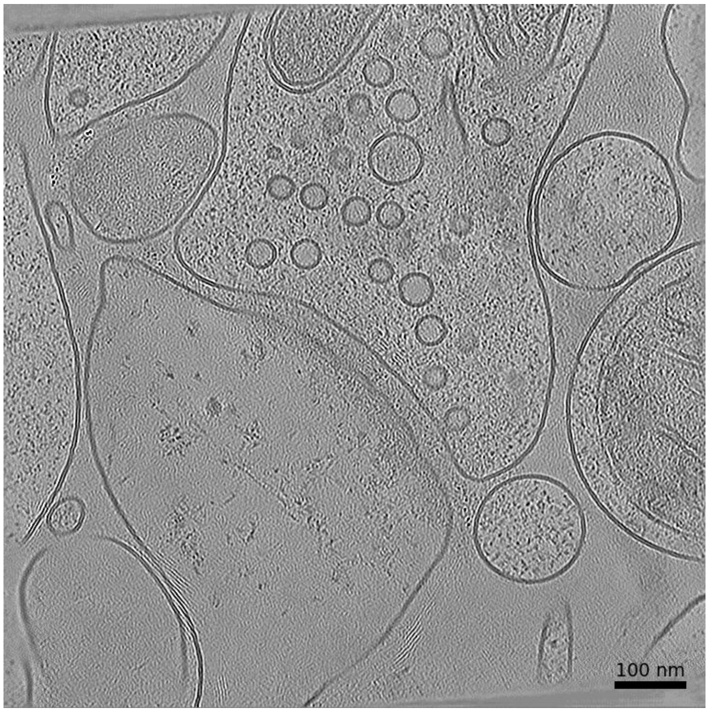


### Membrane Segmentation & Cleft Position

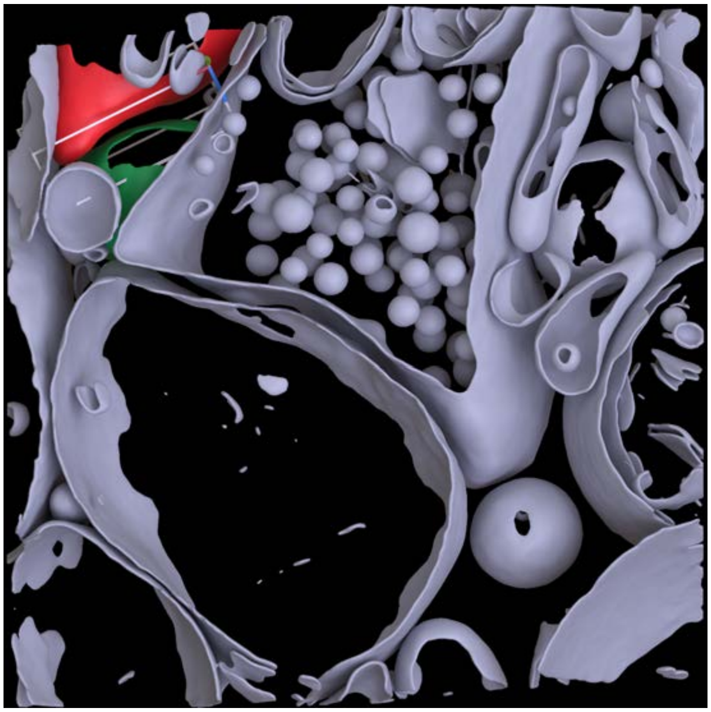

### Cleft MIP

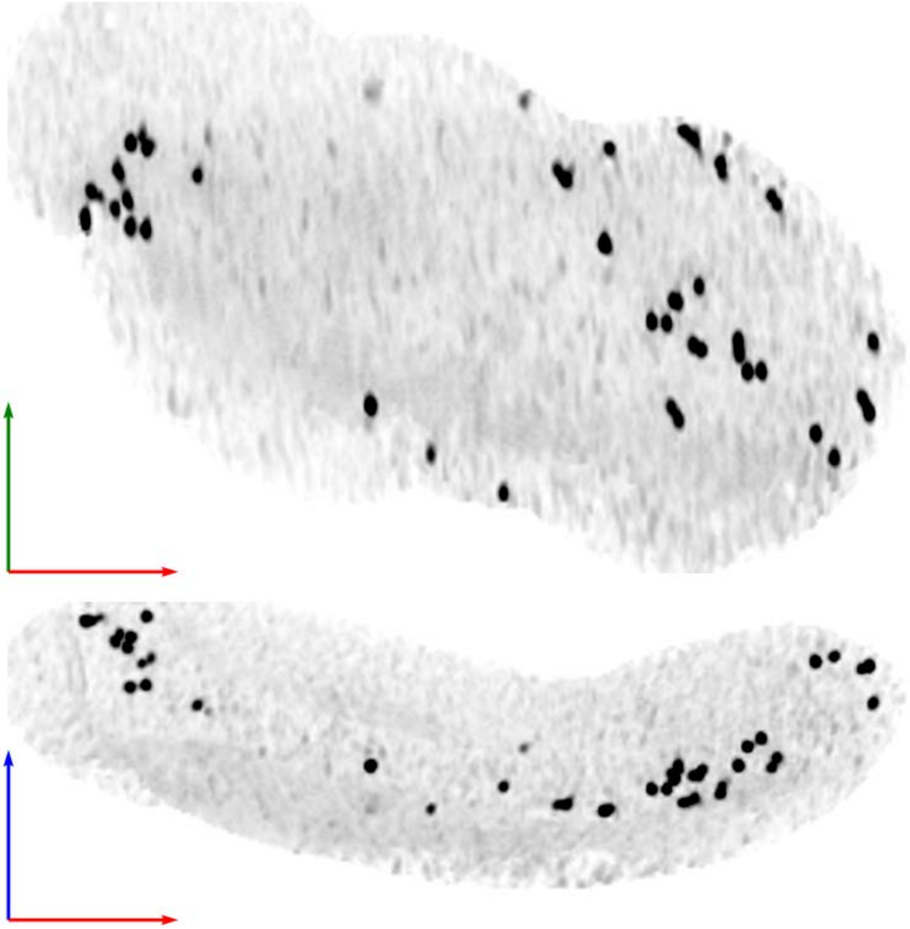

| Tomogram ID: 20260127_AMmilled50-3_Position_13_3 | SET: Dual 15F1 monomer & 5F11 dimer |
|---|---|
| Cleft ID: 1 | Type: tissue |

**Denoised Tomogram Slice**

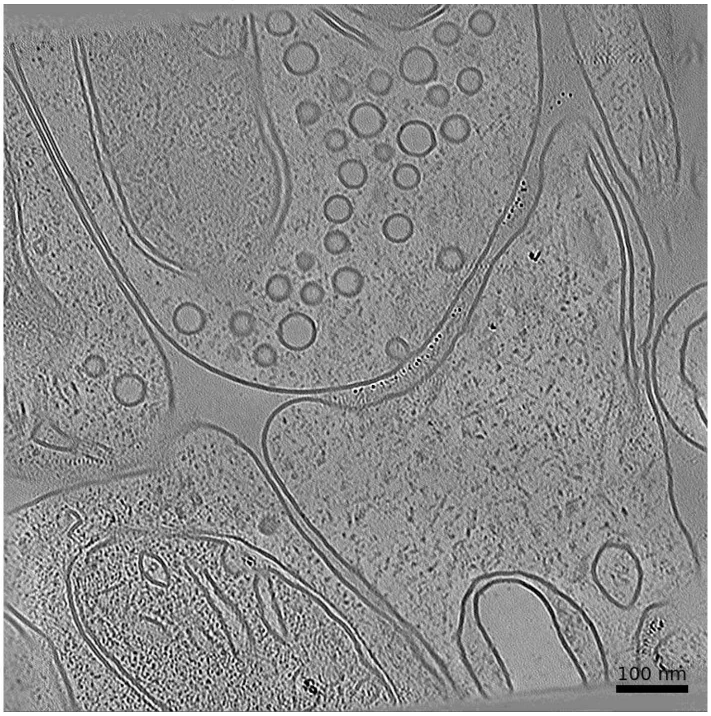


**Membrane Segmentation & Cleft Position**

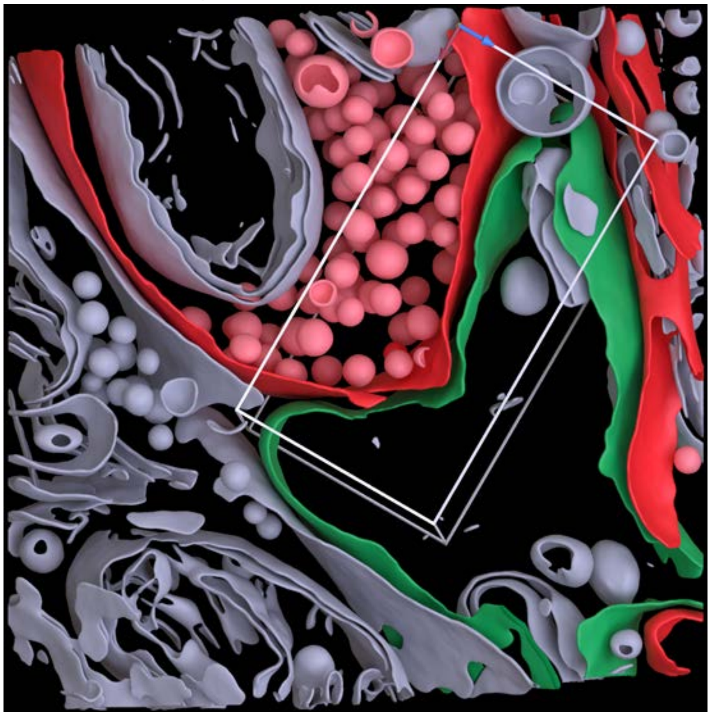

**Cleft MIP**

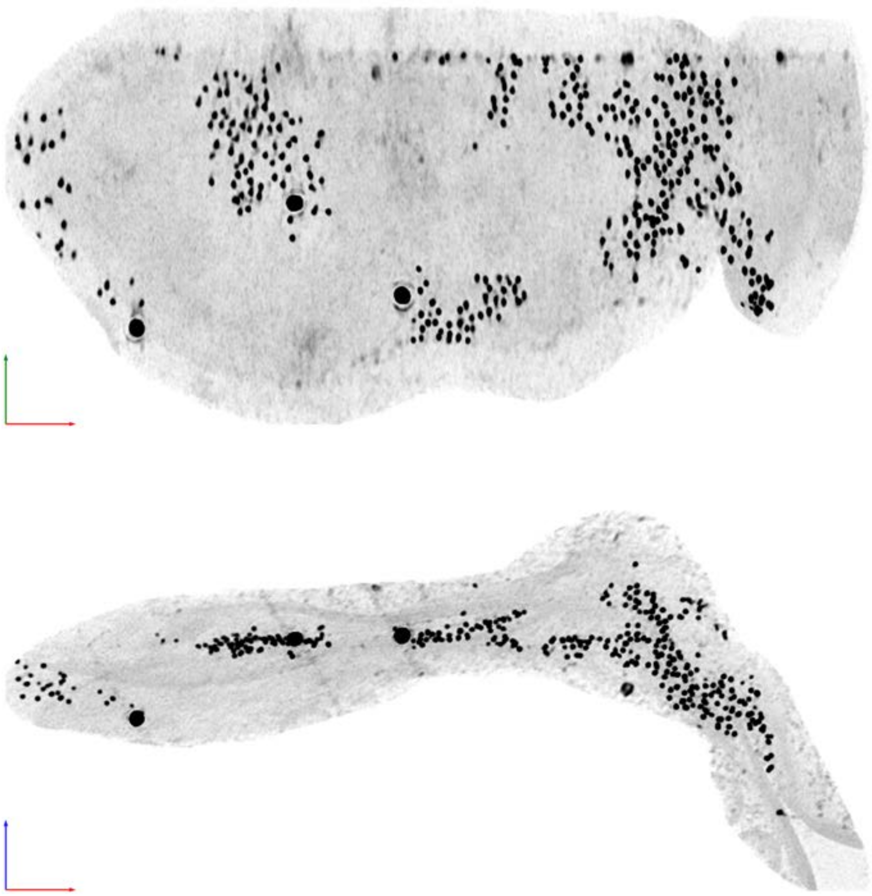

| Tomogram ID: 20260127_AMmilled50-3_Position_13_3 | SET: Dual 15F1 monomer & 5F11 dimer |
|---|---|
| Cleft ID: 1 | Type: tissue |

**Denoised Tomogram Slice**

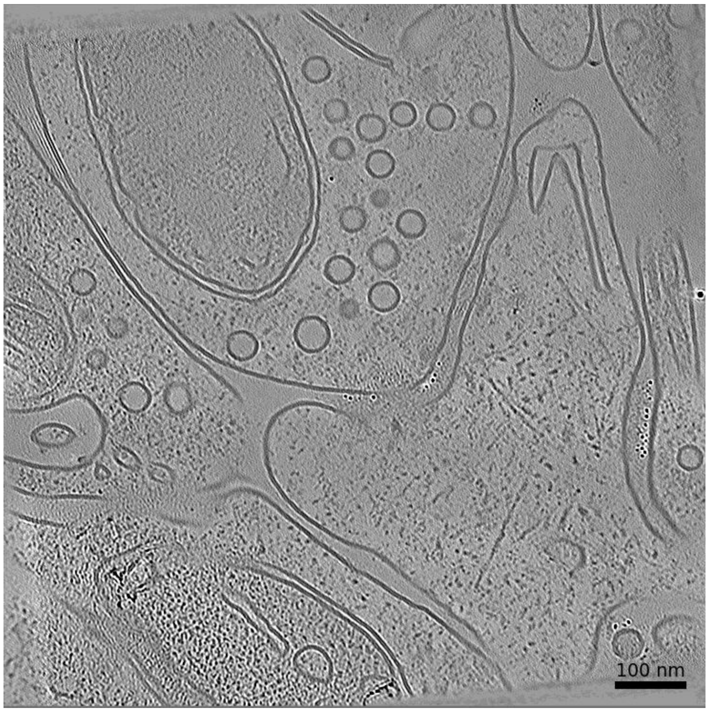


**Membrane Segmentation & Cleft Position**

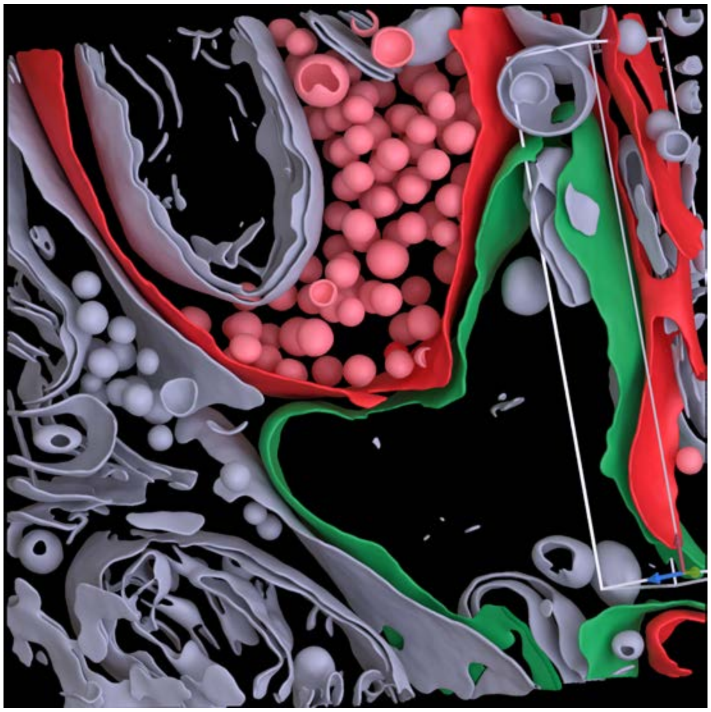

**Cleft MIP**

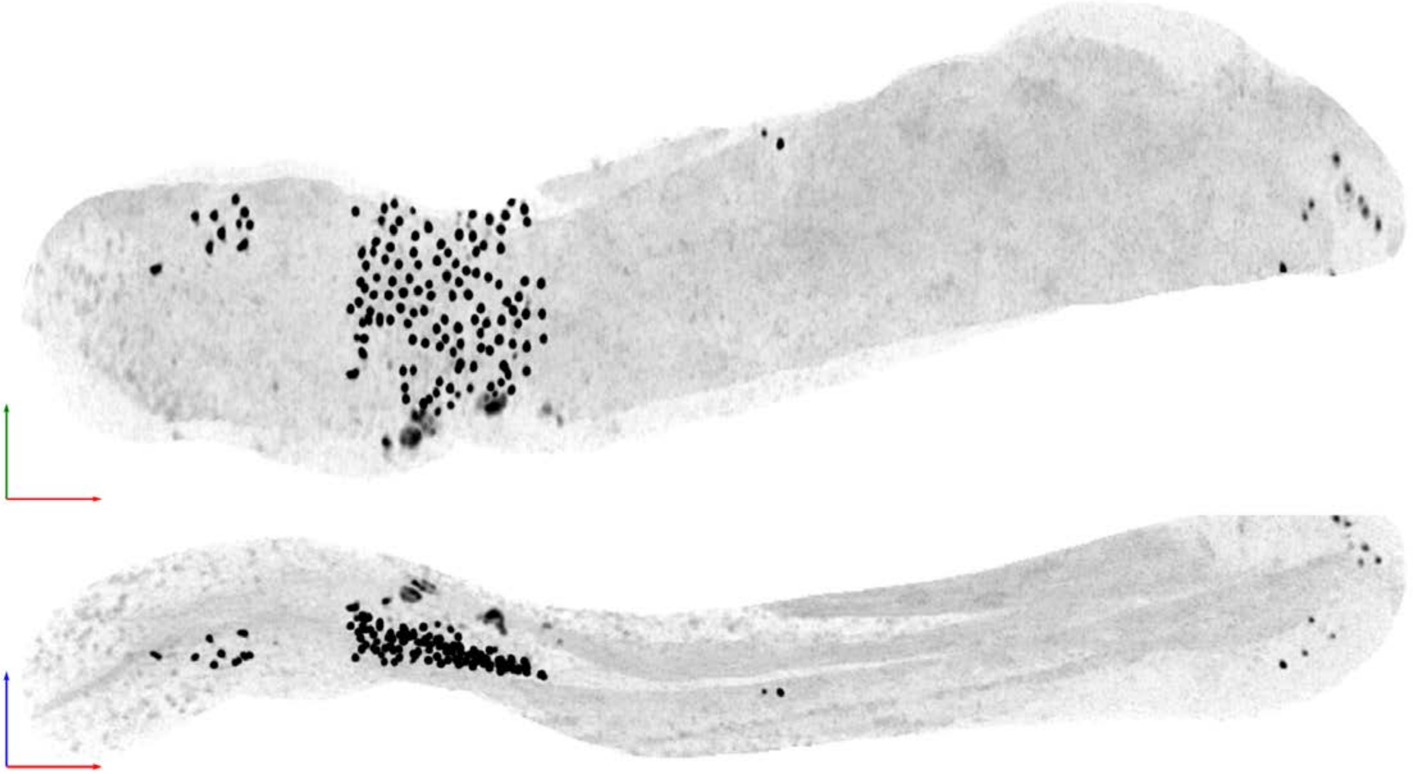

| Tomogram ID: 20260127_AMmilled50-3_Position_13_3 | SET: Dual 15F1 monomer & 5F11 dimer |
| --- | --- |
| Cleft ID: 1 | Type: tissue |

### Denoised Tomogram Slice

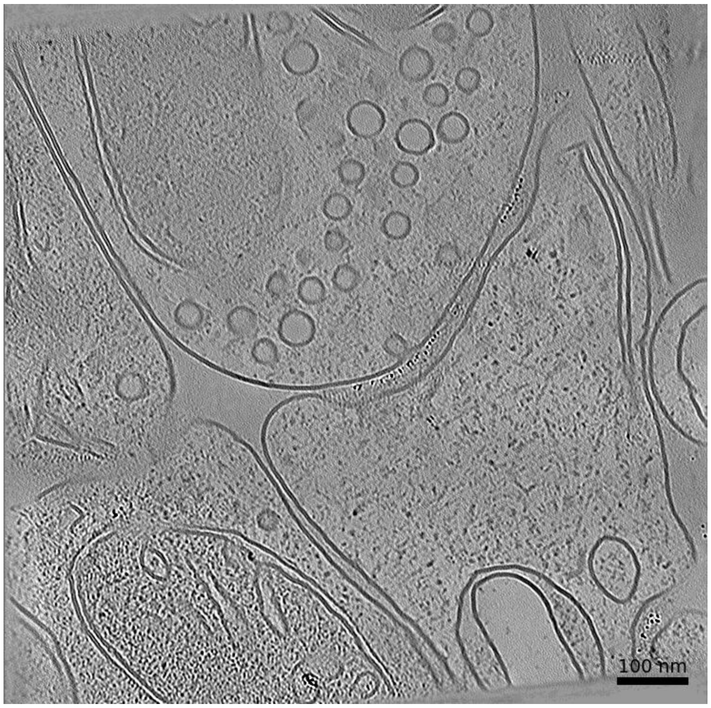


### Membrane Segmentation & Cleft Position

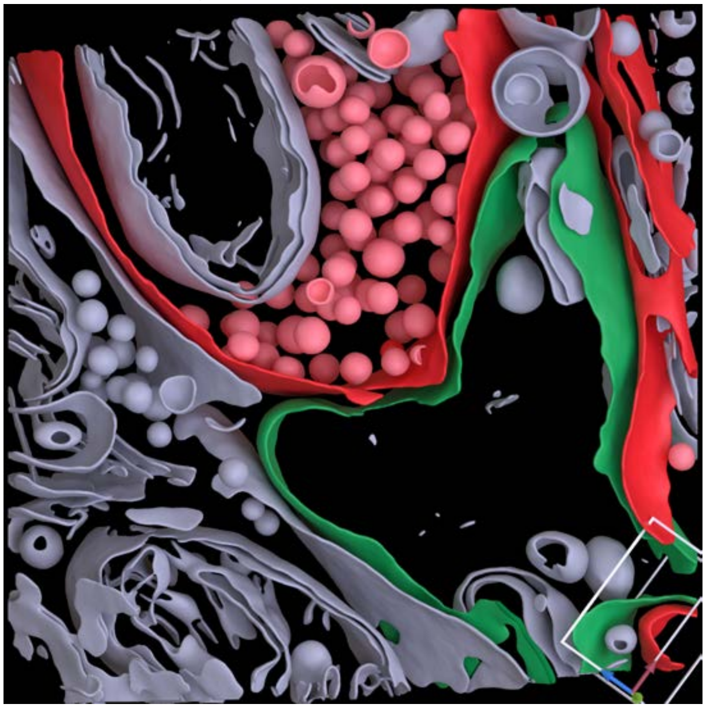

### Cleft MIP

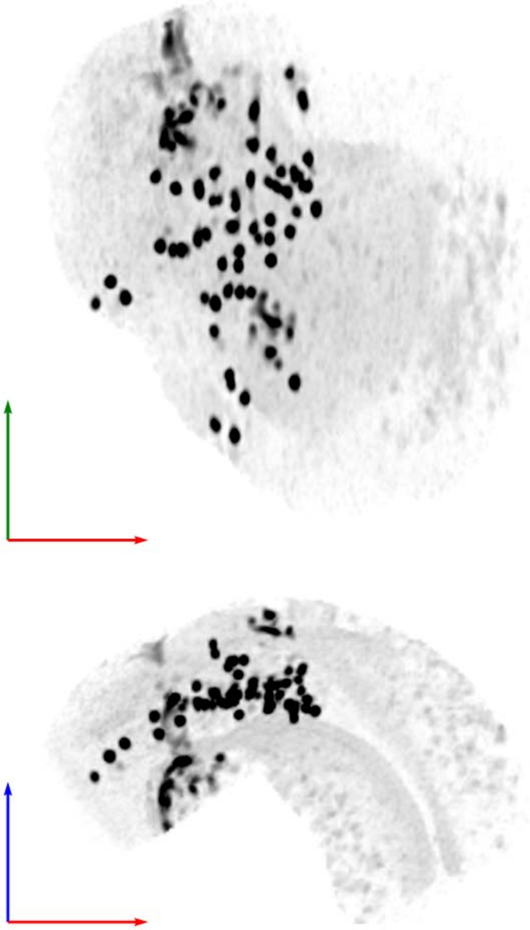

**Tomogram ID: 20260127_AMmilled50-3_Position_40** **SET: Dual 15F1 monomer & 5F11 dimer**

Cleft ID: 0 Type: tissue

### Denoised Tomogram Slice

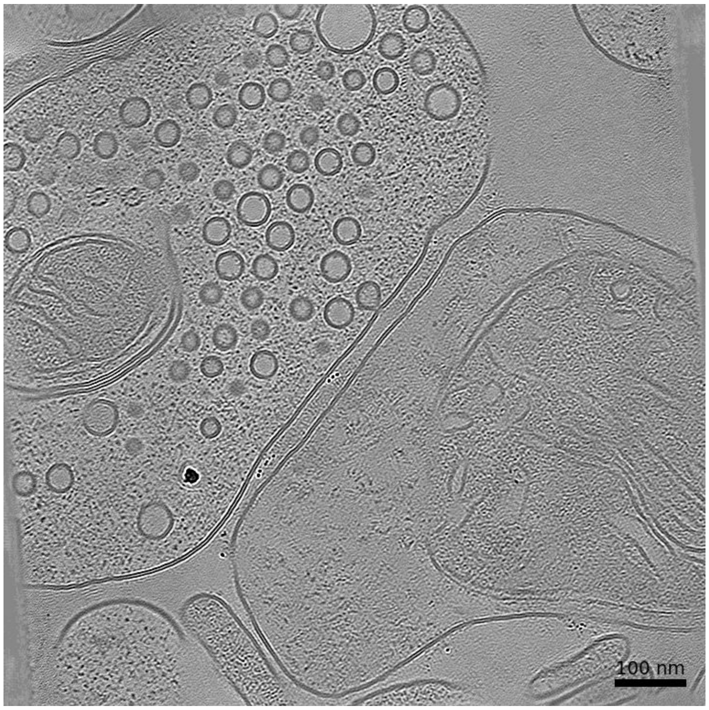


### Membrane Segmentation & Cleft Position

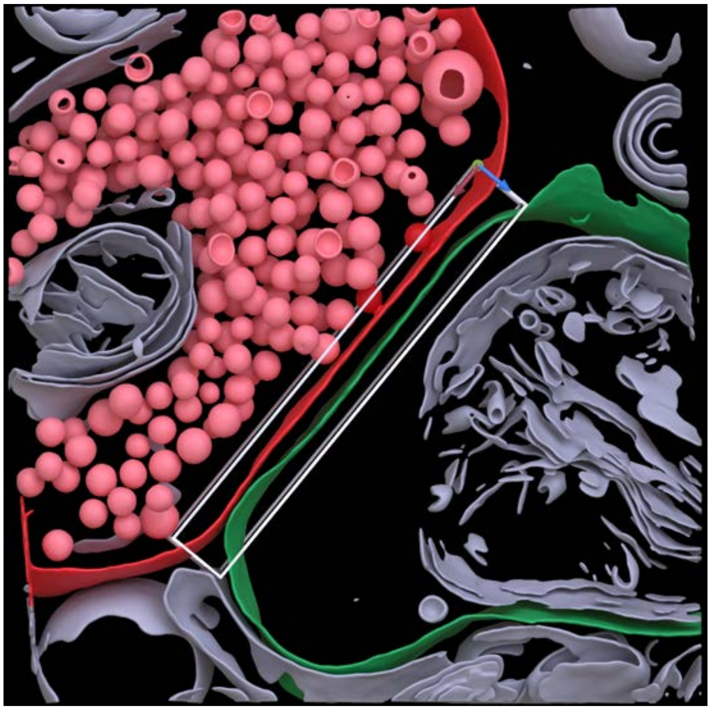

### Cleft MIP

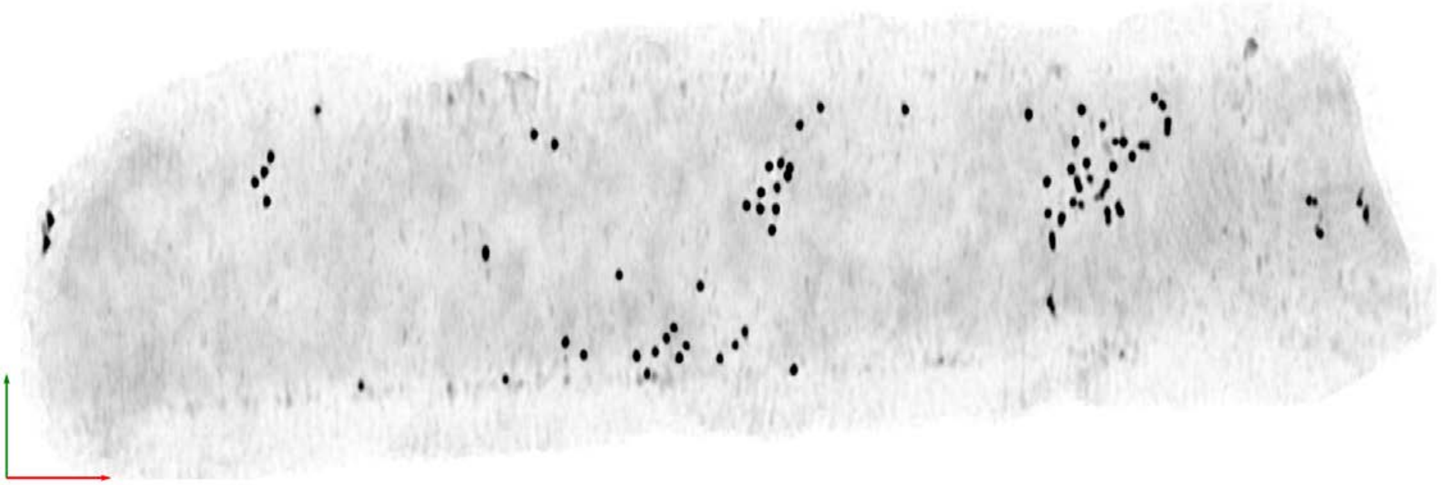

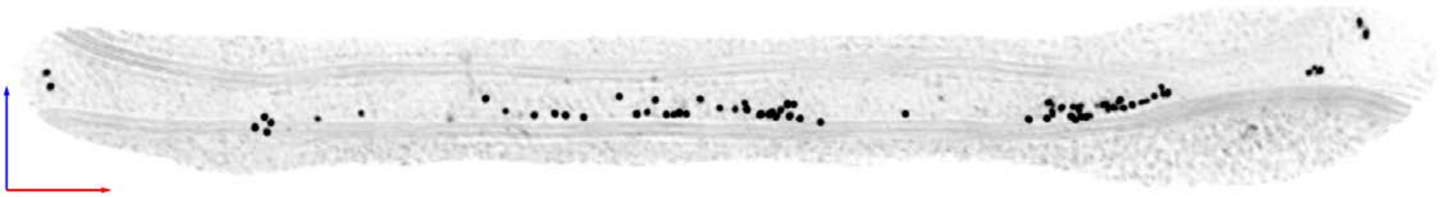

| Tomogram ID: 20260127_AMmilled50-3_Position_57_2 | SET: Dual 15F1 monomer & 5F11 dimer |
| --- | --- |
| Cleft ID: 0 | Type: semi-tissue |

**Denoised Tomogram Slice**

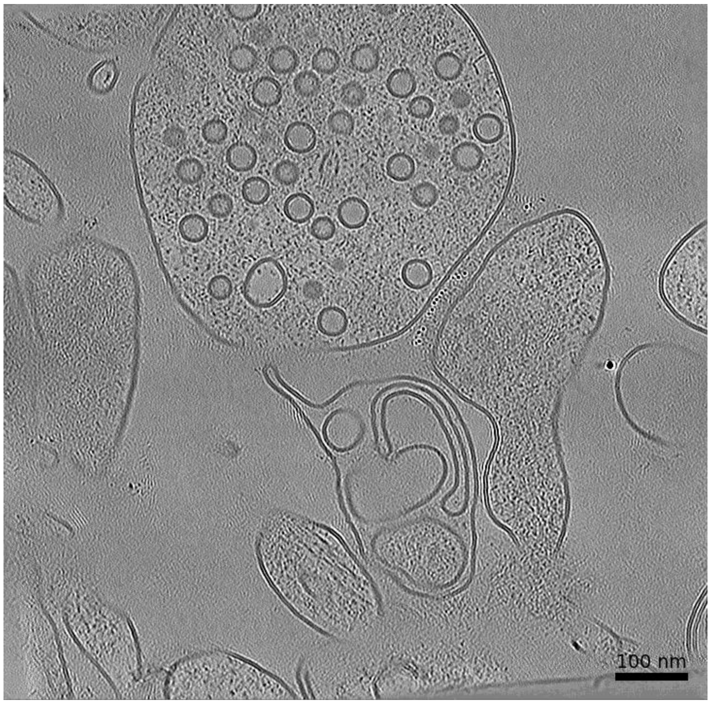


**Membrane Segmentation & Cleft Position**

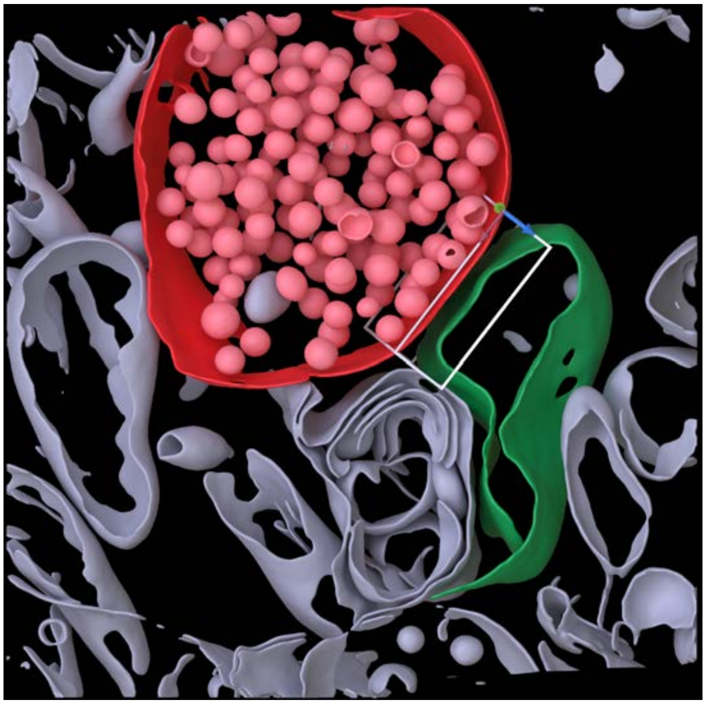

**Cleft MIP**

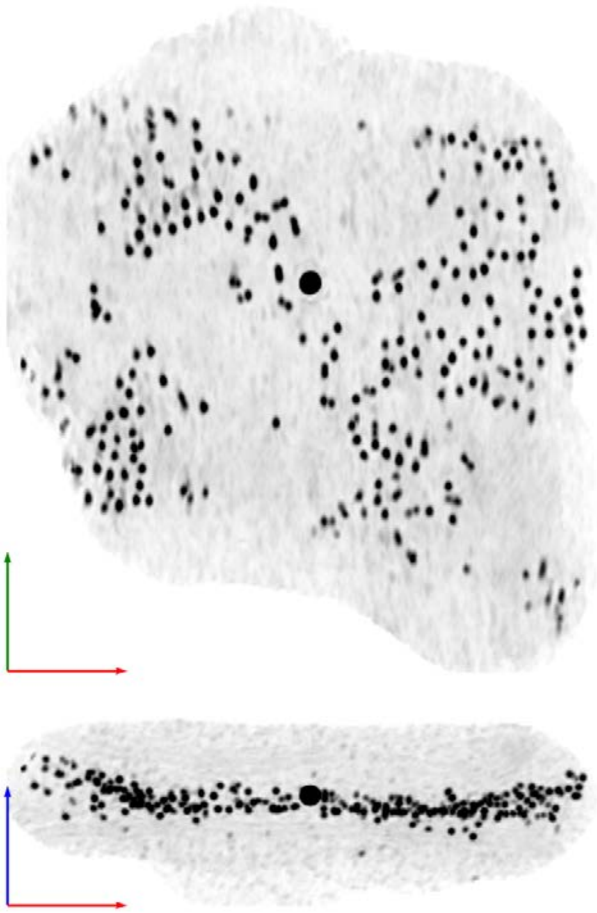

| Tomogram ID: 20260128_AMmilled50-2_Position_2_2 | SET: Dual 15F1 monomer & 5F11 dimer |
| --- | --- |
| Cleft ID: 0 | Type: tissue |

**Denoised Tomogram Slice**

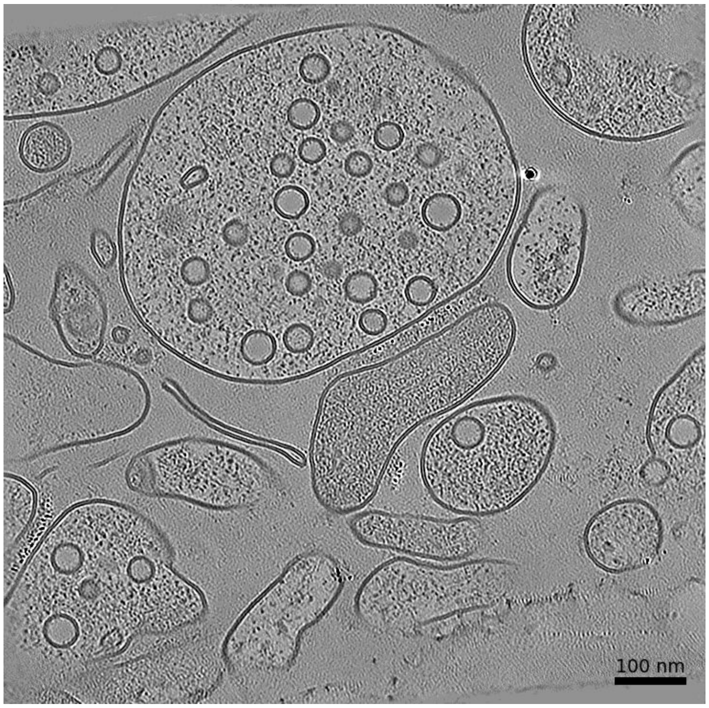


**Membrane Segmentation & Cleft Position**

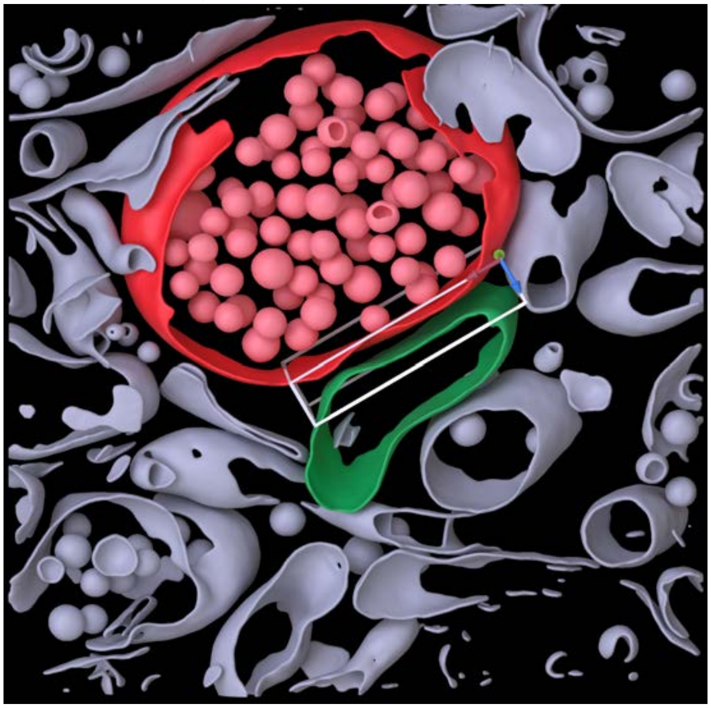

**Cleft MIP**

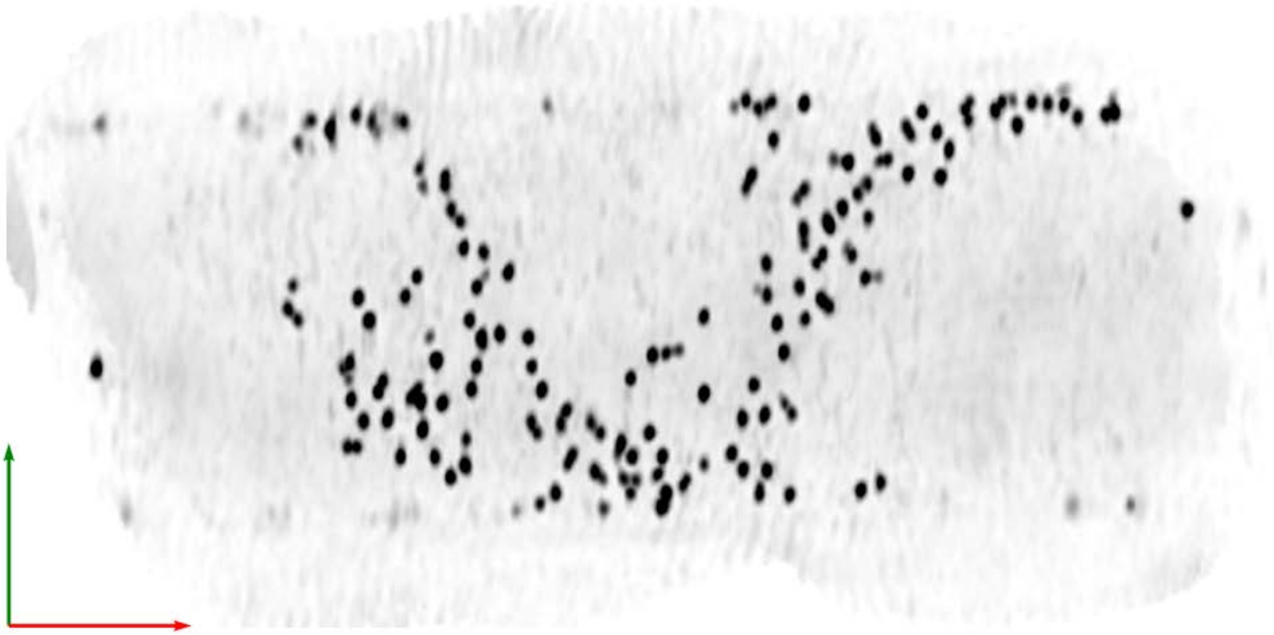

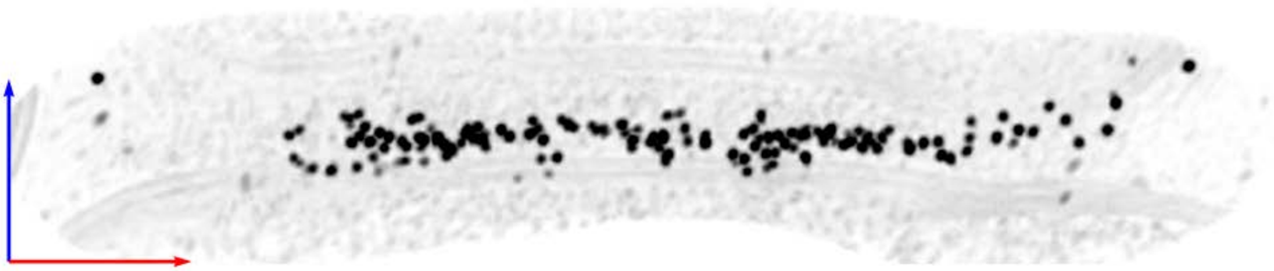

| | |
|---|---|
| **Tomogram ID: 20250227_AMmilled25-3_Position_16** | **SET: no labeling** |
| Cleft ID: 0 | Type: tissue |

**Denoised Tomogram Slice**

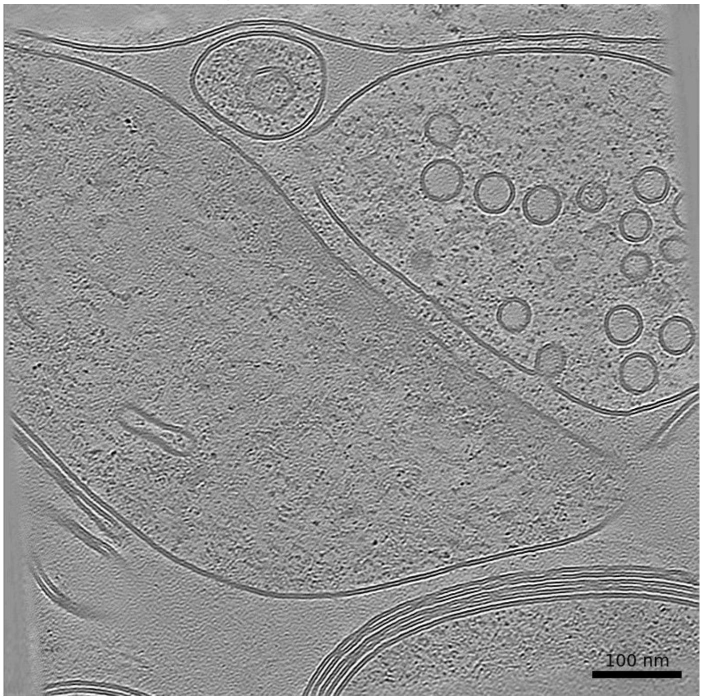


**Membrane Segmentation & Cleft Position**

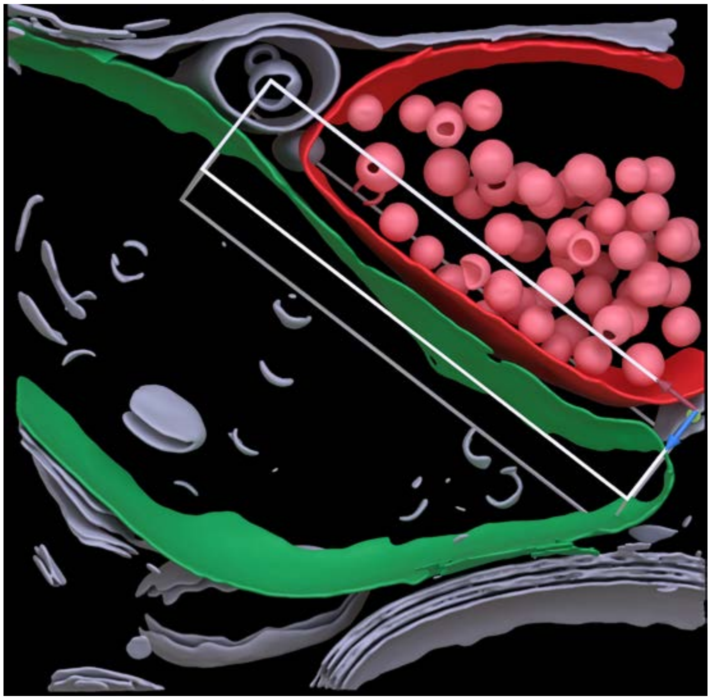

**Cleft MIP**

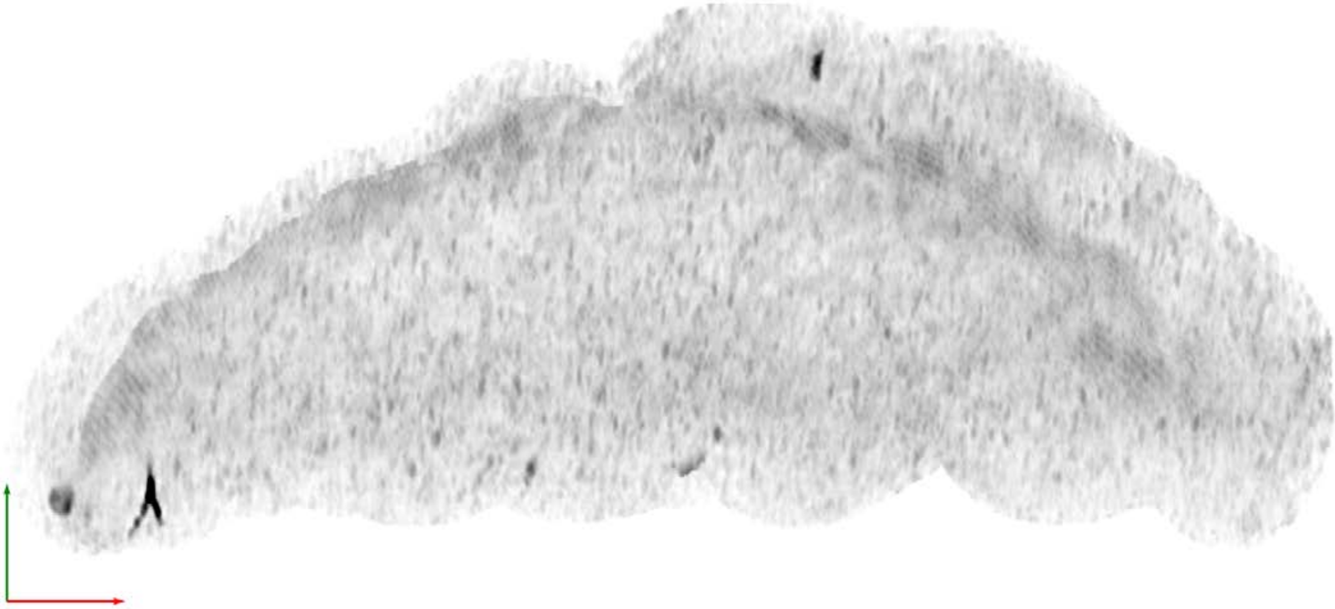

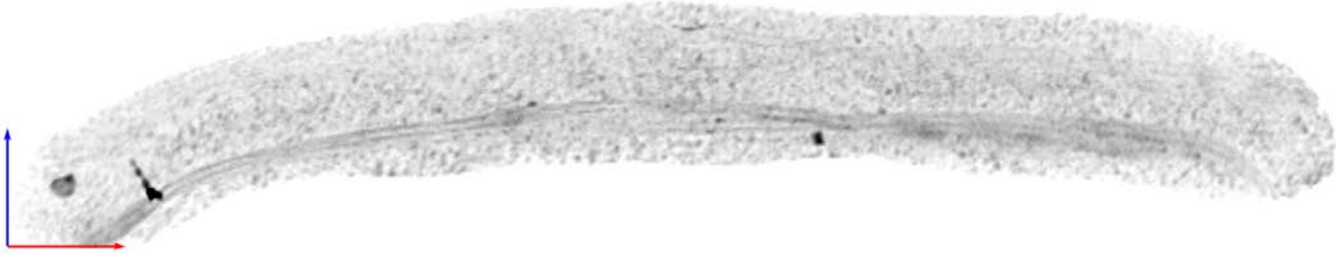

**Tomogram ID: 20250227_AMmilled25-3_Position_64** **SET: no labeling**

Cleft ID: 0 Type: tissue

### Denoised Tomogram Slice

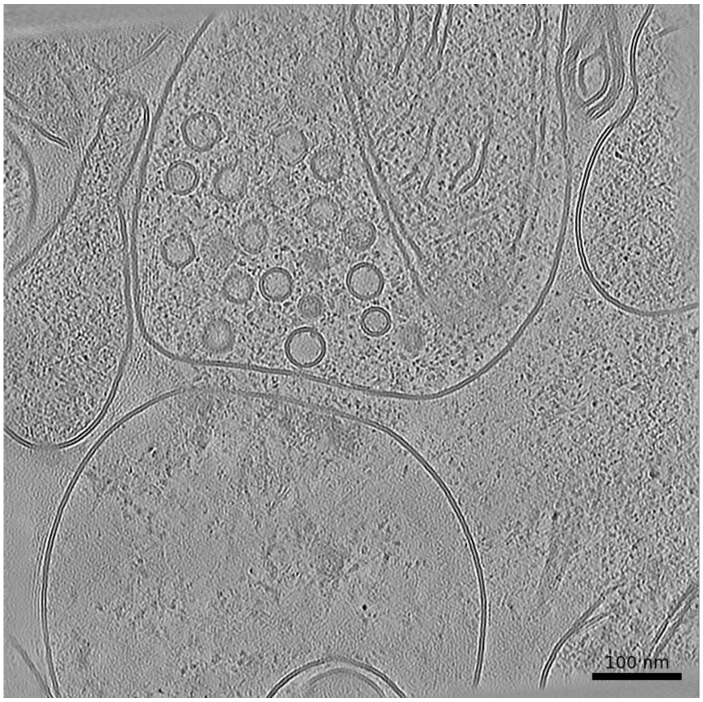


### Membrane Segmentation & Cleft Position

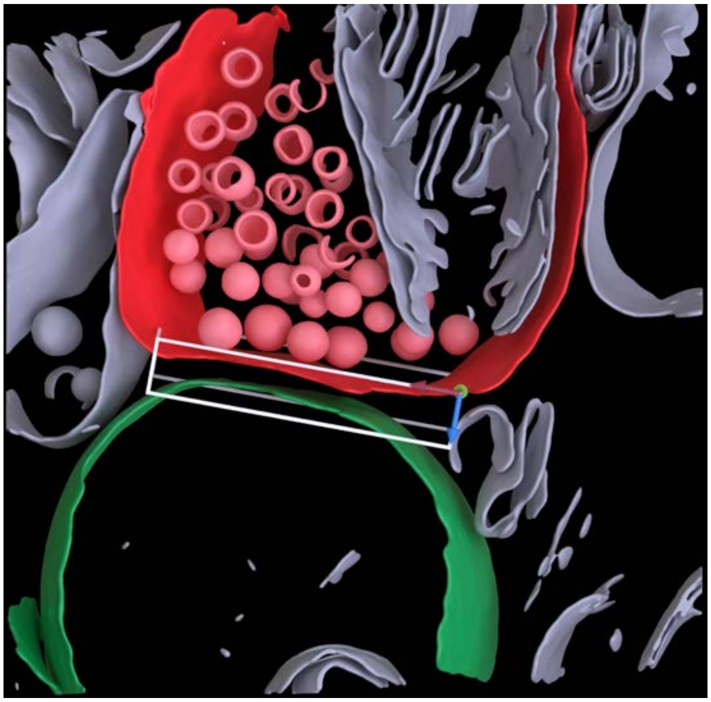

### Cleft MIP

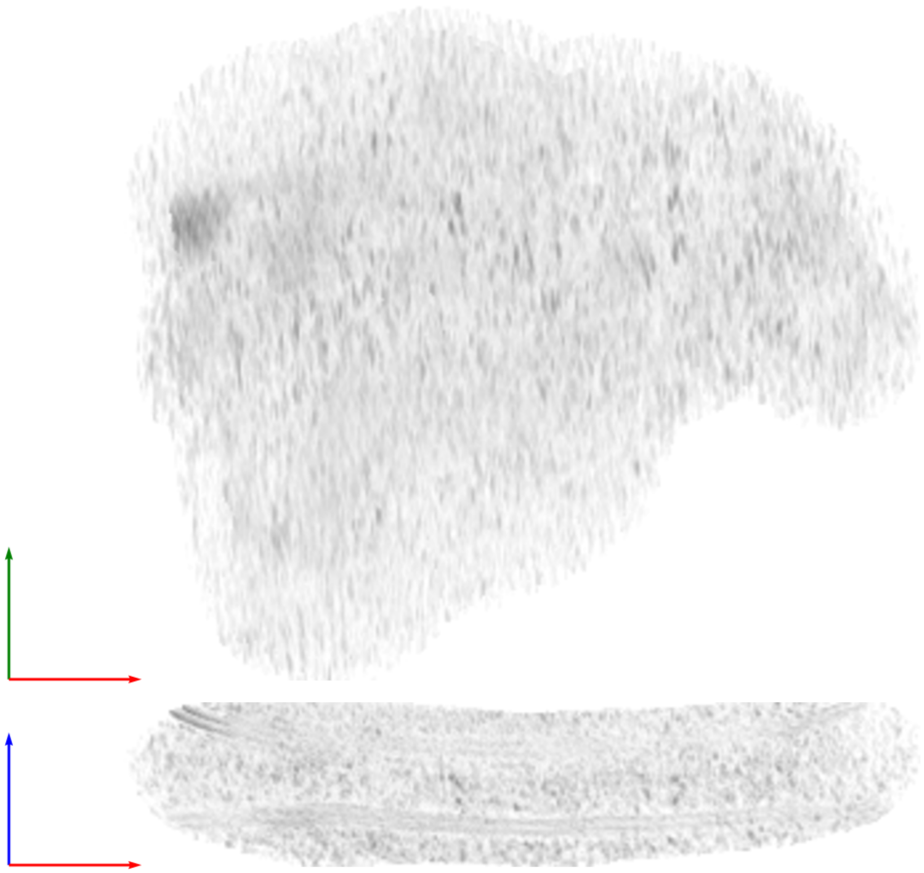

**Tomogram ID: 20250227_AMmilled25-3_Position_69_3** **SET: no labeling**

Cleft ID: 0 Type: tissue

**Denoised Tomogram Slice**

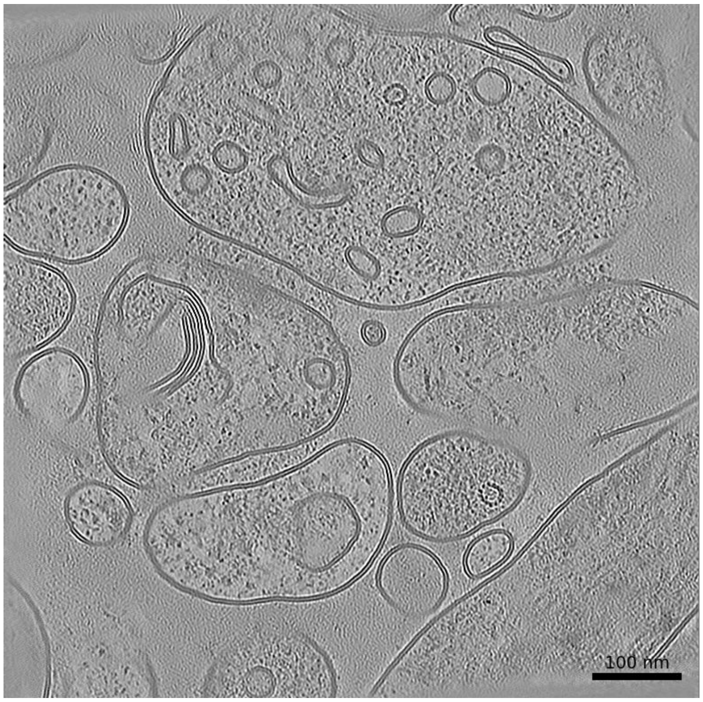


**Membrane Segmentation & Cleft Position**

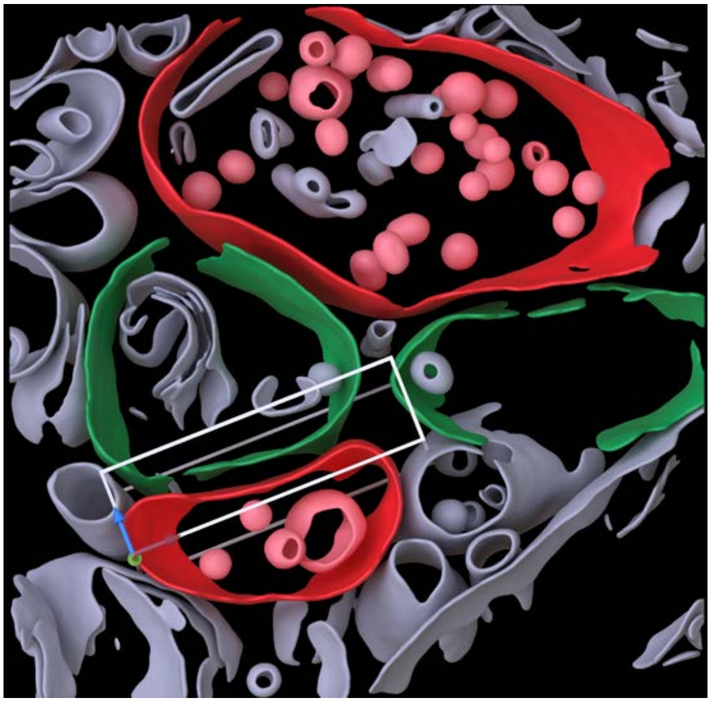

**Cleft MIP**

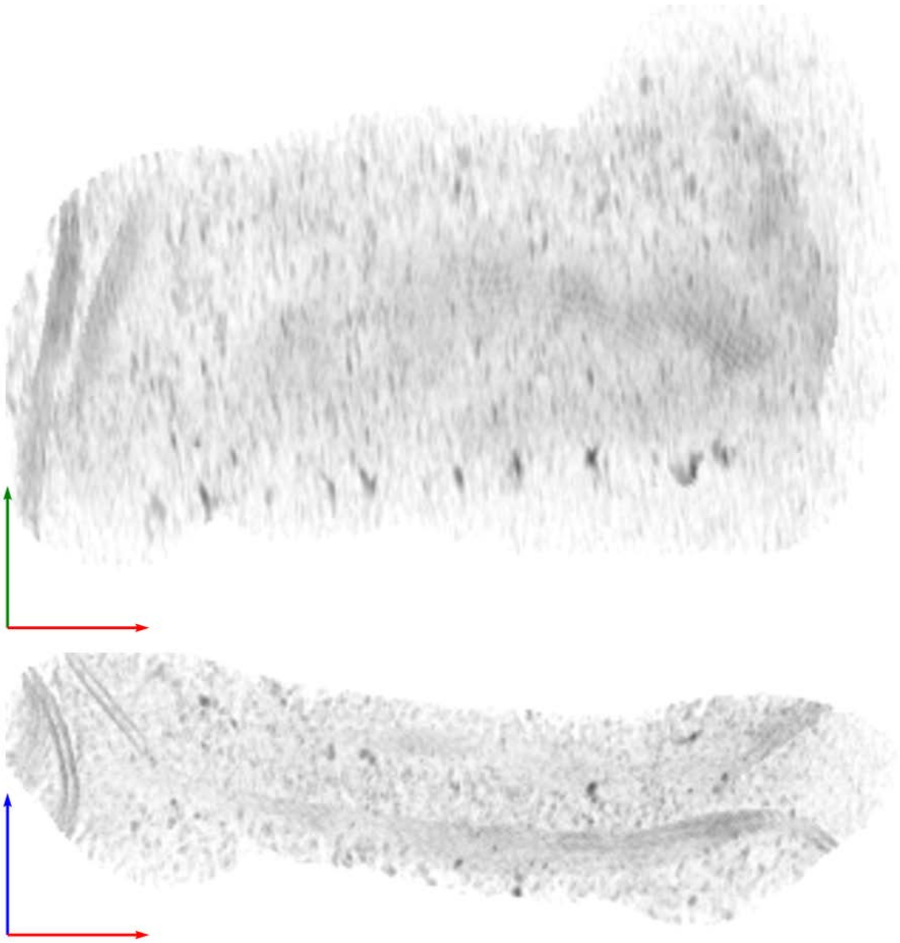

**Tomogram ID: 20250227_AMmilled25-3_Position_69_3** **SET: no labeling**

Cleft ID: 1 Type: tissue

**Denoised Tomogram Slice**

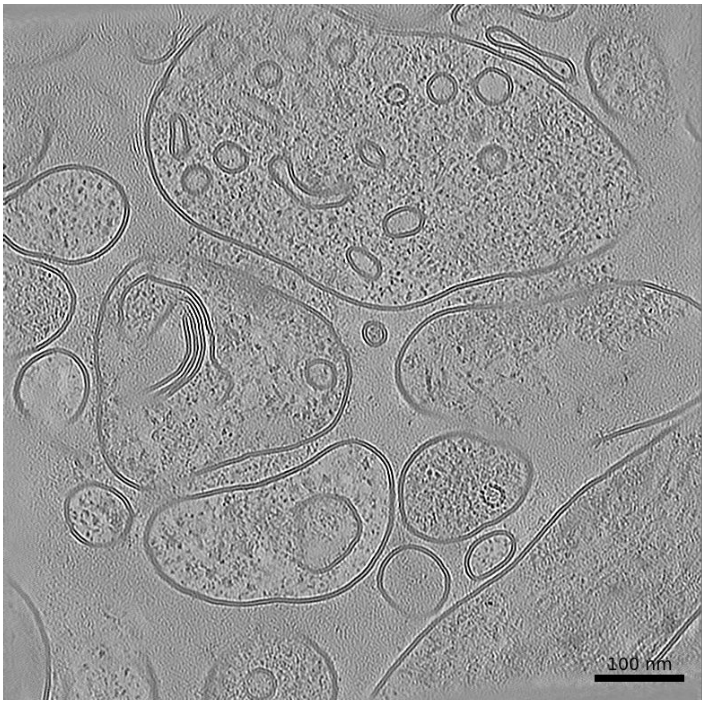


**Membrane Segmentation & Cleft Position**

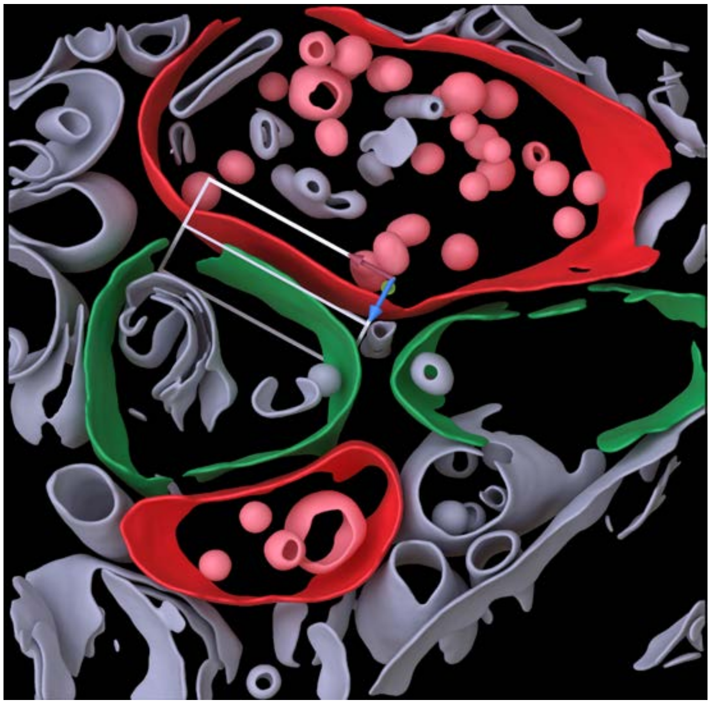

**Cleft MIP**

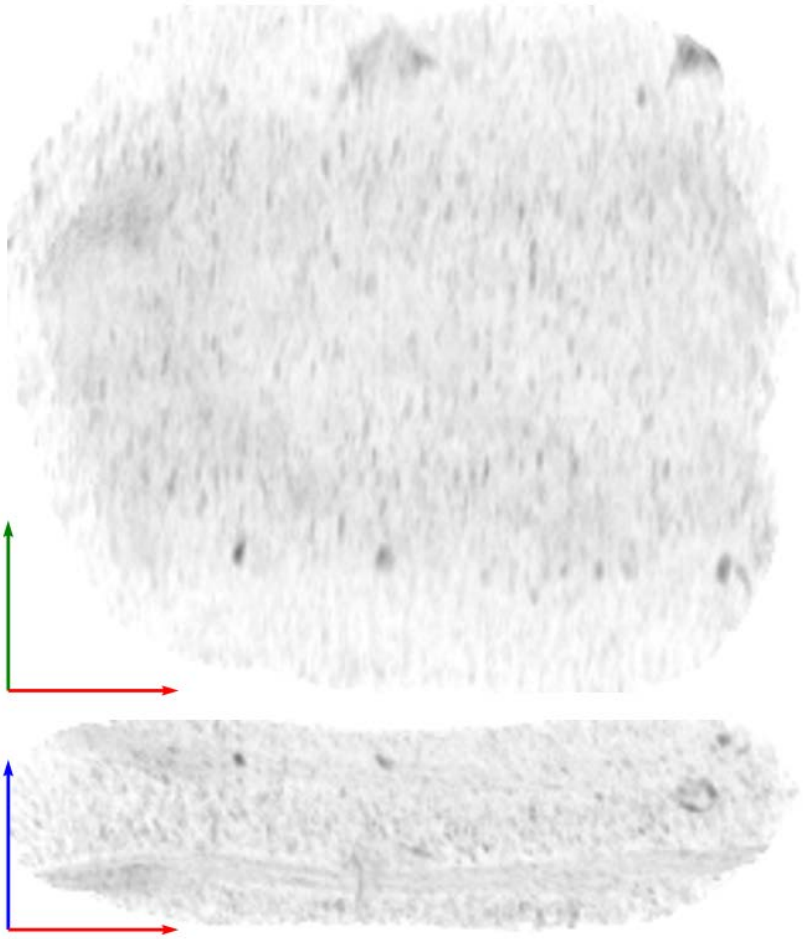

| Tomogram ID: 20250227_AMmilled25-3_Position_69_3 | SET: no labeling |
| --- | --- |
| Cleft ID: 2 | Type: tissue |

**Denoised Tomogram Slice**

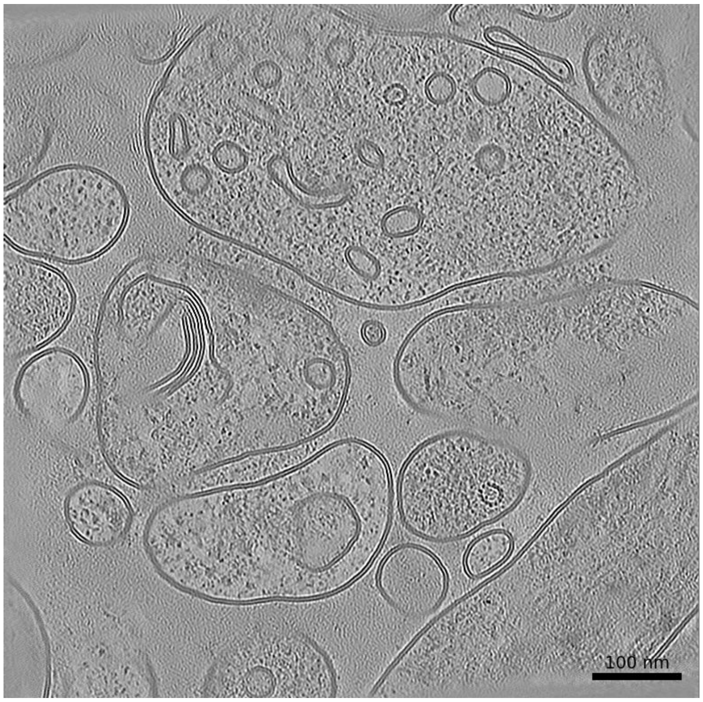


**Membrane Segmentation & Cleft Position**

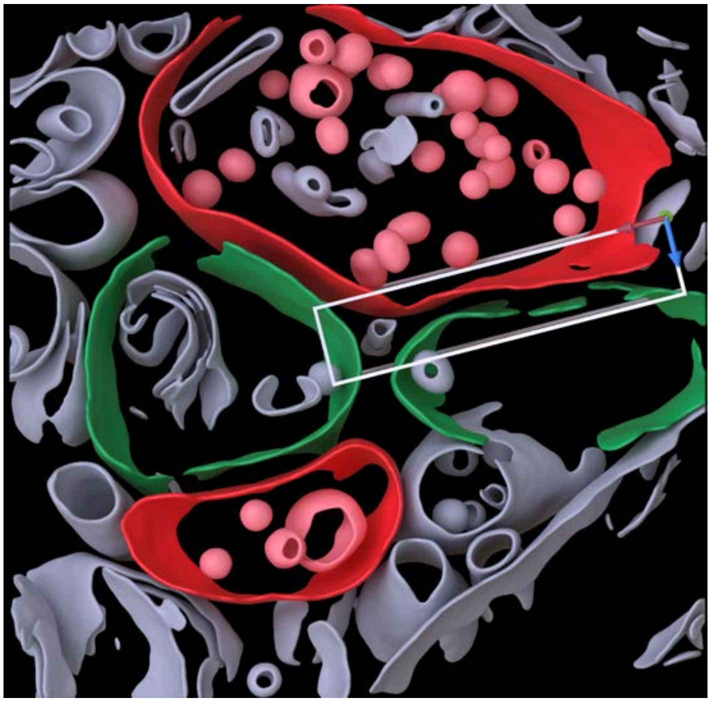

**Cleft MIP**

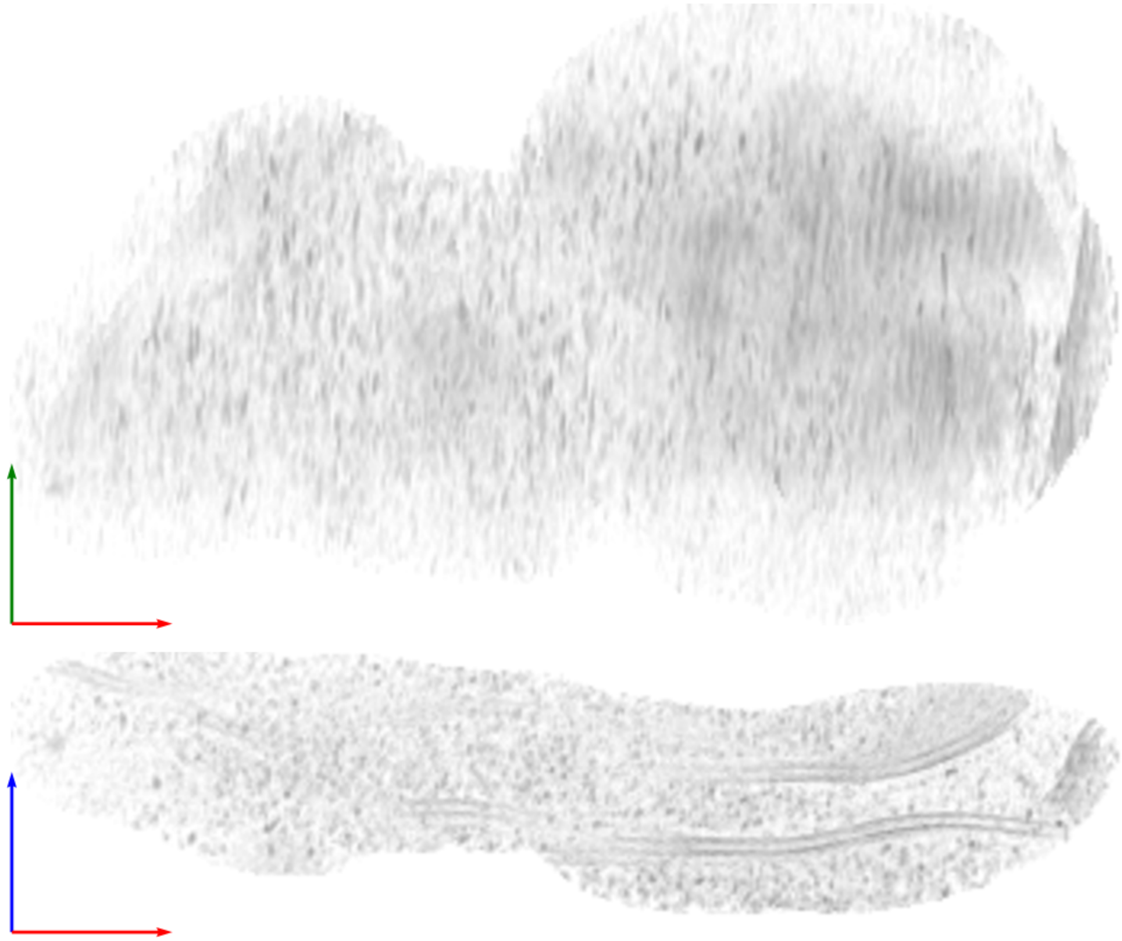

**Tomogram ID: 20250227_AMmilled25-3_Position_79** **SET: no labeling**

Cleft ID: 0 Type: tissue

**Denoised Tomogram Slice**

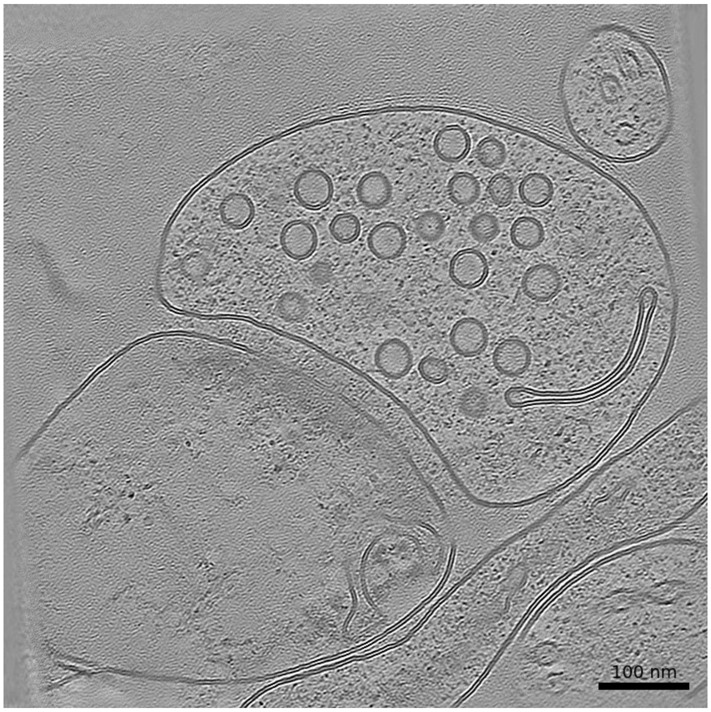


**Membrane Segmentation & Cleft Position**

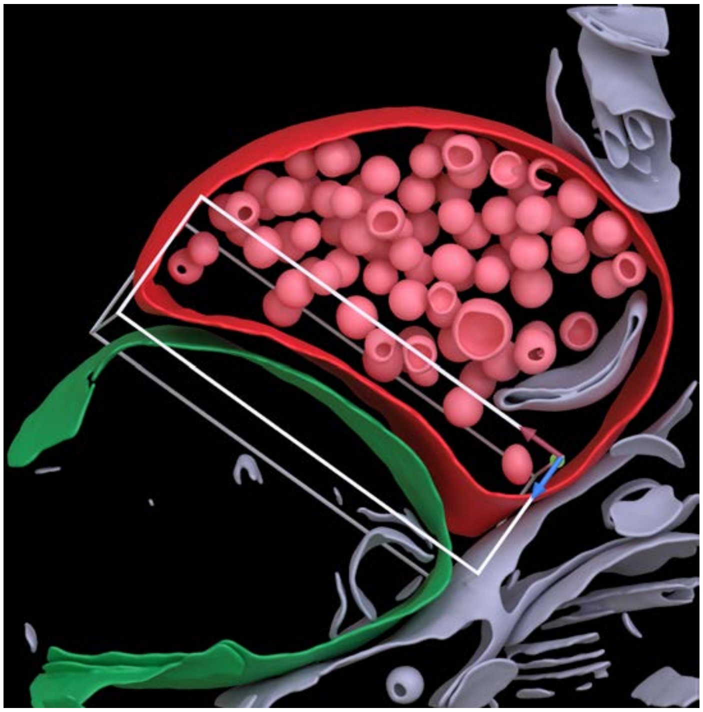

**Cleft MIP**

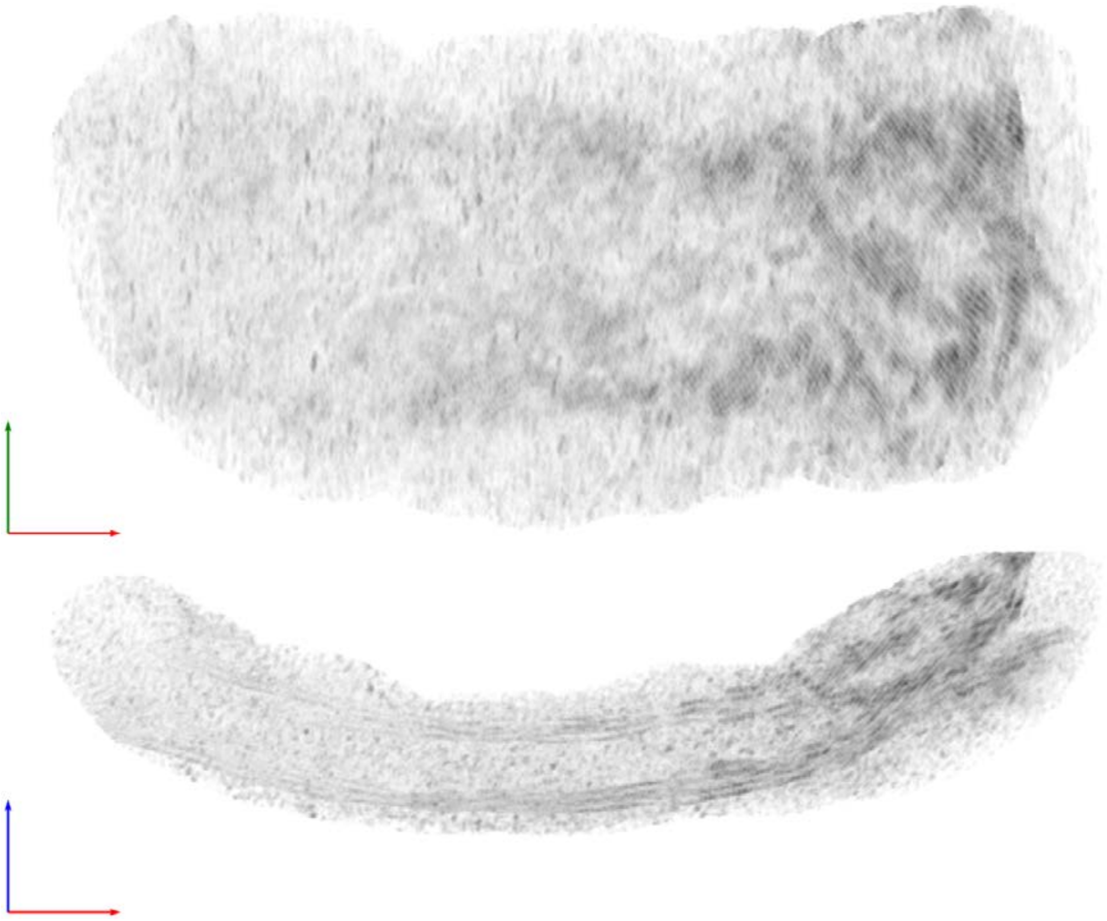

| Tomogram ID: 20250228_AMmilled25-2_Position_8_2 | SET: no labeling |
| --- | --- |
| Cleft ID: 0 | Type: synaptosome |

**Denoised Tomogram Slice**

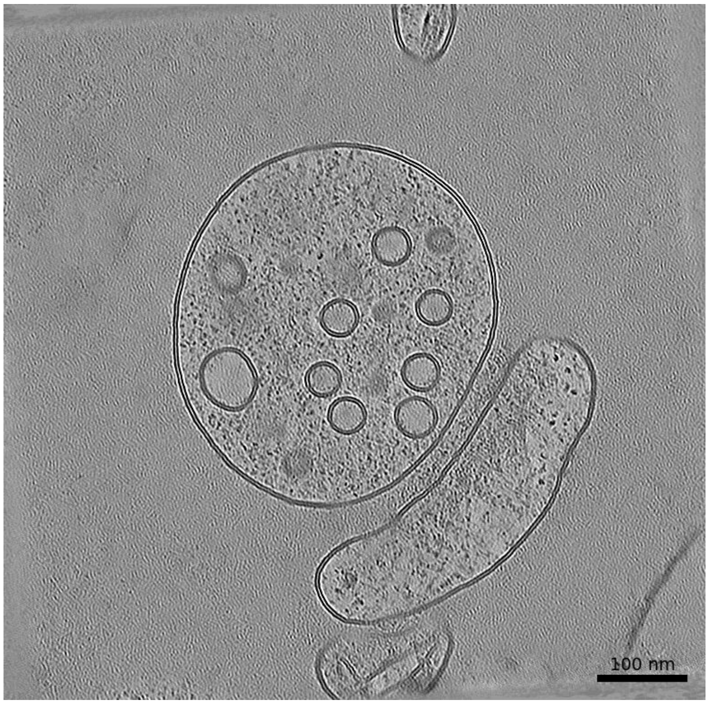


**Membrane Segmentation & Cleft Position**

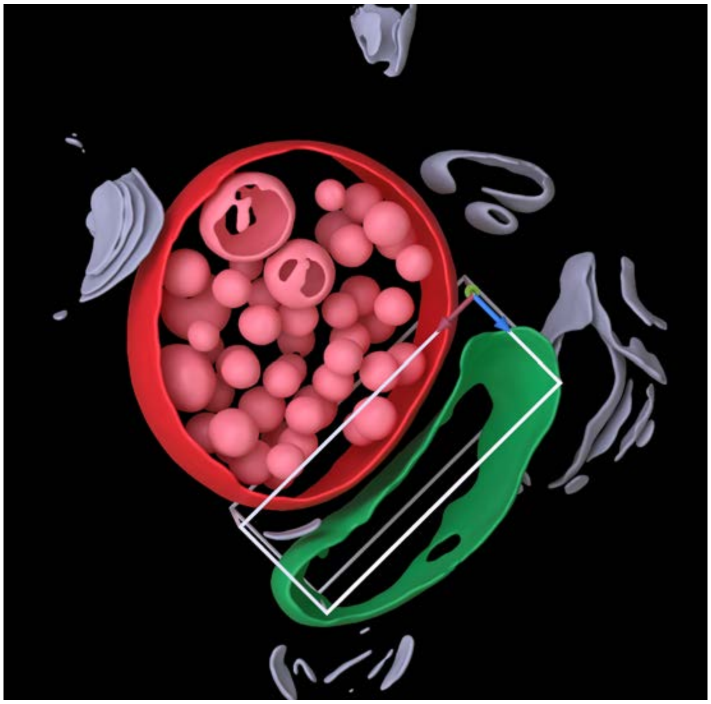

**Cleft MIP**

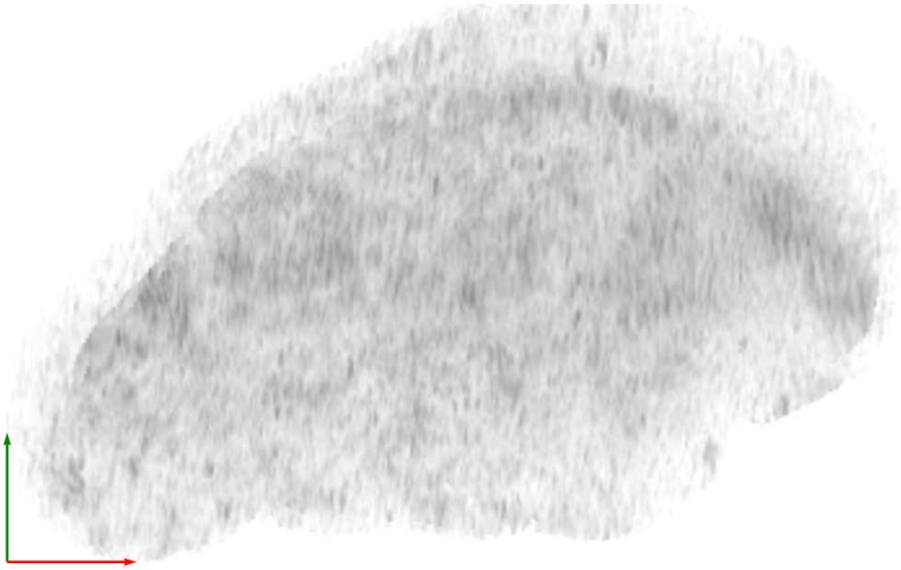

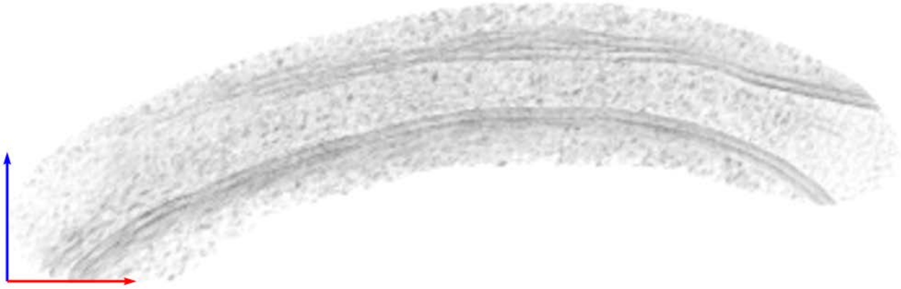
